# The evolution of DNA methylation and its relationship to sociality in insects

**DOI:** 10.1101/062455

**Authors:** Adam J. Bewick, Kevin J. Vogel, Allen J. Moore, Robert J. Schmitz

**Affiliations:** Department of Genetics, University of Georgia, Athens, GA 30602, USA; Department of Entomology, University of Georgia, Athens, GA 30602, USA

**Keywords:** DNA methylation, whole genome bisulfite sequencing, social behavior, phylogenetic comparisons, molecular evolution

## Abstract

DNA methylation contributes to gene and transcriptional regulation in eukaryotes, and therefore has been hypothesized to facilitate the evolution of flexible traits such as sociality in insects. However, DNA methylation is sparsely studied in insects. Therefore, we documented patterns of DNA methylation across a wide diversity of insects. Furthermore, we tested the hypothesis that the DNA methylation system will be associated with presence/absence of sociality among insects. We also predicted that underlying enzymatic machinery is concordant with patterns of DNA methylation. We found DNA methylation to be widespread, detected in all orders examined except Diptera (flies). Whole genome bisulfite sequencing showed that orders differed in levels of DNA methylation. Hymenopteran (ants, bees, wasps and sawflies) had some of the lowest levels, including several potential losses. Blattodea (cockroaches) show all possible patterns, including a potential loss of DNA methylation in a eusocial species whereas solitary species had the highest levels. Phylogenetically corrected comparisons revealed no evidence that supports evolutionary dependency between sociality and DNA methylation. Species with DNA methylation do not always possess the typical enzymatic machinery. We identified a gene duplication event in the maintenance DNA methyltransferase 1 (DNMT1) that is shared by some hymenopteran, and paralogs have experienced divergent, non-neutral evolution. This diversity and non-neutral evolution of underlying machinery suggests alternative DNA methylation pathways may exist. Altogether, DNA methylation is highly variable in insects and is not a universal driver of social behavior. Future, functional studies are required to advance our understanding of DNA methylation in insects.

## INTRODUCTION

DNA methylation is recognized as an important chromatin modification providing structural integrity and proper regulation of the genome for many species. In animals DNA methylation typically occurs at CG sites, established *de novo* by DNA methyltransferase 3 (DNMT3), and maintained by the maintenance methyltransferase DNMT1 (Kim et al. 2008; Goll and Bestor 2005; Cheng and Blumenthal 2008). Homologous to DNMT1 and 3 is DNMT2: a tRNA^Asp^ DNA methyltransferase, which does not contribute to the DNA methylome (Goll et al. 2006). Despite CG DNA methylation conservation across the tree of life (Feng et al. 2010; Zemach et al. 2010; Huff and Zilberman 2014), the understanding of its contribution to genome function and complex traits is limited. Although it has been proposed that heritable epigenetic variation is an important contributor to evolutionary change, direct evidence for this hypothesis is sparse. In insects, DNA methylation is implicated in behavioral plasticity and social behavior, especially eusociality in Hymenoptera (Yan et al. 2014; Yan et al. 2015). At a mechanistic level, it is difficult to test the role of DNA methylation in social behavior because the generation of mutants depleted of DNA methylation are not available. As an alternative, comparative methods have been used to infer conserved function of DNA methylation (Elango et al. 2009; Bonasio et al. 2012). Comparisons between pairs of species within the order Hymenoptera (ants, bees, wasps and sawflies) support the association between DNA methylation and presence of non-reproductive castes (Elango et al. 2009; Bonasio et al. 2012; Patalano et al. 2013). Although others have observed little association between social behavior and DNA methylation (Libbrecht et al. 2016; Standage et al. 2016). However, comparative epigenomics in insects has been taxonomically piecemeal, and examining potential influences on insect sociality beyond eusocial traits have yet to be performed.

Here we test the hypothesis that DNA methylation is associated with the evolution of sociality, more broadly defined (Wilson 1971; Costa 2006) across insects and independent of phylogenetic history. We define sociality at as is commonly done (Wilson 1971; Costa 2006) – solitary (none), communal (group living without cooperation or care), subsocial (overlapping generations and parental care), and eusocial (overlapping generations, parental care, and reproductive division of labor). Because studies of DNA methylation in insects are currently taxonomically sparse, we first define the extent that DNA methylation occurs in insects. We tested for the presence of DNA methylation using whole genome bisulfite sequencing (WGBS) and utilized previously published data to generate a dataset comprising of 41 species of insects from six orders, including three orders where eusociality has evolved. To expand our taxonomic sampling and cover insect orders and species where sequencing has not yet been done, we documented the presence or absence of DNA methylation using *C_p_G_o/E_* in 123 species of insects from 11 orders. The measure *C_p_G_o/E_* relies on natural, spontaneous demanination of methylated cytosines to thymines, and robustly recovers presence/absence of DNA methylation (see Materials and Methods). We then used both continuous (WGBS) and discrete (*C_p_G_o/E_*) data in explicit phylogenetic comparative tests that take into account evolutionary history and non-independence of species. We tested our hypothesis at two broad levels of sociality; the evolution of any level of sociality (from solitary to eusocial) and the evolution of division of labor associated with eusociality. Finally, we examined the presence, distribution and evolution of DNA methyltransferases across the diversity of insects. We found widespread presence but diverse patterns of DNA methylation across the insect tree of life. Contrary to a prediction of a role for DNA methylation facilitating social evolution, we found no association between sociality and DNA methylation in insects. Instead we suggest that the presence of DNA methylation in a variety of insects suggests a much broader function of this chromatin modification. This is supported by the diverse patterns of presence/ absence of DNA methyltransferases, including duplication of DNMT1 in some groups and lack of DNMT3 in others. Functional studies are now needed to elucidate the role of DNA methylation in insects, and our work will facilitate identifying species suitable for such studies.

## RESULTS AND DISCUSSION

### DNA methylation levels are highly variable across insects

Estimates of DNA methylation from WGBS from 41 species from six orders (Blattodea [n=9], Hemiptera [n=5], Hymenoptera [n=12], Coleoptera [n=3], Lepidoptera [n=4], and Diptera [n=8]) revealed diverse levels of DNA methylation. DNA methylation was found in all insect orders except Diptera and in social and solitary species (Fig. 1; Table S1). DNA methylation levels within coding regions are higher than genome-wide levels for all species, which suggests a conserved pattern of DNA methylation across insects (Fig. 1; Table S1). Higher levels of DNA methylation within coding regions compared to genome-wide might partially reflect highly conserved protein coding genes in insects having the highest levels of DNA methylation (Glastad et al. 2014). DNA methylation levels within commonly studied hymenopteran are some of the lowest compared to other species sampled, and three species (*Aphidius ervi*, *Microplitis demolitor* and *Microplitis mediator*) are identified as having completely absent or extremely reduced levels of DNA methylation (Fig. 1; Table S1). This loss or extreme reduction in DNA methylation is similar to what is observed in the eusocial species *Polistes dominula* (Hymenoptera) (Standage et al. 2016). Levels of DNA methylation genome-wide and within coding regions were overall higher in the order Blattodea (Fig. 1). Genome-wide and coding levels of DNA methylation were highest in the solitary cockroaches in the genus *Blattella*, *B. asahinai* and *B. germanica*, respectively. Furthermore, drastic differences in levels of DNA methylation were observed between the highly eusocial species *Reticulitermes flavipes* and *R. virginicus* within Blattodea (Fig. 1). Conversely, similar levels of DNA methylation were observed between closely related *Pemphigus spp*. of aphids (Hemiptera) with differing social behaviors (Fig. 1). Additionally, the solitary species *Acyrthosiphon pisum* had the highest levels of DNA methylation within in the Hemiptera. Overall, DNA methylation levels were variable in insects, with some species’ levels reaching those observed in plants (Feng et al. 2010; Zemach et al. 2010; Niederhuth and Bewick et al. 2016; Takuno et al. 2016), where its contribution to complex traits still remains largely unknown.

**Figure 1.**
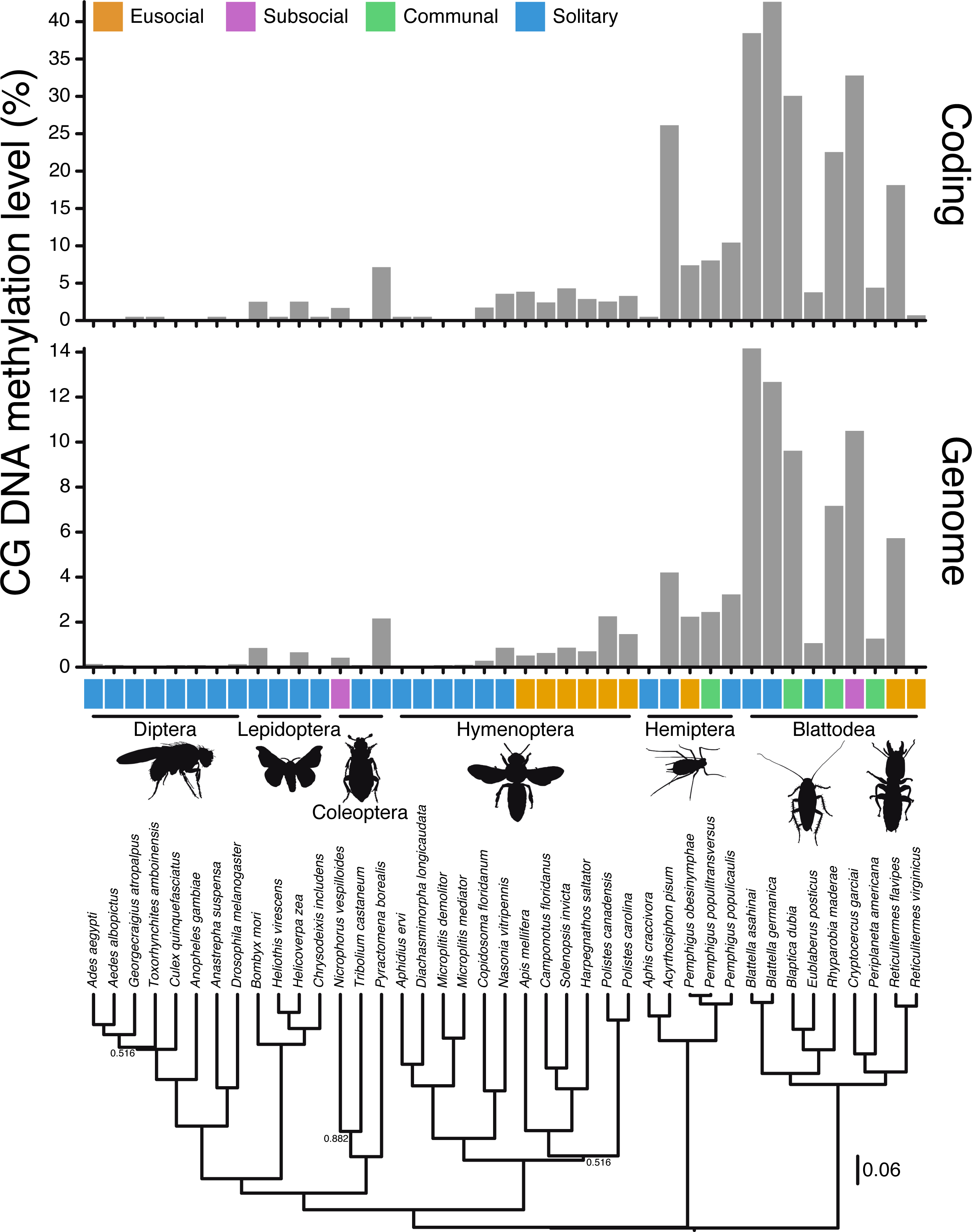
WGBS reveals extensive variation of DNA methylation in insects. (A) Genomic levels of DNA methylation in insects ranges from zero (all Diptera examined) to ∼14% (*Blattella asahinai*). Higher ranges are observed for coding regions; zero to ∼42% (*Blattella germanica*). Overall, levels are highest in Blattodea, and do not always associate with social species. (B) A species tree constructed from nuclear and mitochondrial loci, which was used in Phylogenetic Generalized Least Squares (PGLS) analysis. Results from this analysis revealed that there is no correlation between social behavior and DNA methylation (Table 1). Values at nodes are posterior probabilities <0.95; all blank nodes have ≥0.95 posterior probability.

### DNA methylation is present across the insect tree of life

DNA methylation, signified by a bimodal distribution of *C_p_G_o/E_* values, is a robust estimator for the presence/absence of DNA methylation in insect genomes. When compared to estimates of DNA methylation from WGBS, *C_p_G_o/E_* accurately estimates presence/absence of DNA methylation in 18/18 (100%) species (see Materials and Methods, and Table S1). Therefore, to taxonomically expand on insect orders and species, the distribution of *C_p_G_o/E_* of 123 insect species from 11 orders were investigated for signatures of DNA methylation. Similarly to our WGBS data, DNA methylation was identified in insect species from all orders except Diptera (Fig. 2A and B). Species belonging to the order Diptera comprised ∼46.3% (57/123) of the total number of species investigated, which was the largest number of species sampled from one order. The absence of bimodal distribution from members of both the Nematocera and Brachycera sub-orders suggests that DNA methylation was lost early on in dipteran evolution. The evolution of DNA methylation is deeply rooted in insects, going as far back as the Ordovician and the divergence of Diplura (∼489.84-436.68 MYA (Misof et al. 2014)), but is most likely older.

**Figure 2.**
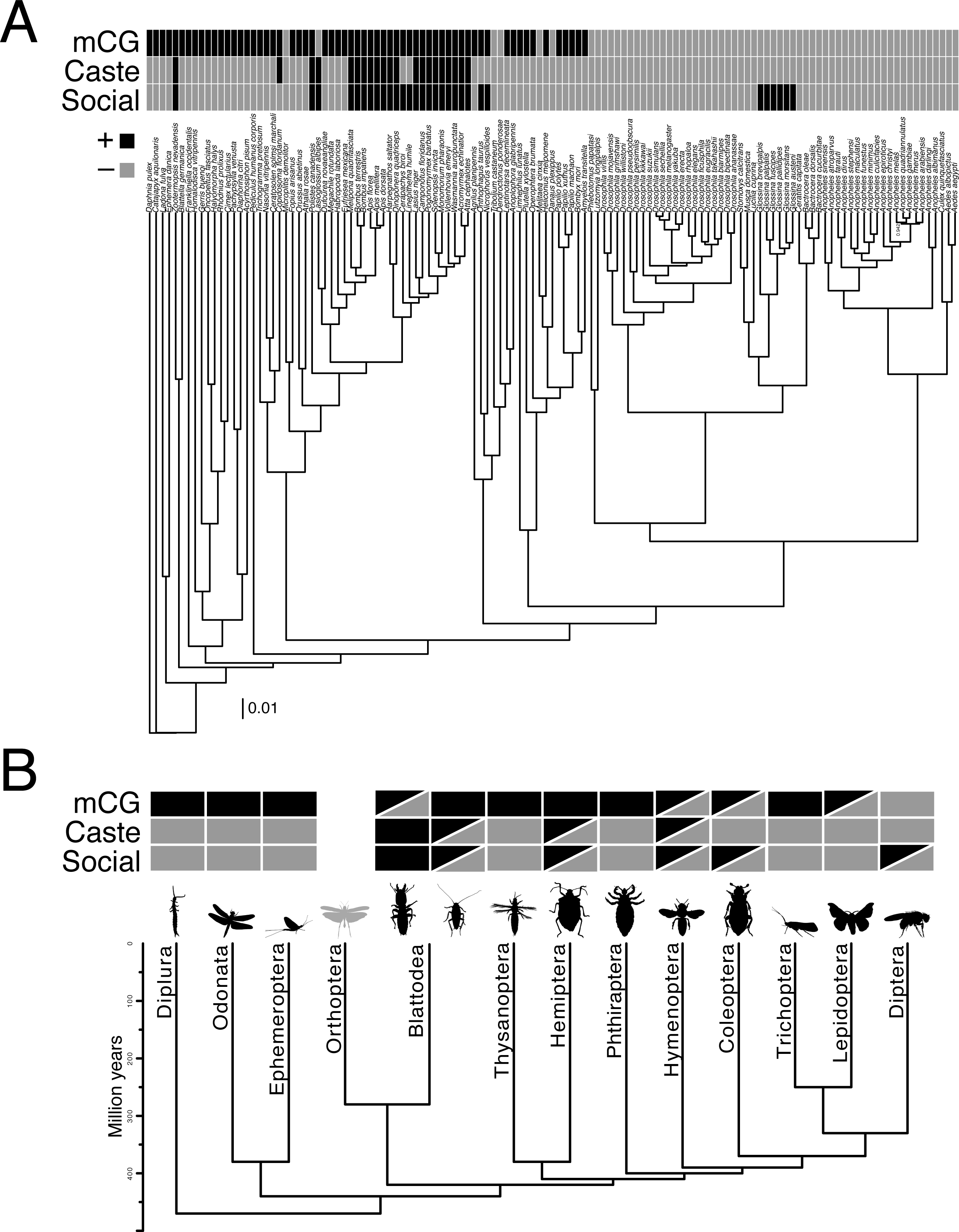
DNA methylation in insects is not always associated with social behavior. (A) Relationships of 123 insect species, and the outgroups *Catajapyx aquilonaris* (Dipluran) and *Daphnia pulex* (Crustacea) investigated with sociality, division of labor and DNA methylation scored as a binary (presence/absence) trait. The tree was constructed from 58 nuclear protein coding loci, and was used in Pagel’s test for evolutionary dependence. All nodes except one had posterior probability of 1.00, which is indicated on the phylogeny. (B) A chronogram of insect order relationships with sociality, division of labor and DNA methylation scored as binary (presence/absence) trait. The chronogram was modified from Misof et al. 2014. For (A) and (B) traits are represented as shaded boxes above each species or order. Half-filled boxes indicate the trait is variable within the corresponding order.

DNA methylation was found in insects exhibiting the complete range of social behavior based on modality of *C_p_G_o/E_* (Table S1). This includes insects classified as solitary, communal (group living), subsocial (parental care), and eusocial (cooperative brood care, overlapping generations and reproductive castes – references (Wilson 1971; Costa 2006)). For example, the solitary thrip *Frankliniella occidentalis* had an extremely pronounced bimodal *C_p_G_o/E_* distribution compared to the eusocial honey bee *Apis mellifera* (Fig. S1). The order Hymenoptera contains species that are solitary or eusocial; however, evidence for DNA methylation based on *C_p_G_o/E_* did not always co-occur with eusociality, and *vice versa* (Table S1). Moreover, all species examined within the order Lepidoptera are solitary, yet 8/10 showed evidence for DNA methylation based on the distribution of *C_p_G_o/E_*, respectively (Table S1). Similarly, 3/4 solitary coleopteran species were expected to have DNA methylation based on bimodality of *C_p_G_o/E_*. Interestingly, the six Diptera in the genus *Glossina* have sub-social behavior in the form of parental care (e.g., Table S1, the tsetse fly, *Glossina morsitans* morsitans (International Glossina Genome Initiative et al. 2014)), yet do not have DNA methylation based on the distribution of *C_p_G_o/E_*. This is contrary to the burying beetle *Nicrophorus vespilloides*, which has DNA methylation based on WGBS and *C_p_G_o/E_* (Cunningham et al. 2015), and also provides parental care through provisioning developing offspring (Walling et al. 2008) (Table S1). Overall, based on modality of *C_p_G_o/E_* DNA methylation is found in all insect orders examined except Diptera.

### Sociality and DNA methylation are not evolutionary dependent

DNA methylation has been proposed to control many aspects of sociality including behavior expressed in social interactions, caste determination, and learning and memory (reviewed in (Li-Byarlay and Hongmei 2016)). Other investigators have suggested that empirical evidence for caste-specific DNA methylation in social insects is weak (Libbrecht et al. 2016). However, relatedness of species has not been considered in previous studies, and is necessary to control for non-independence of species relationships (Felsenstein 1985). If sociality is dependent on DNA methylation we would expect a gain in DNA methylation to result in either a gain of social behavior or a predisposition to evolve social behavior. Thus, if sociality is dependent on DNA methylation we would expect a phylogenetically-corrected correlation between these two traits. We do not find this. Sociality – including division of labor – is not evolutionary dependent on DNA methylation (Table 1). This lack of dependence holds true when DNA methylation is categorized as a discrete or continuous trait, and sociality is categorized as presence/absence or into multiple discrete classes (Fig. 1; Fig. 2A; Table 1). Although there are differences in transition rates between social behavior and DNA methylation states, they are not dependent on one another (Table 1; Fig. S2). For example, transitions from social to solitary occur more frequently than solitary to social when DNA methylation is fixed (Fig. S2). Also, transitions from solitary to social, *in lieu* of DNA methylation, occur more frequently than transitions from social to solitary (Fig. S2). Furthermore, based on Akaike information criterion (AIC), an independent model of evolution for each trait is preferred (Table 1). We further show that these traits are not dependent when 35 species within Hymenoptera are considered (Table 1). As dipterans composed a large fraction of species investigated in this study and do not have DNA methylation (Fig. 1 and 2), we were concerned that they may be biasing our analyses. However, when all species within Diptera are removed we similarly recover a lack of dependence between sociality and DNA methylation (Table 1). Together this evidence supports the hypothesis that sociality and DNA methylation are not evolutionary dependent.

### Evolution of DNMT1, DNMT2 and DNMT3 and DNA methylation

DNMT1 is found in all orders of insects investigated in this study except Diptera (Fig. 3A; Table S2). Only DNMT2 is found in Diptera, which reflects the lack of DNA methylation in the genomes of all species belonging to this order (Fig. 2B; Table S1). The loss of DNA methylation most parsimoniously occurred once in all Diptera ∼206.28-107.26 MYA (Misof et al. 2014), which likely coincided with the loss of DNMT1 and/or DNMT3.

**Figure 3.**
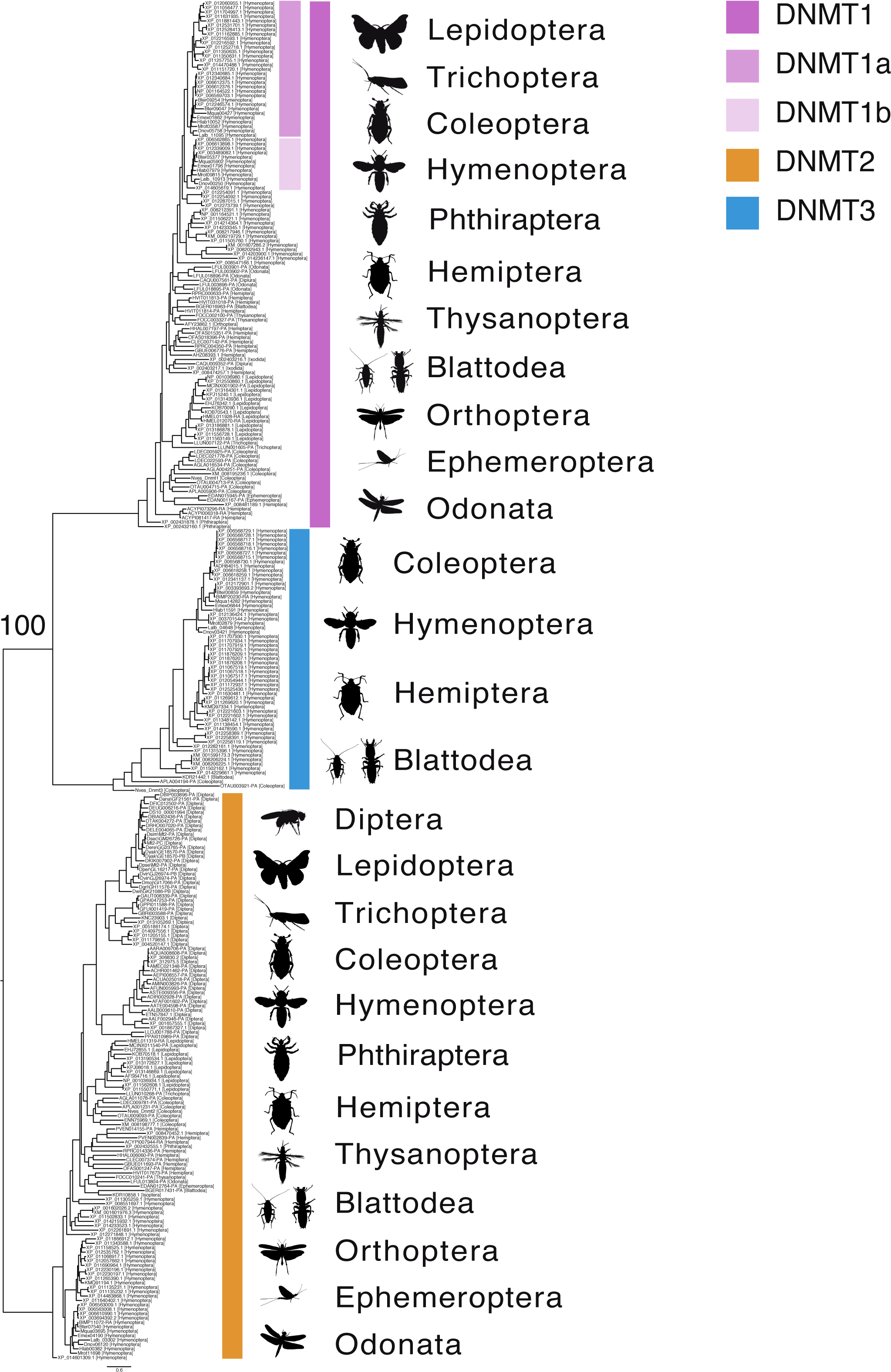
Evolution of DNMT1, 2 and 3 across insects. (A) Relationships of DNMT1, 2, and 3 in insects, Diplura and Ixodida (Arachnida). DNMT2 can be found in all insect orders investigated, while DNMT1 and 3 are more order-poor. The insect order for each sequence is provided in square brackets following the GenBank or genome annotation accession number. The tree was rooted to DNMT2.

Only DNMT1 and DNMT2 are found in lepidopteran species (Fig. 3A; Table S2). This observation supports the hypothesis that DNMT3 was lost from species belonging to this order between ∼177.99-116.45 MYA (Misof et al. 2014). Also, the sister order to Lepidoptera, Trichoptera, also only possess DNMT1 and DNMT2. Therefore, the timing of this loss may be older, however this order is represented by only one species in our data. Despite missing DNMT3 from assembled transcriptomes or genomes – as in the lepidopteran *Bombyx mori* – evidence for DNA methylation is still observed from WGBS (Table S1) data and from distributions of *C_p_G_o/E_* values (Fig. S1) and WGBS (Table S1) (Xiang et al. 2010). Similarly, 2/7 Coleoptera and 9/10 Hemiptera only possess DNMT1 and DNMT2, and DNA methylation is expected to be present given bimodality of *C_p_G_o/E_* (Table S1).

Several duplications were observed in the DNMT1 clade, but none coincide with presence or absence of DNA methylation or sociality. A duplication event shared by many hymenopteran gave rise to what is referred to as DNMT1a and DNMT1b (Fig. 4). Differences in codon and amino acid alignment tree topology suggest all hymenopterans or bees and ants shared the duplication, respectively (Fig. 4A; Fig S4). The amino acid alignment tree topology has higher node support than the codon alignment, which provides stronger support for the duplication event being shared by the superfamilies Apoidea (bees) and family Vespoidea (ants and some wasps) and places the timing for this duplication ∼123-225 MYA (Ronquist et al. 2012) (Fig. 4A; Fig. S4). Species relationships suggest reciprocal losses among families within the superfamily Vespoidea: a loss of DNMT1a in Vespidae (some wasps) and a loss of DNMT1b in Formicidae (ants). Whereas DNMT1a and DNMT1b can both be found in the superfamily Apoidea (families Apidae, Halictidae, and Megachilidae). Chalcid wasps (family Chalcidoidea) experienced at least two superfamily-specific duplication events, as suggested by the monophyletic groups containing *Nasonia vitripennis* (Fig. 4A). Additional species-specific duplications – inparalogs – were observed in orders Hemiptera and Lepidoptera (Fig. 3), but these could represent allelic variation of DNMT1.

**Figure 4.**
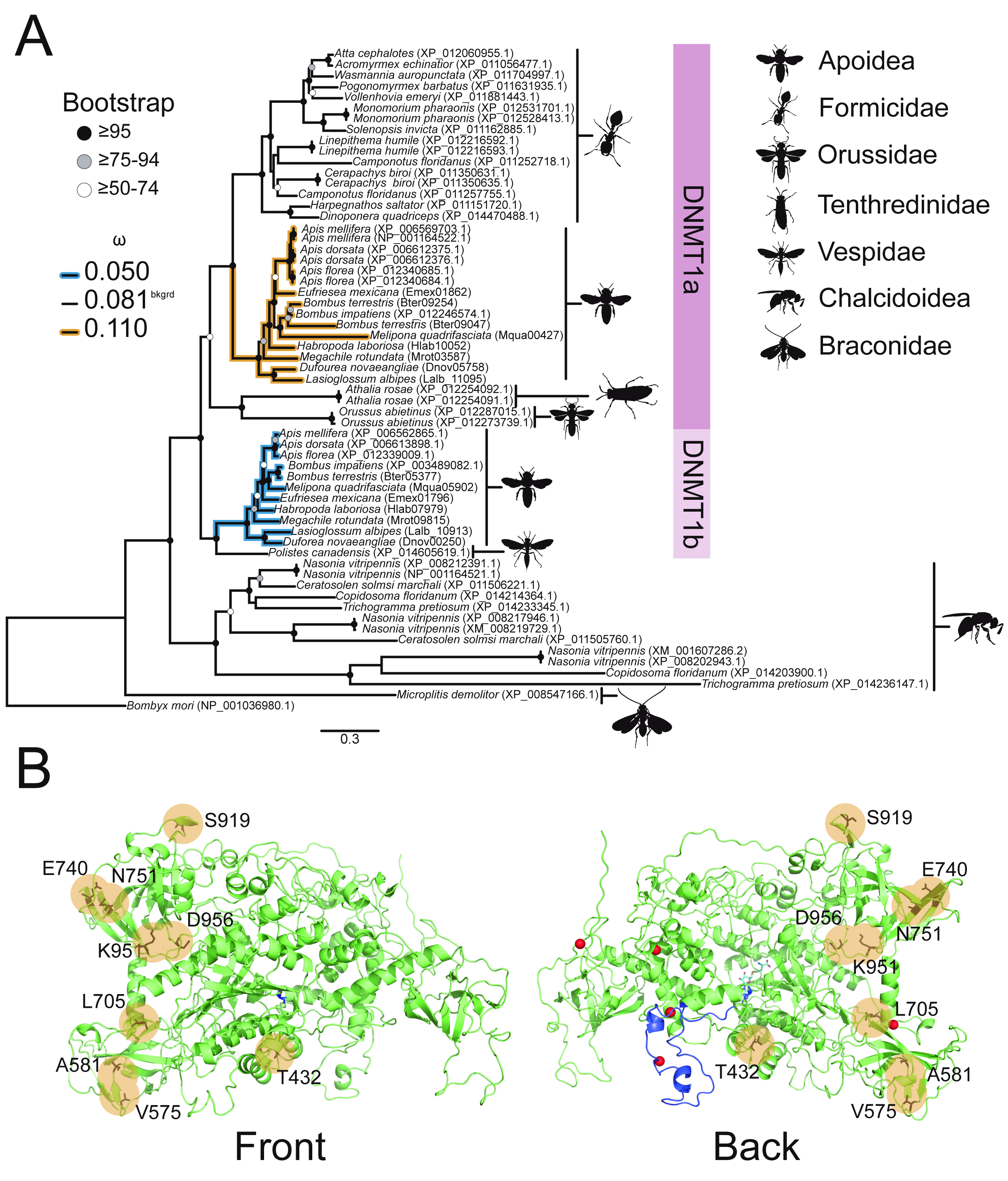
Divergent non-neutral evolution of DNMT1a and b in Apoidea. (A) Hypothesized relationships among DNMT1 in Hymenoptera suggests a duplication event shared by the superfamilies Apoidea (bees) and Vespoidea (ants and some wasps) gave rise to what is referred to as DNMT1a and b (Fig. S4). DNMT1a appears to have been lost from Vespidae (some wasps) and DNMT1b from Fromicidae (ants), whereas both DNMT1a and DNMT1b were retained in Apoidea (families Apidae, Halictidae, and Megachilidae). Divergent selection between DNMT1a and DNMT1b in Apoidea suggests the former is under relaxed purifying selection and the latter is under purifying selection. *Bombyx mori* (Lepidoptera) was used to root the tree, and was excluded from PAML analyses. (B) Several sites in DNMT1b were identified as under positive selection (yellow circles), with one site (T432) in the CXXC zinc finger domain, and three (V575, A581, S919) in the BAH domain. The crystal structure of *Apis mellifera* DNMT1b was predicted from *Mus musculus* DNMT1. Details including red spheres, dark and light blue colouring specify Zn^2+^ ions, CXXC domain and s-adenosyl-l-homocysteine, the secondary product from the DNA methylation reaction it performs, according to *Mus musculus* DNMT1, respectively. Bootstrap support values are characterized as shaded circles. *dN/dS* (ω) values are for the most preferred branch model.

Gene duplication is a source of genetic novelty and functional divergence, which can be characterized by non-neutral evolution (Lynch and Conery 2000). Divergent selection is observed between DNMT1a and DNMT1b in Apoidea (bees), with DNMT1a having experienced relaxed purifying selection and DNMT1b purifying selection (Fig. 4). Furthermore, positively selected sites were identified in DNMT1b, and these include one site (T432) in the CXXC zinc finger domain, and three (V575, A581, S919) in the BAH domain (Fig. 4). DNMT1 paralogs might maintain DNA methylation differently, including efficiency or rate, and spatially and/or temporally. The contribution of each paralog to the maintenance of the DNA methylome is unknown, and functional studies are required to address the fates of these duplicated genes.

DNMT3 is the most order-poor of the DNA methyltransferases, and was only identified in species belonging to Blattodea, Coleoptera, Hemiptera, and Hymenoptera (Fig. 3A; Table S2). For species with annotated genomes, DNMT3 is always accompanied by the presence of DNMT1 (Table S2). Given the scarcity of DNMT3 in insects and positive association between the presence and levels of DNA methylation and DNMT1 only, suggests DNMT3 may be dispensable for DNA methylation or DNMT1 compensates for DNMT3. However, DNMT1 is missing protein domains typically associated with metazoan *de novo* DNA methyltransferases. For example, DNMT1 does not contain a PWWP domain (PF00855), which interacts with DNA and histone lysine modified nucleosomes (Qiu et al. 2010; Qin et al. 2014). PWWP domain containing proteins exist (e.g., *Bombyx mori*), however none of these proteins contain a C-5 cytosine-specific DNA methylase domain (PF00145), which is required for DNA methylation (Bestor and Verdine 1994; Goll et al. 2005; Cheng et al. 2008). Based on these observations three possible alternative mechanisms explain the presence of DNA methylation *in lieu* of typical enzymes: (i) *de novo* DNA methylation does not occur via a DNMT-like protein; (ii) DNA methylation is not reprogrammed during embryogenesis and is robustly maintained by DNMT1 during each cell replication; or (iii) as has been shown in a plant species (Bewick, Li, Niederhuth, Willing et al. 2016; Bewick et al. 2016), non-neutral evolution could have occurred at DNA methyltransferases, which affects their ability to methylate or maintain methylation. However, alternative mechanisms might exist, and functional tests of DNA methylation mechanisms in insects warrant further investigation.

## CONCLUSION

We have conducted the largest phylogenetic investigation of DNA methylation in insects. Our investigation suggests that DNA methylation is too variable in insects to support a single, common role in regulating traits expressed in social interactions. DNA methylation is common in insects, but the level of DNA methylation is fluid and is not more likely to be associated with the presence of sociality. Although this does not preclude a role for DNA methylation in regulating caste in eusocial insects, one possibility is that DNA methylation in most insects that display sociality is a downstream consequence of upstream environment, genetic or hormonal factors (Kapheim et al. 2015; Wallberg et al. 2016). Furthermore, the gain, loss, and duplication of DNMT1 suggests that there may be novel functions and justifies further studies using RNAi or CRISPR/Cas9 to mutate DNA methyltransferases (Gilles et al. 2015). For now, the universal role of DNA methylation in insects remains elusive, and is likely to remain so until functional tests are made to advance our understanding of this evolutionary conserved and important chromatin modification.

## MATERIALS AND METHODS

### Tissue collection and DNA extraction

All samples were collected from established laboratory colonies with the following exceptions. *Cryptocercus garciai* was collected from northeast Georgia by Brian Forschler, *Polistes carolina* were collected from the campus of the University of Georgia by K. J. Vogel.

DNA extractions were performed on tissues from either freshly sacrificed insects or ethanol preserved samples. For samples likely to contain significant gut microbial contamination (Blattodea), guts were dissected and discarded. Approximately 10 mg of material was frozen in liquid nitrogen, ground with a pestle. Samples were then extracted with the Qiagen DNEasy Mini kit following the manufacturer’s instructions for animal tissues.

### Behavior classification

Insect species were classified into one of four social behavior categories based on the presence or absence of well-defined traits (Wilson 1971; Costa 2006). *Eusocial* insects must share a common nest site, exhibit cooperative rearing of young, reproductive division of labor with sterile or less fecund workers, and have overlap of generations. Species that exhibited only parental care or cooperative brood rearing were classified as *subsocial*. Species with individuals of the same generation that use the same composite nest site and do not cooperate in brood care were classified as *communal*.

### Whole-genome bisulfite sequencing and levels of DNA methylation

MethylCseq libraries were prepared according to the following protocol (Urich et al. 2015). Sequencing data for *Acyrthosiphon pisum*, *Aedes aegypti*, *Aedes albopictus*, *Anopheles gambiae*, *Apis mellifera*, *Blattella germanica*, *Camponotus floridanus*, *Copidosoma floridanum*, *Culex quinquefasciatus*, *Dinoponera quadriceps*, *Drosophila melanogaster*, *Harpegnathos saltator*, *Microplitis demolitor*, *Nasonia vitripennis*, *Nicrophorus vespilloides*, *Polistes canadensis*, *Solenopsis invicta*, *Tribolium castaneum* was aligned to their respective genome assembly using the methylpy pipeline (Schultz et al. 2015) (Table S1). *Blattella asahinai* was aligned to its sister species *Blattella germanica*. Similarly, *Microplitis mediator* was aligned to its sister species *Microplitis demolitor*. *Dinoponera quadriceps* was removed from Fig. 1 and associated analyses because of possible genomic sequence contamination with other insect species. Species for MethylC-seq were chosen to maximize taxonomic breadth based on previously published insect DNA methylomes. Thus, species within the same genus with published methylomes were often limited to a single representative. However, *a priori* knowledge of closely related species with differences in social behavior was taken into consideration. Overall, the species chosen for MethylC-seq represent phylogenetically deep and shallow switches for social behavior. Other species, including *Ceratina calcarata, Locusta migratoria* and *Schistocerca gregaria*, were excluded because no genome annotation exists on GenBank.

In brief, reads were trimmed of sequencing adapters using Cutadapt (Martin and Marcel 2011), and then mapped to both a converted forward strand (cytosines to thymines) and converted reverse strand (guanines to adenines) using bowtie (Langmead et al. 2009). Reads that mapped to multiple locations, and clonal reads were removed. Weighted DNA methylation was calculated for each sequence context (CG, CHG and CHH) by dividing the total number of aligned methylated reads by the total number of methylated plus un-methylated reads. Slight differences in levels of DNA methylation can be found between this and previously published studies (e.g., Bonasio et al. 2012). These differences are shared by all studies and are most likely due to technical and methodological differences. The majority of these differences are negated within this study due to the use of identical methods applied to all samples, including mapping of MethylC-Seq reads, categorization of coding sequences, and estimates of DNA methylation levels. Thus, levels of DNA methylation are comparable within this study. Furthermore, the discrete categorization of DNA methylation is not affected by slight differences between different social castes or individuals of a population, as genome-wide reductions or gains in DNA methylation are not observed (Elango et al. 2009). For the remaining species, genome-wide levels of DNA methylation were estimated using *FAST^m^C* and the --animal model (Bewick et al. 2015).

CG DNA methylation in insects is enriched in coding sequences (see Lyko et al. 2010; Beeler et al. 2014) and a strong correlation between genome-wide and within coding sequence levels of CG DNA methylation is observed (Fig. S3). Therefore, using genome-wide levels of CG DNA methylation estimated from *FAST^m^C*, we were able to extrapolate levels of CG DNA methylation within coding sequence for species without sequenced genomes (Table S1).

### Measurement of *C*_*p*_*G*_*o*/*E*_ and tests for bimodality

*C_p_G_o/E_* is a metric of CpG dinucleotides normalized by G and C nucleotide content (GC content) and length (bp) of a specific region of interest (e.g., a transcript or protein coding gene) (Patalano et al. 2015). Due to spontaneous deamination of methylated cytosines, genes that are hypermethylated are expected to have a lower *C_p_G_o/E_* value than hypomethylated genes. Thus, in a mixture of genes that are methylated and low to un-methylated, a bimodal distribution of *C_p_G_o/E_* values is expected. Conversely, a unimodal distribution is suggestive of a set of genes that are mostly low to unmethylated. We therefore used this metric to expand our sampling of insect species for the presence of DNA methylation in 11 orders: Odonata (n=1), Ephemeroptera (n=1), Blattodea (n=2), Thysanoptera (n=1), Hemiptera (n=9), Phthiraptera (n=1), Hymenoptera (n=32), Coleoptera (n=7), Trichoptera (n=1), Lepidoptera (n=11), and Diptera (n=57). The *C_p_G_o/E_* value for each gene within 125 total transcriptomes or gene annotations was defined as:

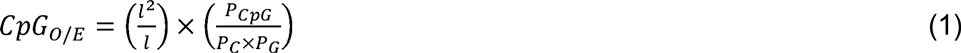

where *P_CpG_*, *P_C_*, and *P_G_* are the frequencies of CpG dinucleotides, C nucleotides, and G nucleotides, respectively, estimated from each gene of length (*l*) in bp. Only exonic sequences of a gene were considered when estimating *C_p_G_o/E_*.

The modality of *C_p_G_o/E_* distributions was tested using Gaussian mixture modeling (mclust v5.2). Two modes were modeled for each *C_p_G_o/E_* distribution, and the subsequent means and 95% confidence interval (CI) of the means were compared with overlapping or non-overlapping CI’s signifying unimodality or bimodality, respectively. Based on the largest set of genomes and WGBS data to date, Gaussian mixture modeling using mclust identified 18/18 (100%) species correctly for presence/absence of DNA methylation within coding regions. Thus, *C_p_G_o/E_* modality tested with Gaussian mixture modeling is a robust and accurate predictor of DNA methylation.

### Phylogenetic comparative methods

The R package phytools (Revell 2011) was used for all phylogenetic comparative methods. Two tests of correlated evolution were conducted. (i) Pagel’s (Pagel 1994) method for detecting correlated evolution of two binary traits. Briefly, Pagel’s method uses a continuous-time Markov model to simultaneously estimate transition rates in pairs of binary characters on a phylogeny. These rates are then used to test whether an independent or dependent model of evolution is preferred using the likelihood ratio test (LRT). DNA methylation was categorized into un-methylated or methylated based on the distribution of *C_p_G_o/E_*, unimodal or bimodal, respectively. Sociality was categorized into social (eusocial, subsocial, and communal) or solitary binary traits. Similarly, the *division of labor* trait was categorized into species with (caste^+^) or without (caste^−^) castes. (ii) Phylogenetic generalized least squares (PGLS) was used to correlate continuously categorized estimates of DNA methylation generated through WGBS and alignment to reference assemblies (methylpy), or non-referenced based methods (*FAST^m^C*) to discretely coded social traits (eusocial, subsocial, communal, and solitary). BEAST 2 (Bouckaert et al. 2014) was used to estimate a multilocus coalescent tree, which was used to control for relatedness of species (non-independence) for both comparative tests. For (i) the phylogenetic tree was estimated from a subset of previously identified orthologous protein coding loci (58 of the >1400 described in Misof et al. 2014). These loci were chosen because each contained no more than 5% missing species for 123 insects, and outgroups *Catajapyx aquilonaris* (Diplura) and *Daphnia pulex* (Crustacea) based on best BLASTp hit to a set of core species (*Acyrthosiphon pisum*, *Anopheles gambiae*, *Acromyrmex echinatior*, *Bombyx mori*, *Drosophila melanogaster*, *Apis mellifera*, *Ixodes scapularis*, *Nasonia vitripennis*, *Pediculus humanus*, *Tribolium castaneum*, *Zootermopsis nevadensis*, *Daphnia pulex*). Loci were aligned using PASTA (Mirarab et al. 2015), and Gblocks was used to identify sections of conserved protein coding sequence (Castresana et al. 2000). For (ii) a smaller number of loci were used due to the obscurity of species and available data on Genbank. Loci used for both trees can be found in Table S3. For both trees, the program Tracer (http://tree.bio.ed.ac.uk/software/tracer/) was used to assess stationarity and effective sample size (ESS; ≥100) of the Markov Chain Monte Carlo (MCMC) chains.

### DNA methyltransferase (DNMT) phylogeny and evolution

DNA methyltransferases (DNMT1, 2, and 3) were curated from 125 insect species through homology searches using BLASTp and previously identified DNMT1, 2, and 3 proteins in *Apis mellifera*, *Bombyx mori* (DNMT1 only), *Drosophila melanogaster* (DNMT2 only), and *Nicrophorus vespilloides*. A series of alignment and phylogenetic estimation steps were conducted eliminate partial and poor sequences, which can affect alignment and subsequently the topology of phylogeny. Protein sequences were aligned using the program PASTA, and back-translated using the CDS sequence to generate an in-frame codon alignment. RAxML with 1000 rapid bootstrap replicates and the GTR+G model of nucleotide substitution was used to generate the phylogeny (Stamatakis 2014). The phylogeny was rooted to the DNMT2 clade in the program FigTree (http://tree.bio.ed.ac.uk/software/figtree/) and exported for stylization. An identical method was used to generate the Hymenoptera-specific DNMT1 gene tree. Nine iterations of alignment and phylogeny construction were performed removing sequences on long branches, which is indicative of low sequence homology. For species with assembled transcriptomes or gene annotations, additional DNA methyltransferases were identified by Interproscan (Jones et al. 2014), and filtering those sequences with a DNA methylase domain (PF00145). These DNA methylase domain-containing sequences were then subjected to the previously mentioned homology searches using BLASTp.

### Codon analysis

Similar methodology as described above was used to construct phylogenetic trees for testing hypotheses on the rates of evolution in a phylogenetic context. However, the program Gblocks was used to identify conserved amino acids in codon alignments. The parameters for Gblocks were kept at the default settings, except we allowed for %50 gapped positions. The program Phylogenetic Analysis by Maximum Likelihood (PAML) was used to test branches (branch test) and sites along branches (branch-site test) for deviations from the background rate of molecular evolution (*dN/dS*; ω) and for deviations from the neutral expectation, respectively (Yang 2007). Positively selected sites were determined by Bayes Empirical Bayes (BEB) score of ≥0.95. Branches tested and a summary of each test can be found in Table S4.

## ACKNOWLEDGEMENTS

We thank Lauren A. Eserman and Ben L. S. Furman for phylogenetic comparative advice, and William T. Jordan for crystal structure visualization. We thank Patrick Abbot, Gaelen Burke, Brian Forschler, Sarah E. Sander, Coby Schal, Kathrin Stanger-Hall, and Michael Strand for tissue and/or insect specimens. We also thank Patrick T. Griffin, Libby McKinney, and Nick A. Rohr for library preparation, and the Georgia Genomics Facility (GGF) for sequencing. Computational resources were provided by the Georgia Advanced Computing Resource Center (GACRC). We thank the i5K initiative and the Baylor College of Medicine Human Genome Sequencing Center (BCM-HGSC) for the use of pilot data. This work was supported by funding from the Office of the Vice President for Research at the University of Georgia to RJS, and by NIH award F32GM109750 to KJV.

## AUTHOR CONTRIBUTIONS

The study was conceived by AJB and RJS. All authors were involved in the experimental design. KJV and AJM curated samples. KJV and AJM classified social behavior for insects used in this study. AJB performed all analyses, with contribution from KJV to the phylogenetic analyses. AJB wrote the manuscript with contributions made by all authors.

**Table 1. Sociality and DNA methylation are not evolutionary dependent.** Upper half: Output from Pagel’s method for evolutionary dependence (implemented in phytools^47^) using the phylogeny and traits from Fig. 2A. The dependent model of trait evolution is not preferred based on the likelihood ratio test (LRT), and the independent model is preferred based on Akaike information criterion (AIC). Lower half: Output from Phylogenetic Generalized Least Squares (PGLS) implemented in phytools) using the phylogeny and traits from Fig. 1. A model of zero (0) phylogenetic signal (*λ*) and evolution under Brownian motion was preferred over a model where *λ* was estimated by maximum likelihood. For both tests, P-values represent the significance of correlations between trait *x* and *y*. P-values for PGLS are given for the preferred model.

## SUPPORTING INFORMATION

**Figure S1. *C_p_G_o/E_* distributions for 123 insect species**. Gaussian mixture modeling was performed in R v2.3.4 to estimate uni- and bimodality. The dashed line represents the mean of each distribution, and the shaded area is the 95% confidence interval around the mean.

**Figure S2. Character state transition matrices from Pagel’s test for evolutionary dependence between two binary traits.** Transition rates between states are given for independent and dependent models of evolution. Greyed rates represent a rate of zero (0) between states.

**Figure S3. Genomic levels of DNA methylation are a good predictor of genic levels of DNA methylation.** Linear regression between genomic and genic levels of DNA methylation for 22 species of insects with sequenced genomes.

**Figure S4. The duplication of DNMT1 in Hymenoptera was shared by ants and bees.** (A) The hypothesized relationships of DNMT1 in Hymenoptera. The maximum likelihood tree was generated using an amino acid alignment and RAxML with the PROTGAMMAWAG model of protein evolution. Selected values at nodes represent bootstrap support from 1000 replicates. (B) Superfamily and family relationships suggest the duplication of DNMT1 in Hymenoptera was shared by ants and bees, and reciprocal losses of paralogs (DNMT1a and DNMT1b) occurred in Formicidae and Vespidae.

**Table S1**. Species and accompanying behavior, genomic, DNA methylation and sequencing information used in this study. Rows shaded in grey represent the 18 species where *C_p_G_o/E_* correctly predicted presence/absence of DNA methylation based on comparisons to WGBS.

**Table S2**. Species and DNA methyltransferase (DNMT) accession id’s used in Figure 3.

**Table S3**. A list of proteins used to estimate multilocus coalescent trees and the best BLAST hit to *Apis mellifera* or *Drosophila melanogaster*.

**Table S4.** Hypotheses tested using Phylogenetic Analysis using Maximum Likelihood (PAML) for DNMT1a and DNMT1b in Apoidea.

---

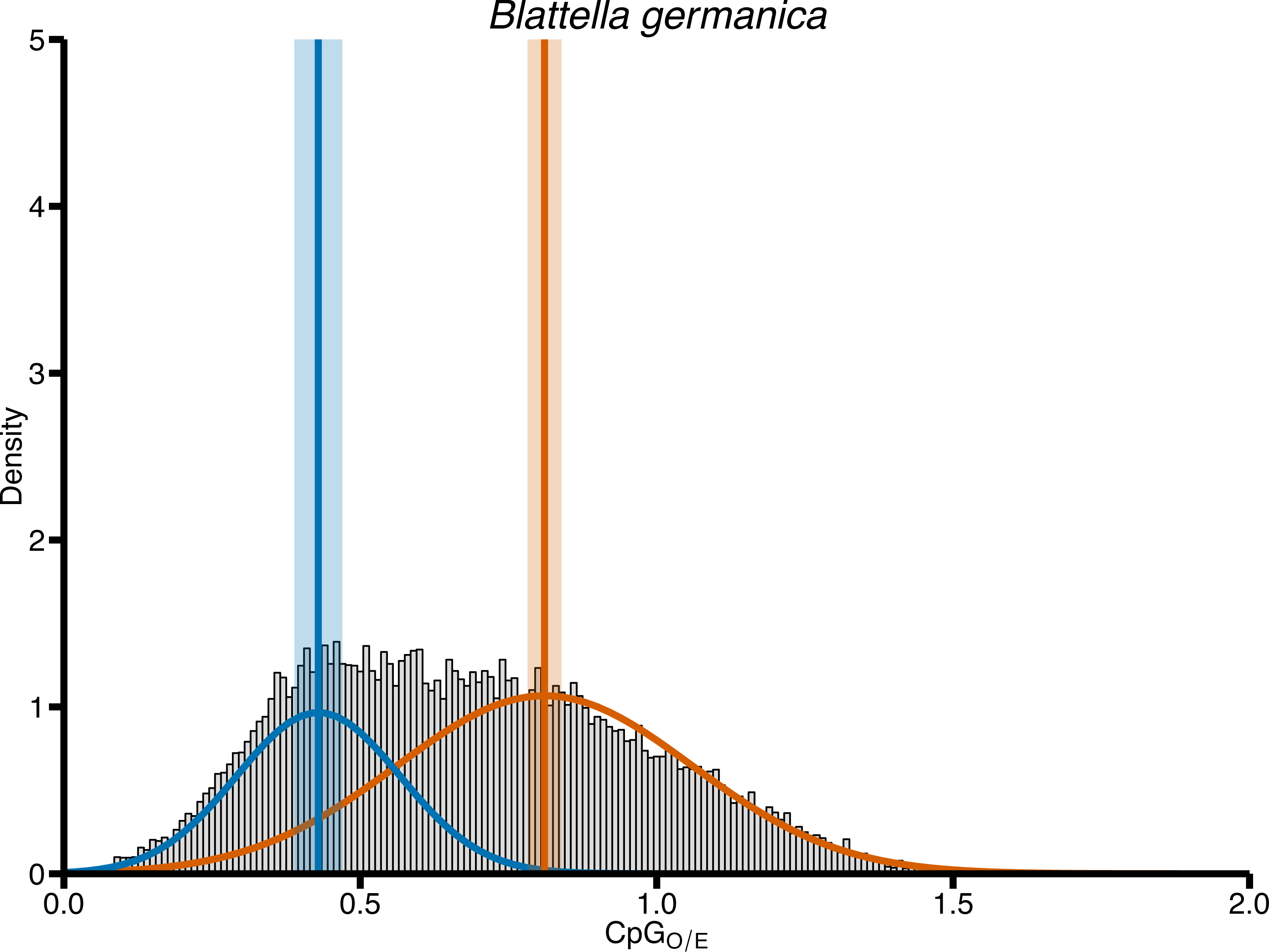

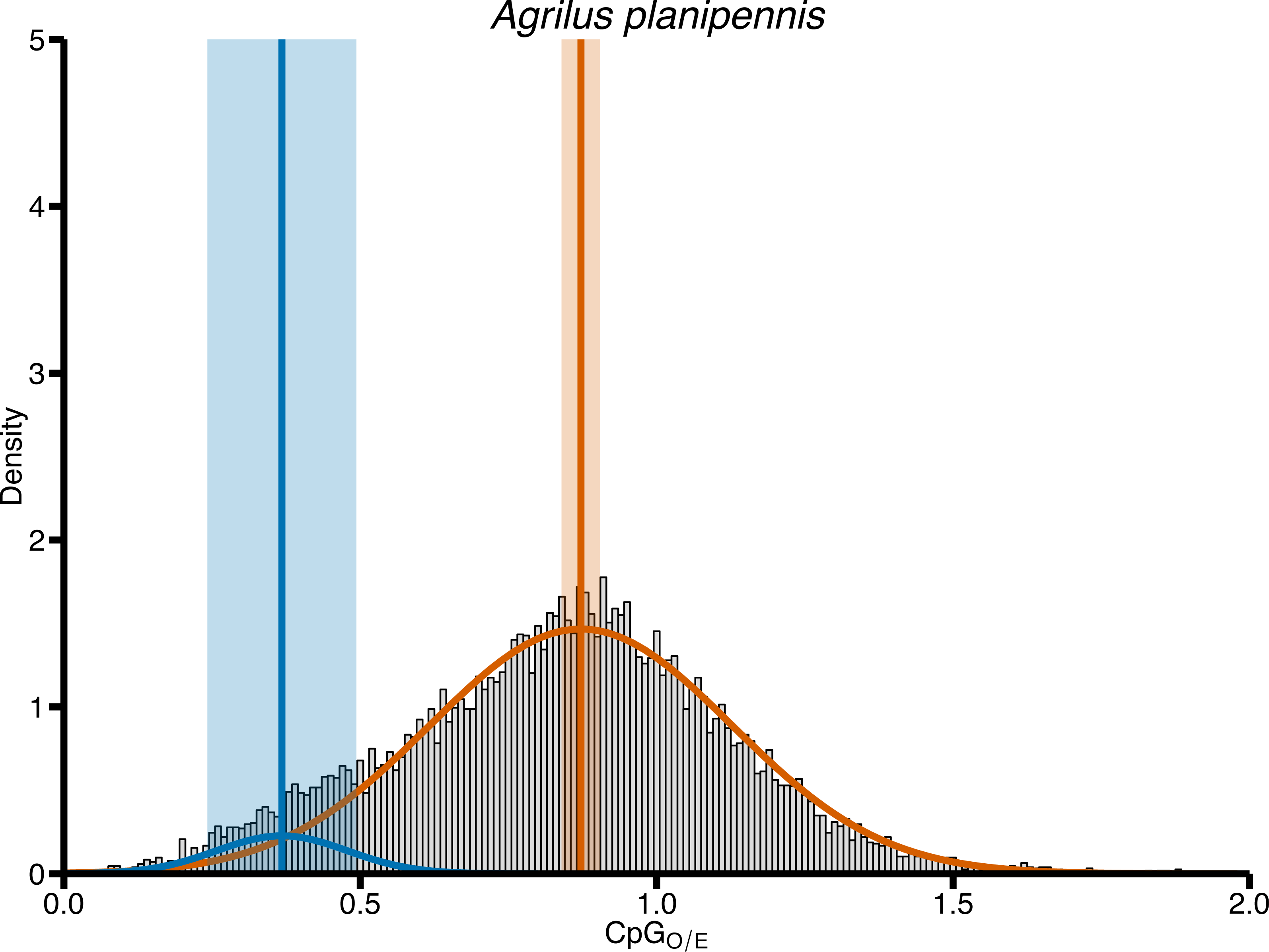

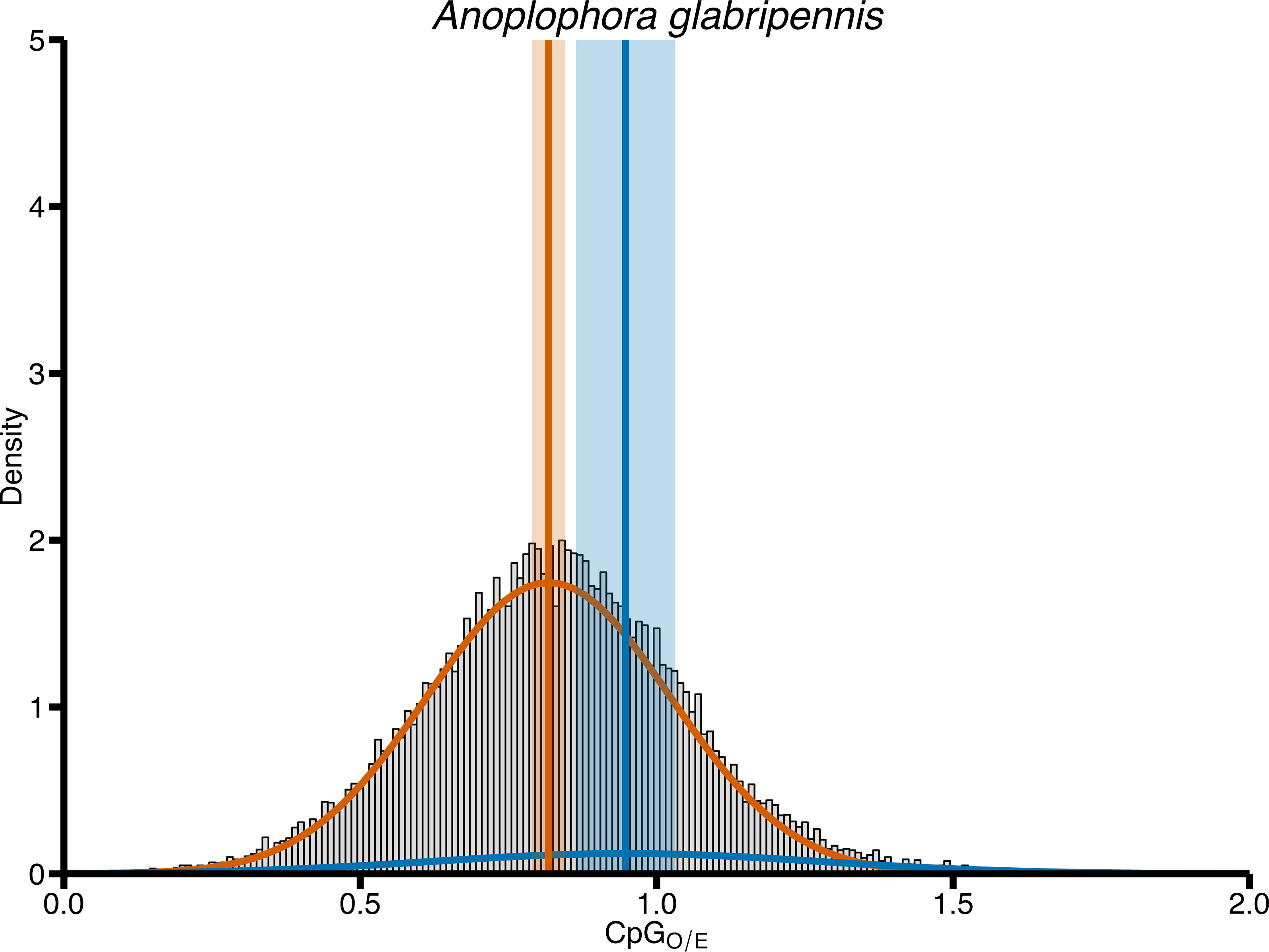

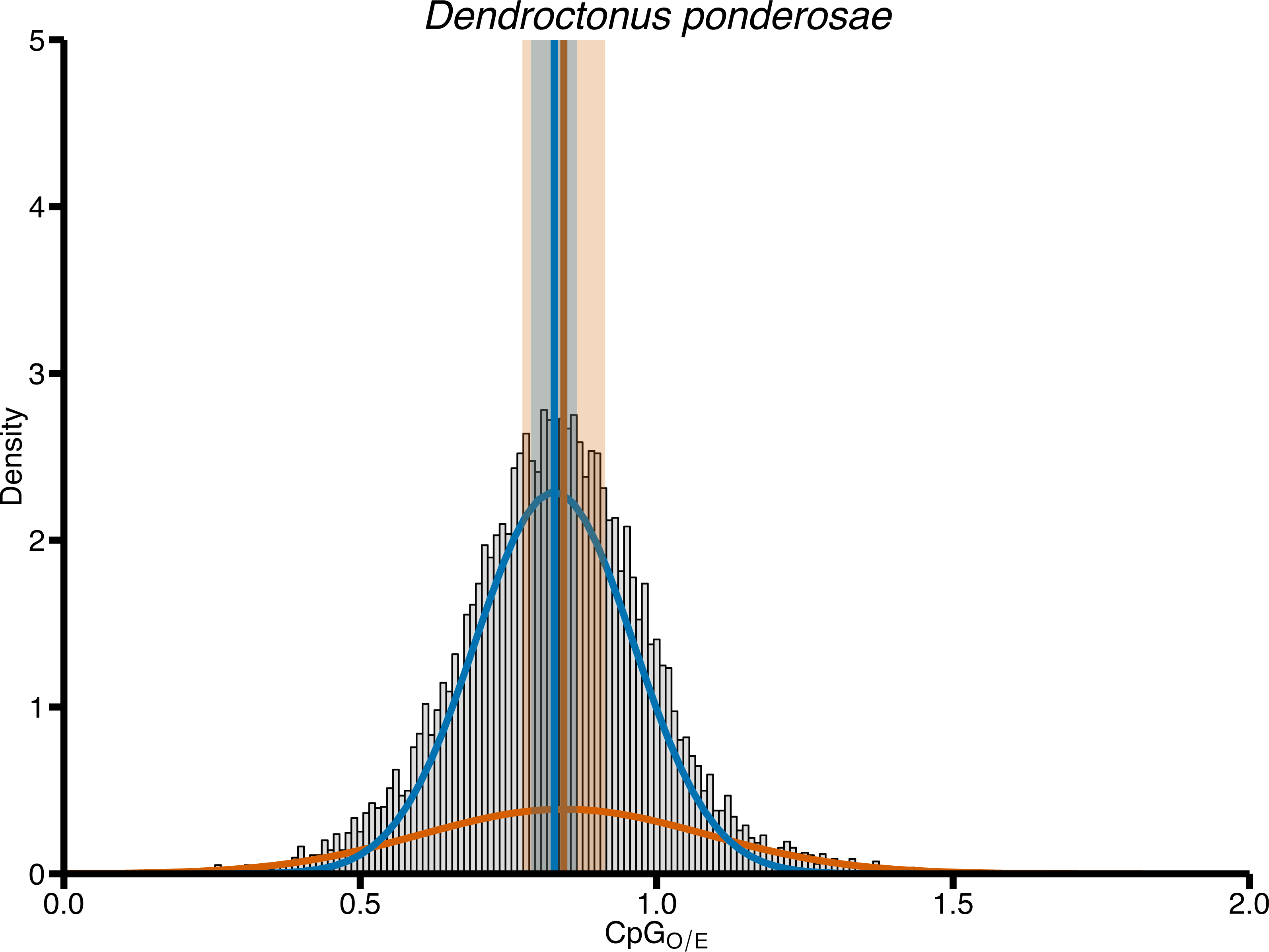

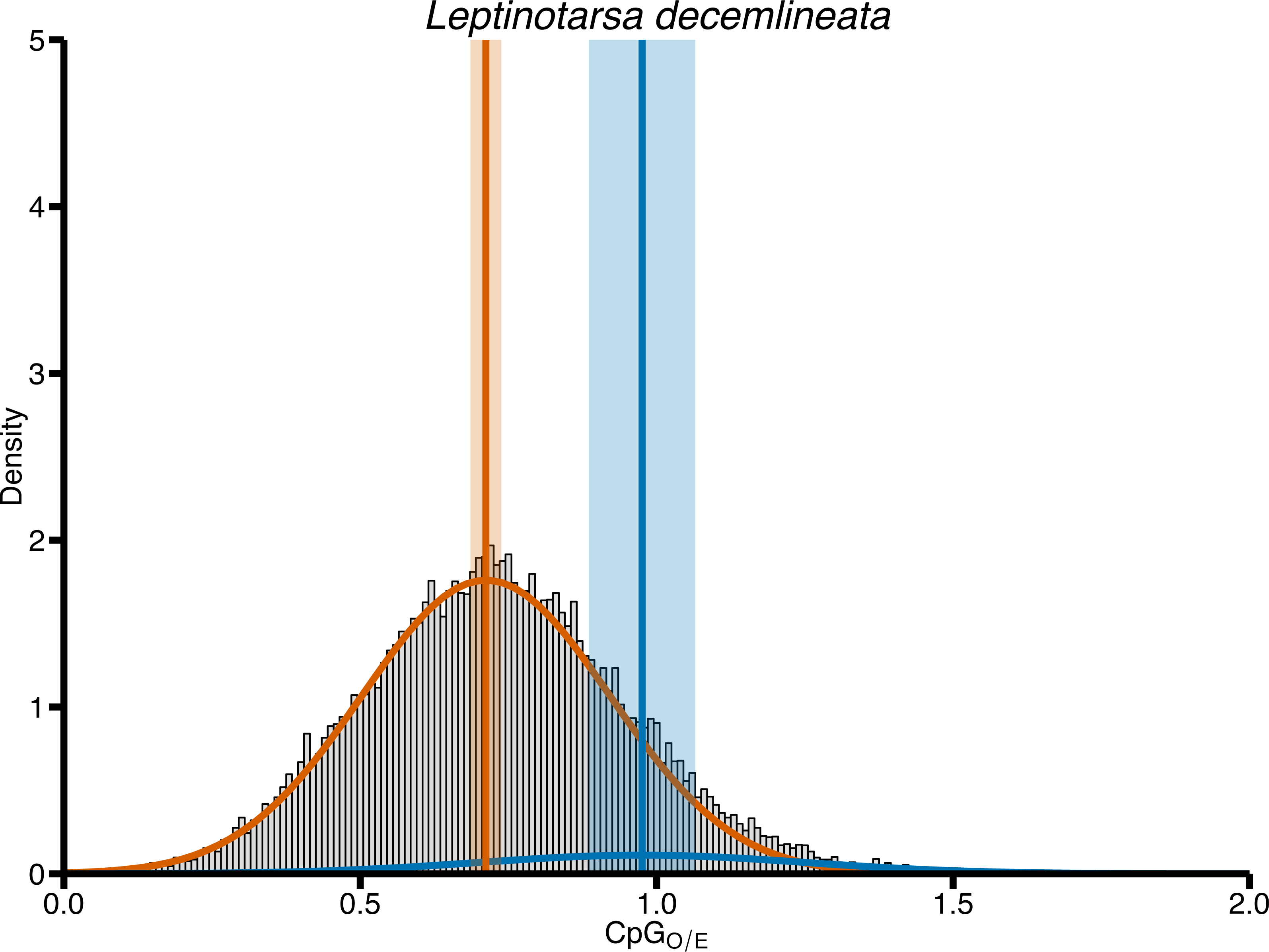

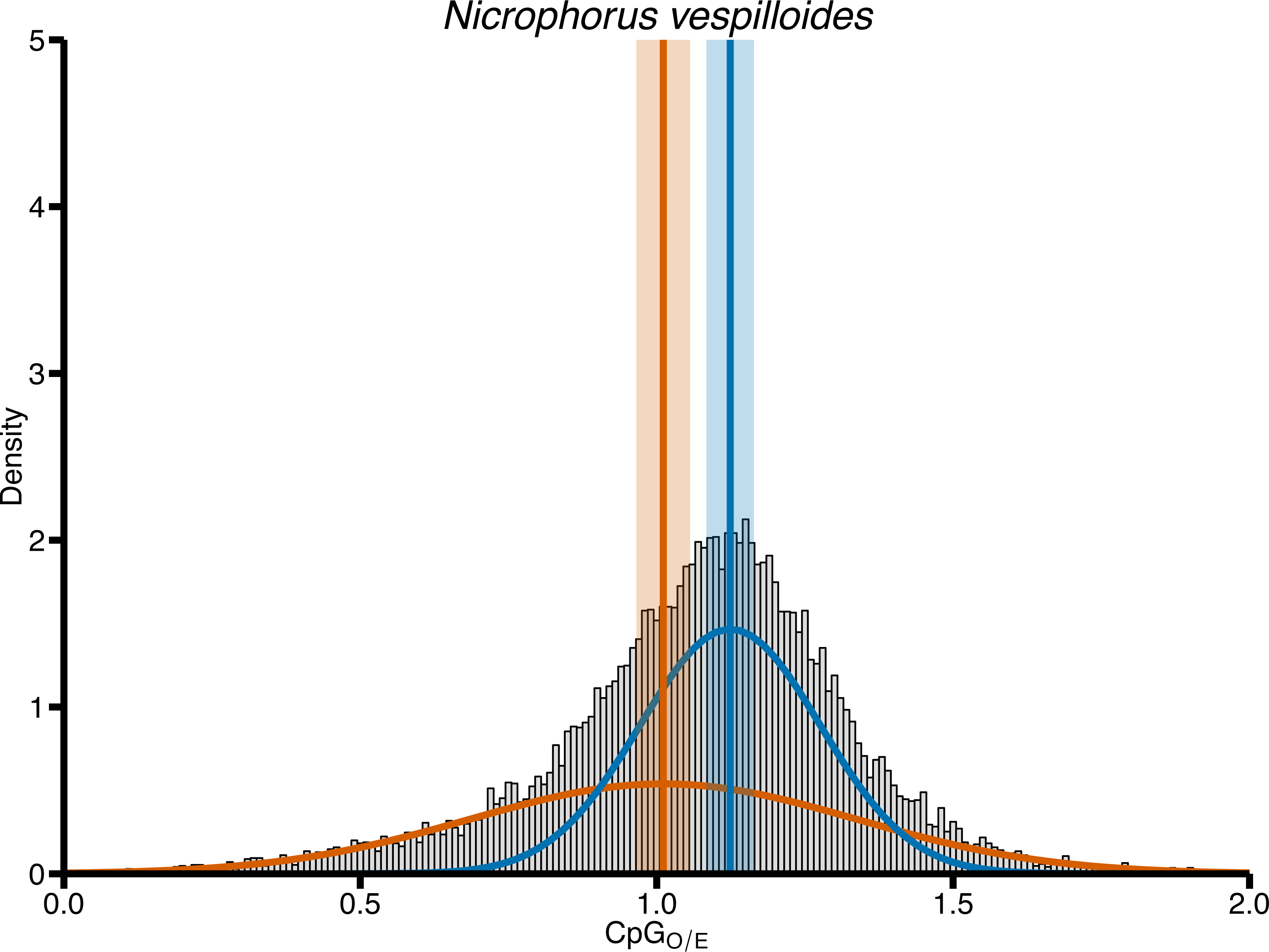

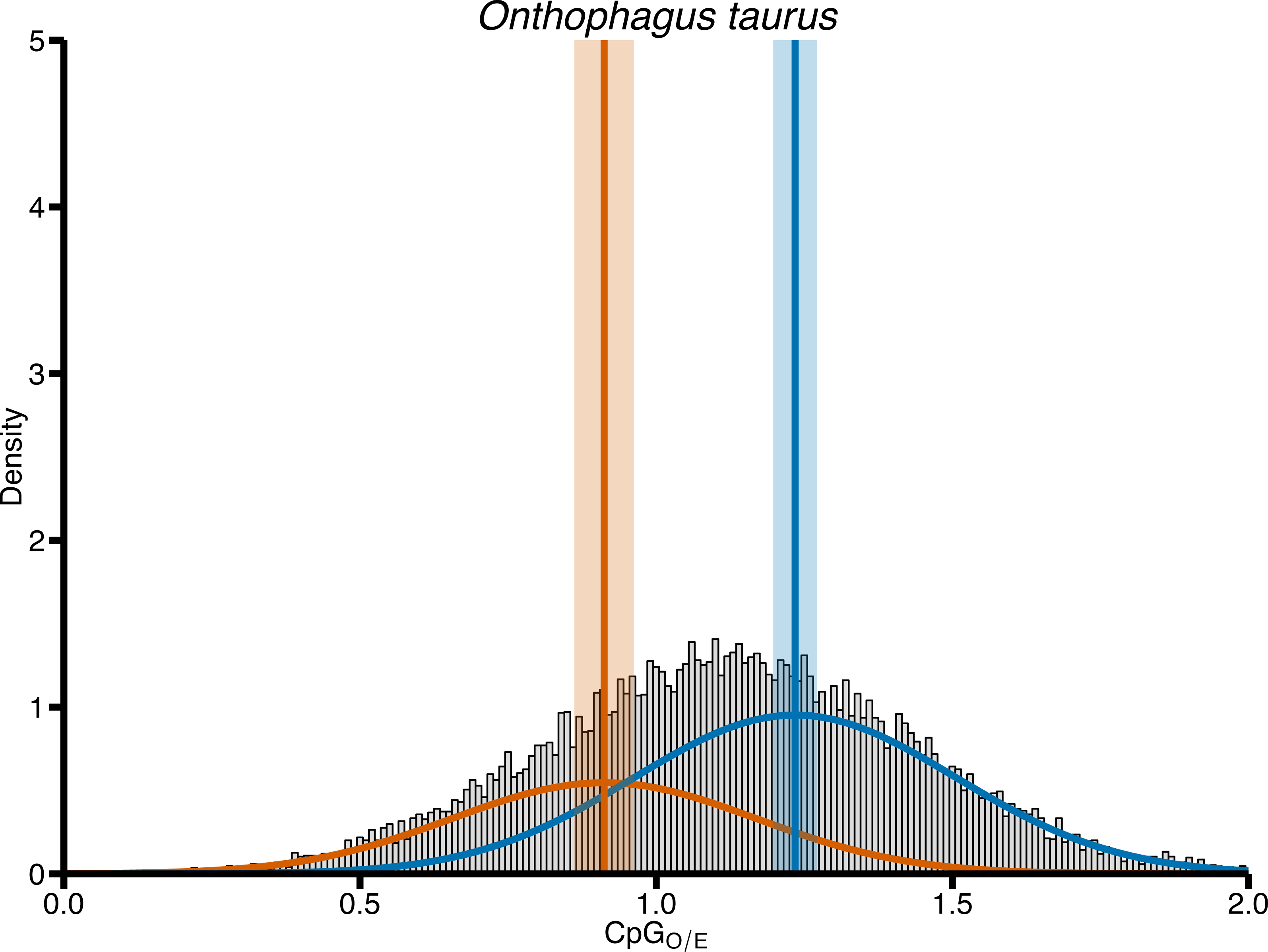

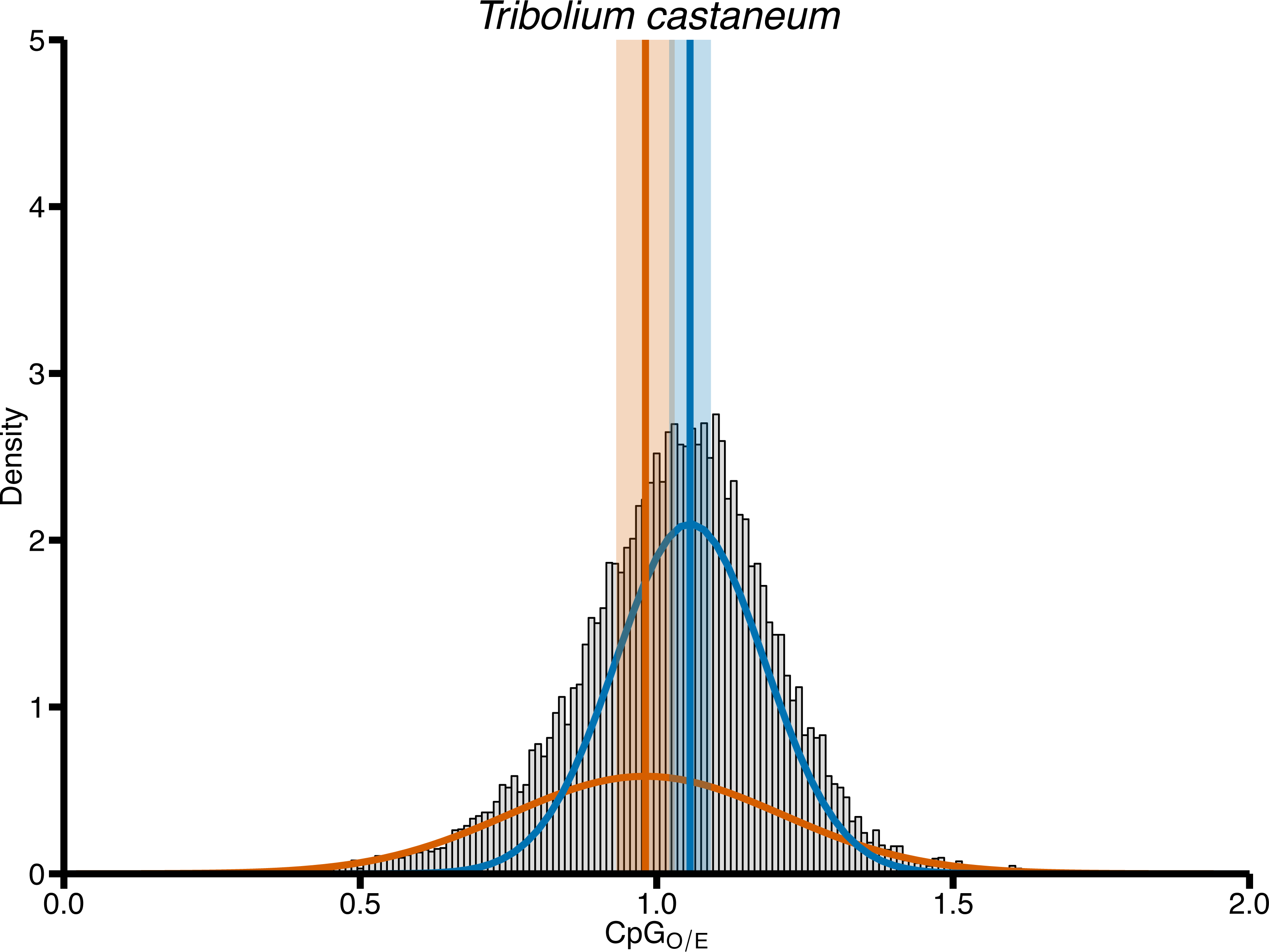

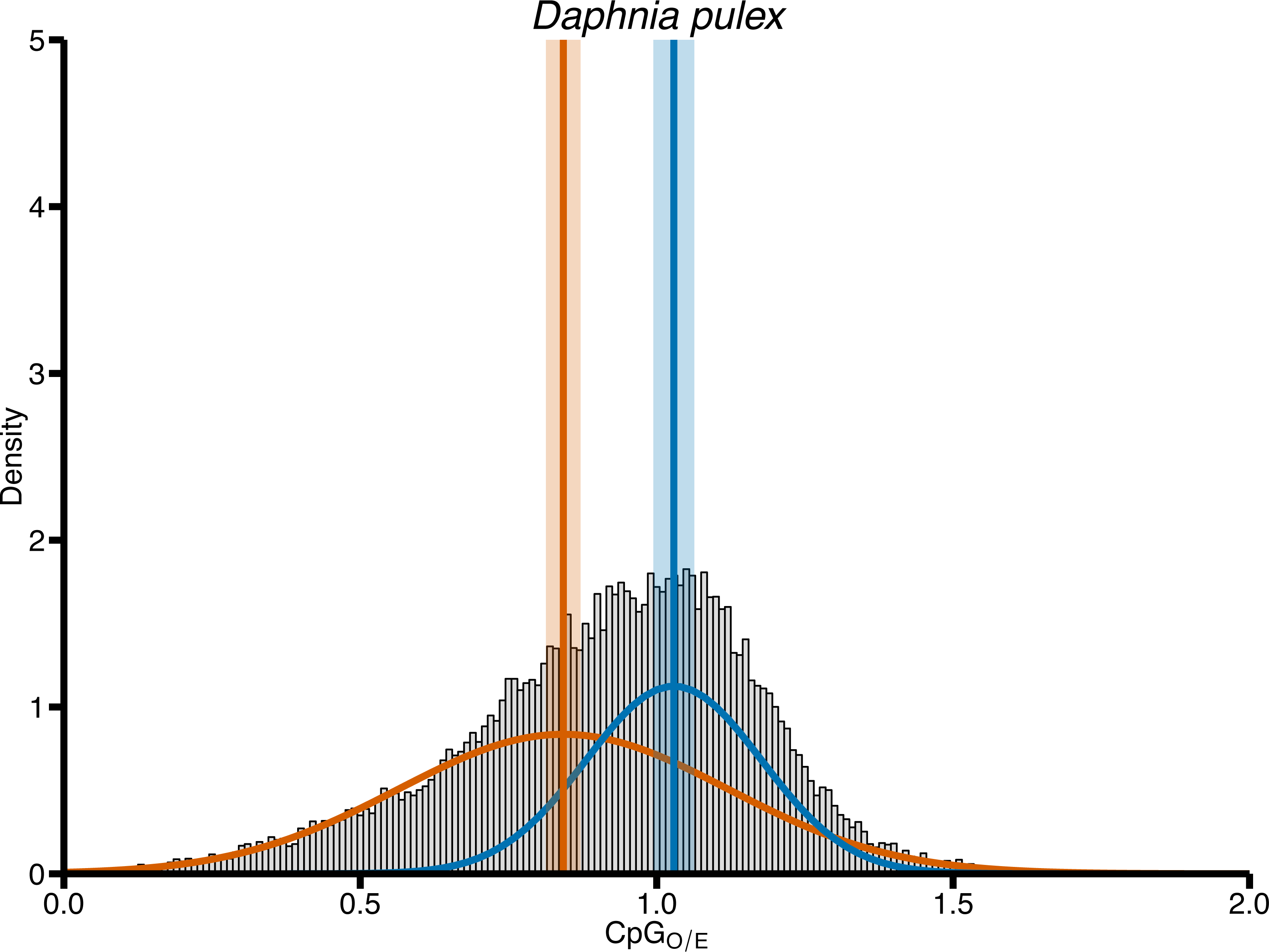

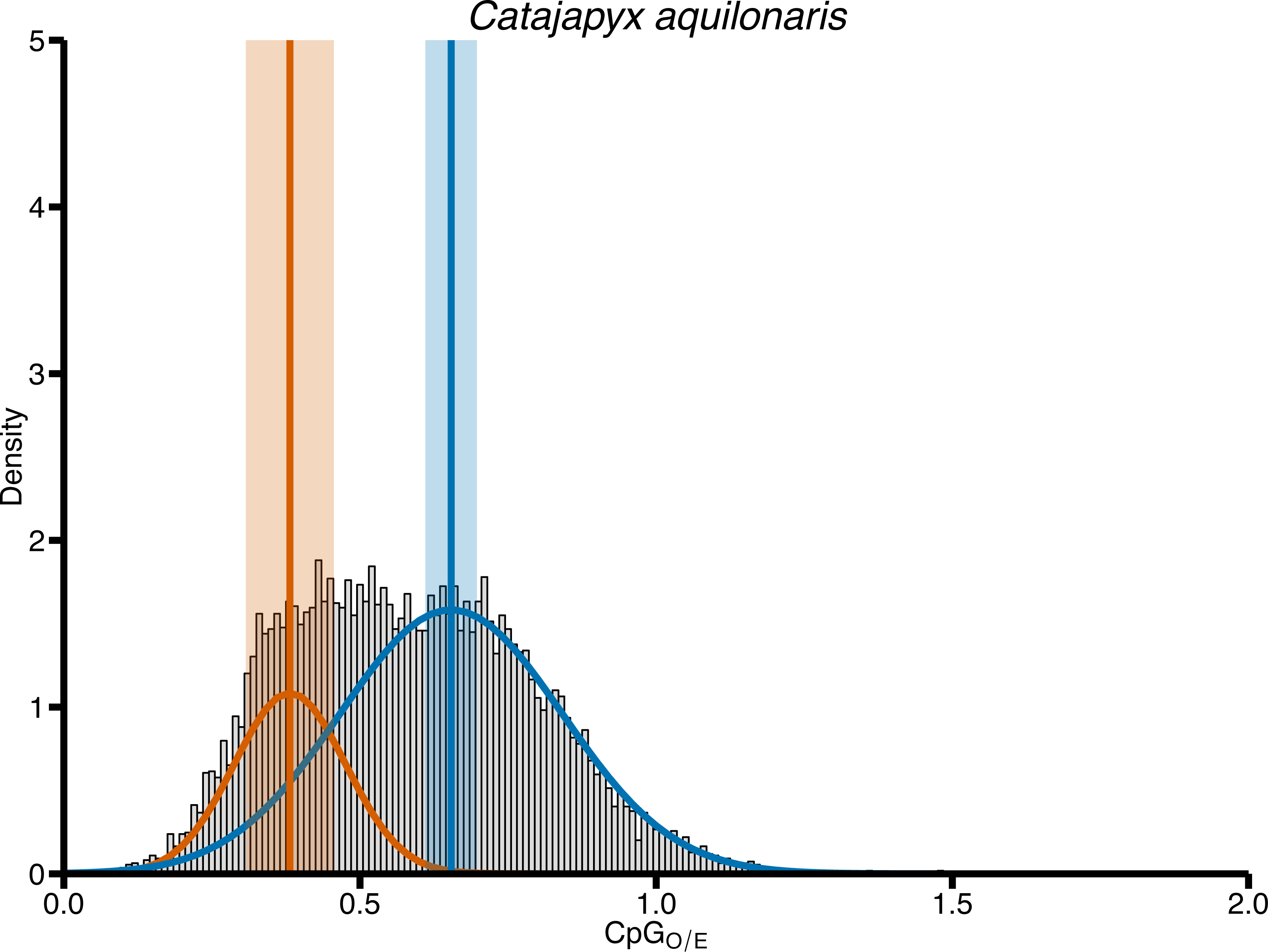

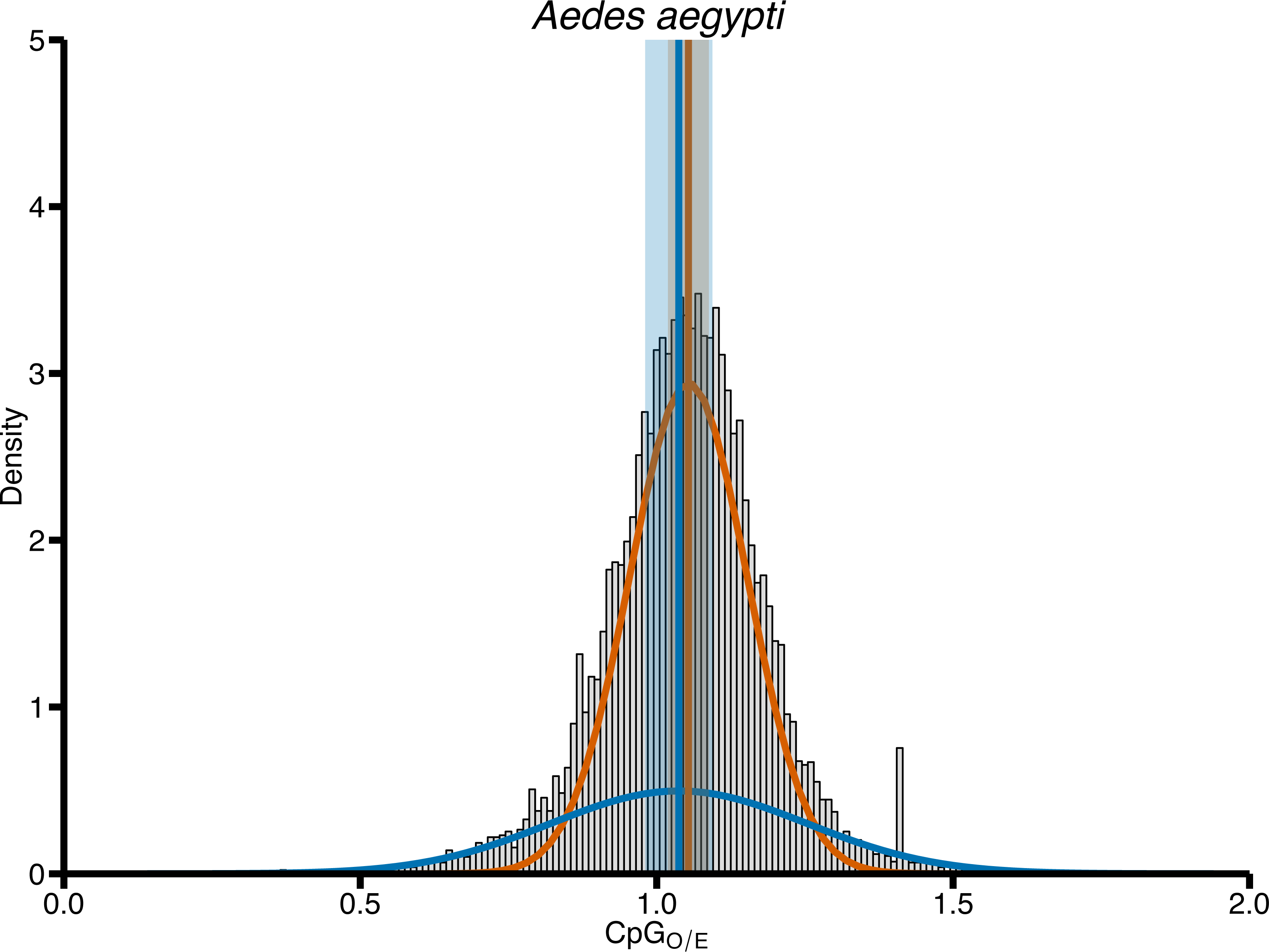

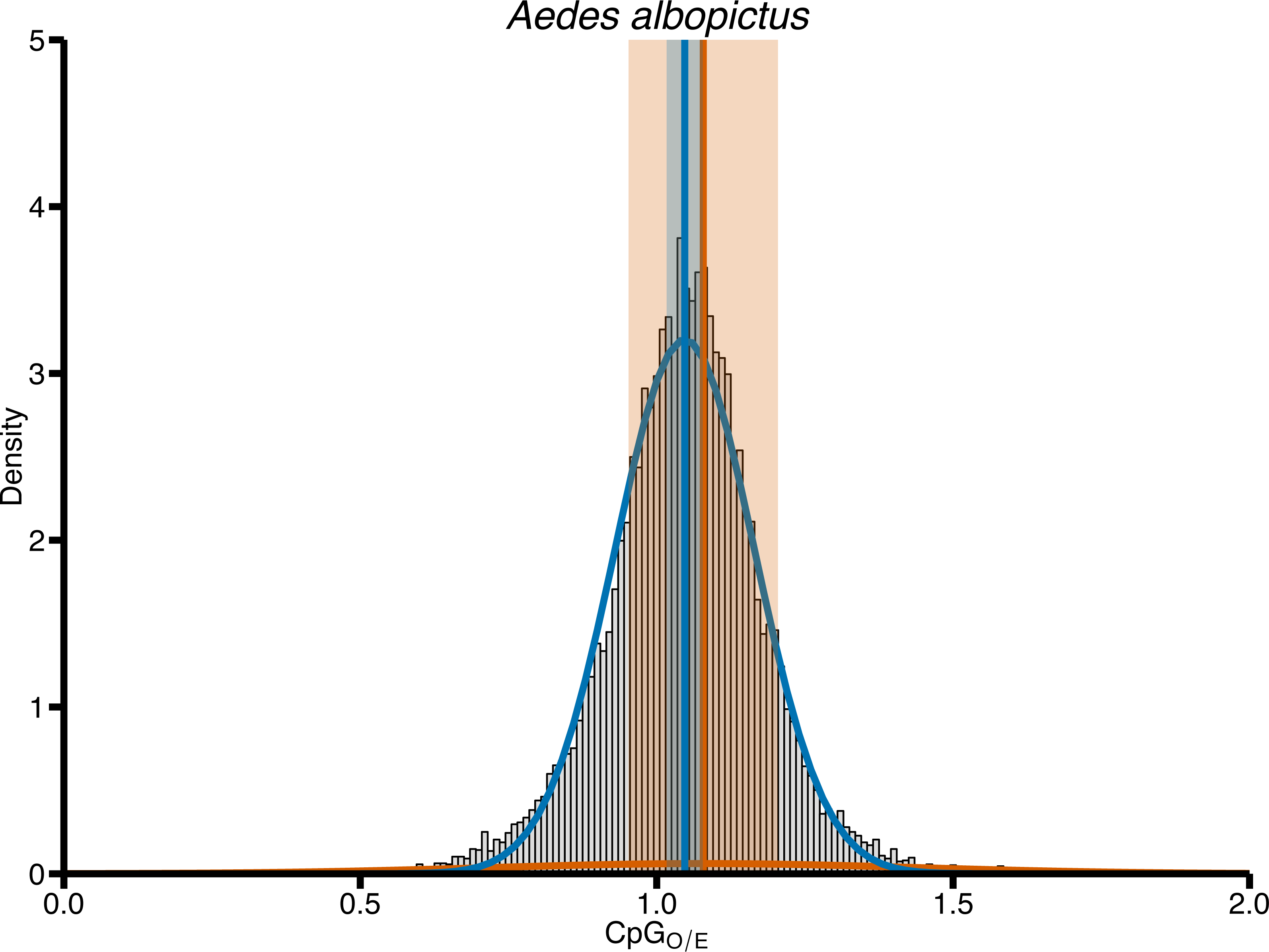

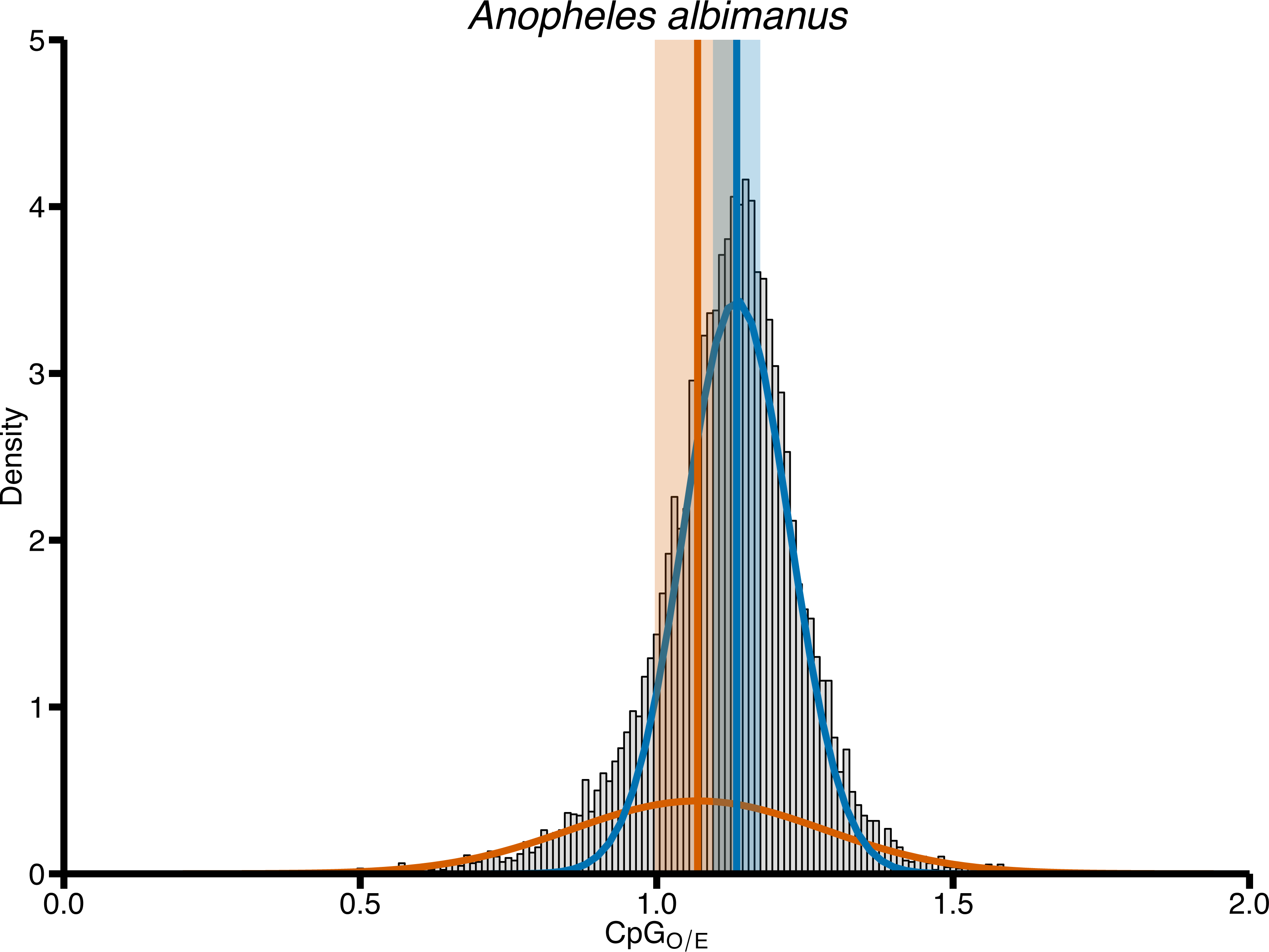

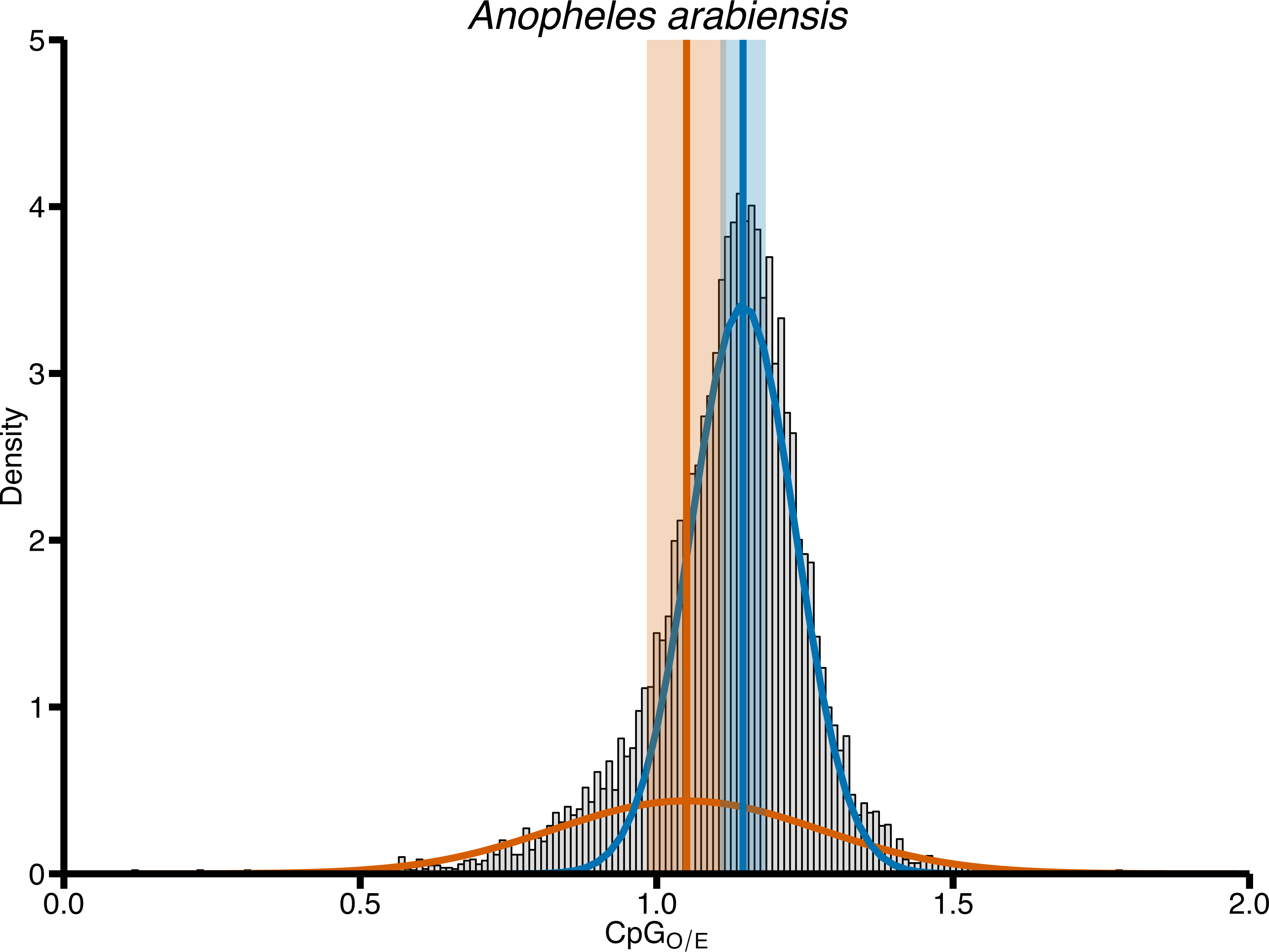

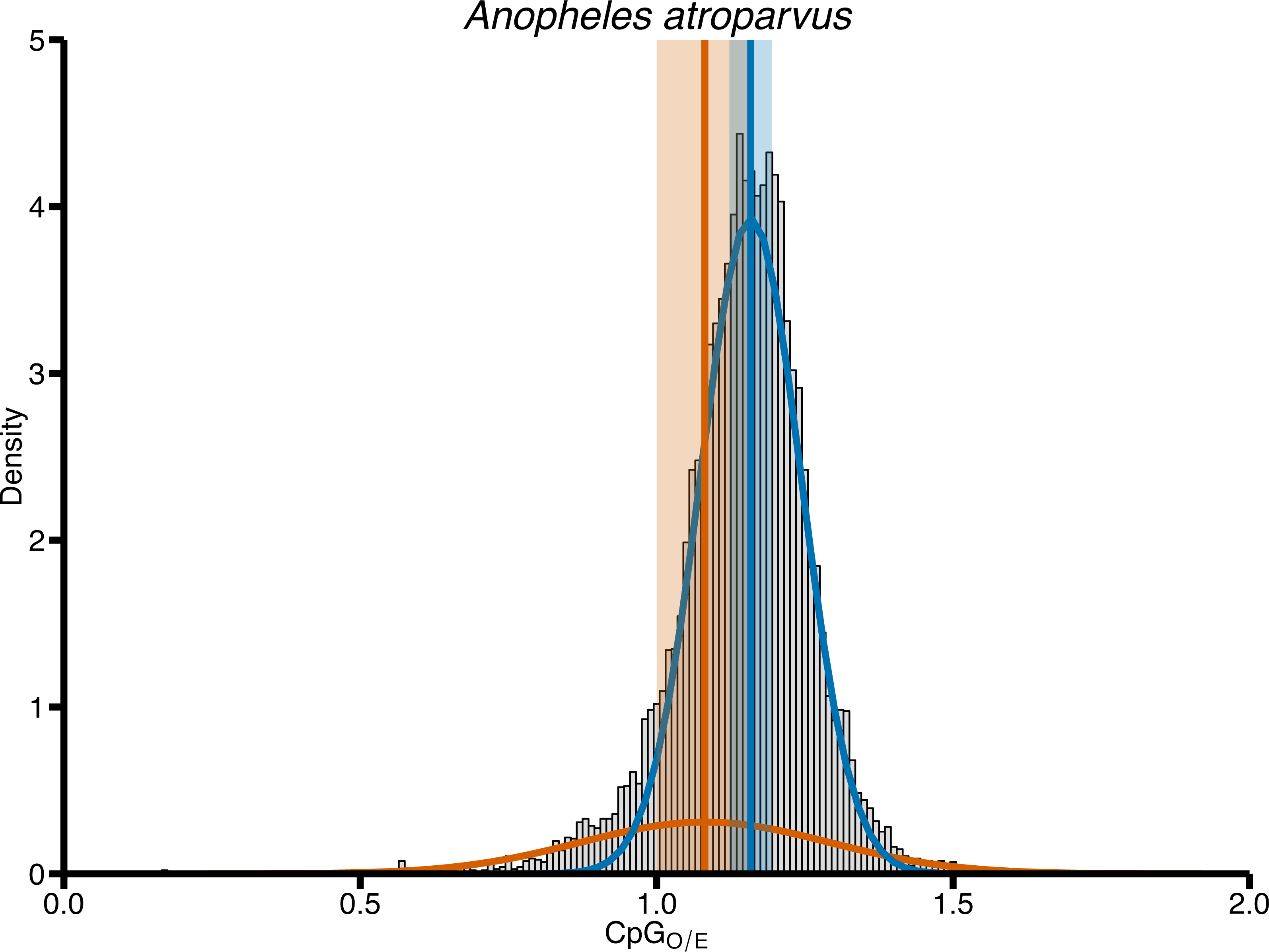

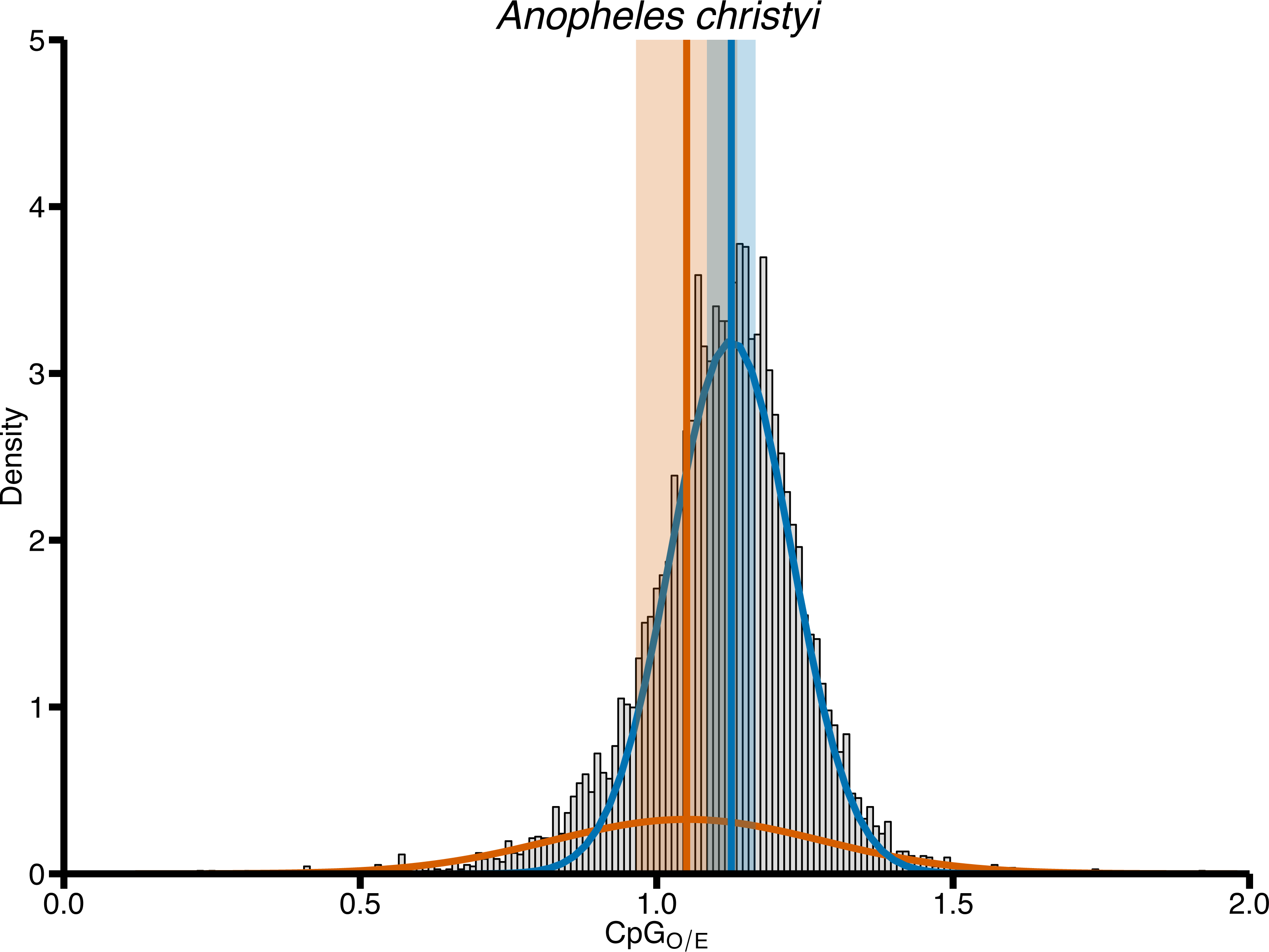

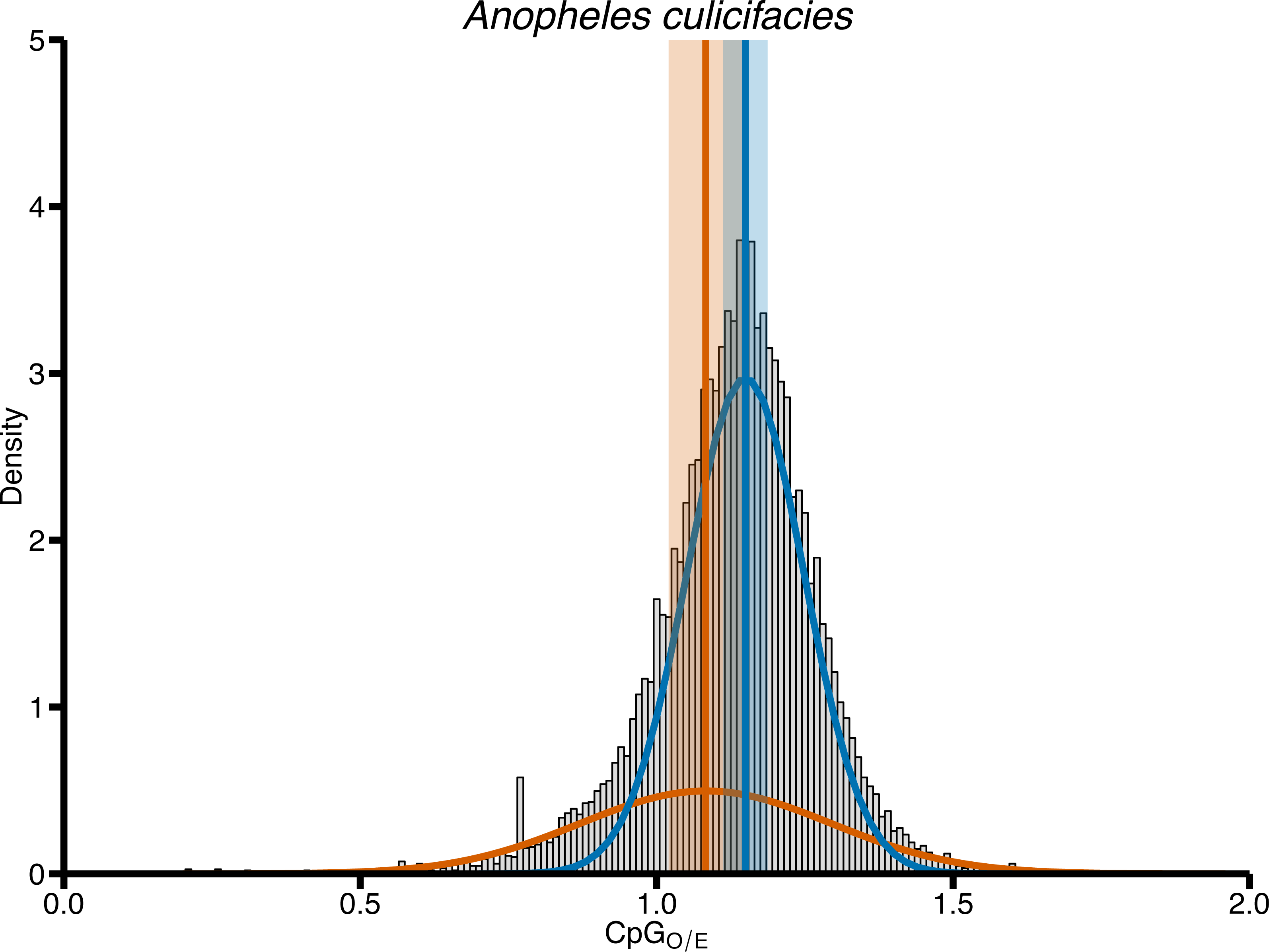

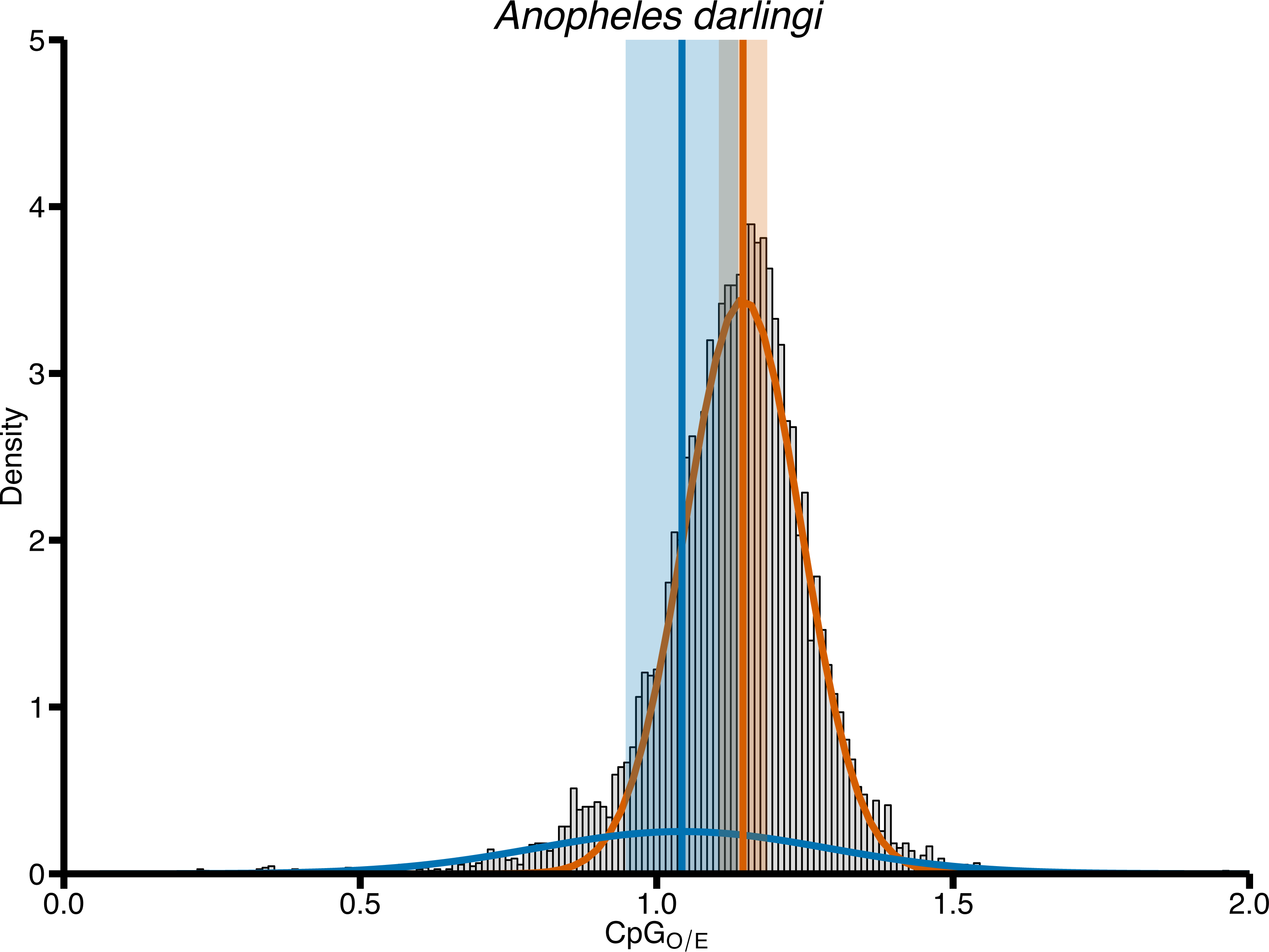

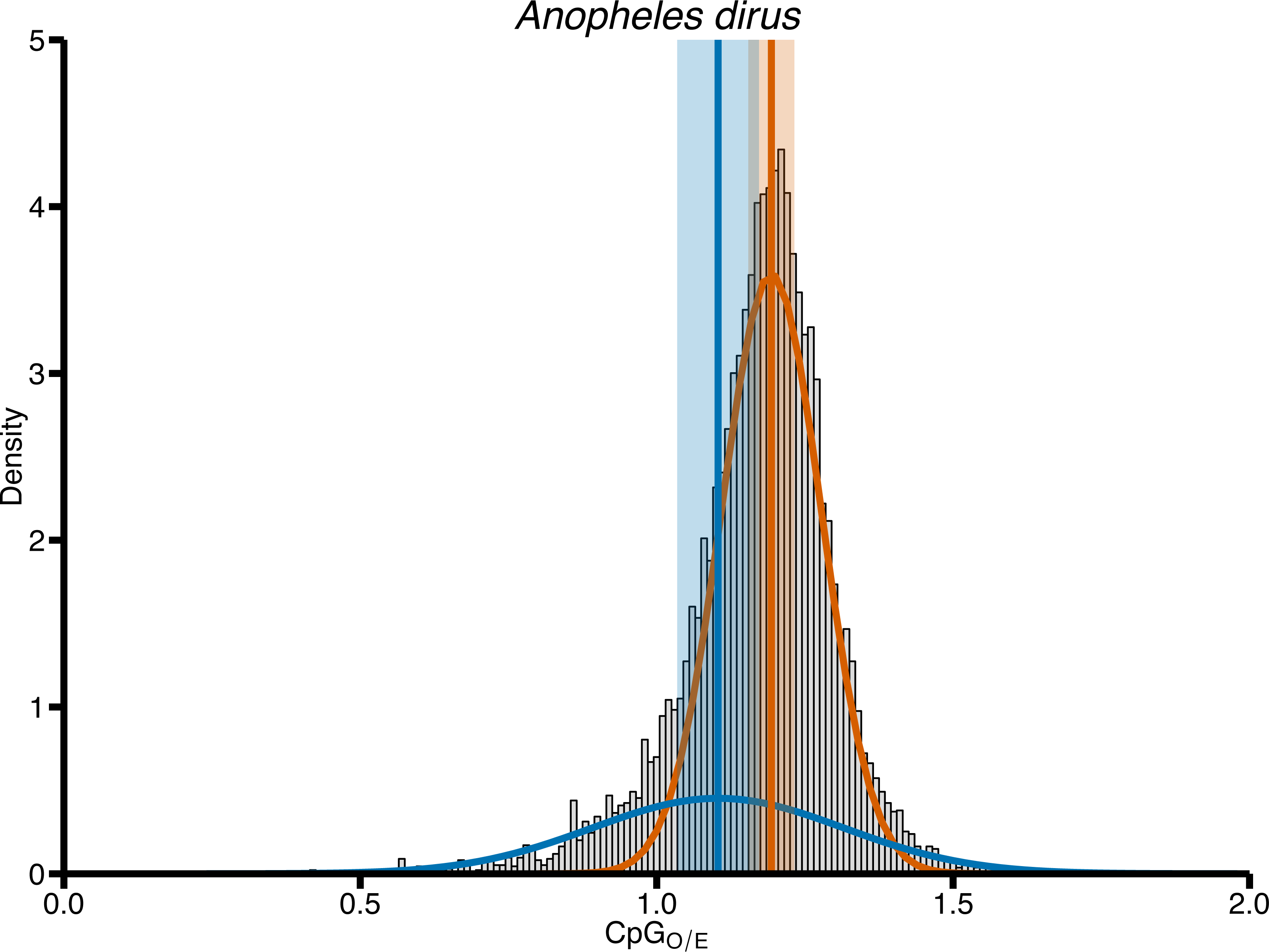

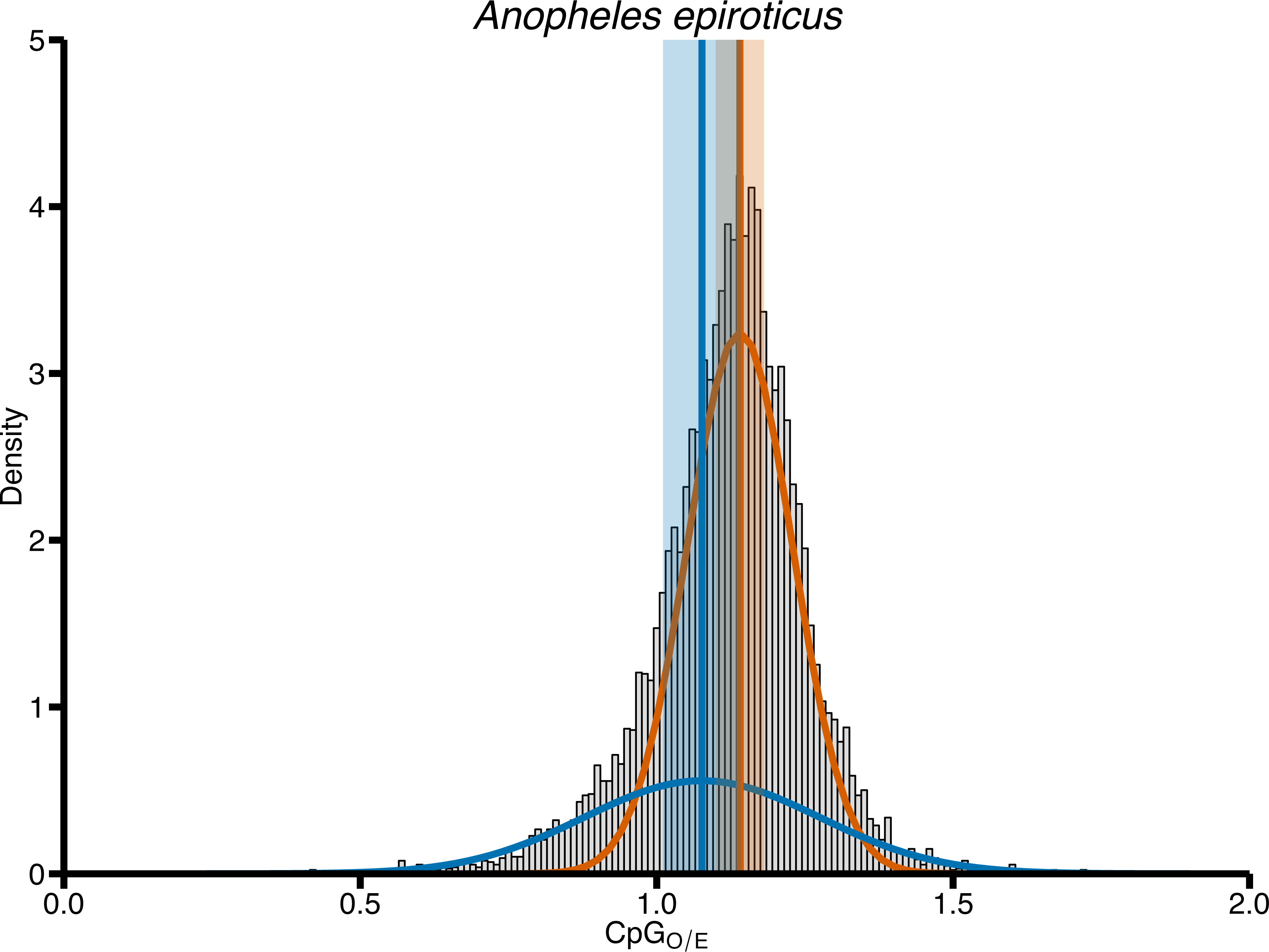

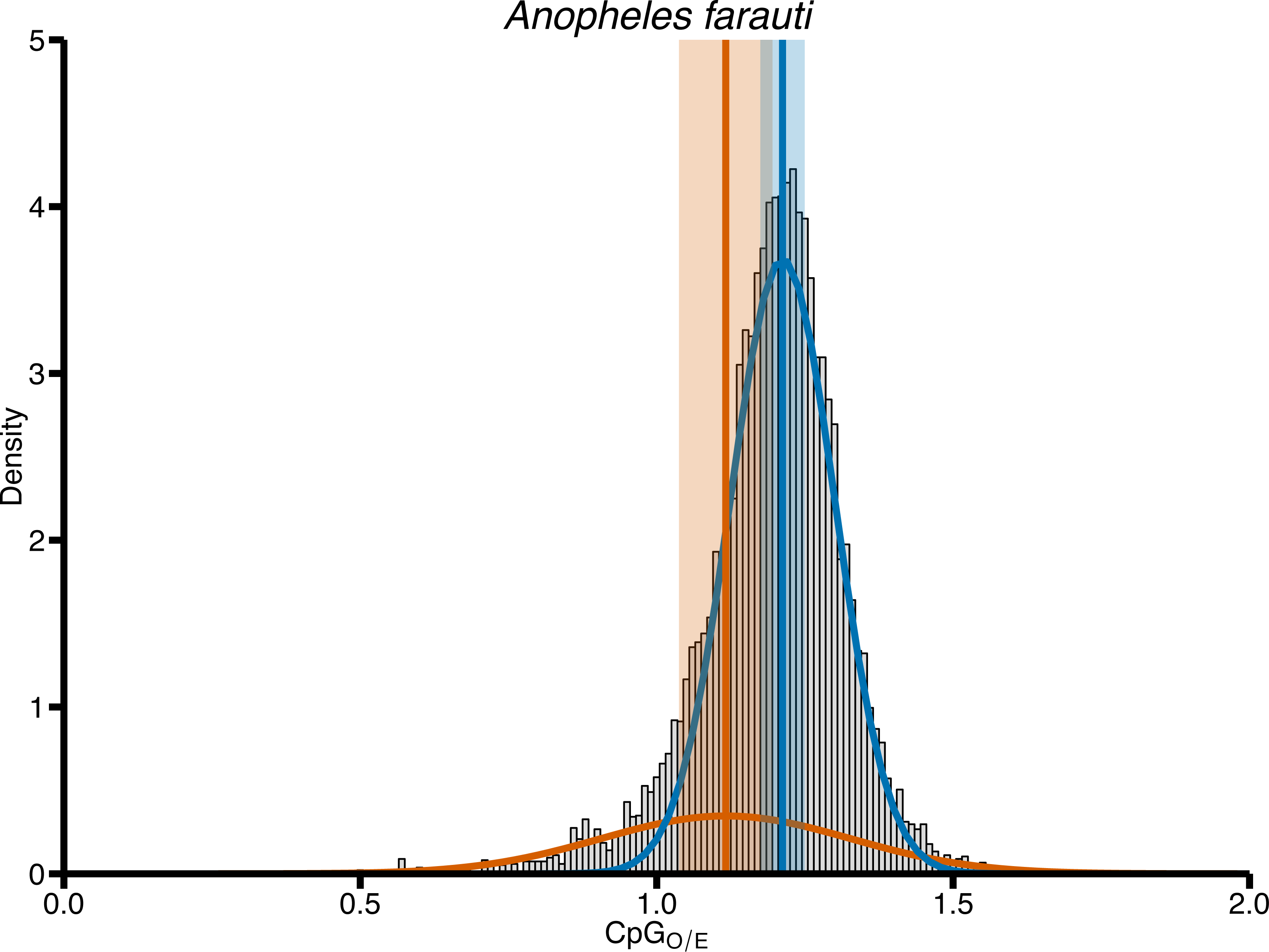

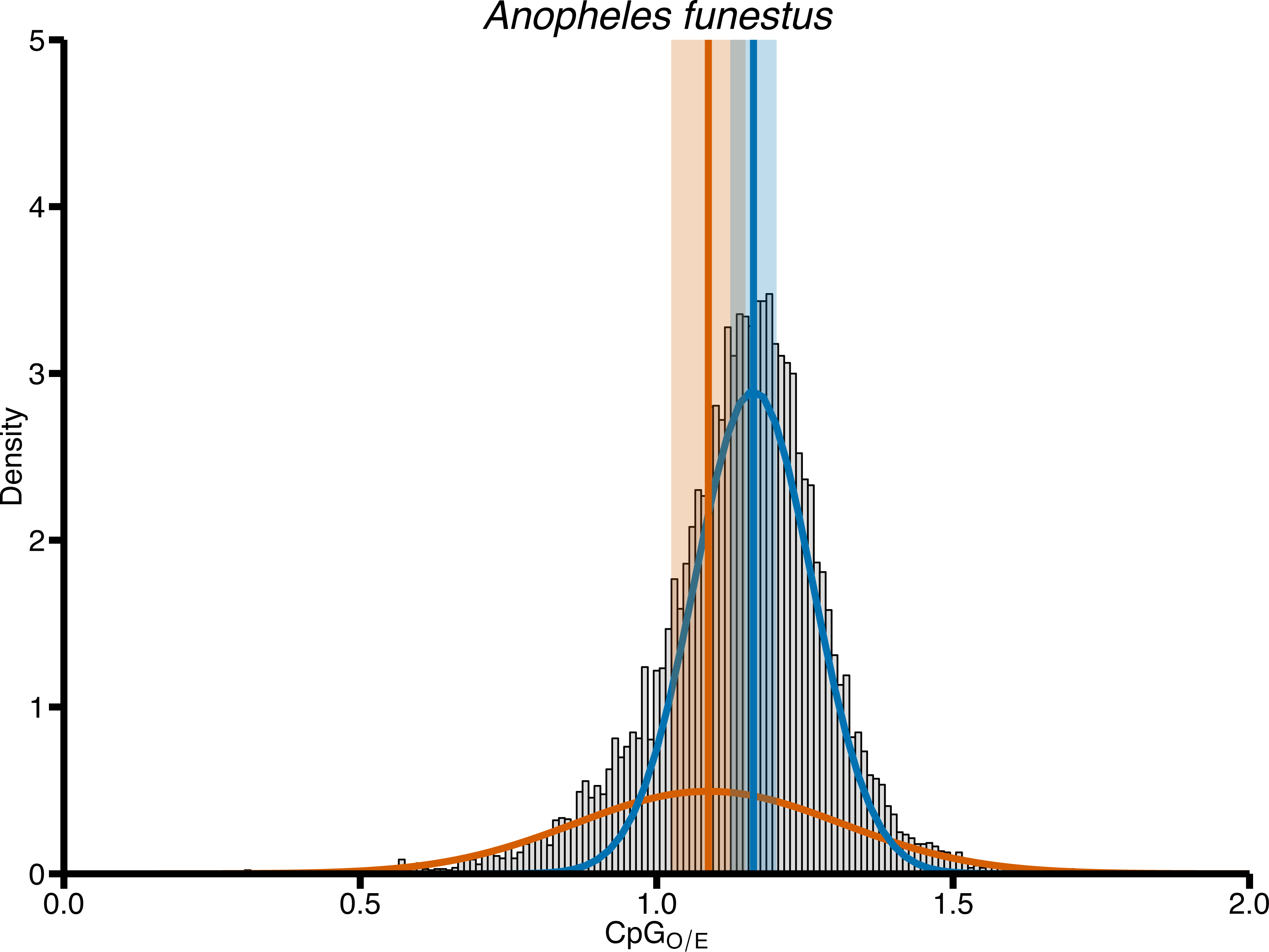

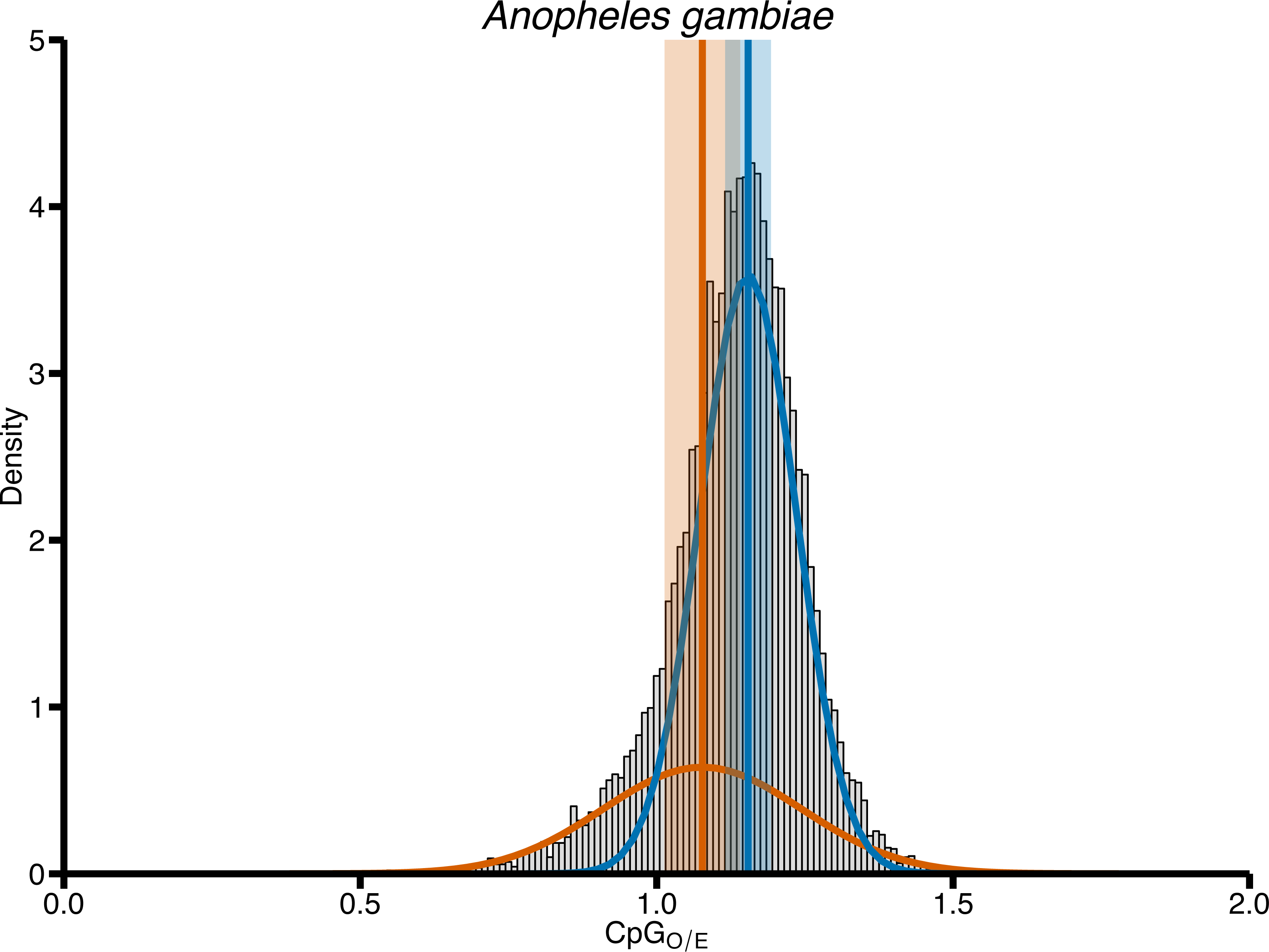

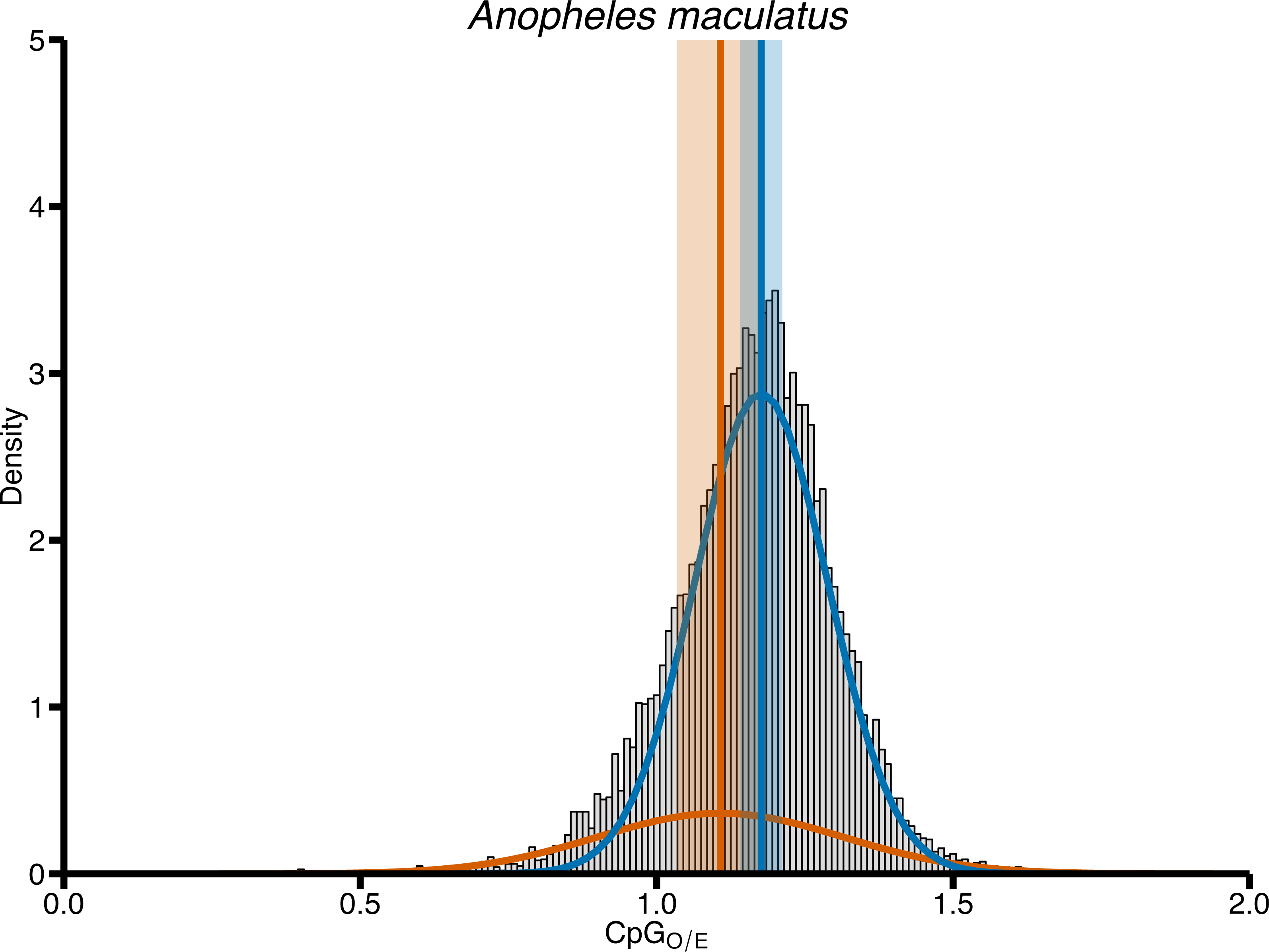

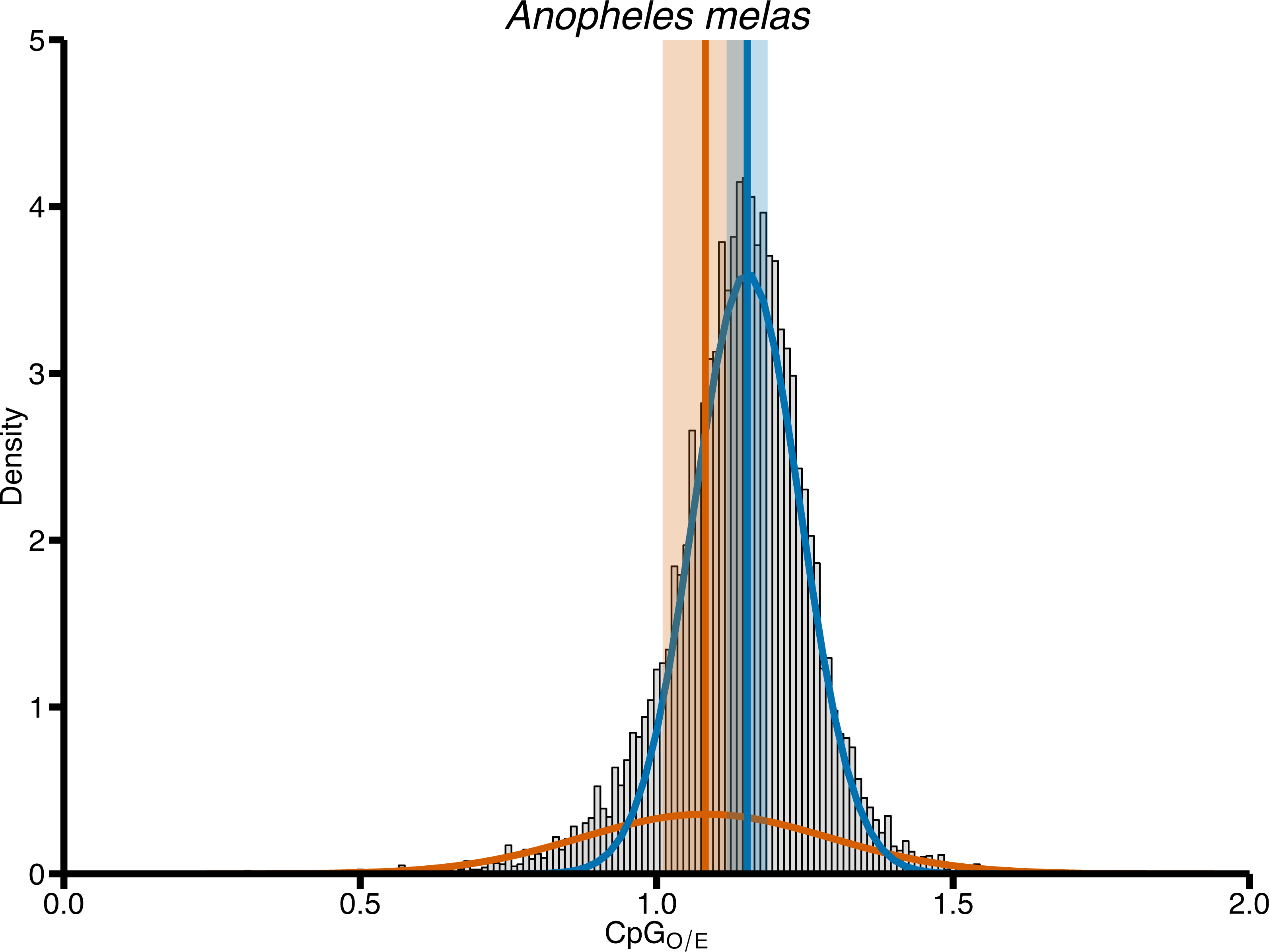

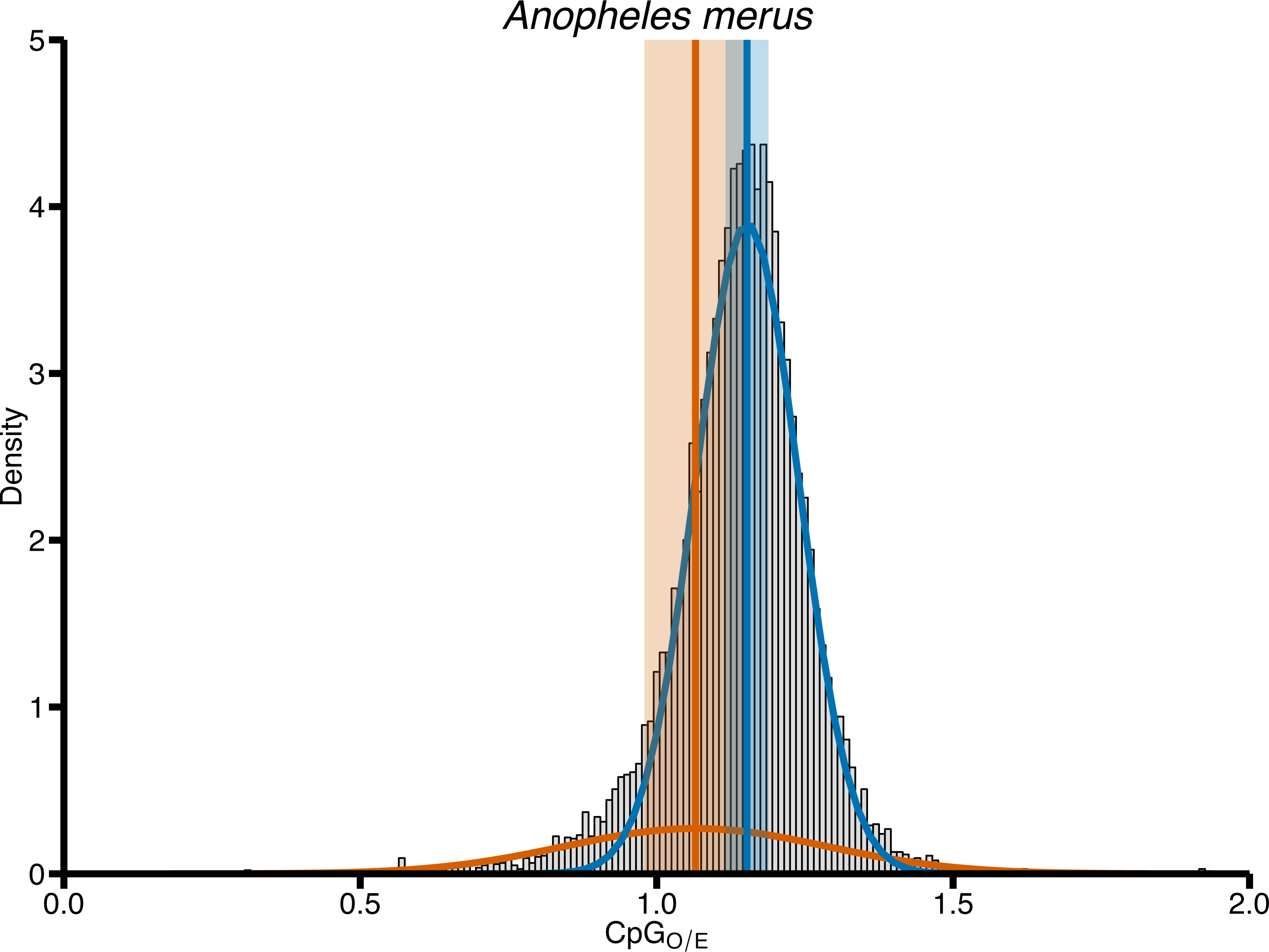

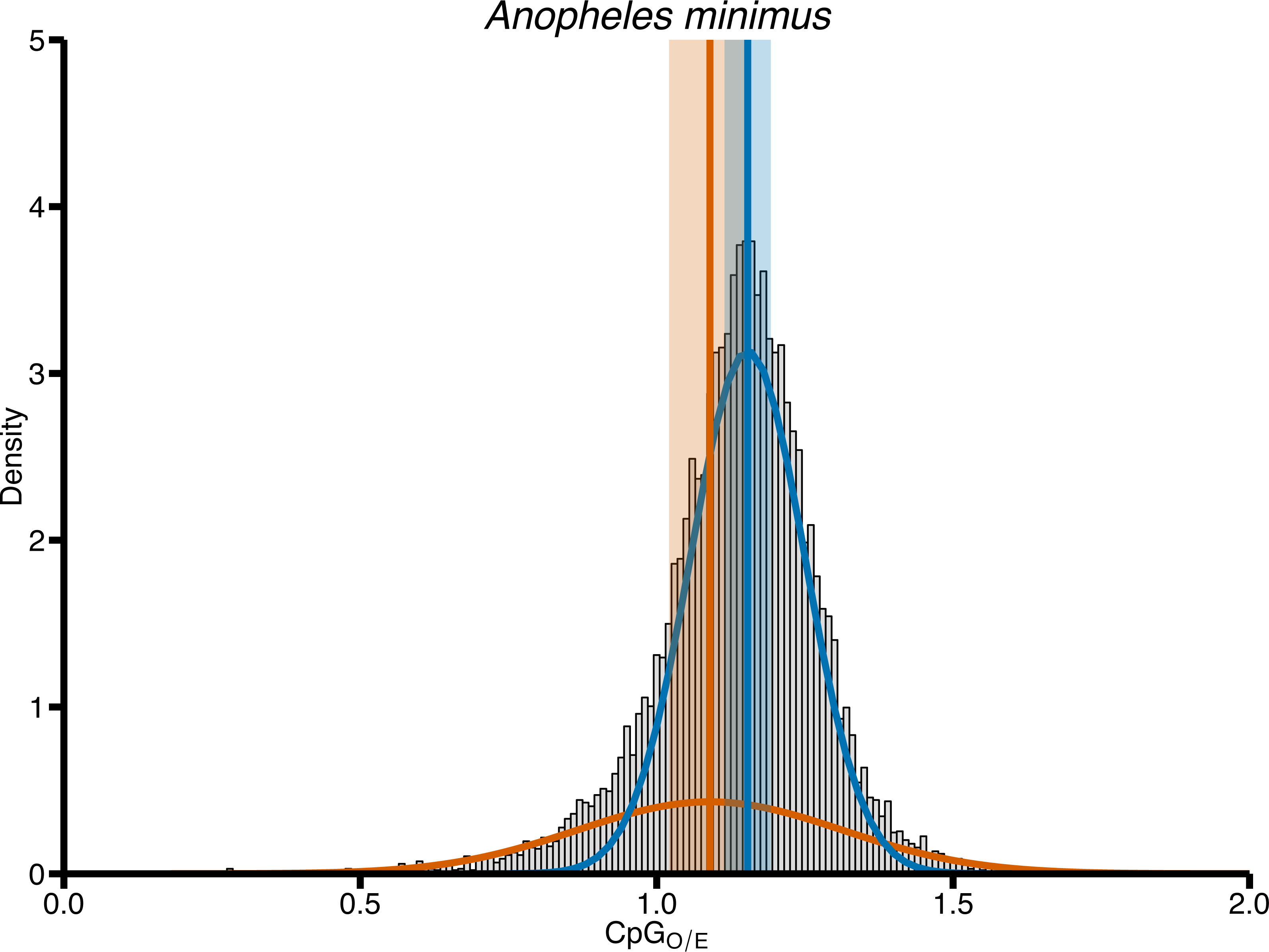

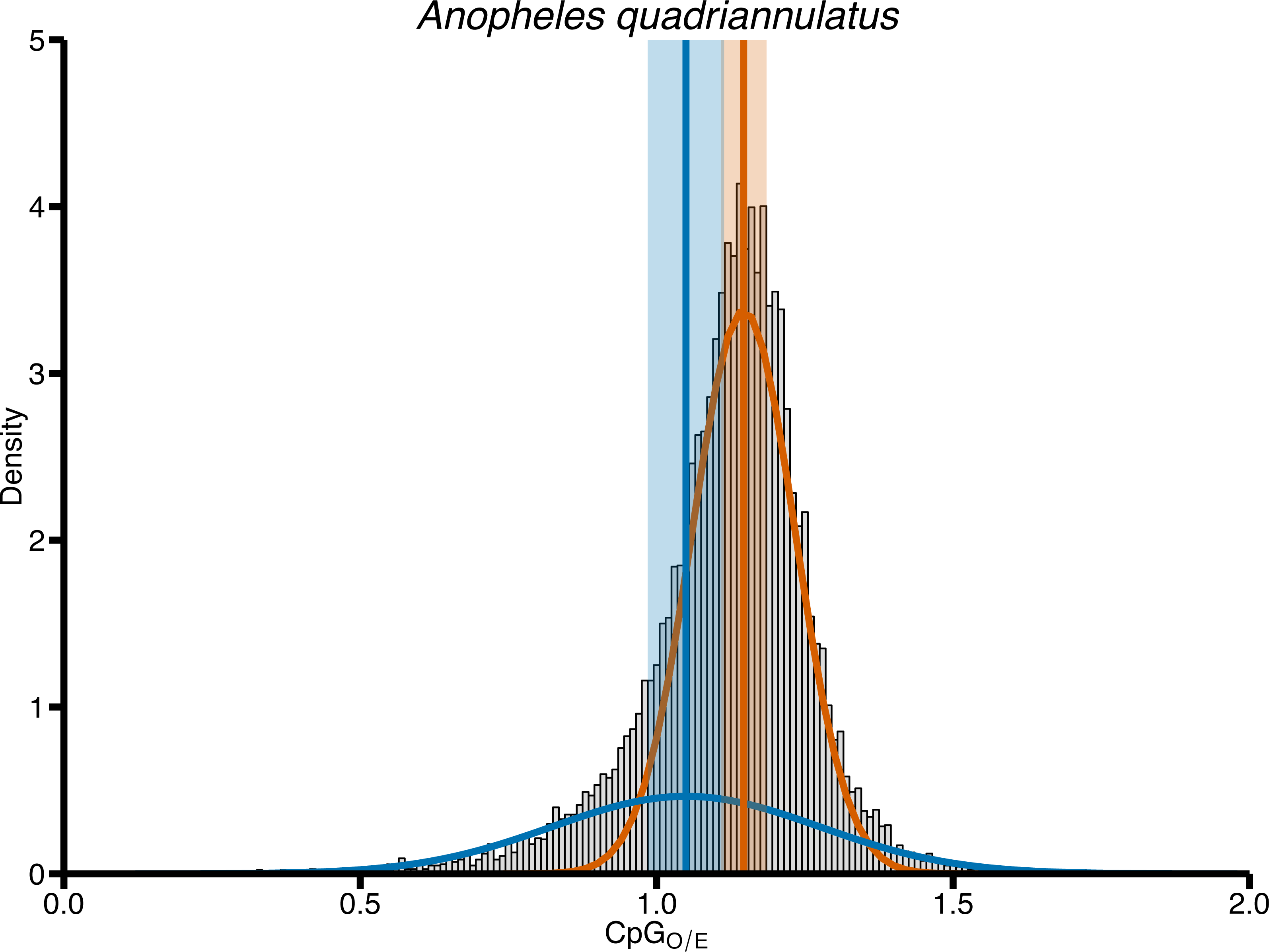

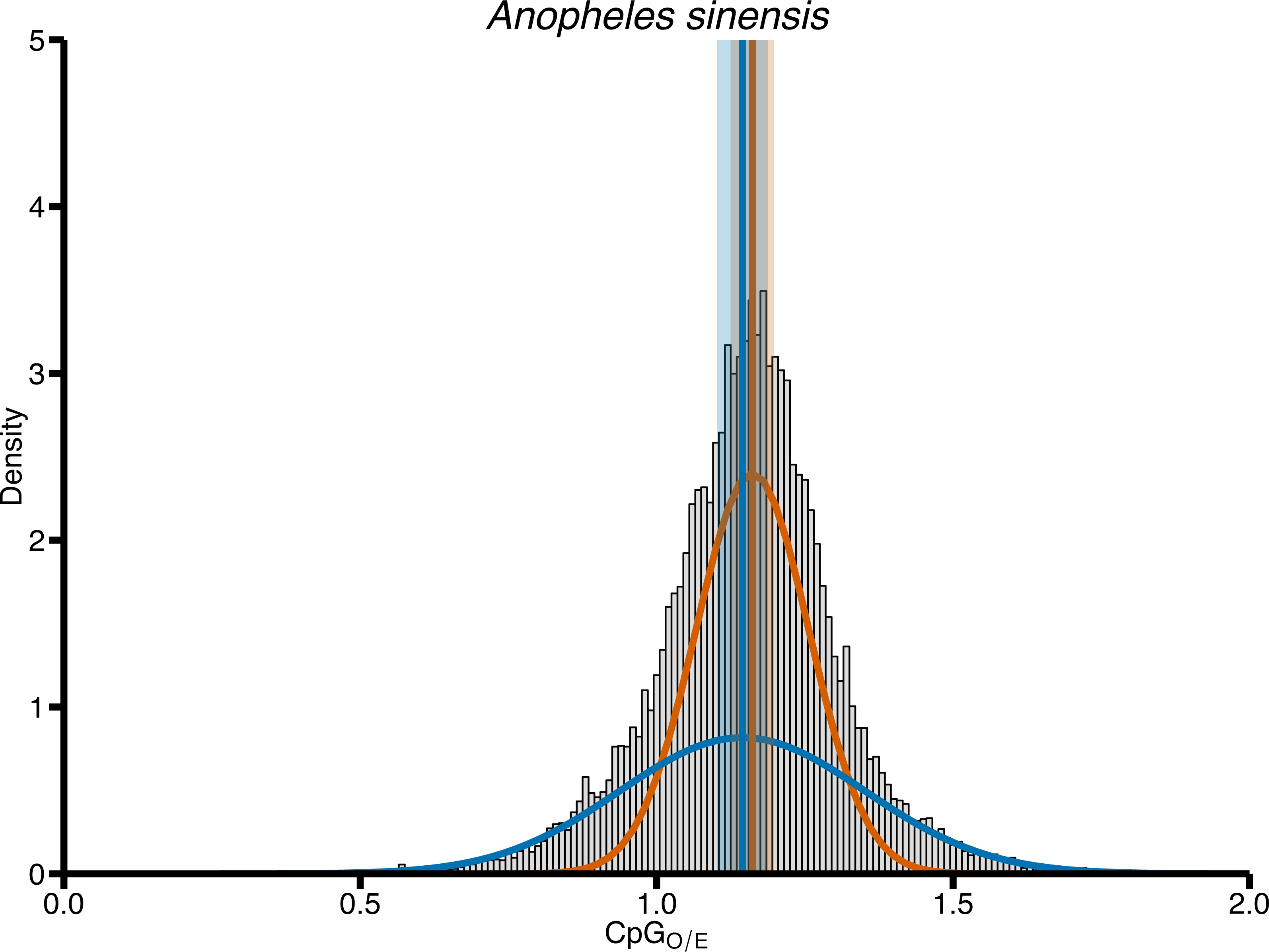

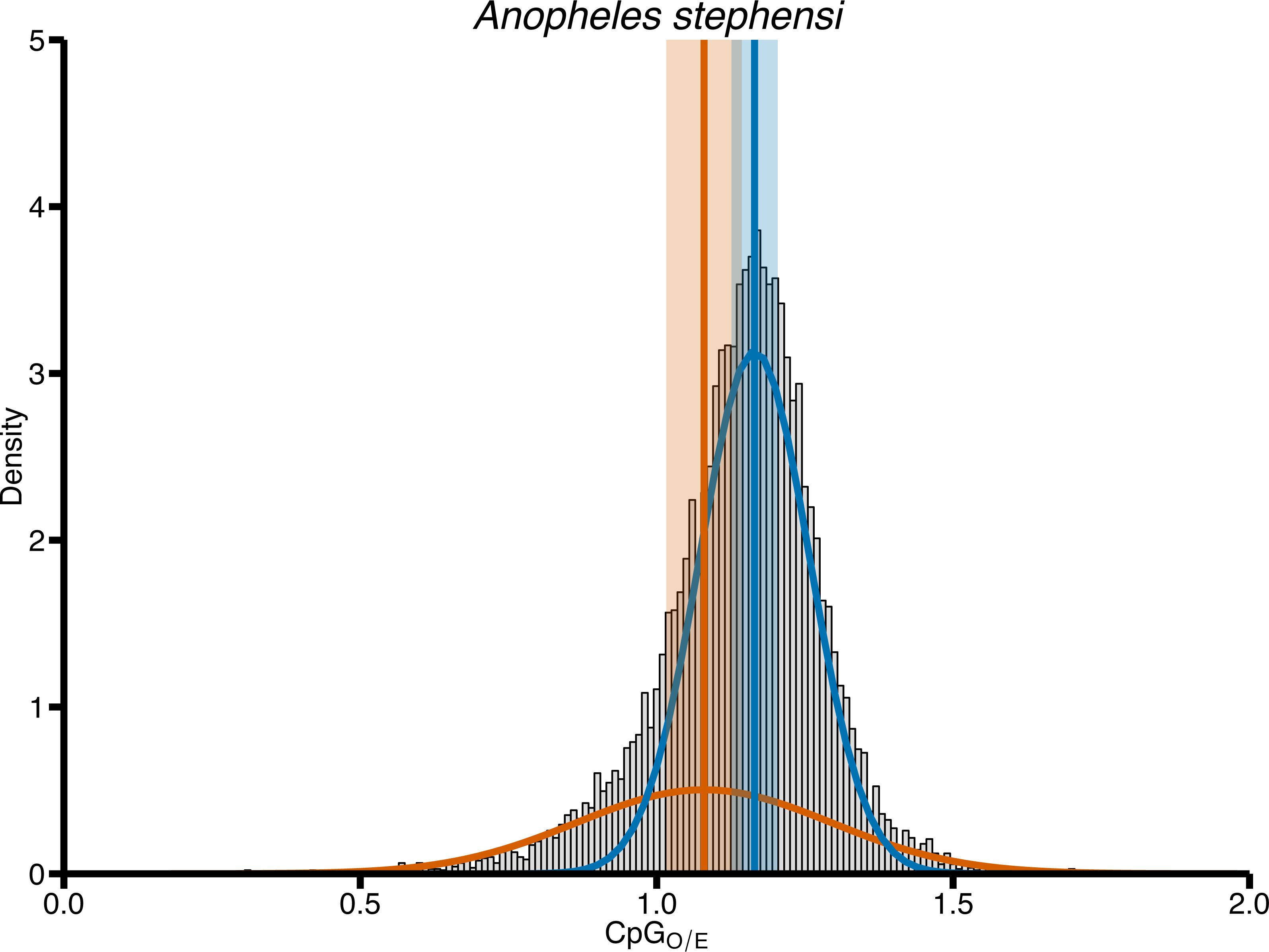

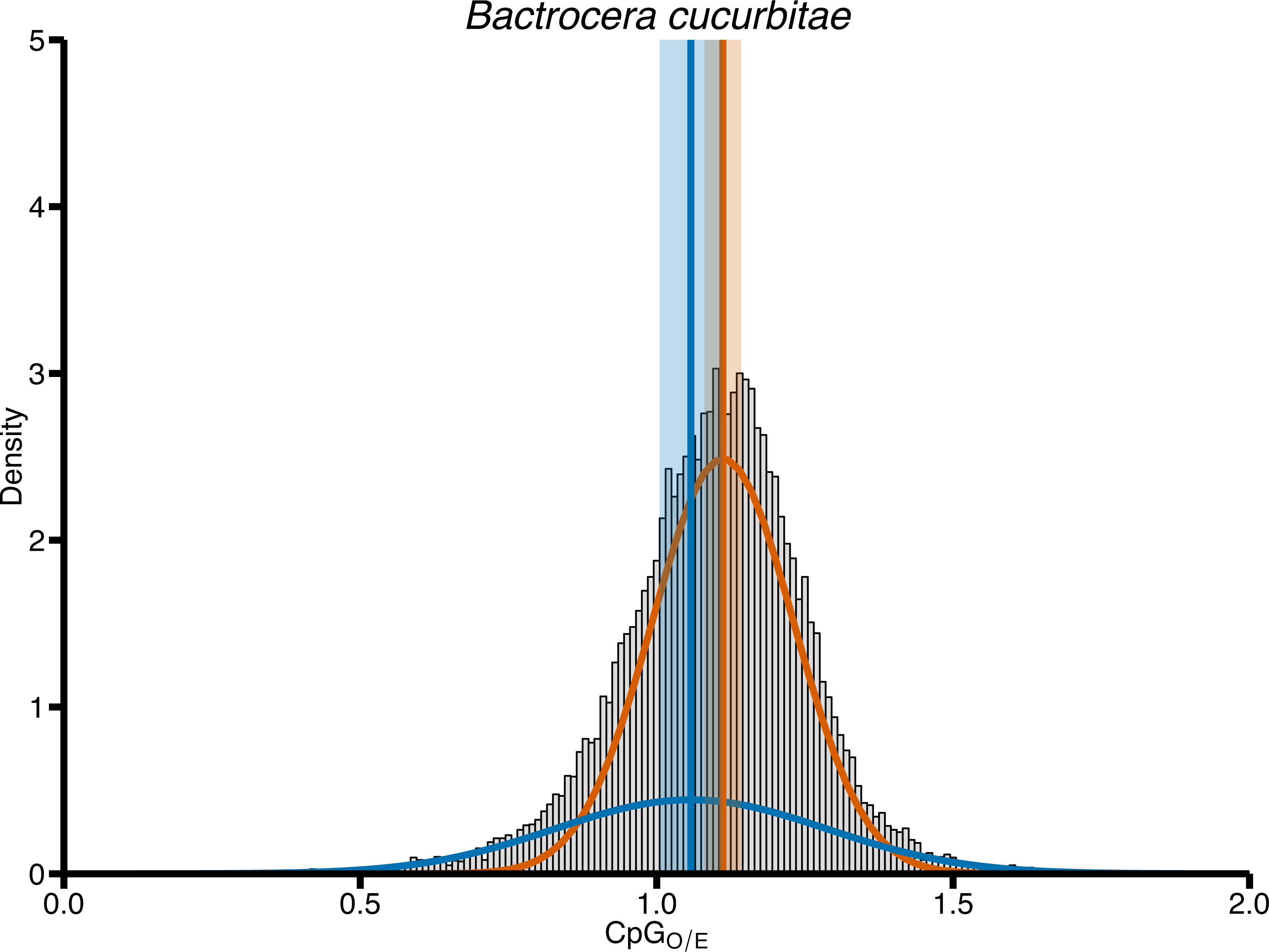

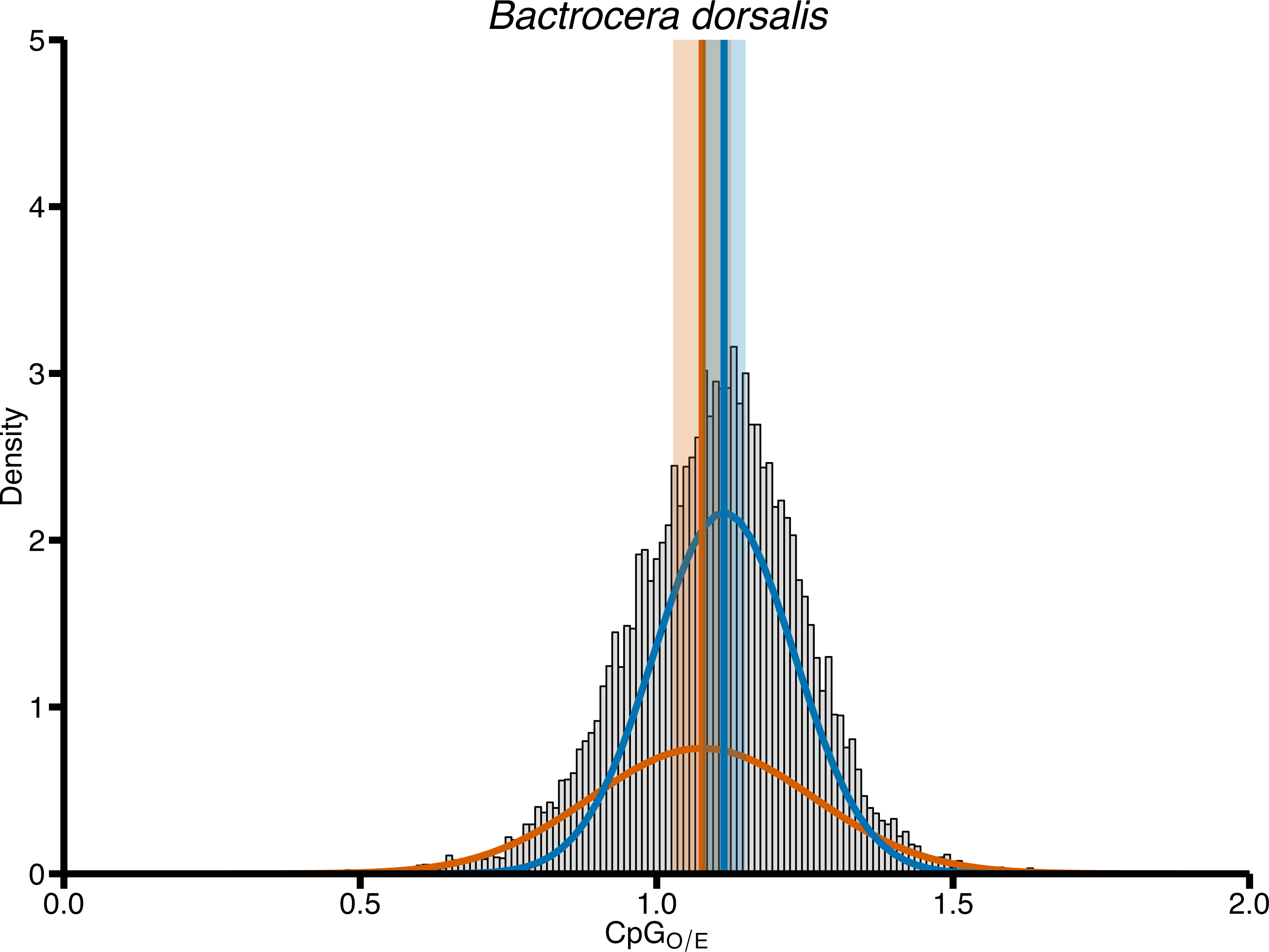

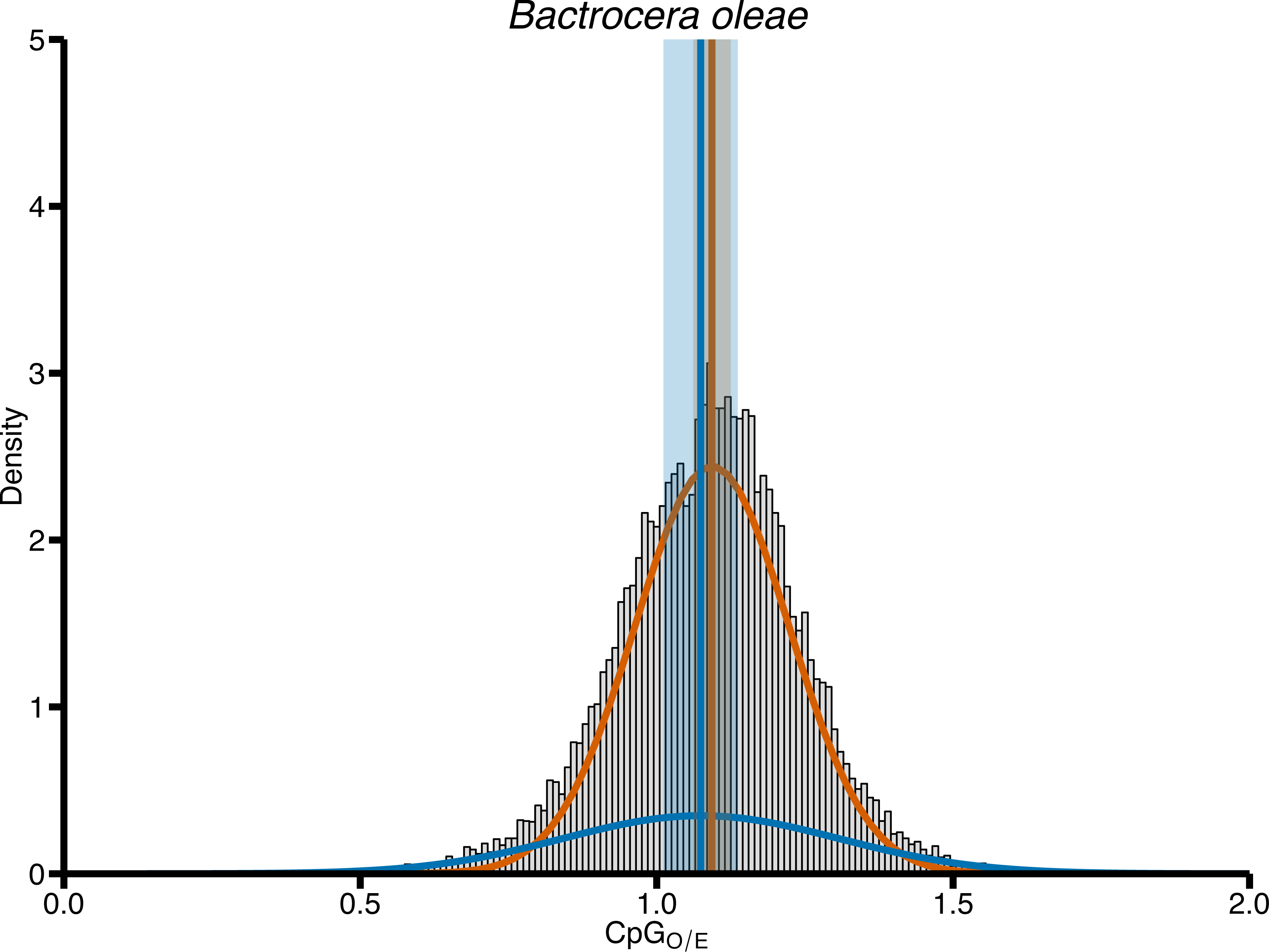

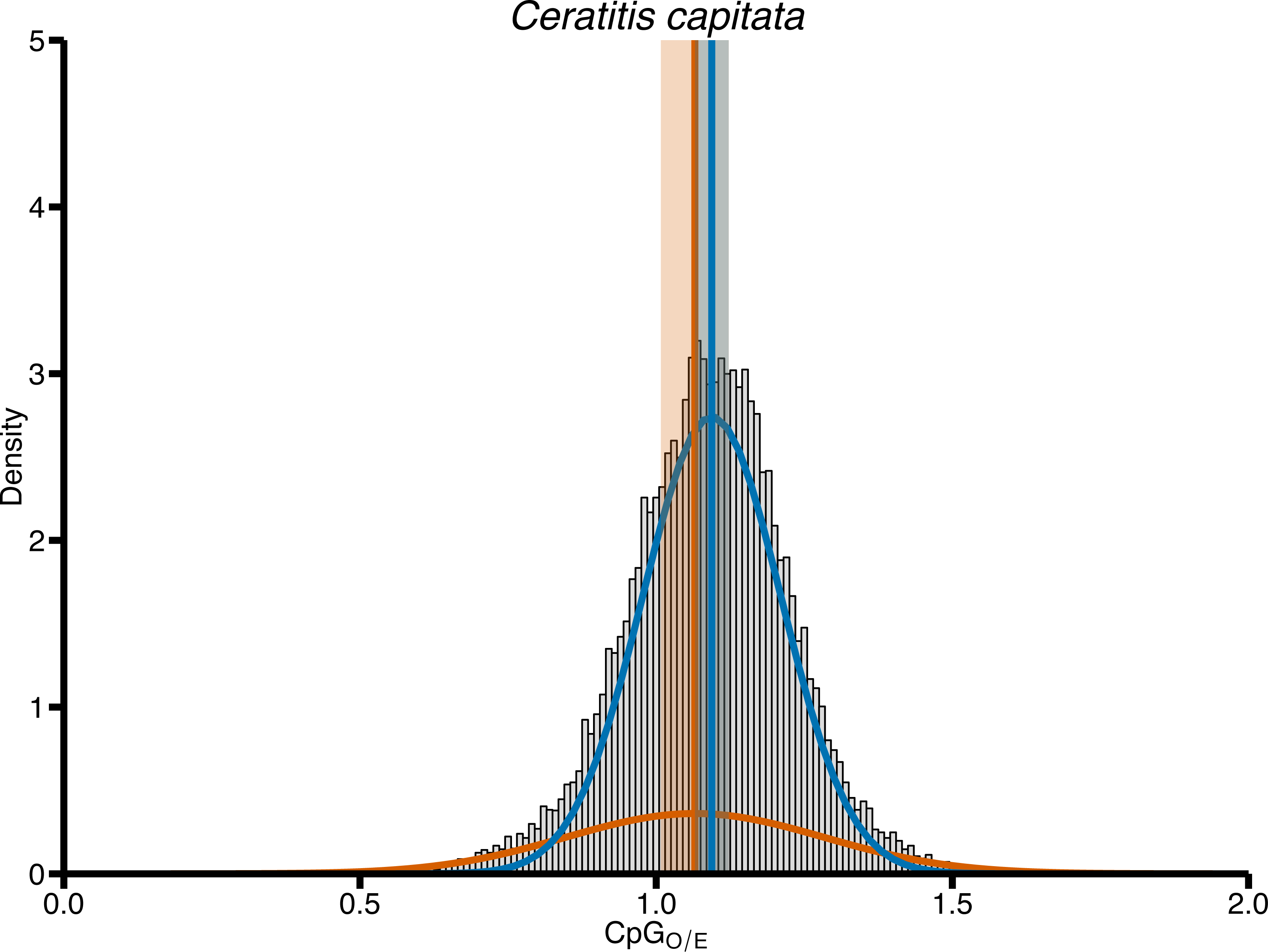

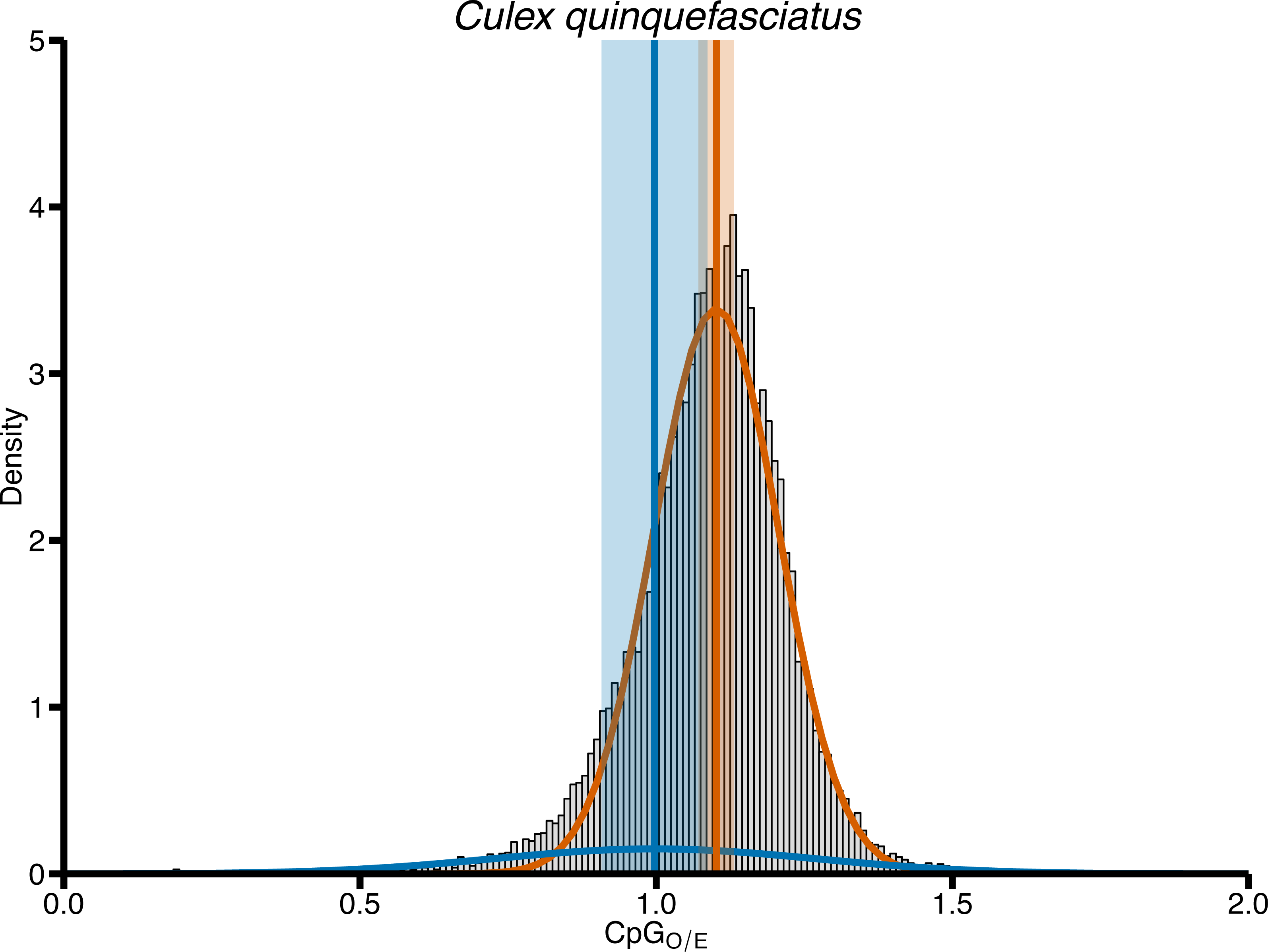

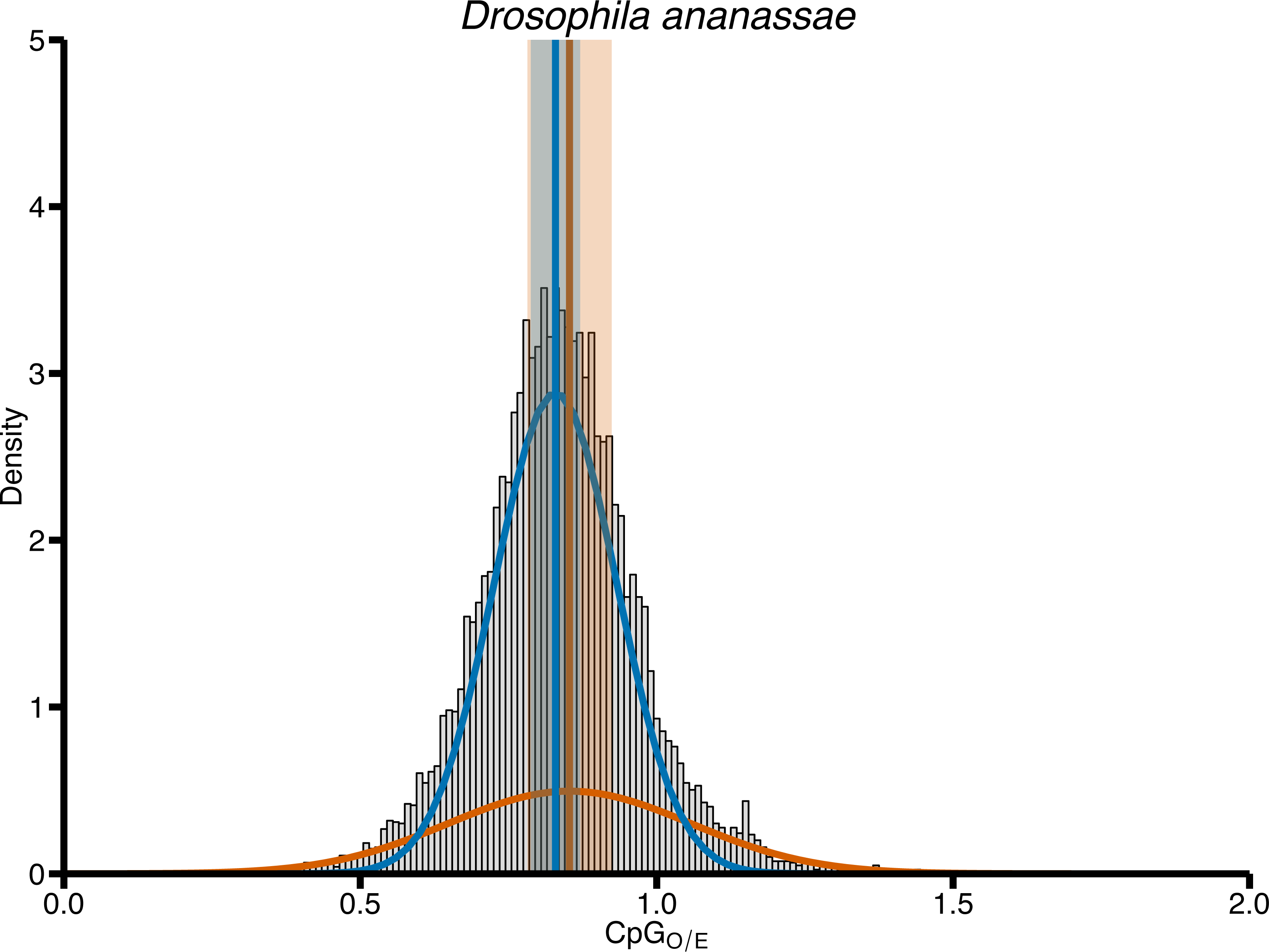

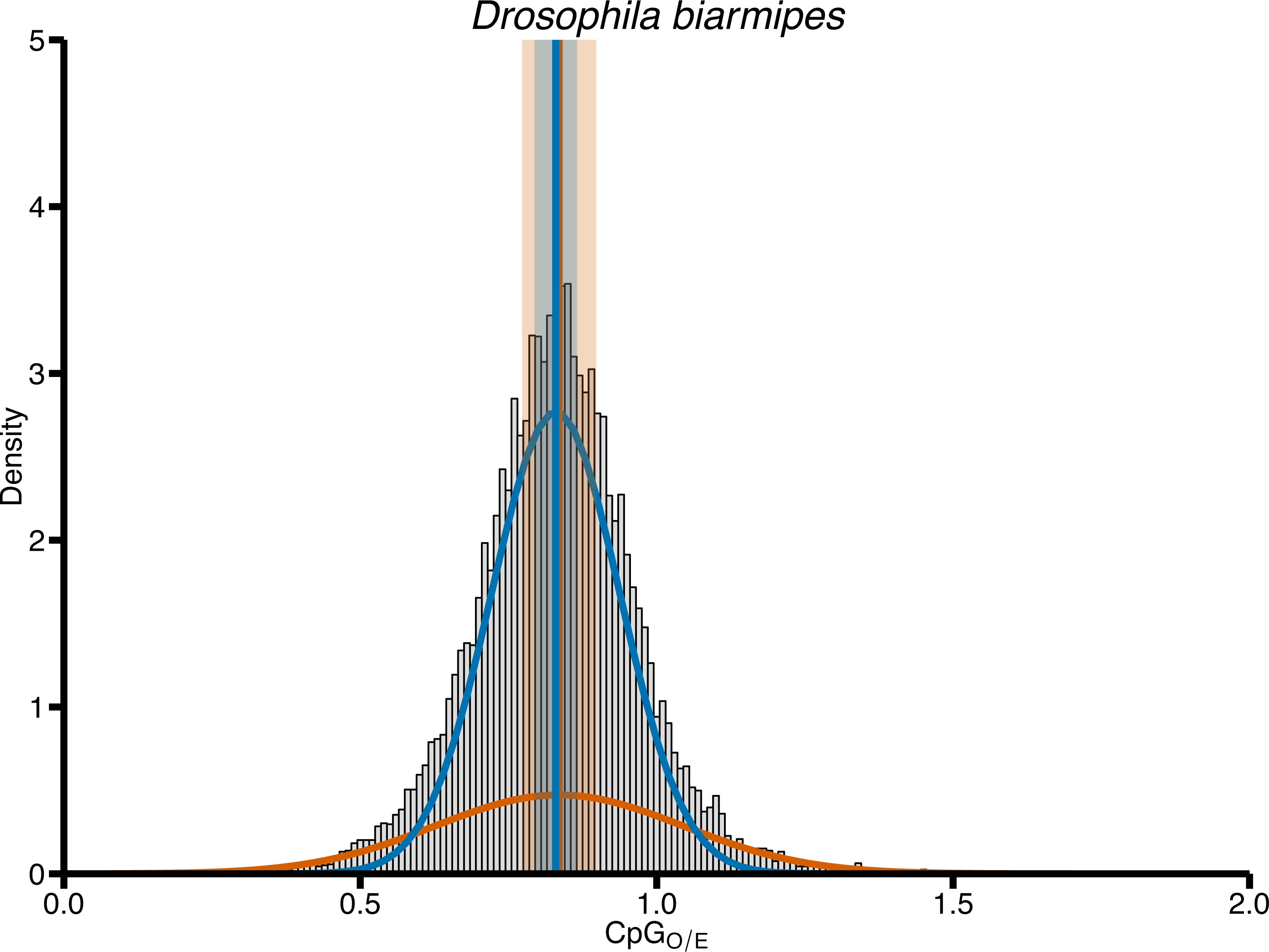

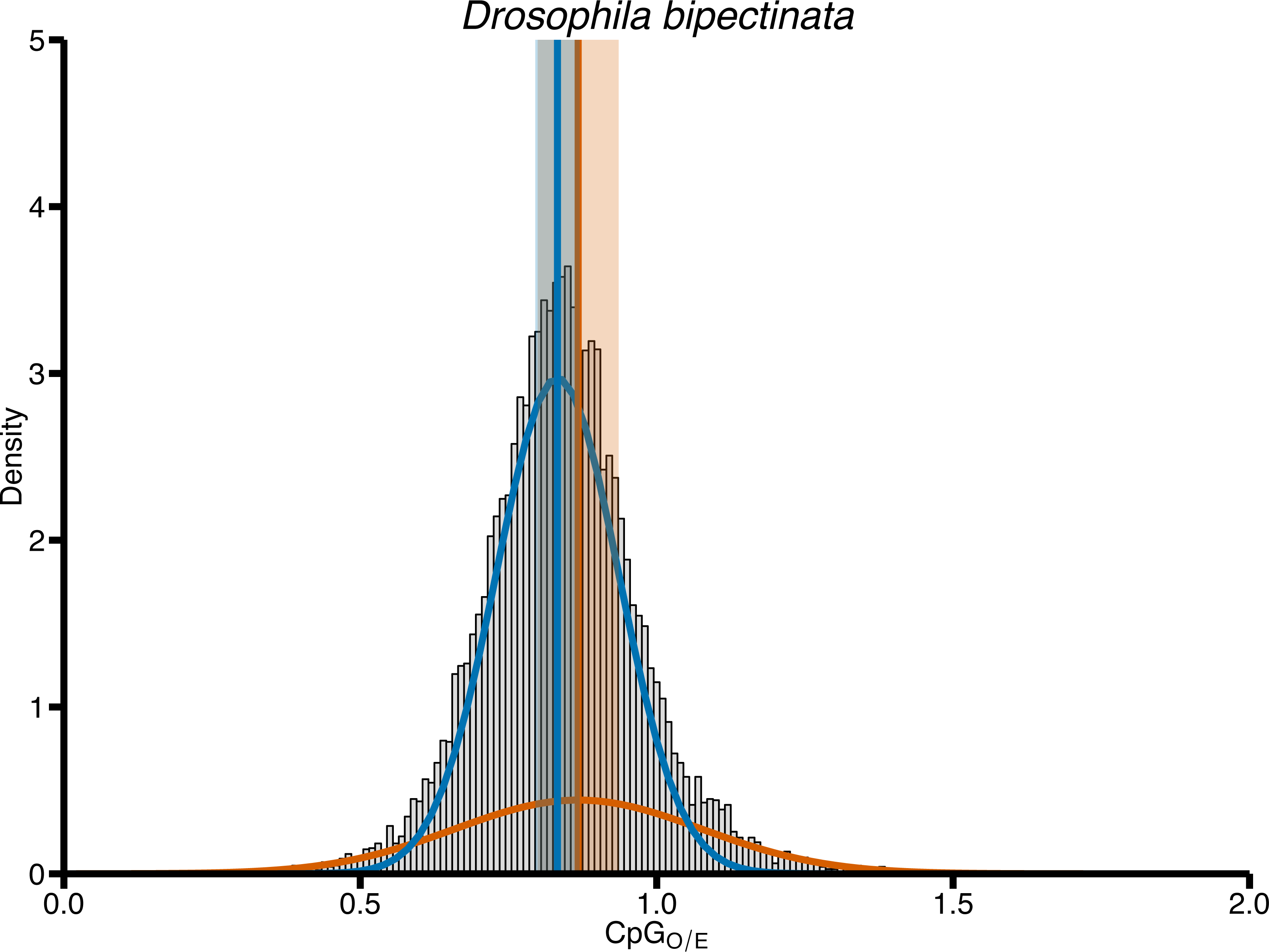

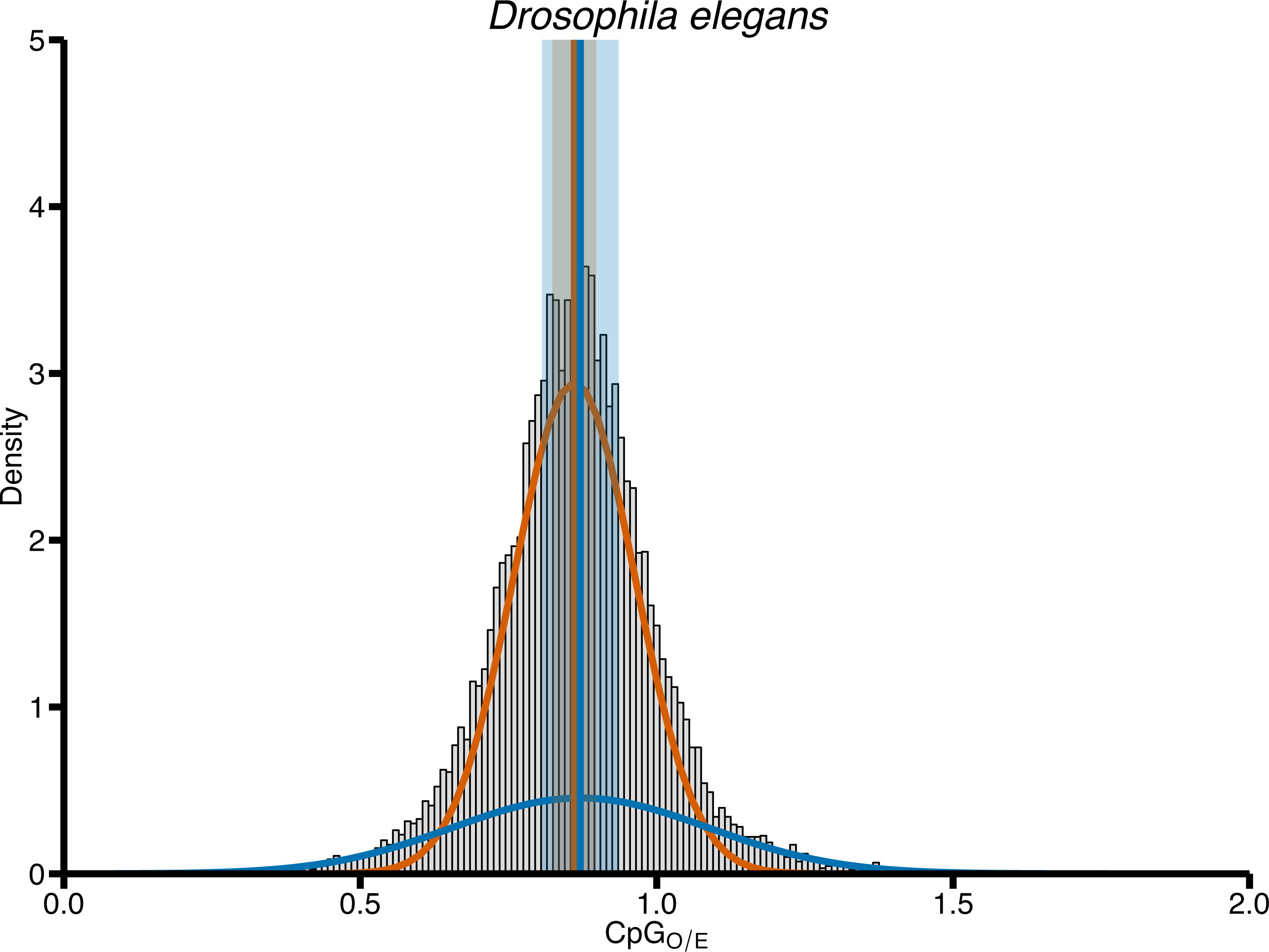

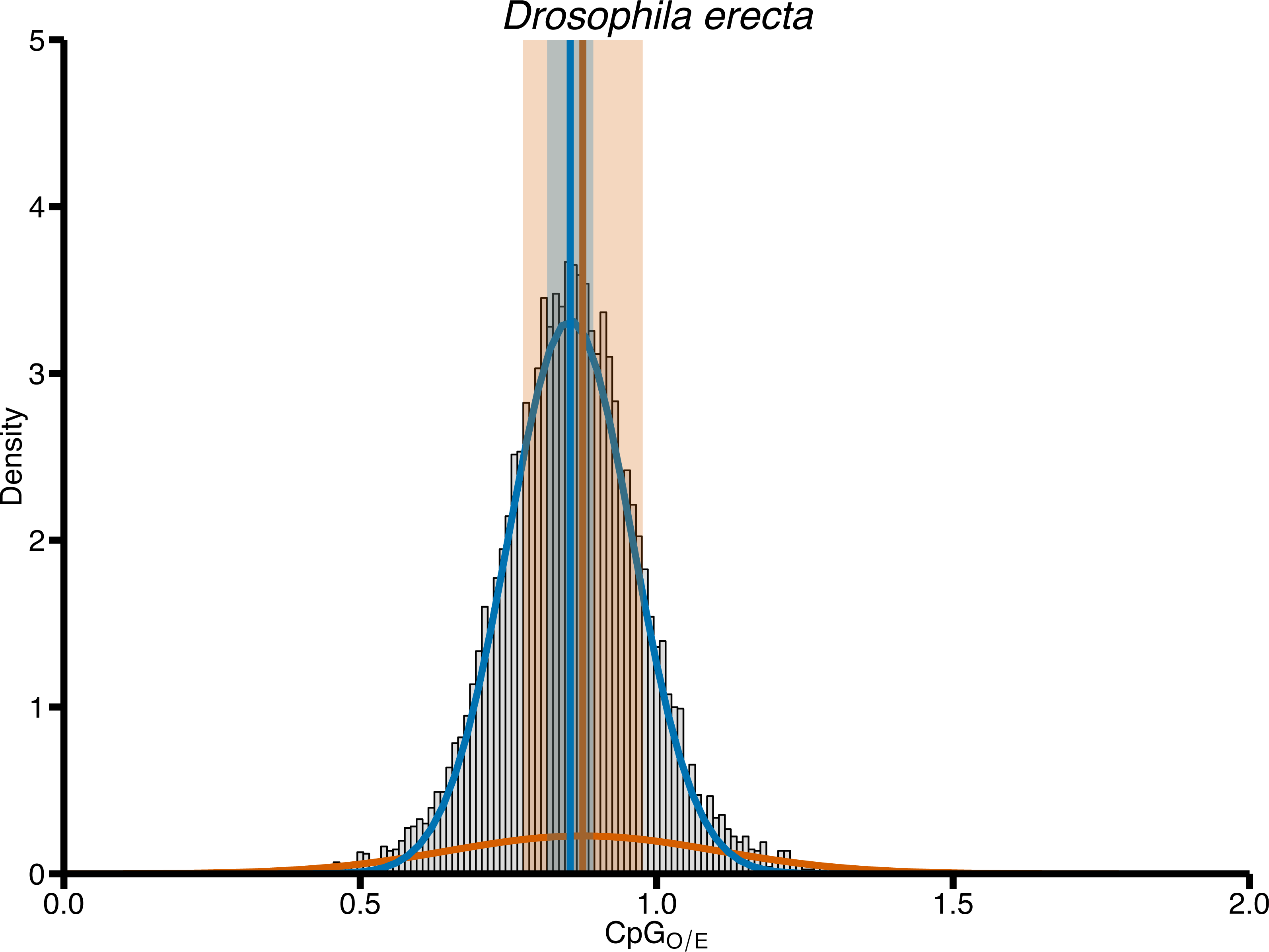

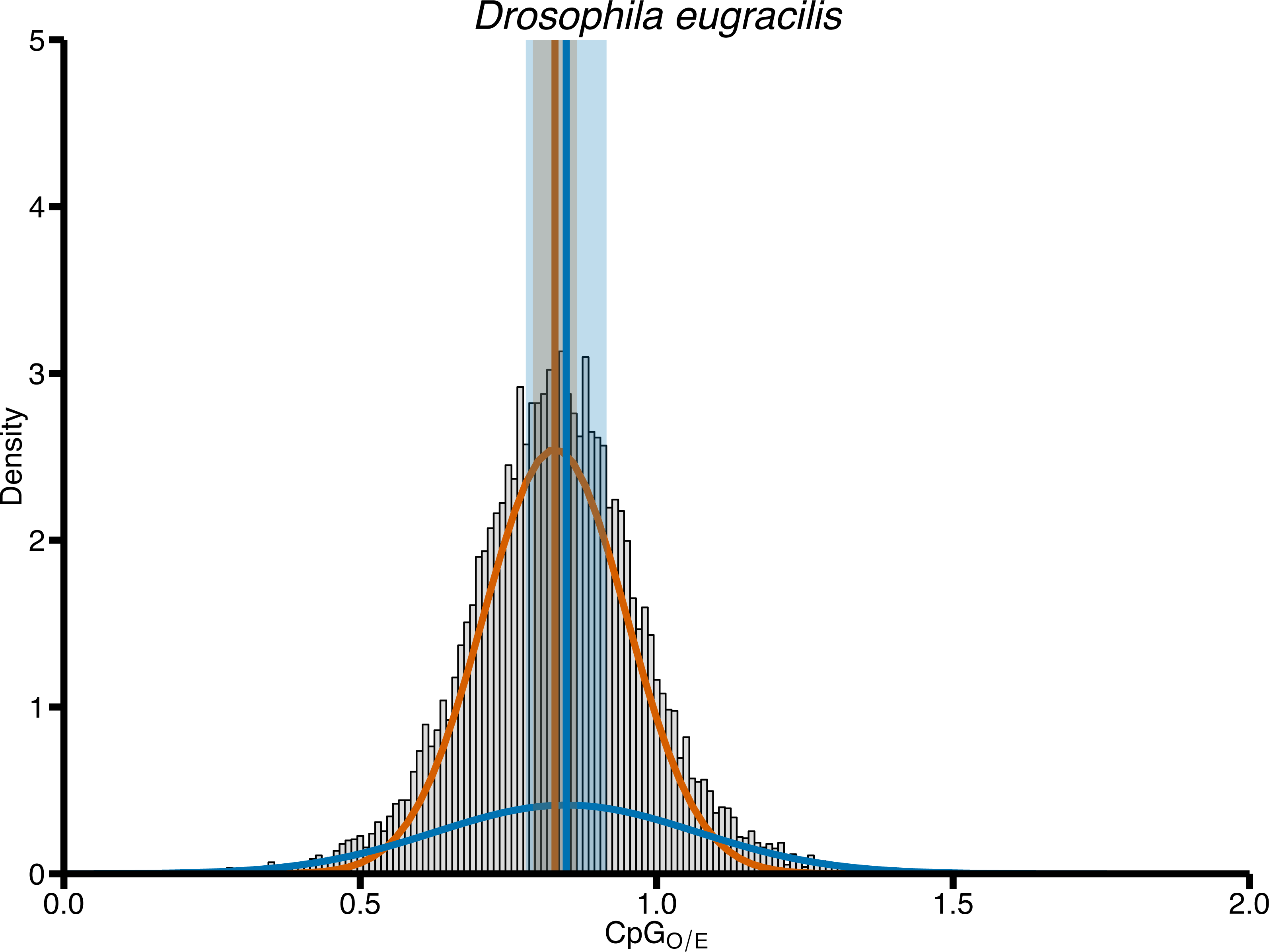

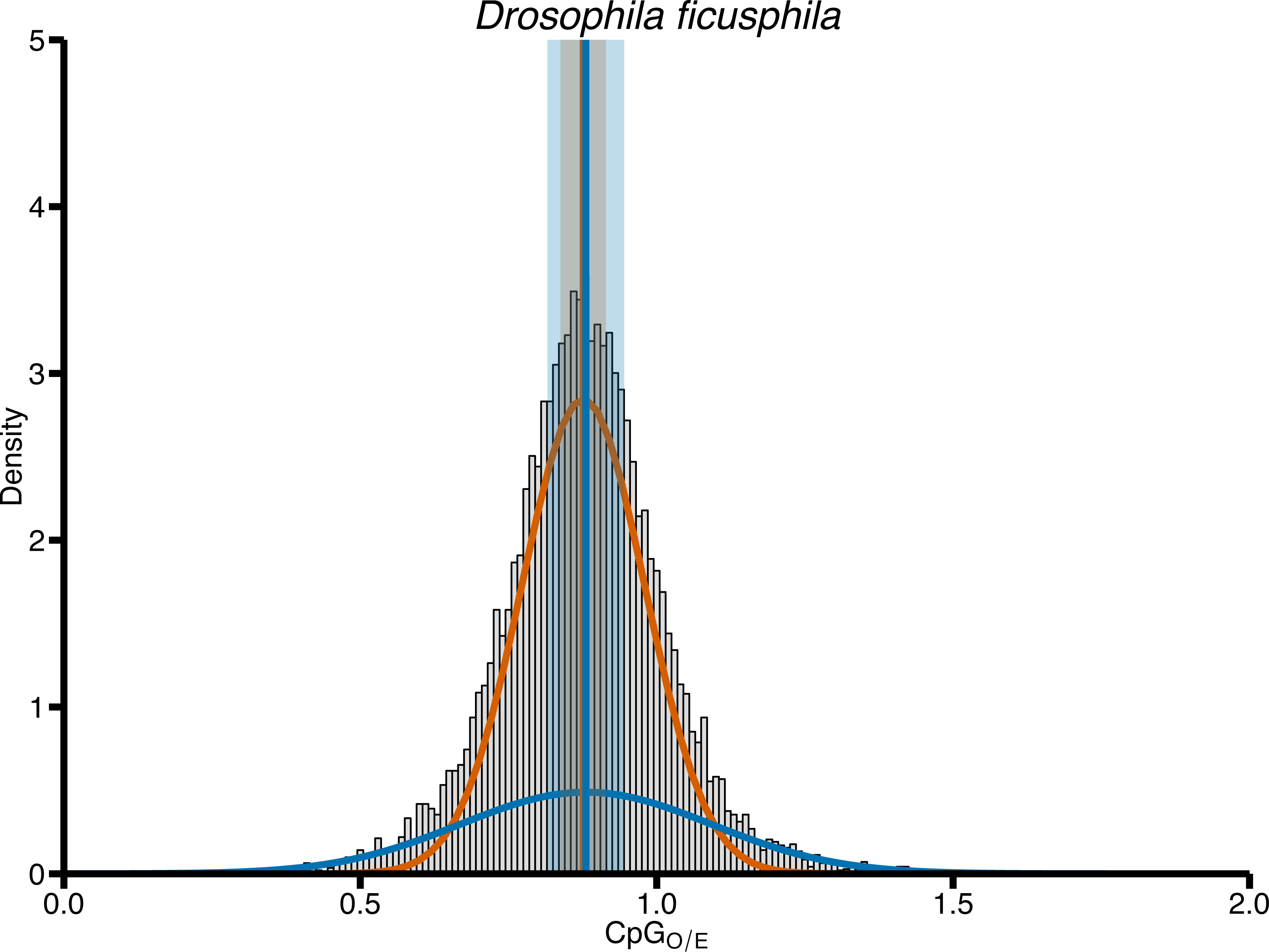

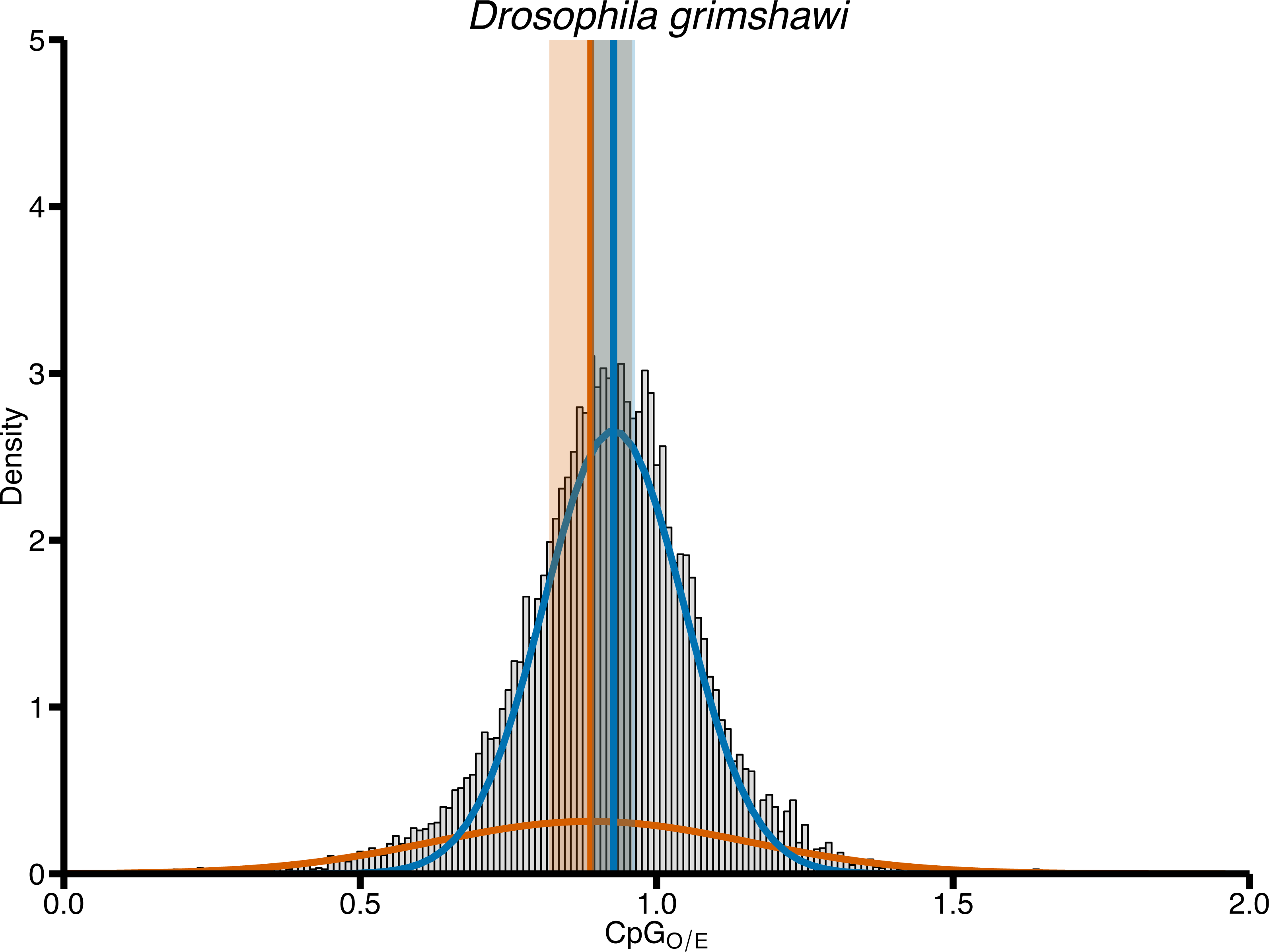

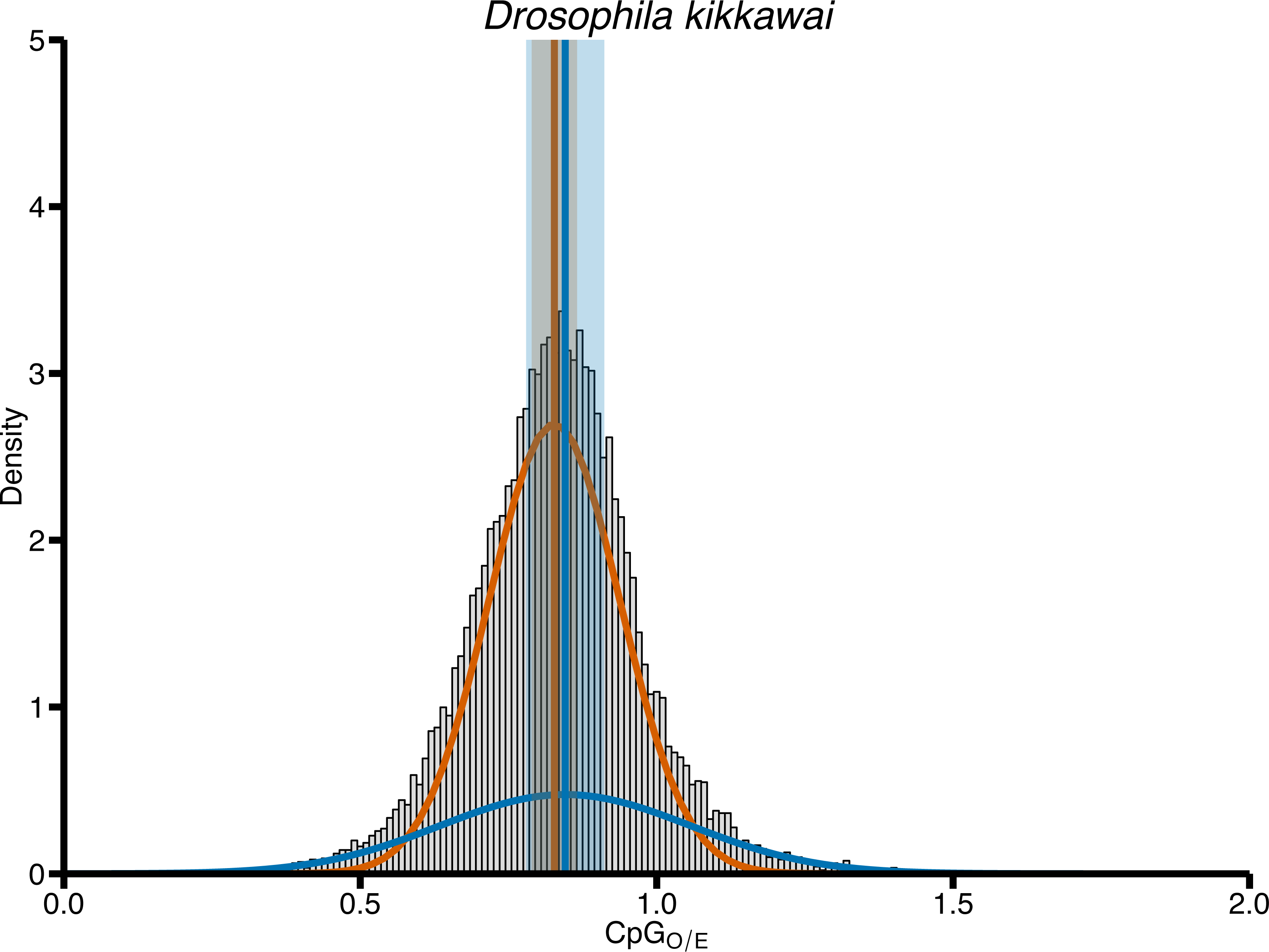

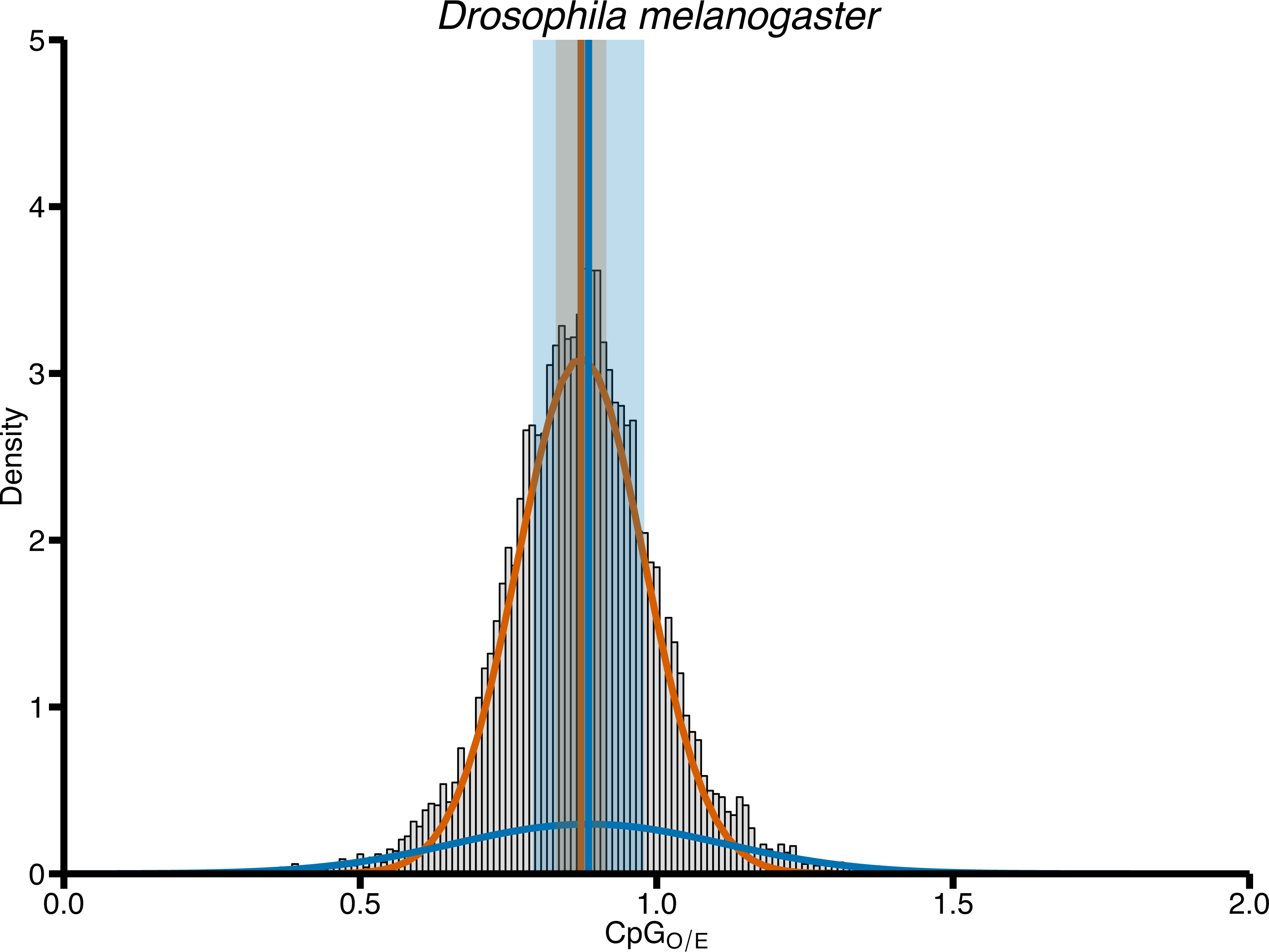

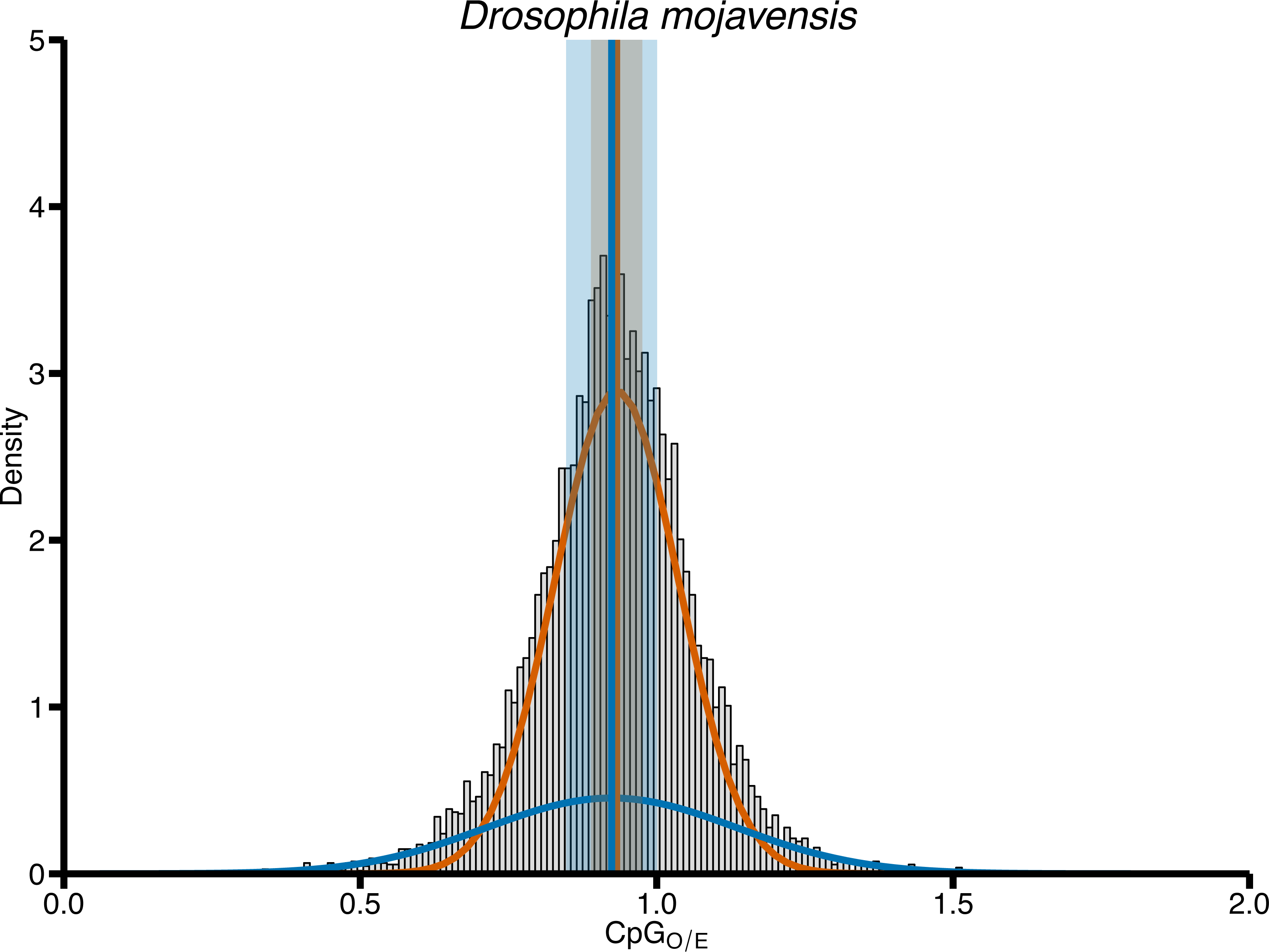

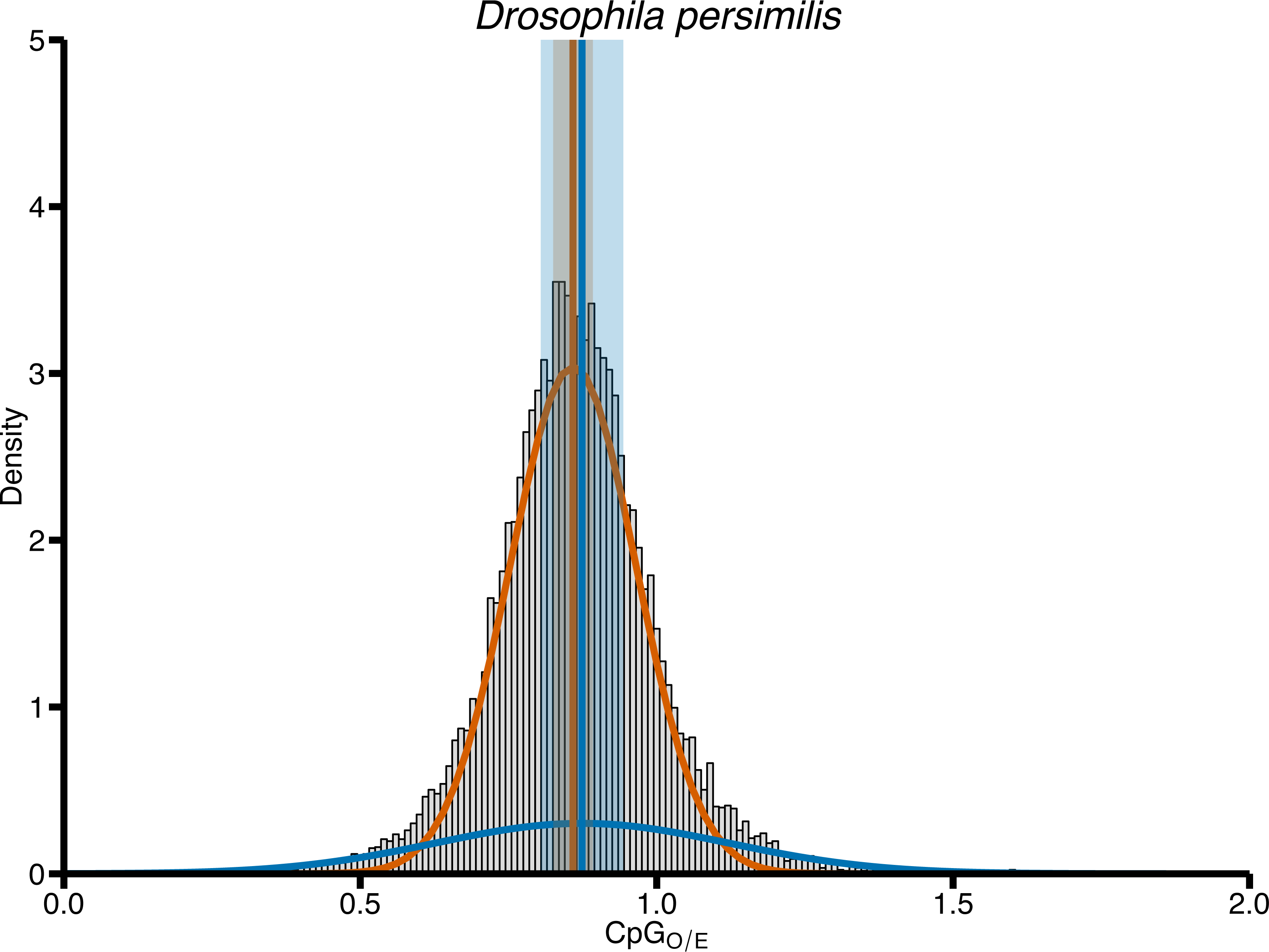

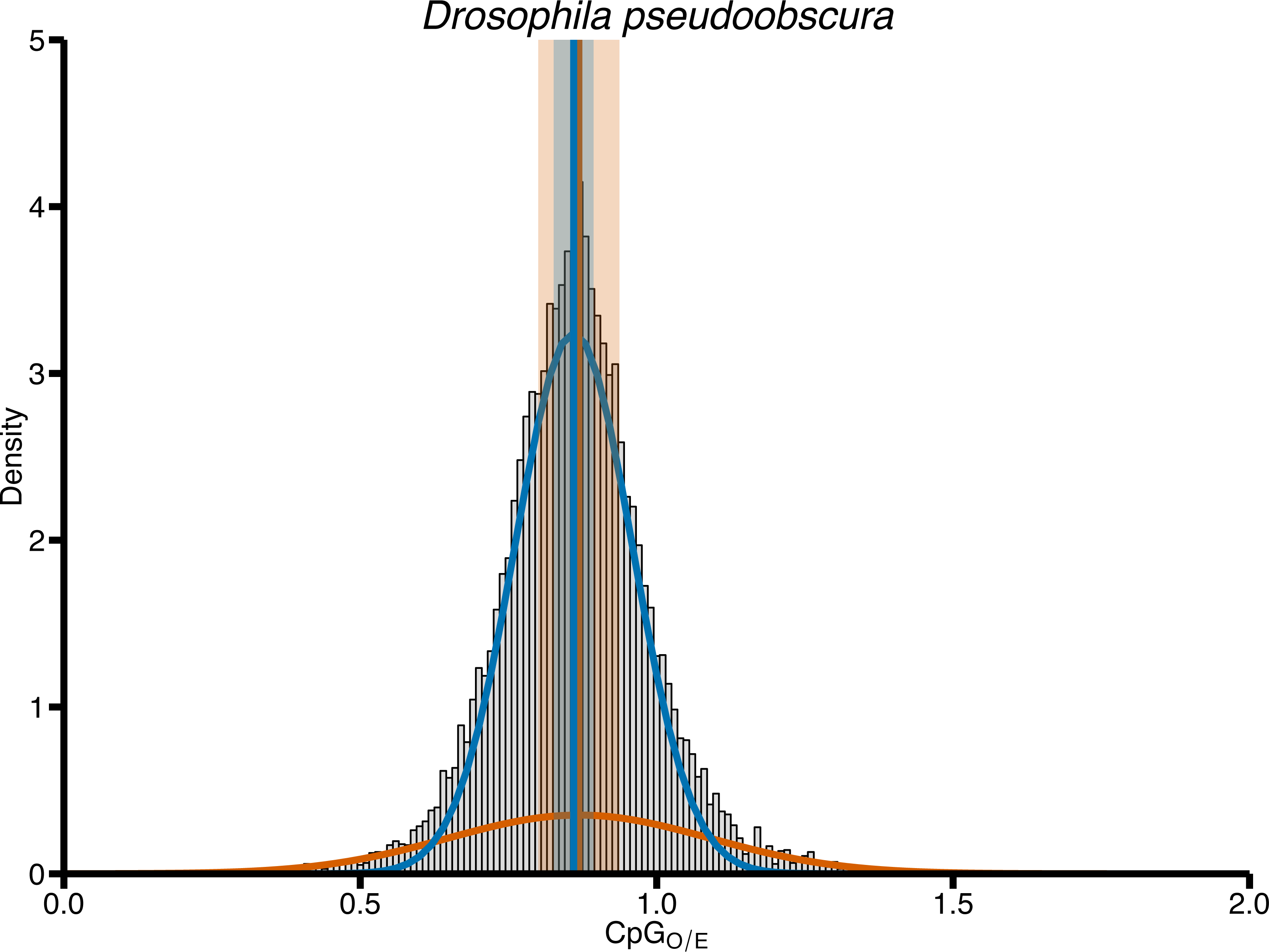

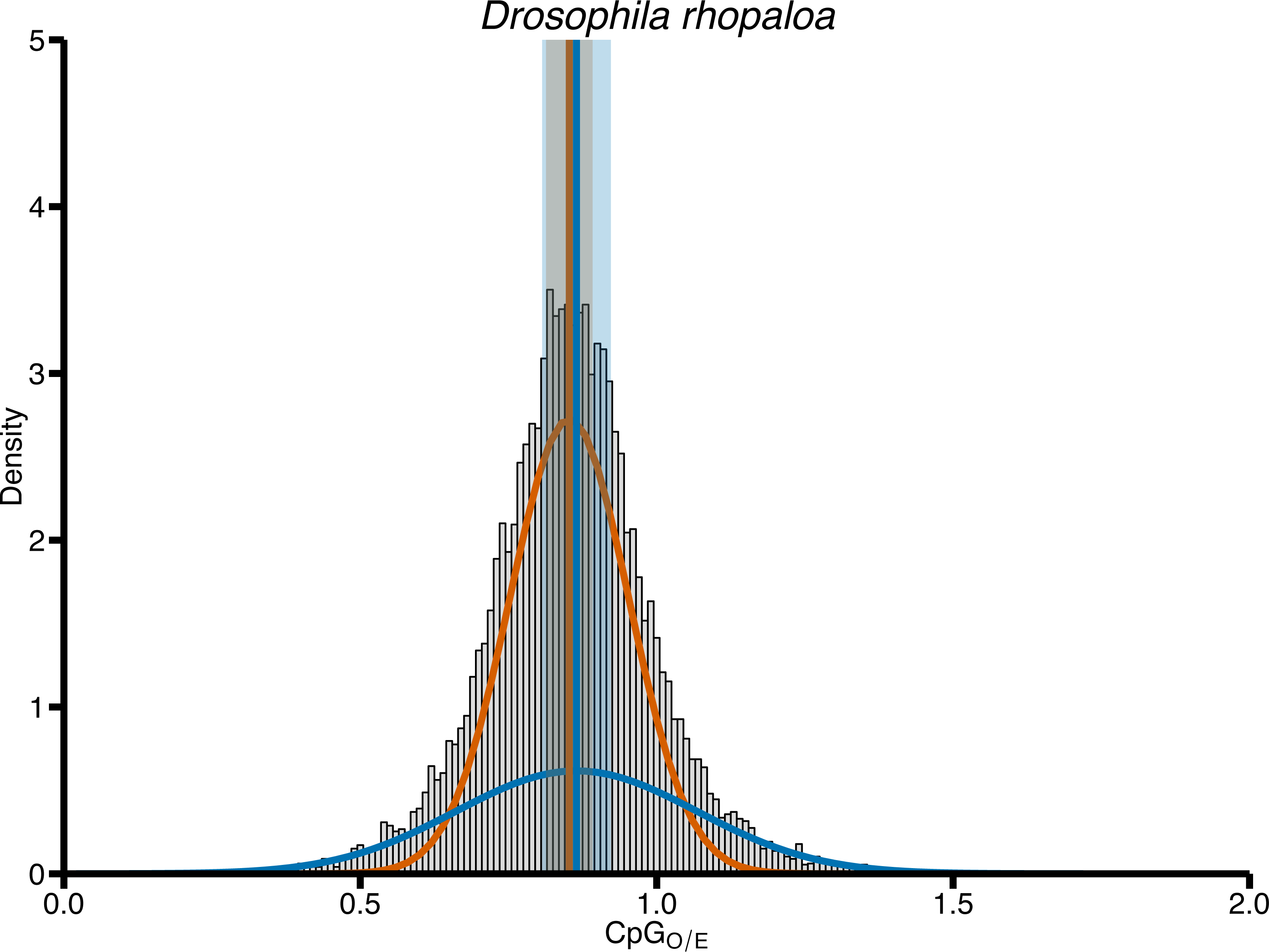

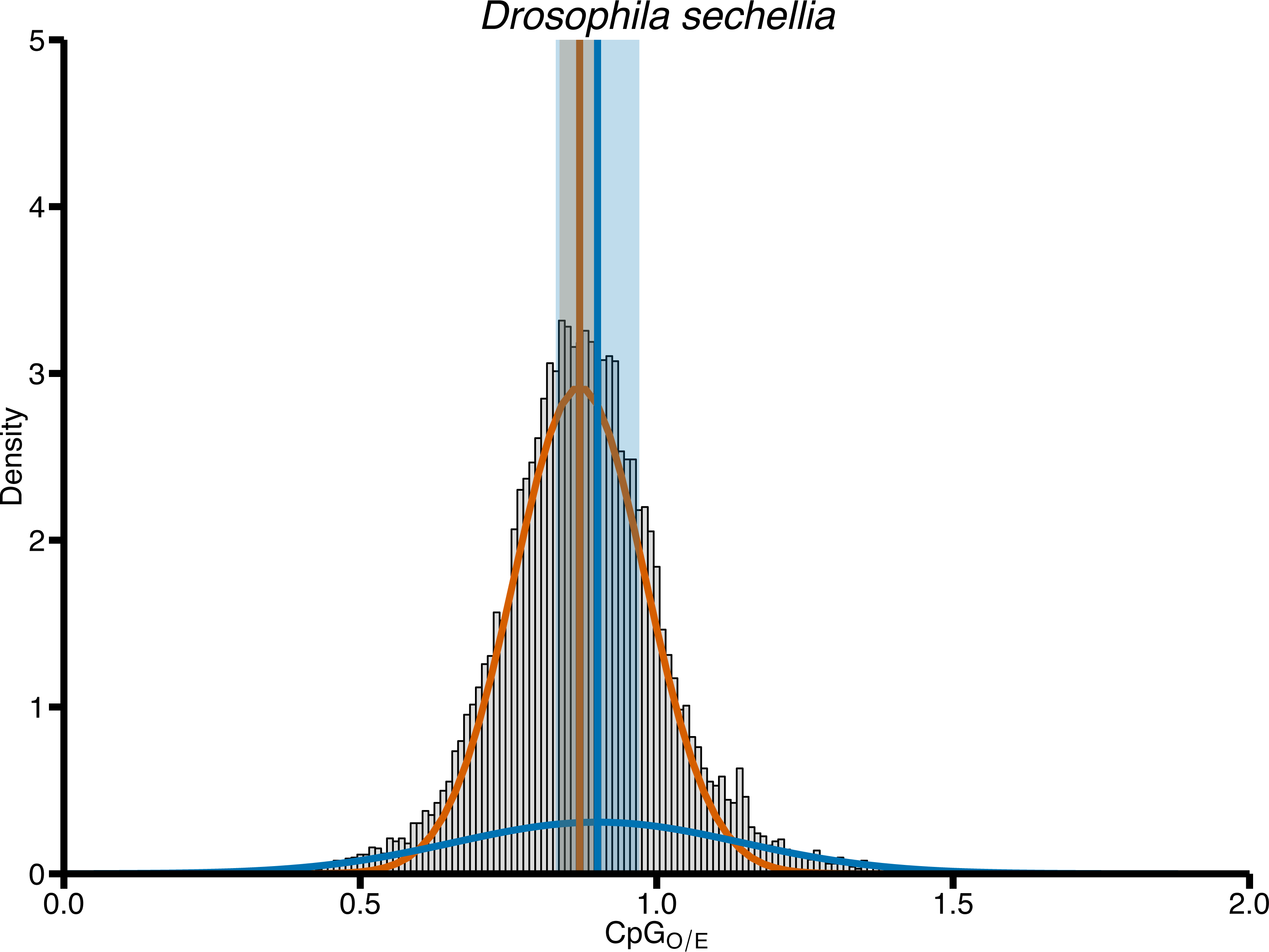

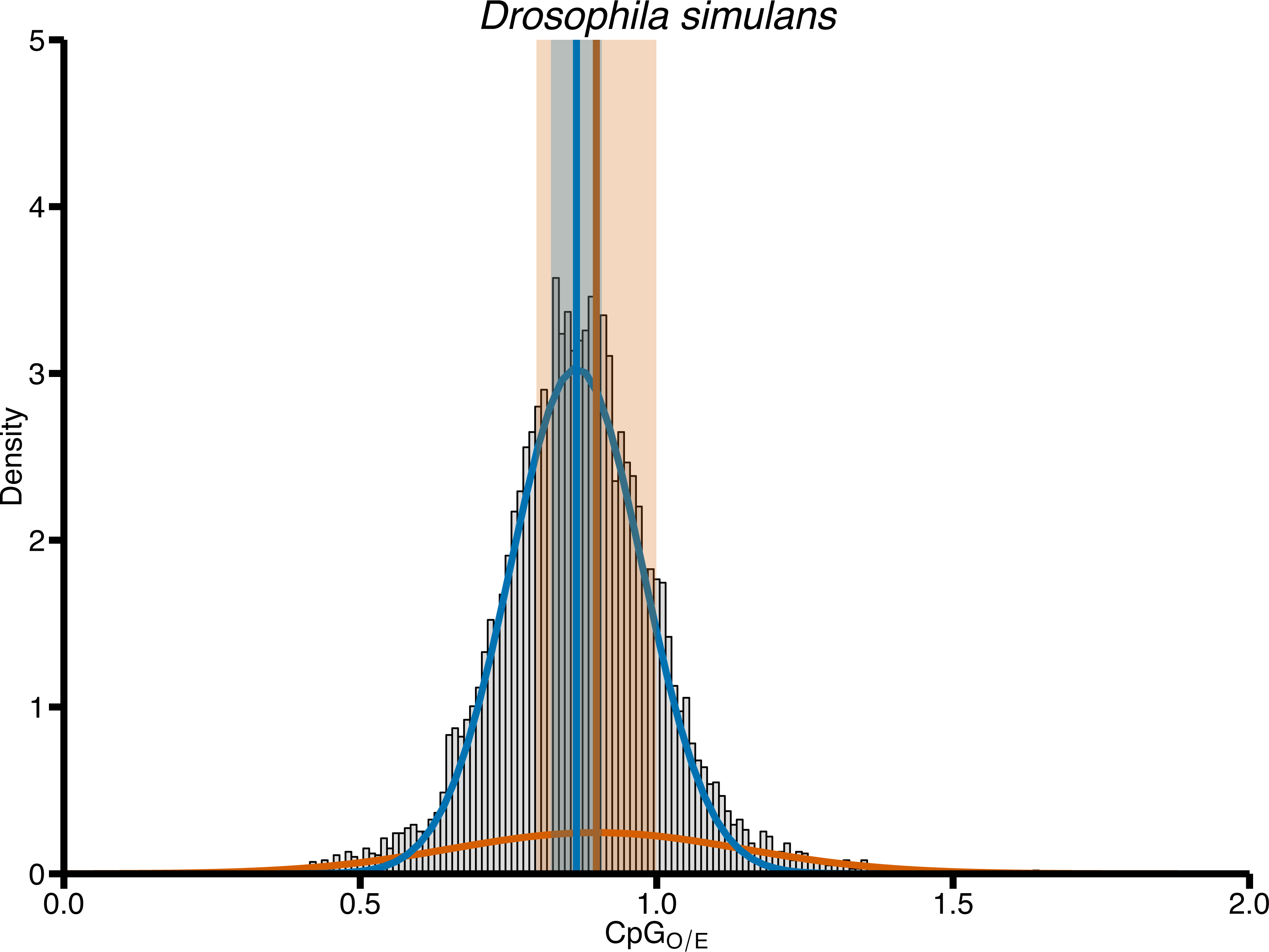

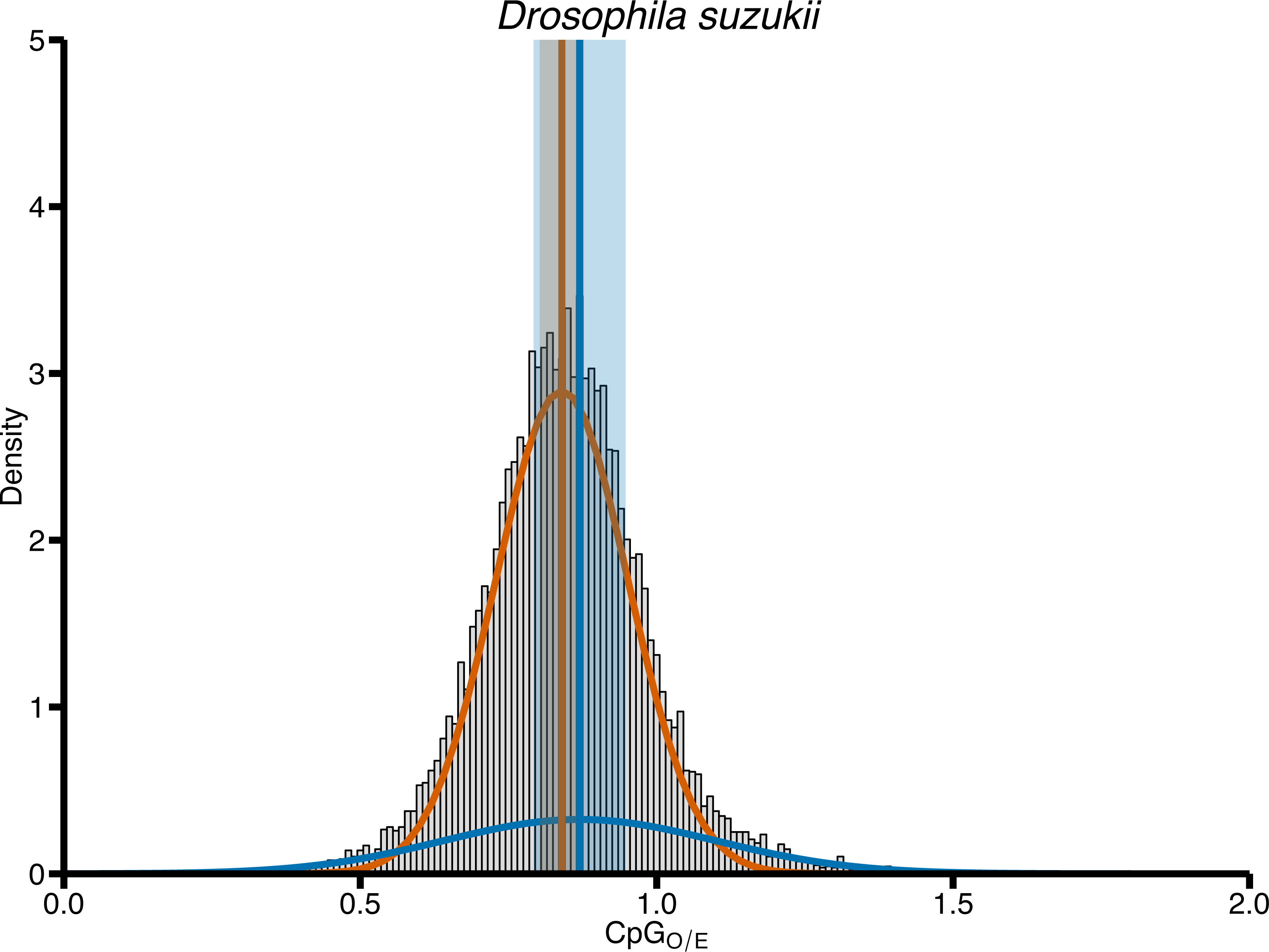

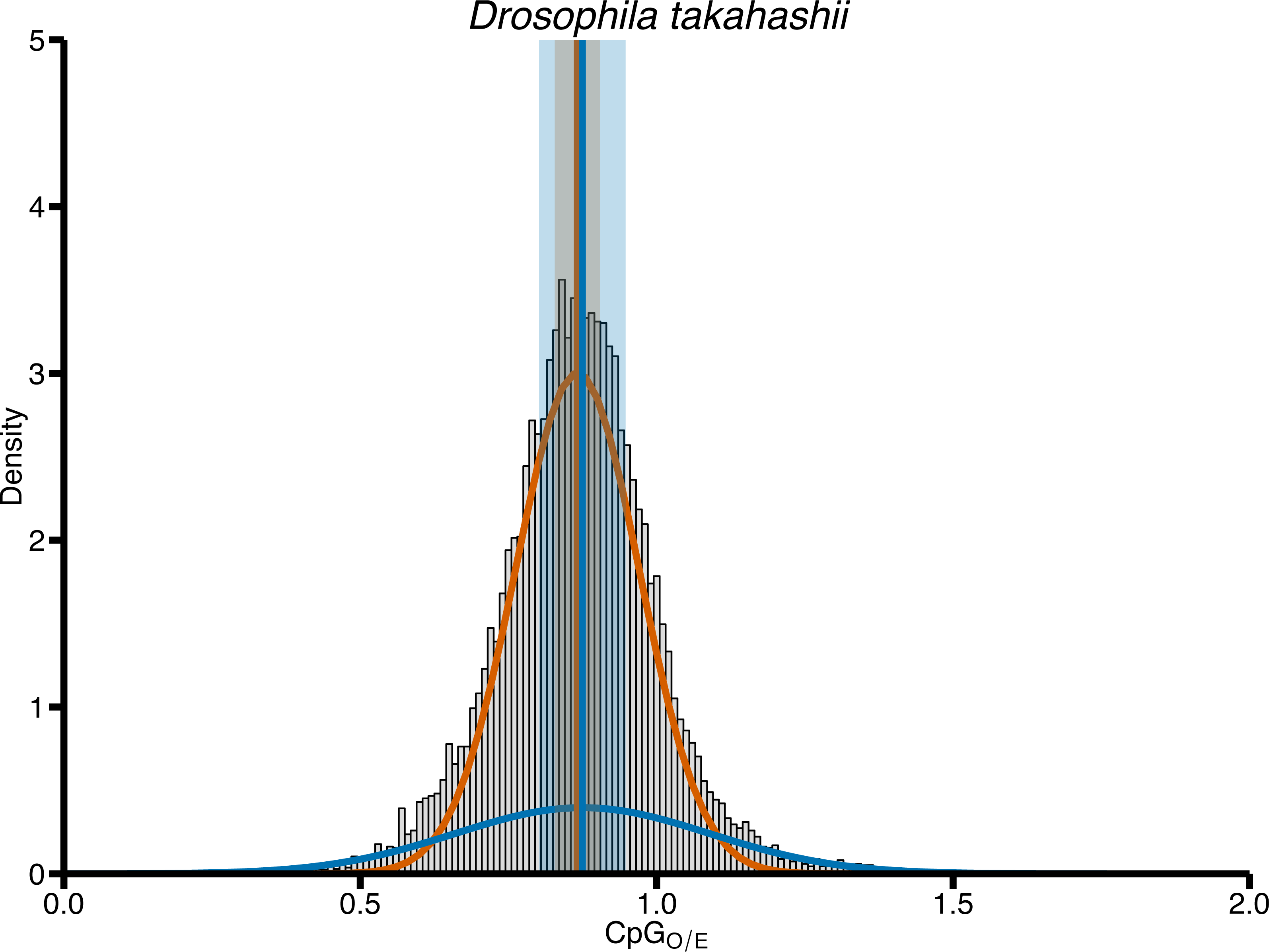

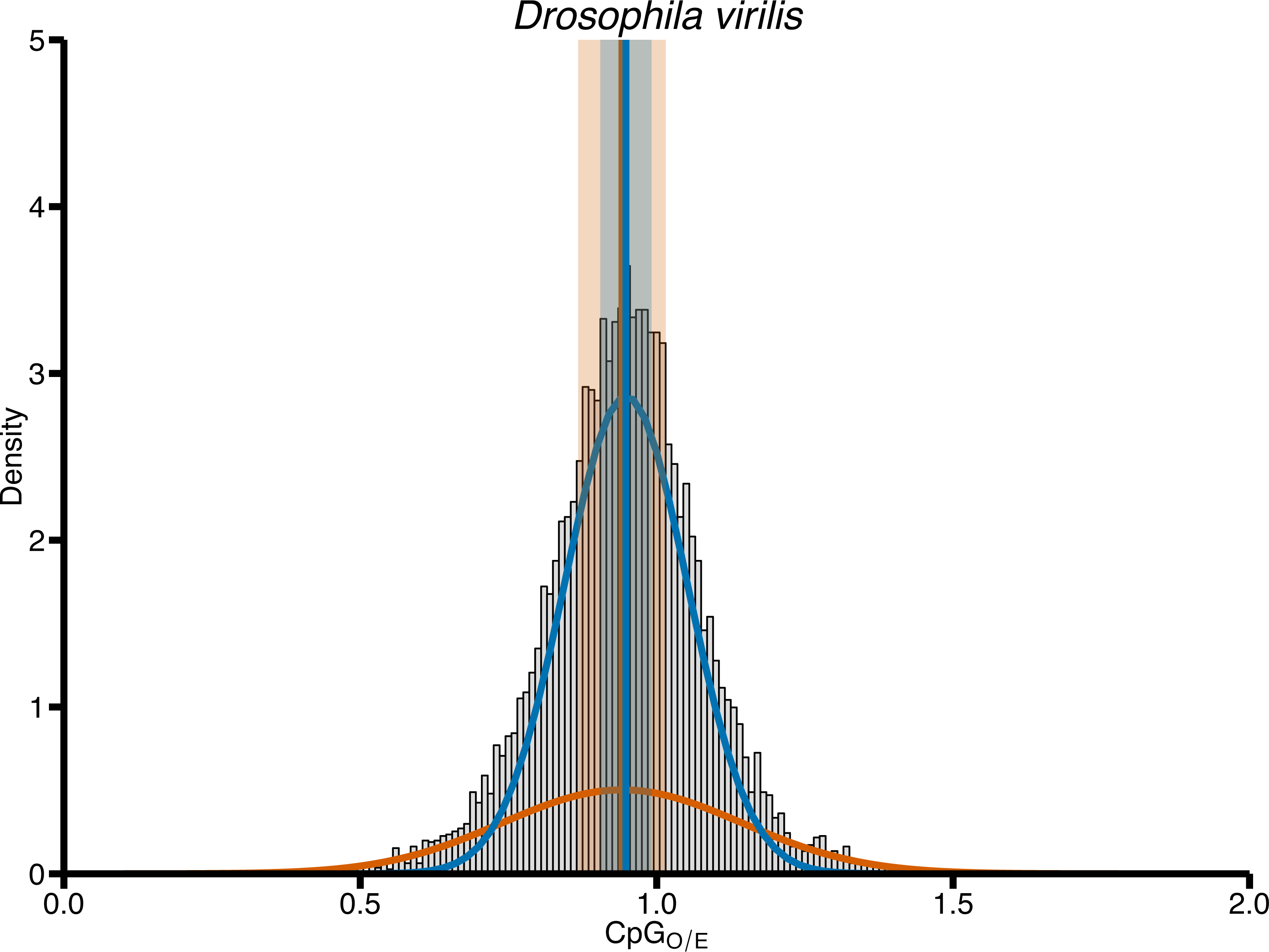

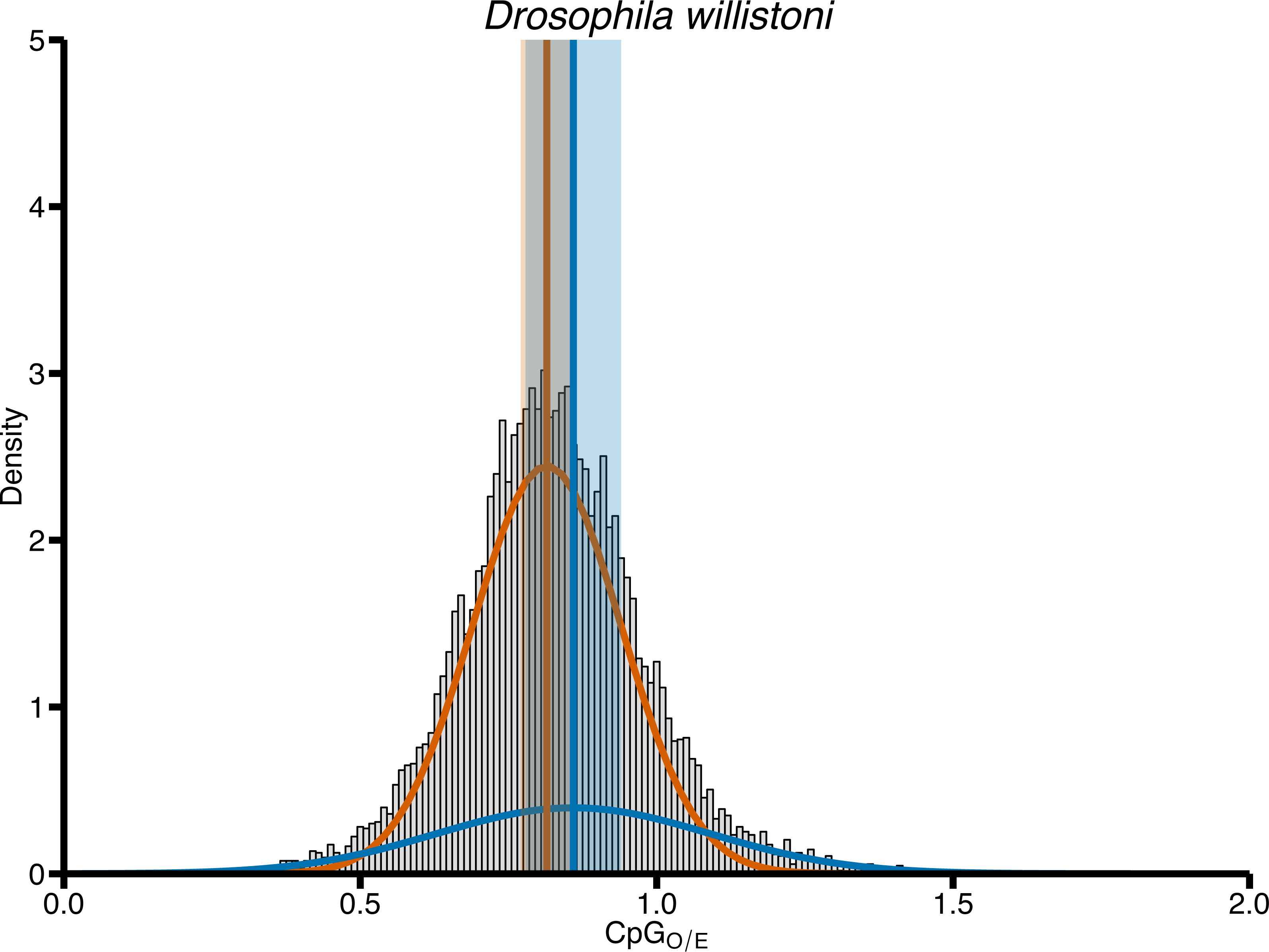

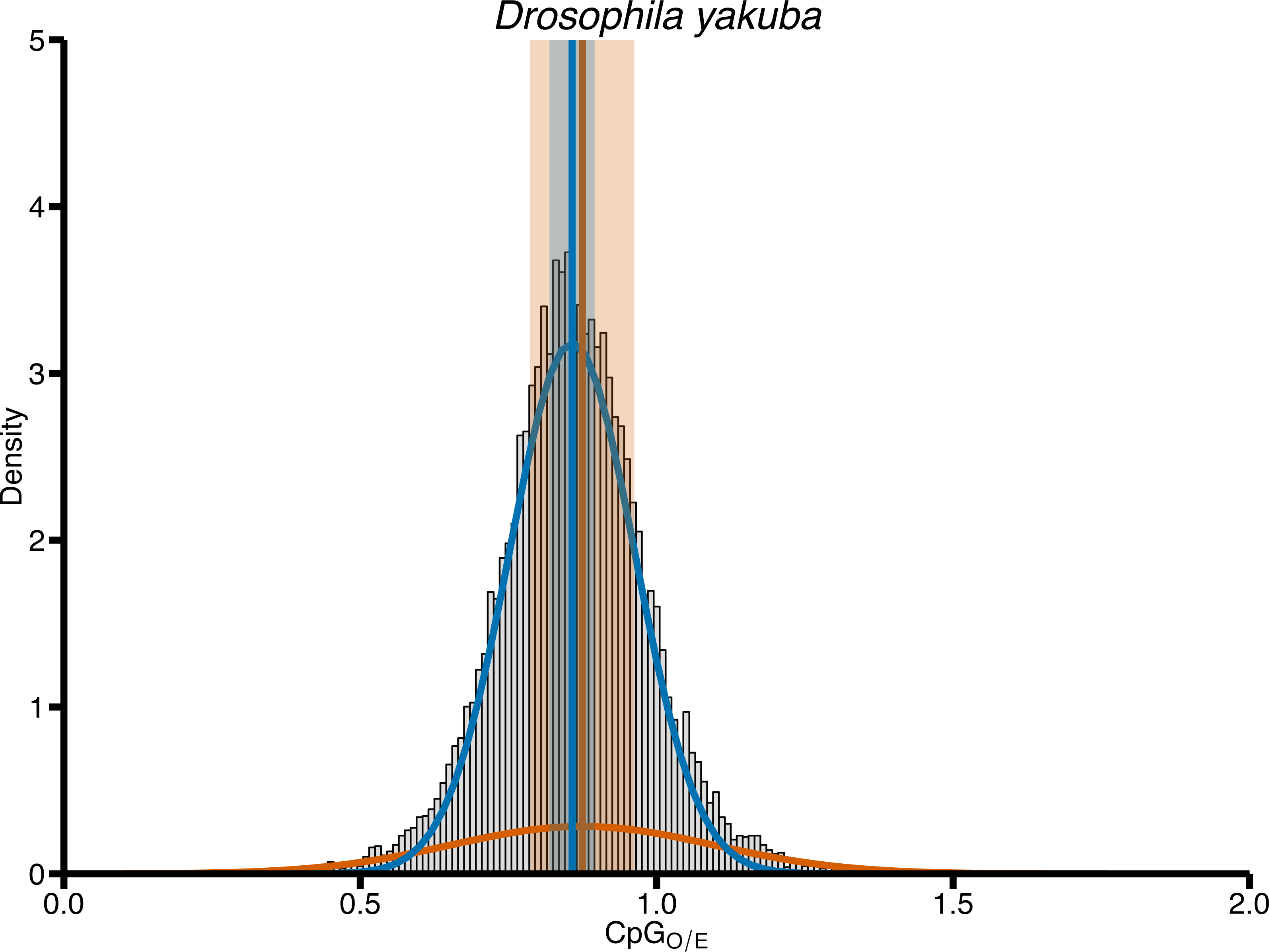

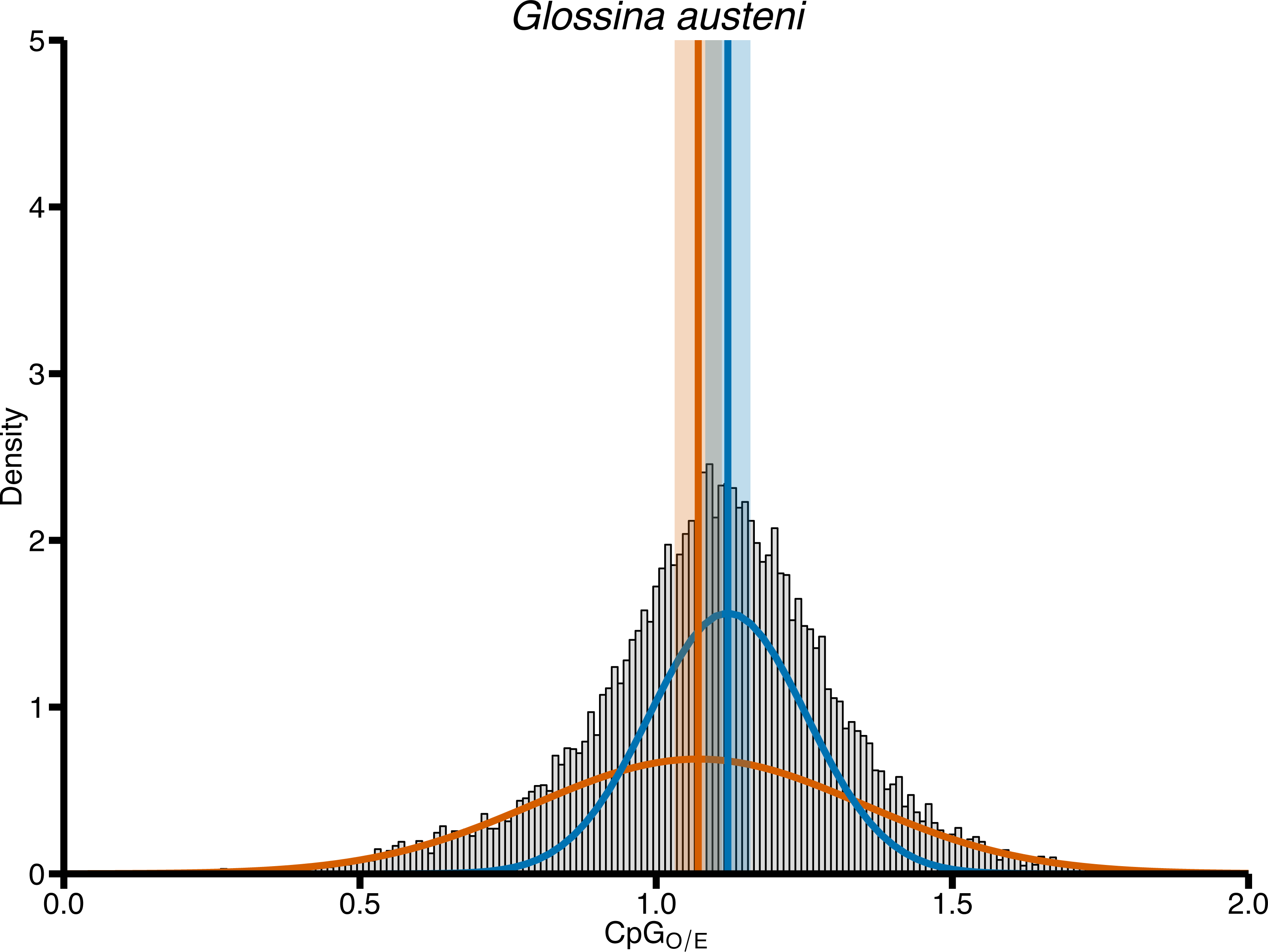

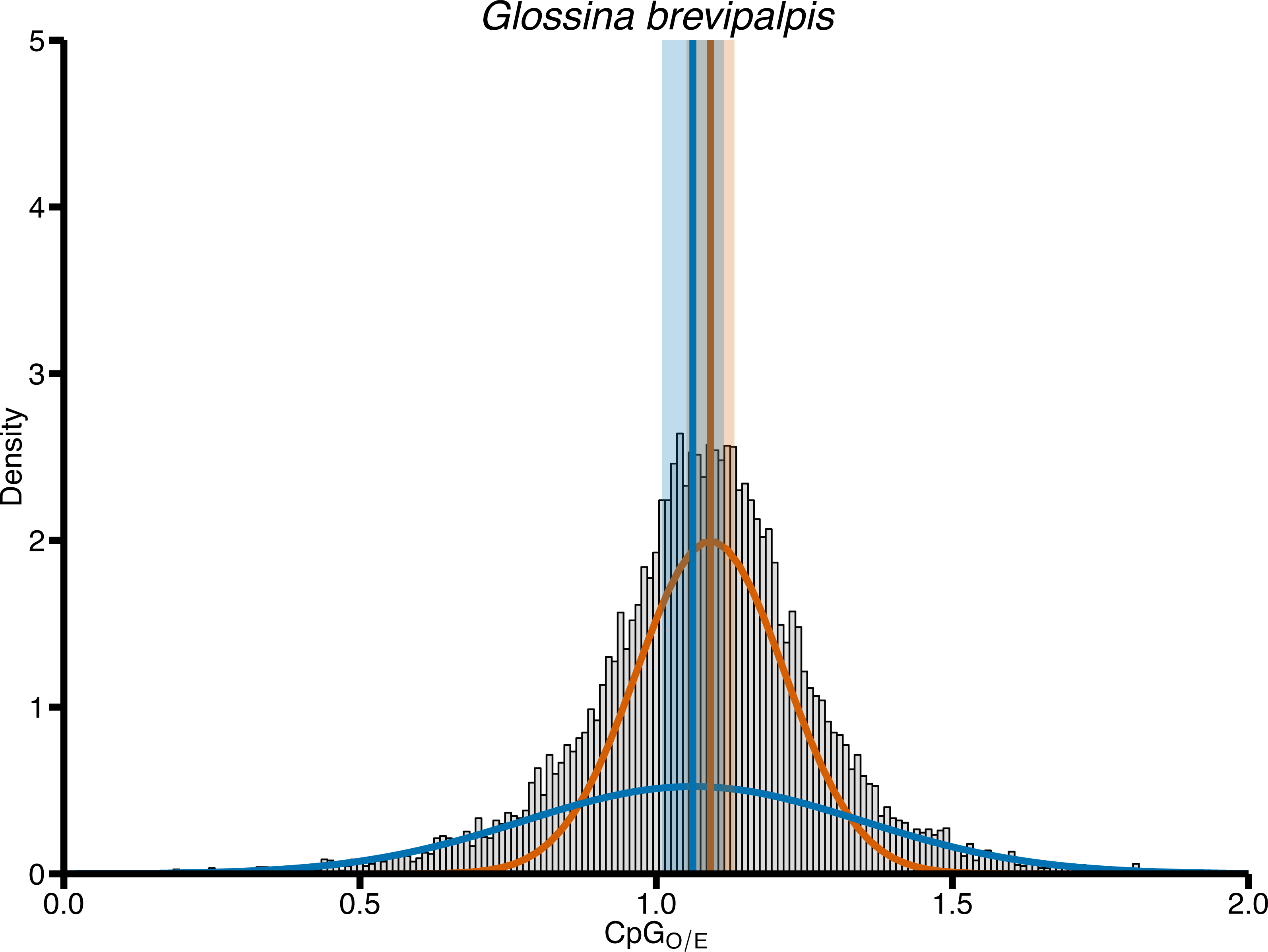

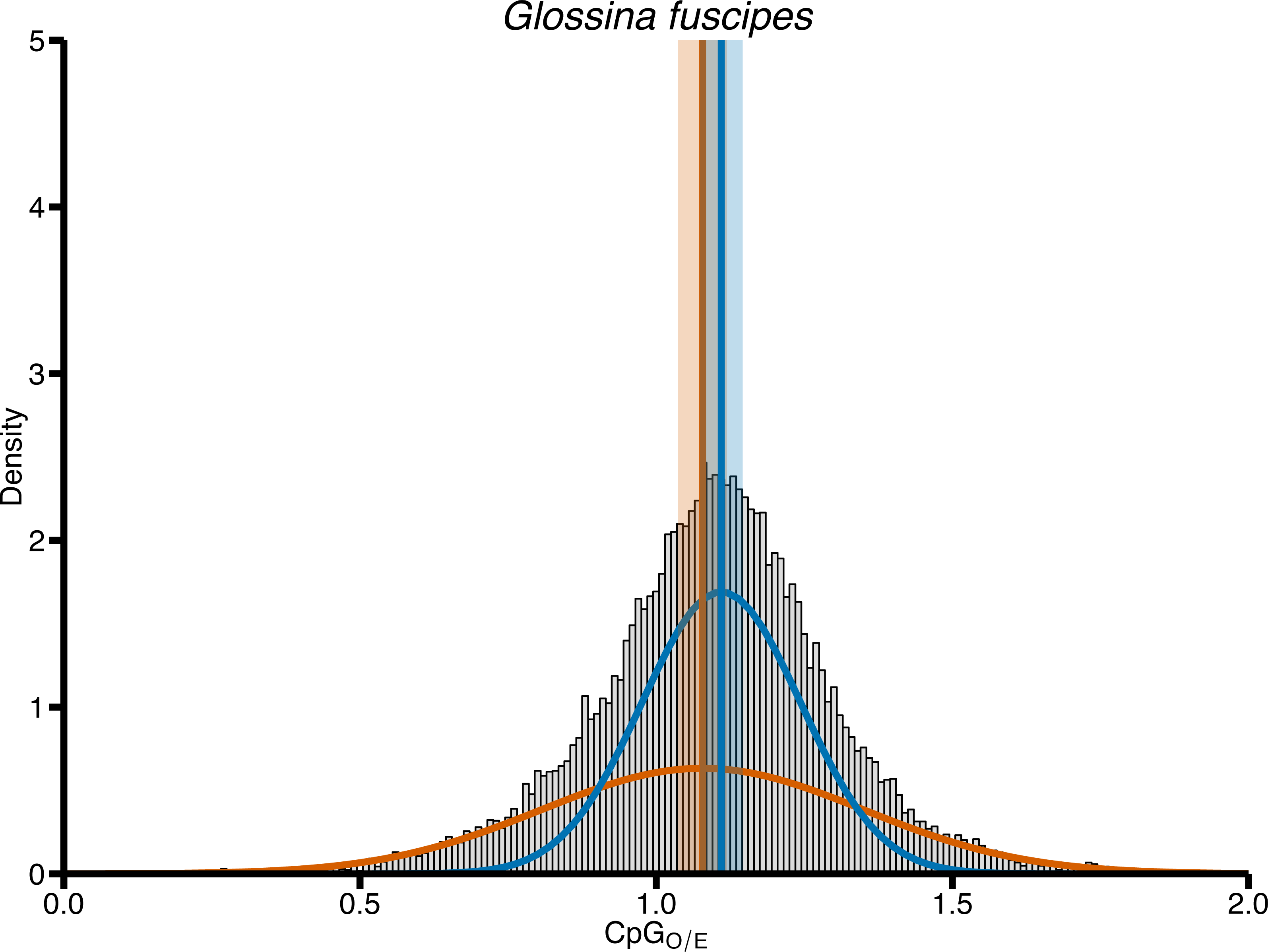

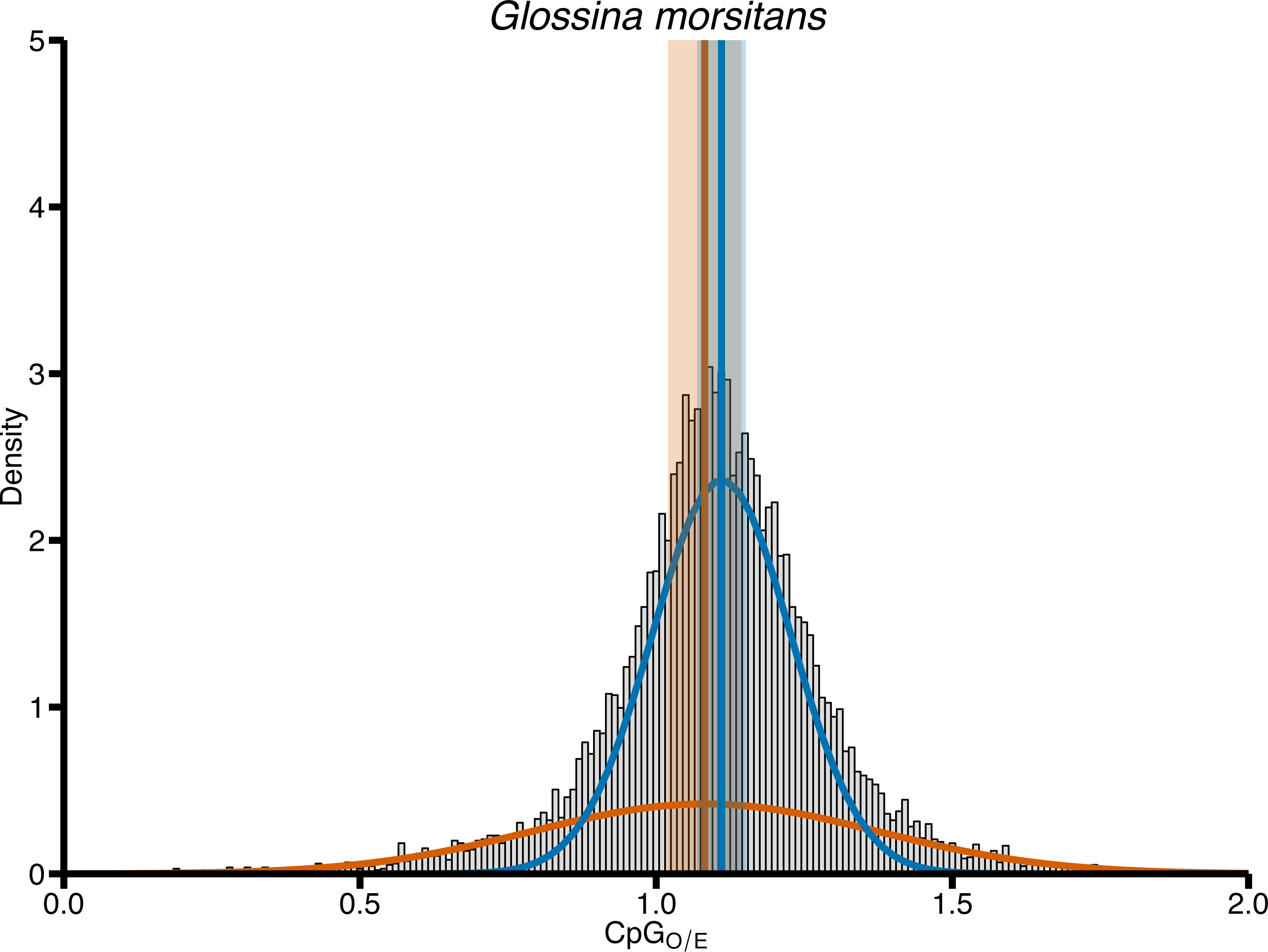

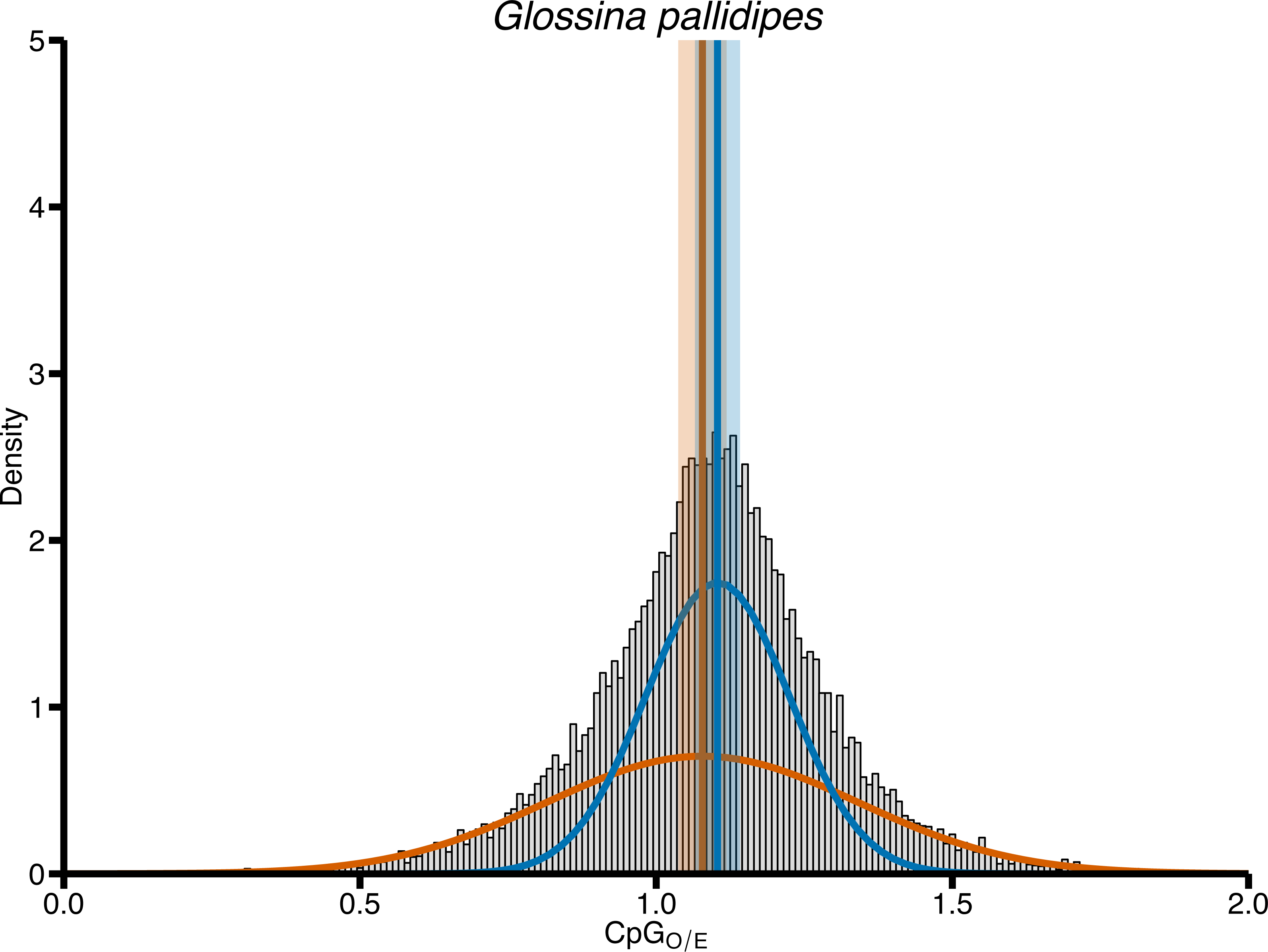

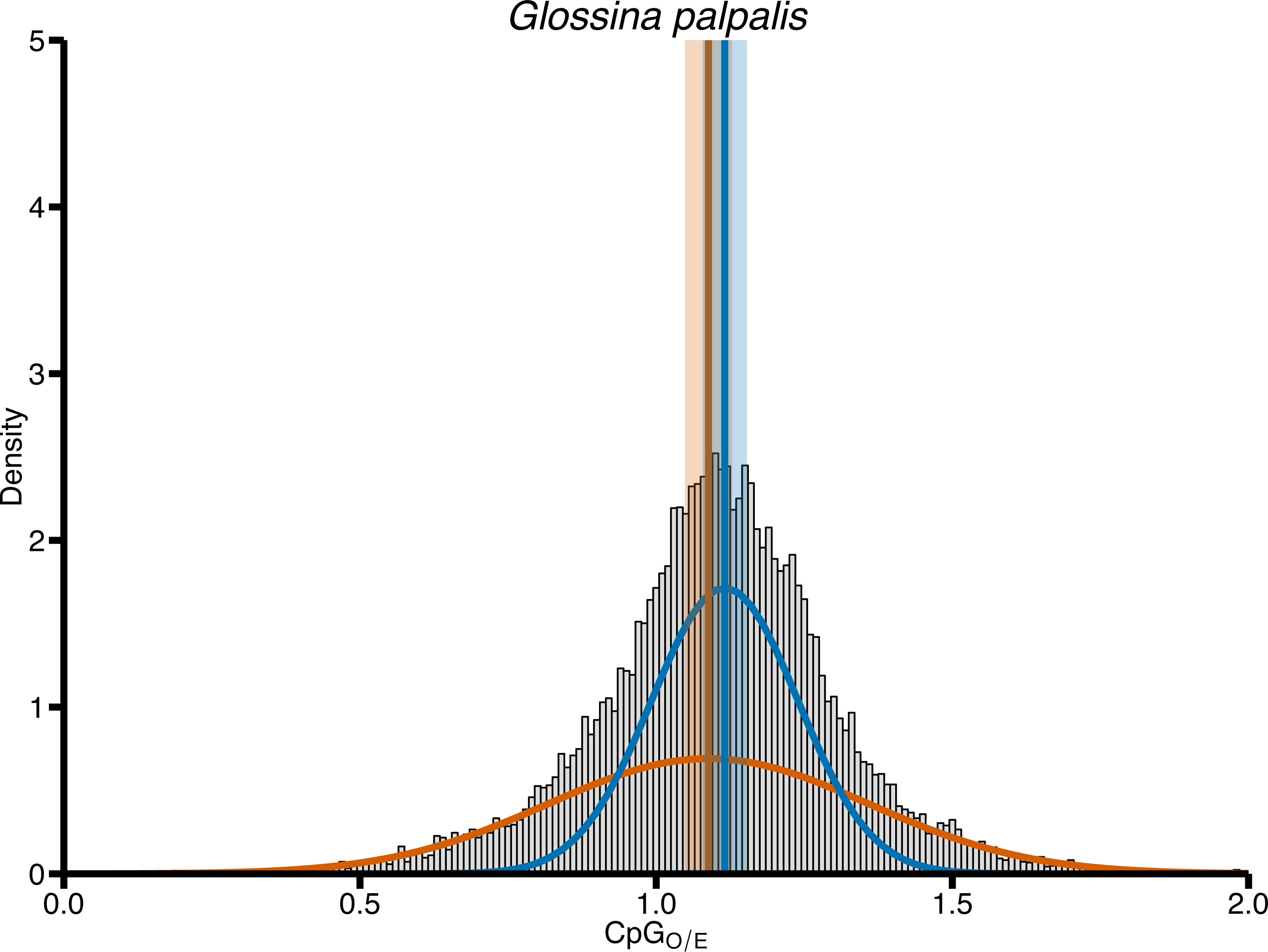

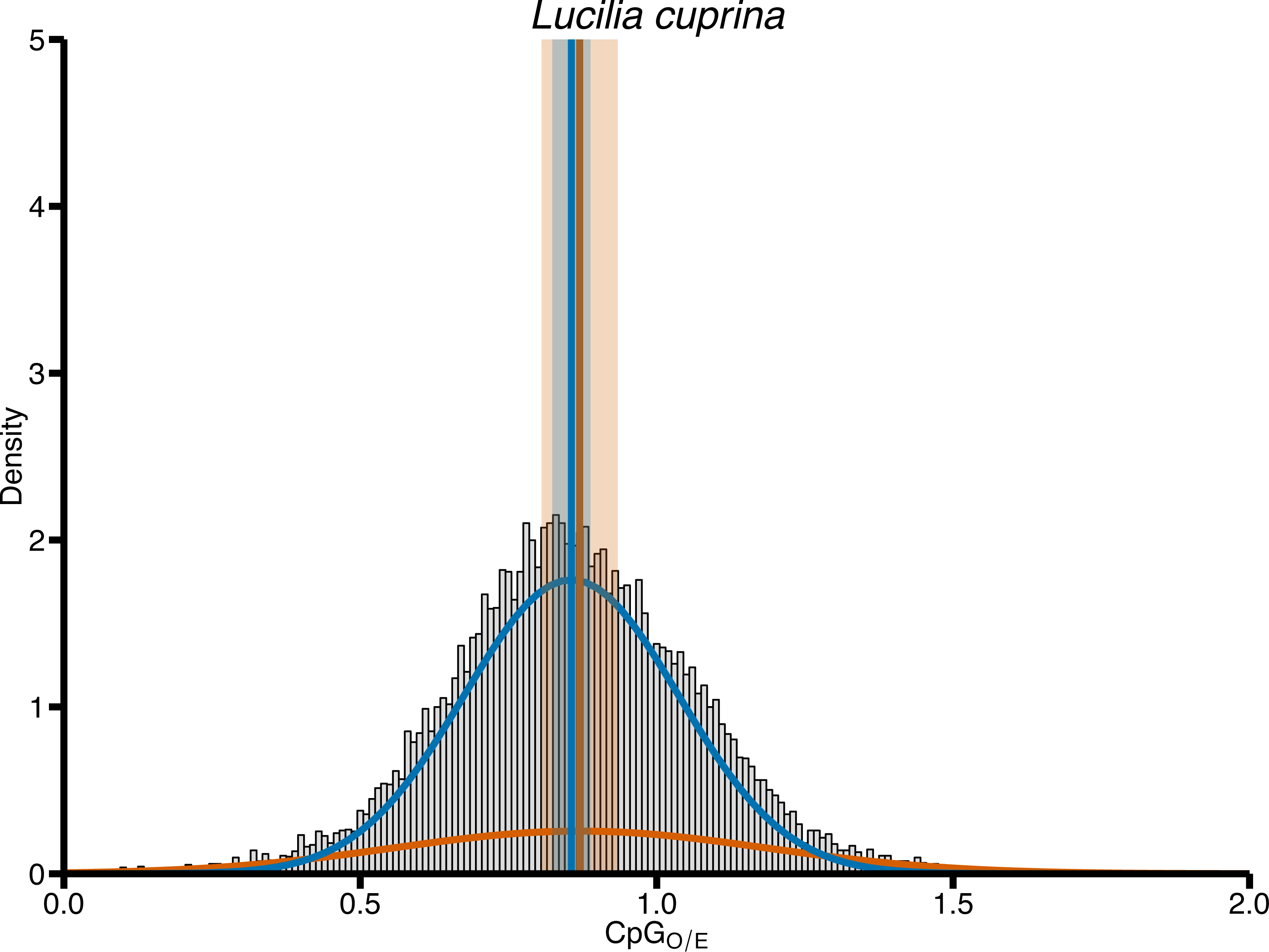

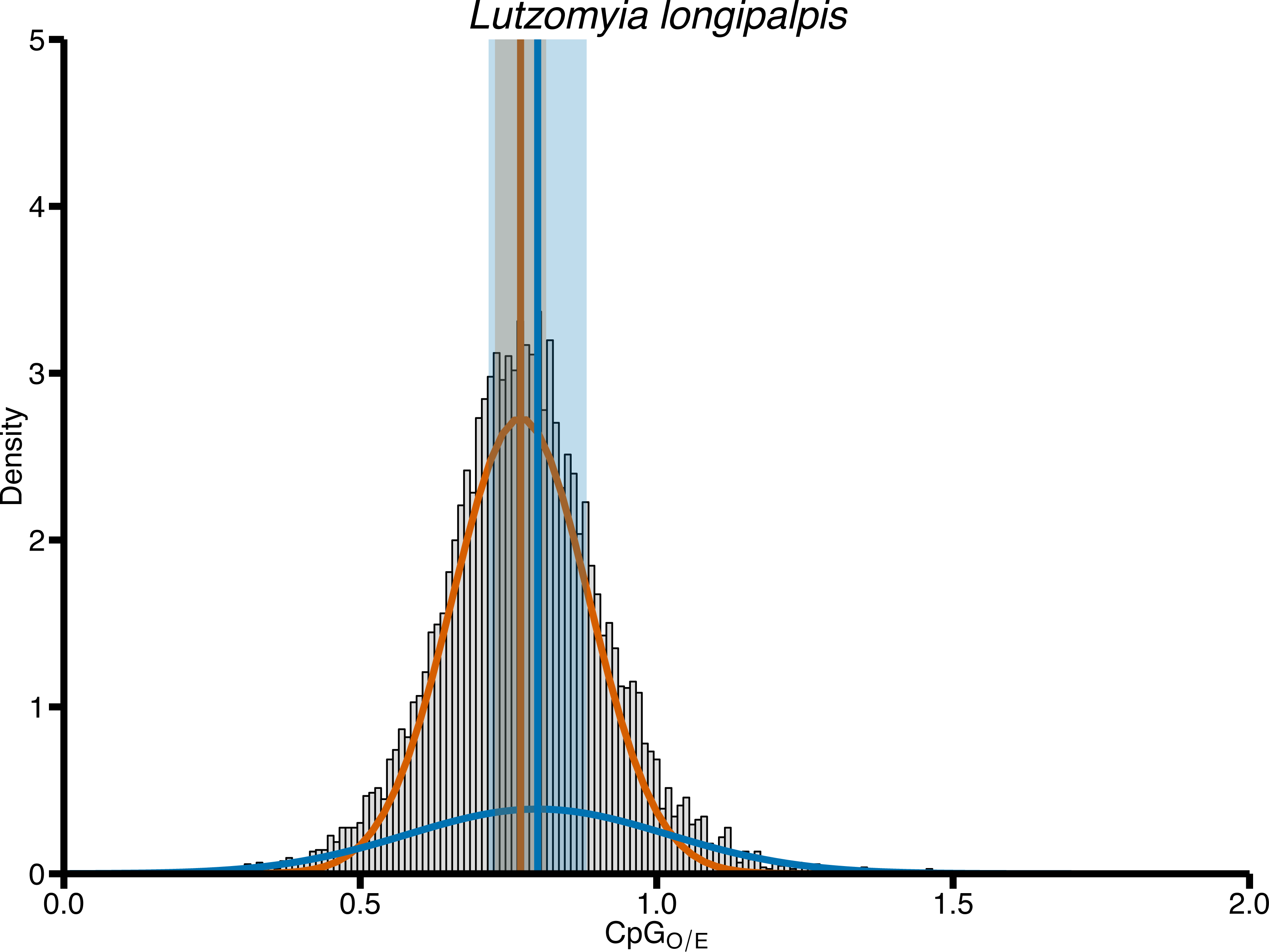

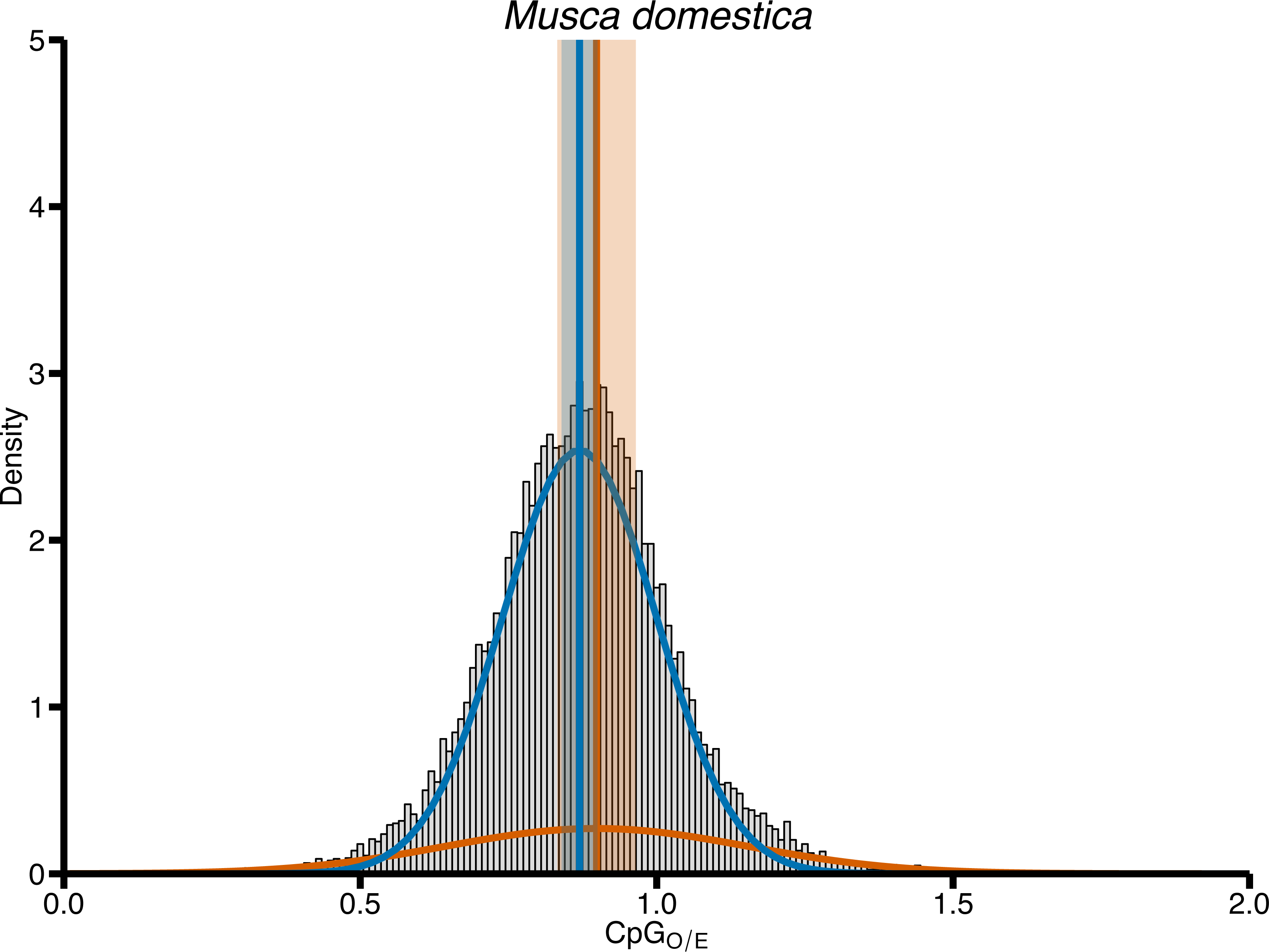

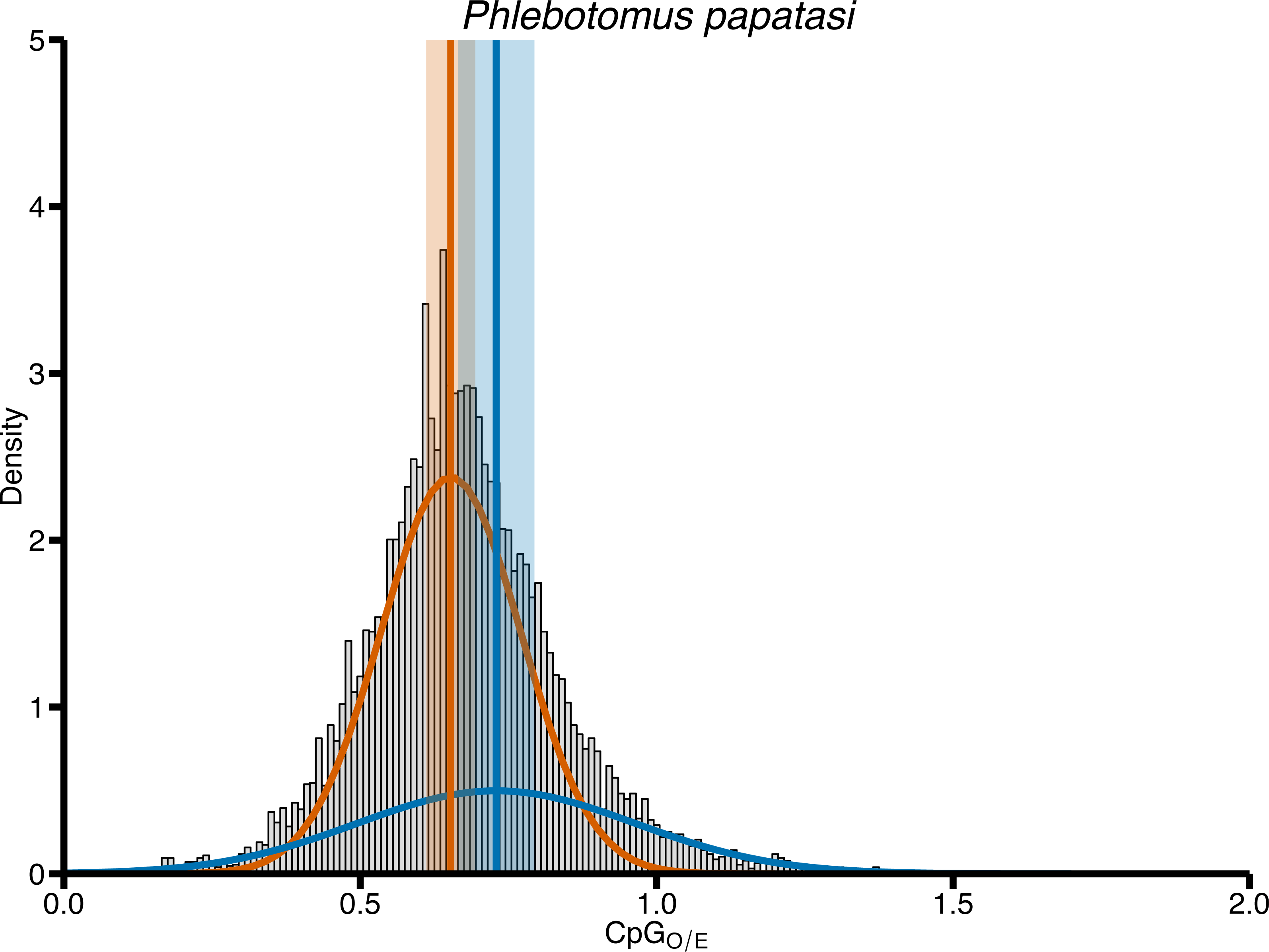

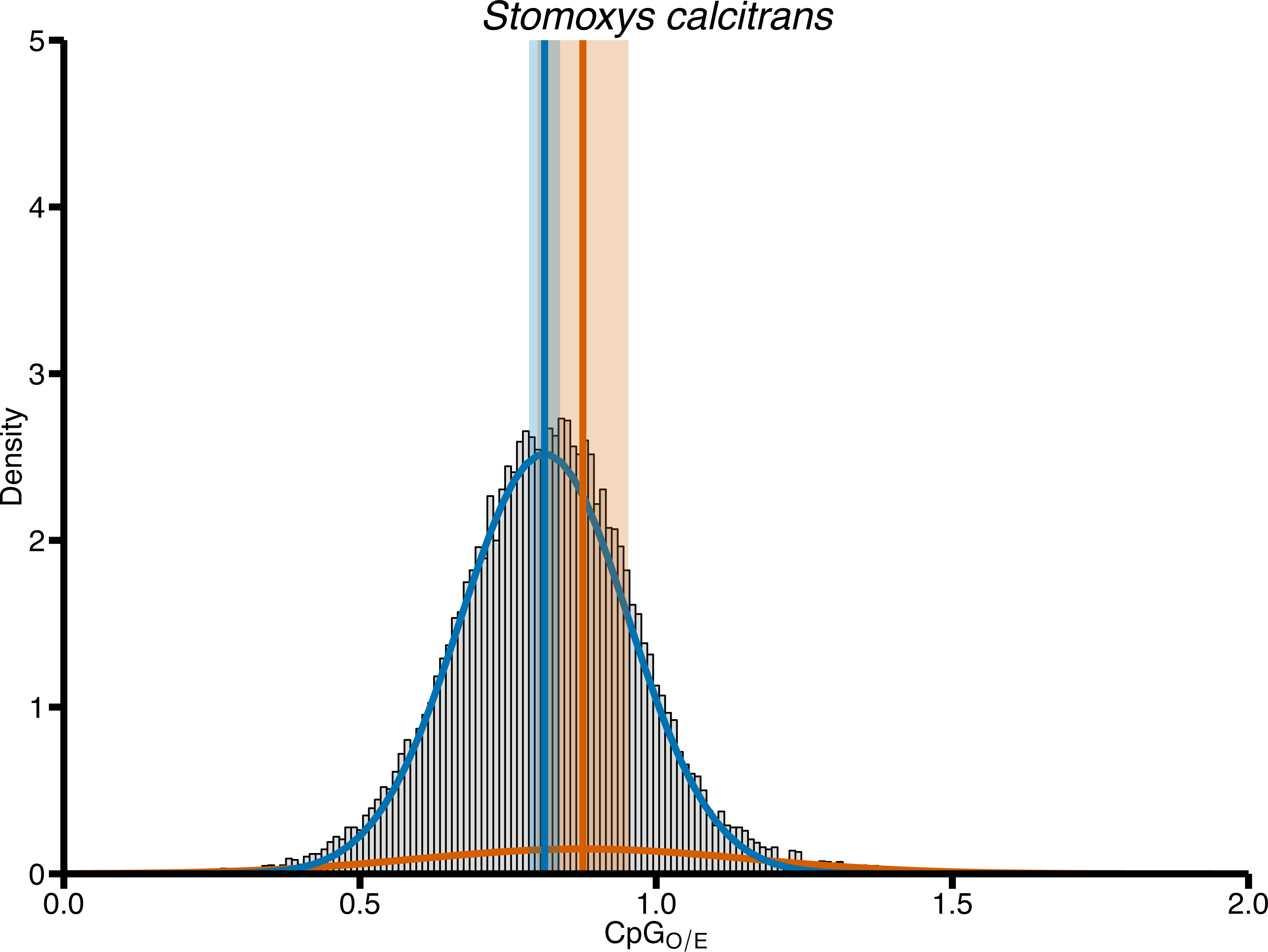

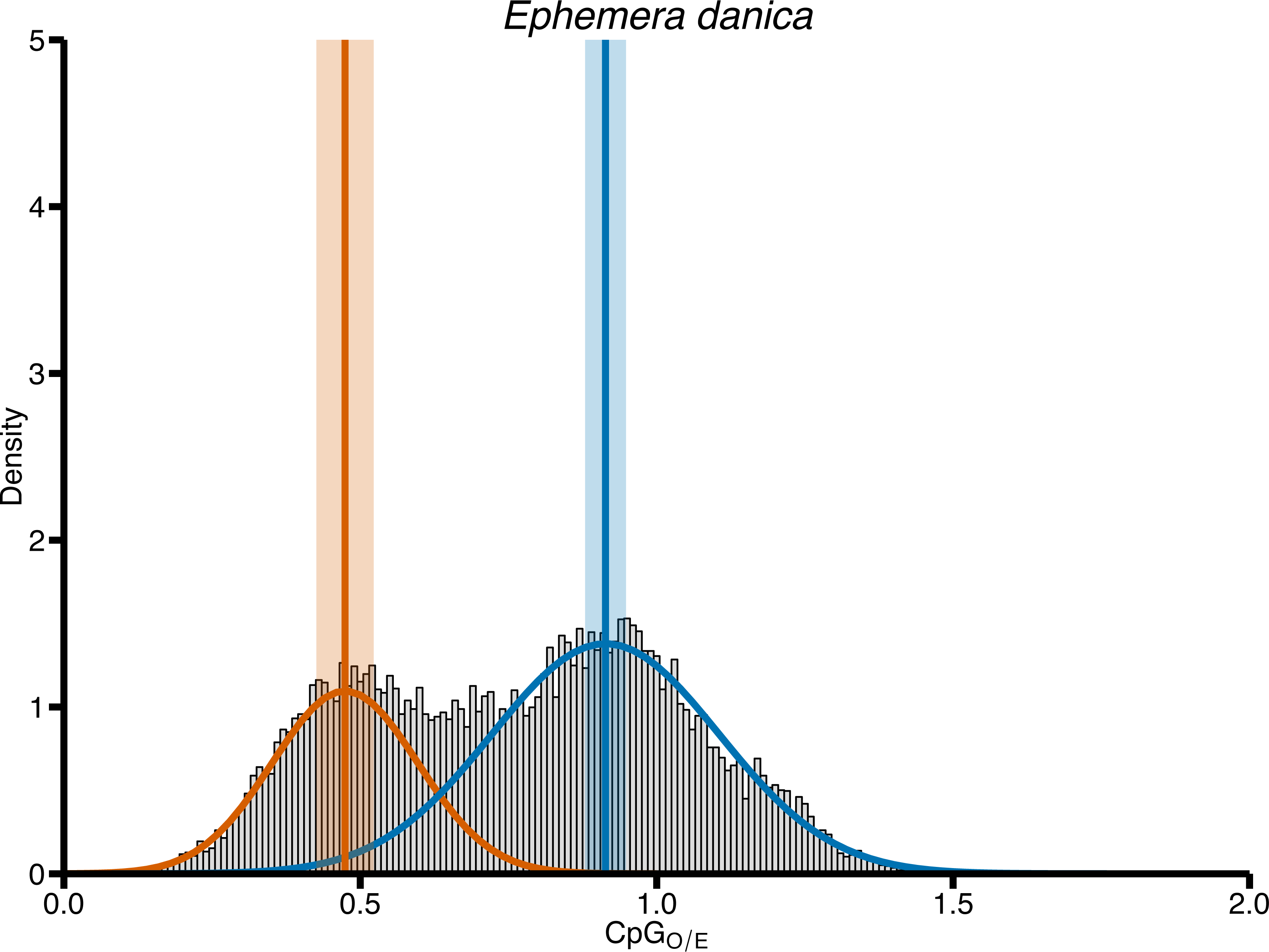

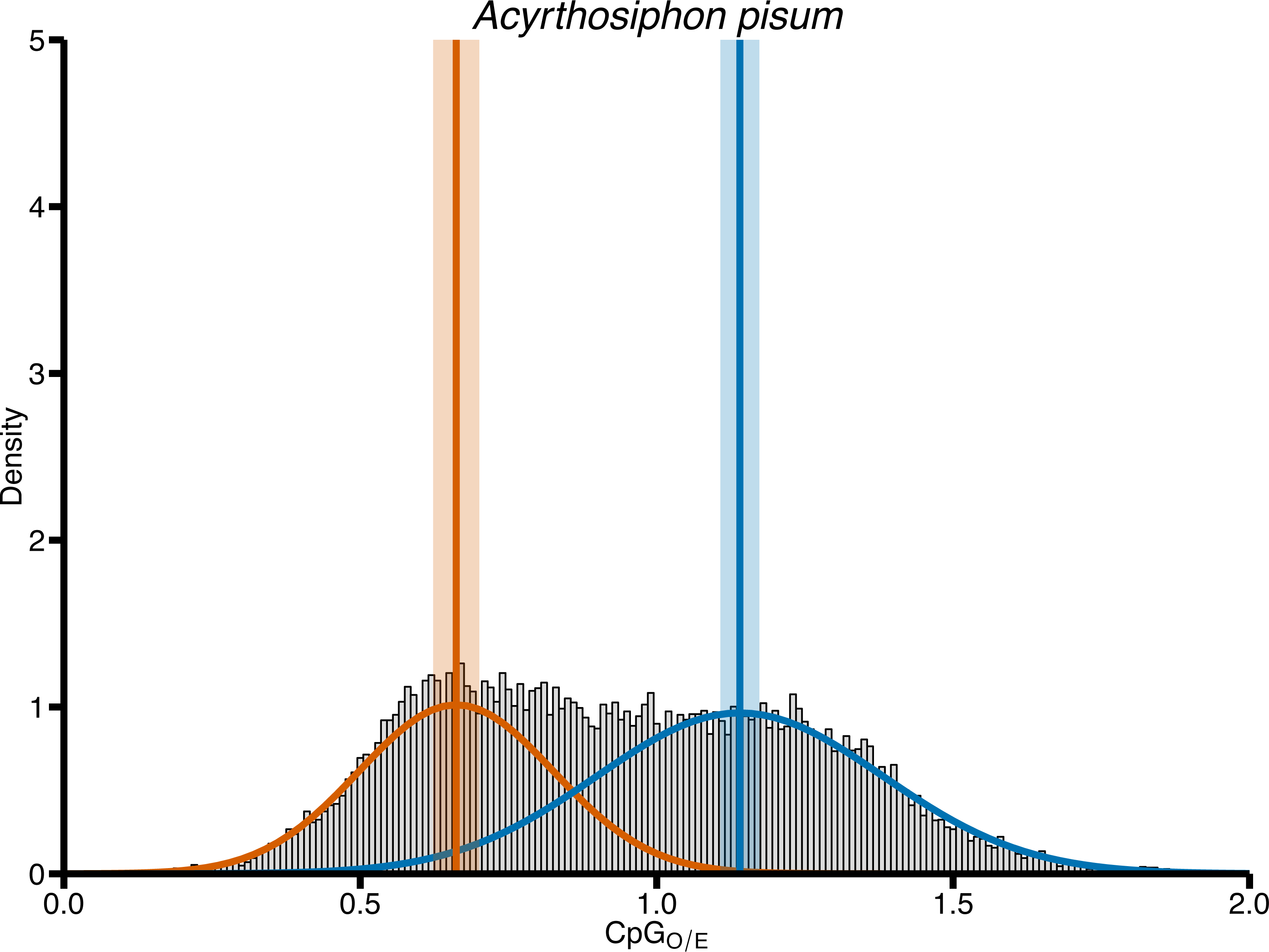

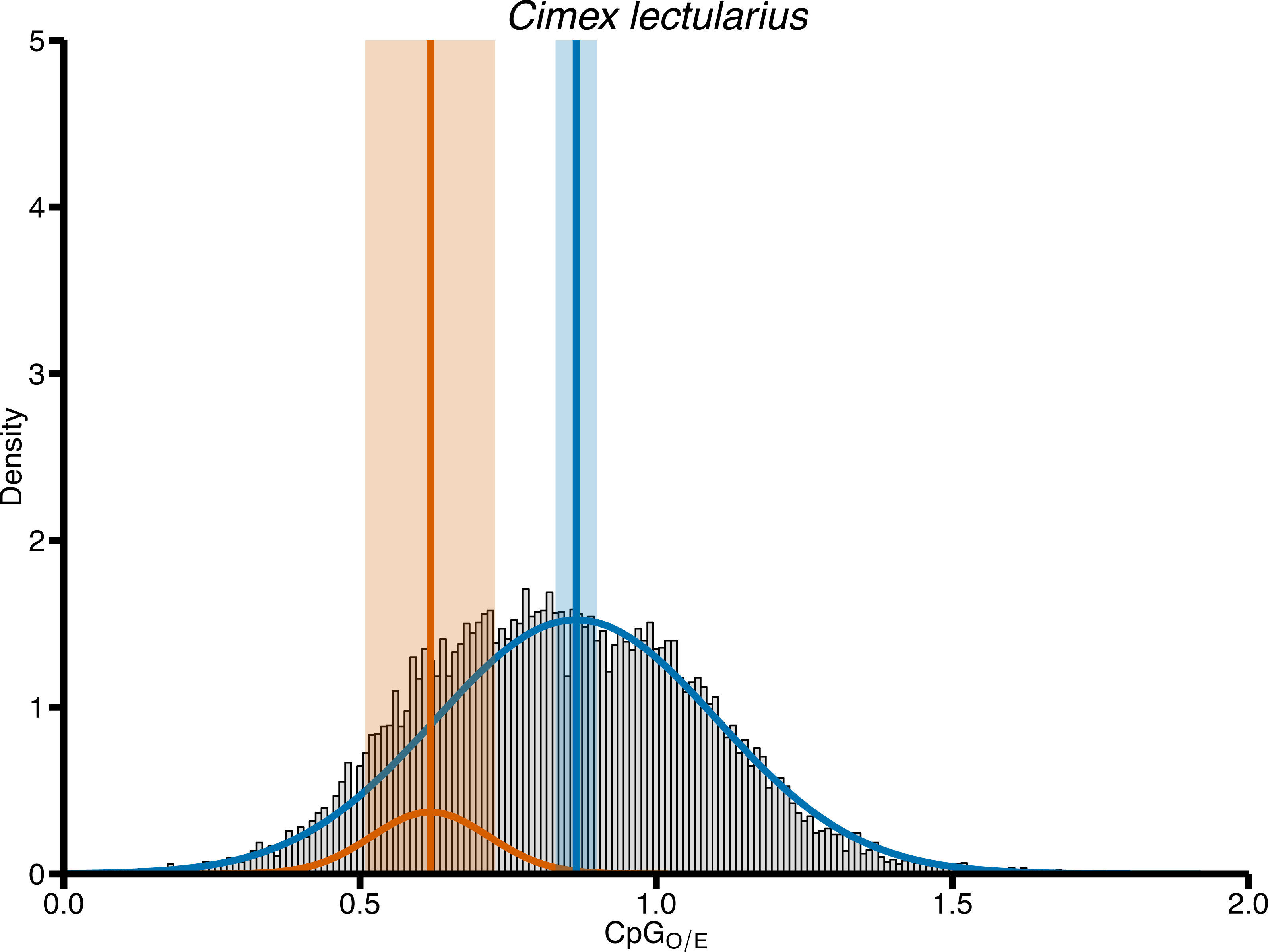

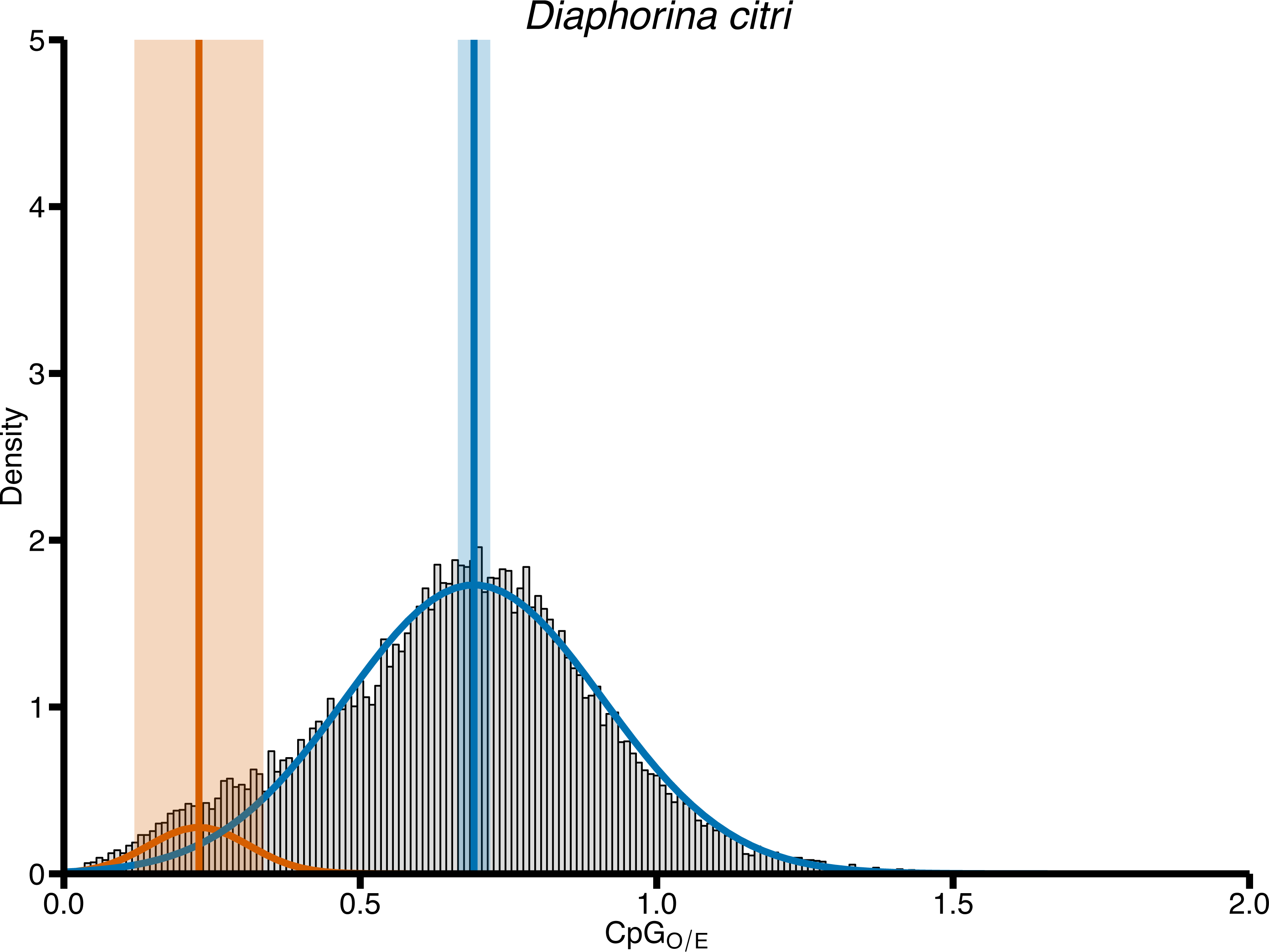

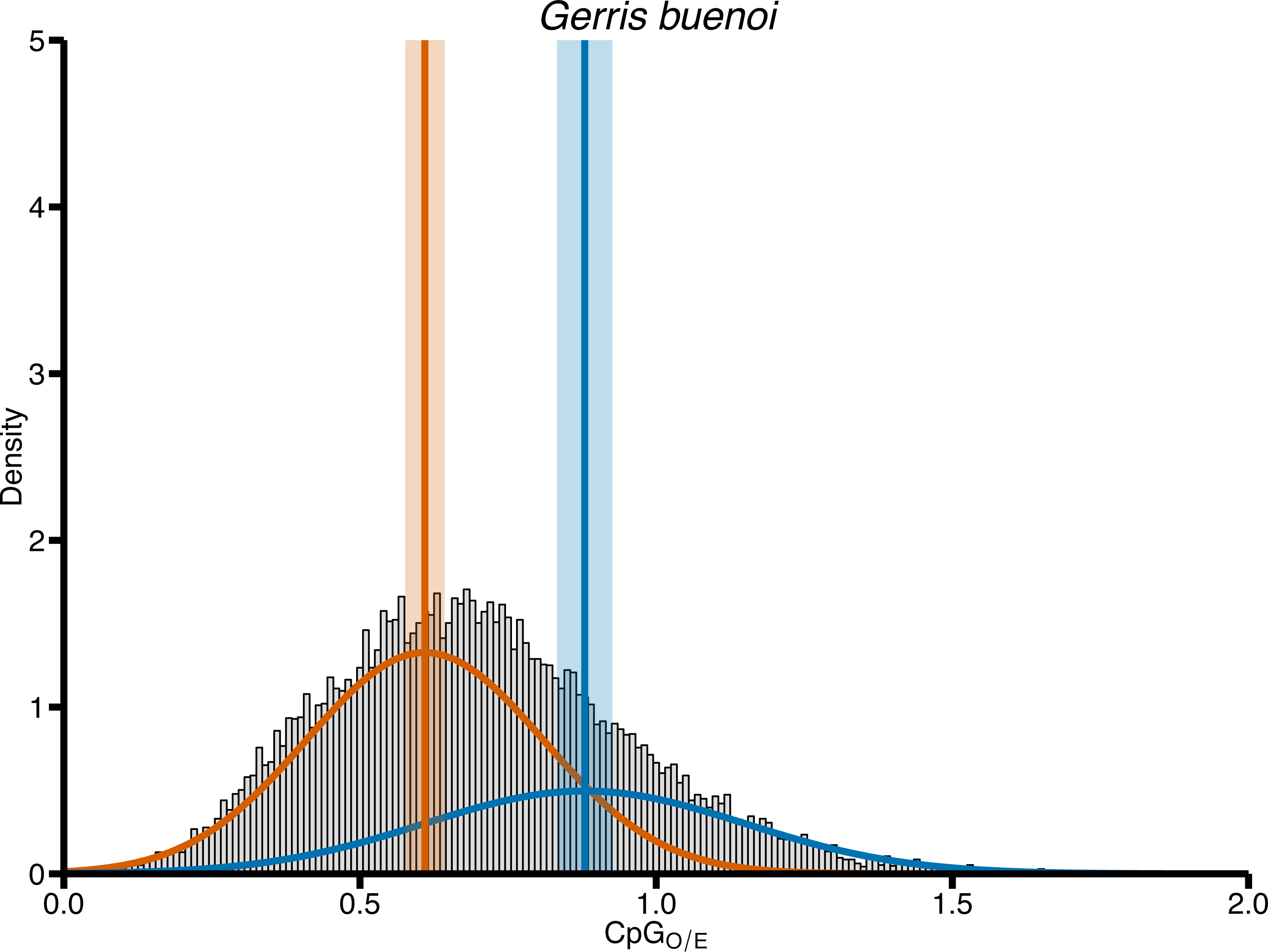

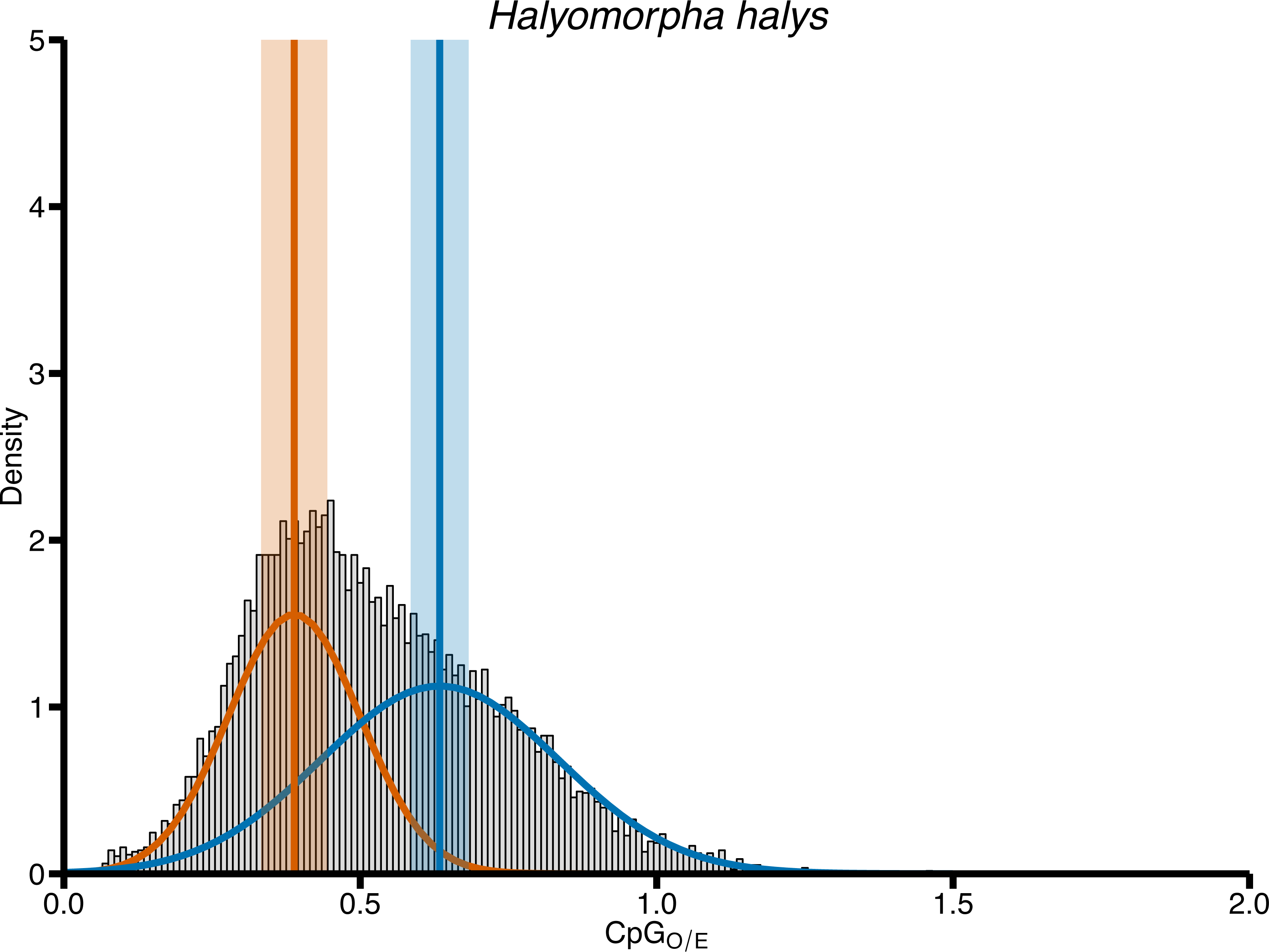

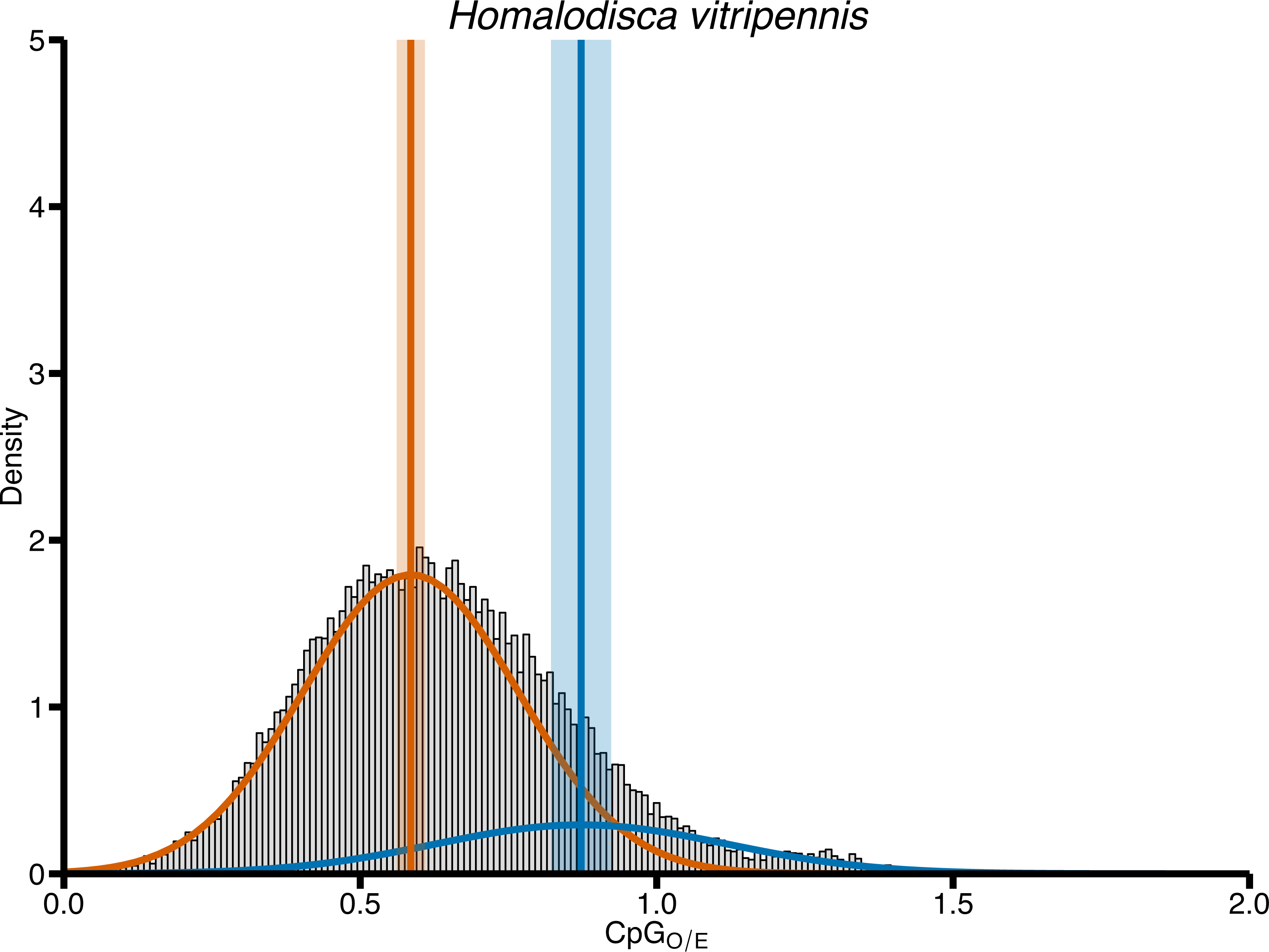

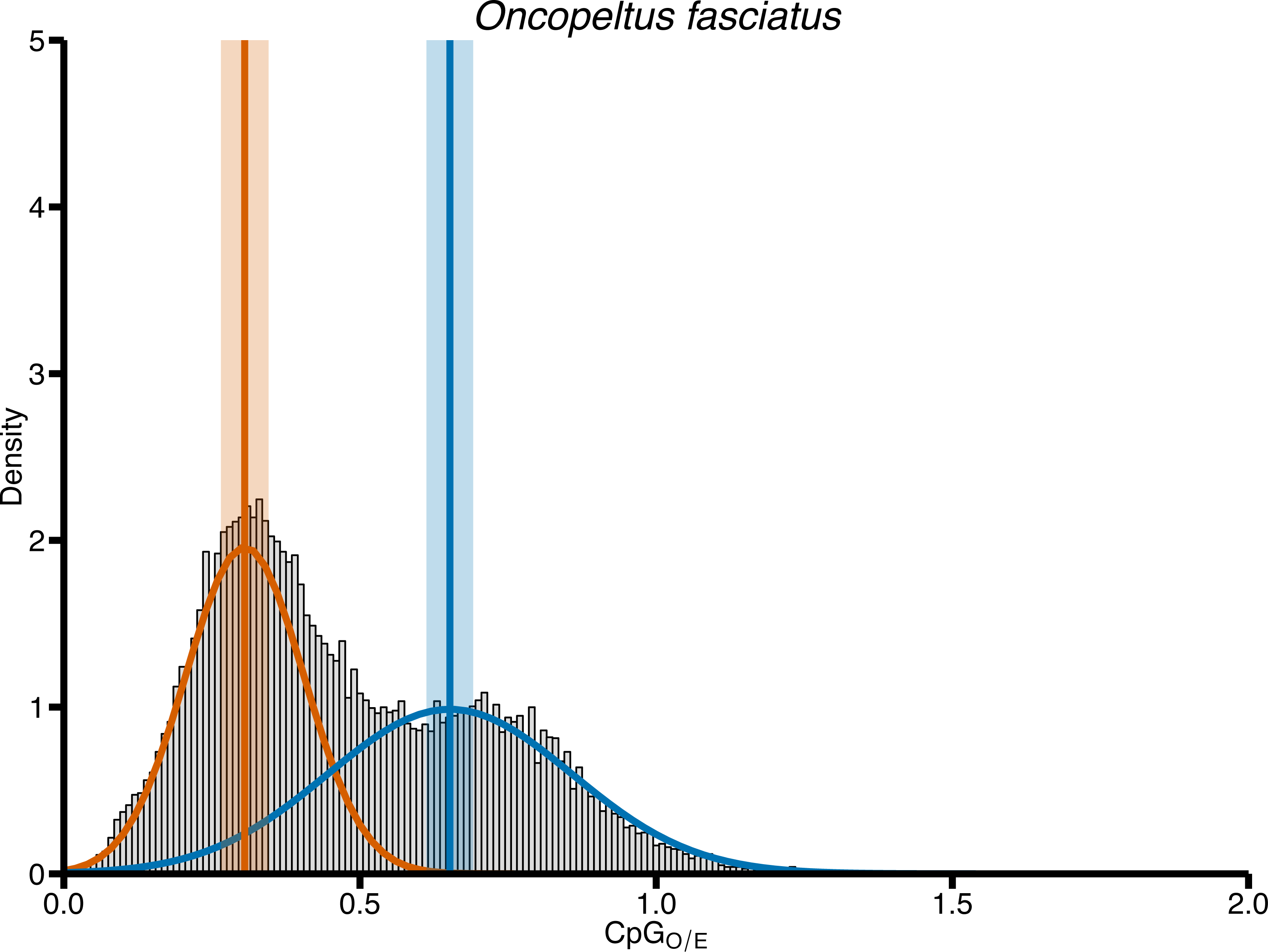

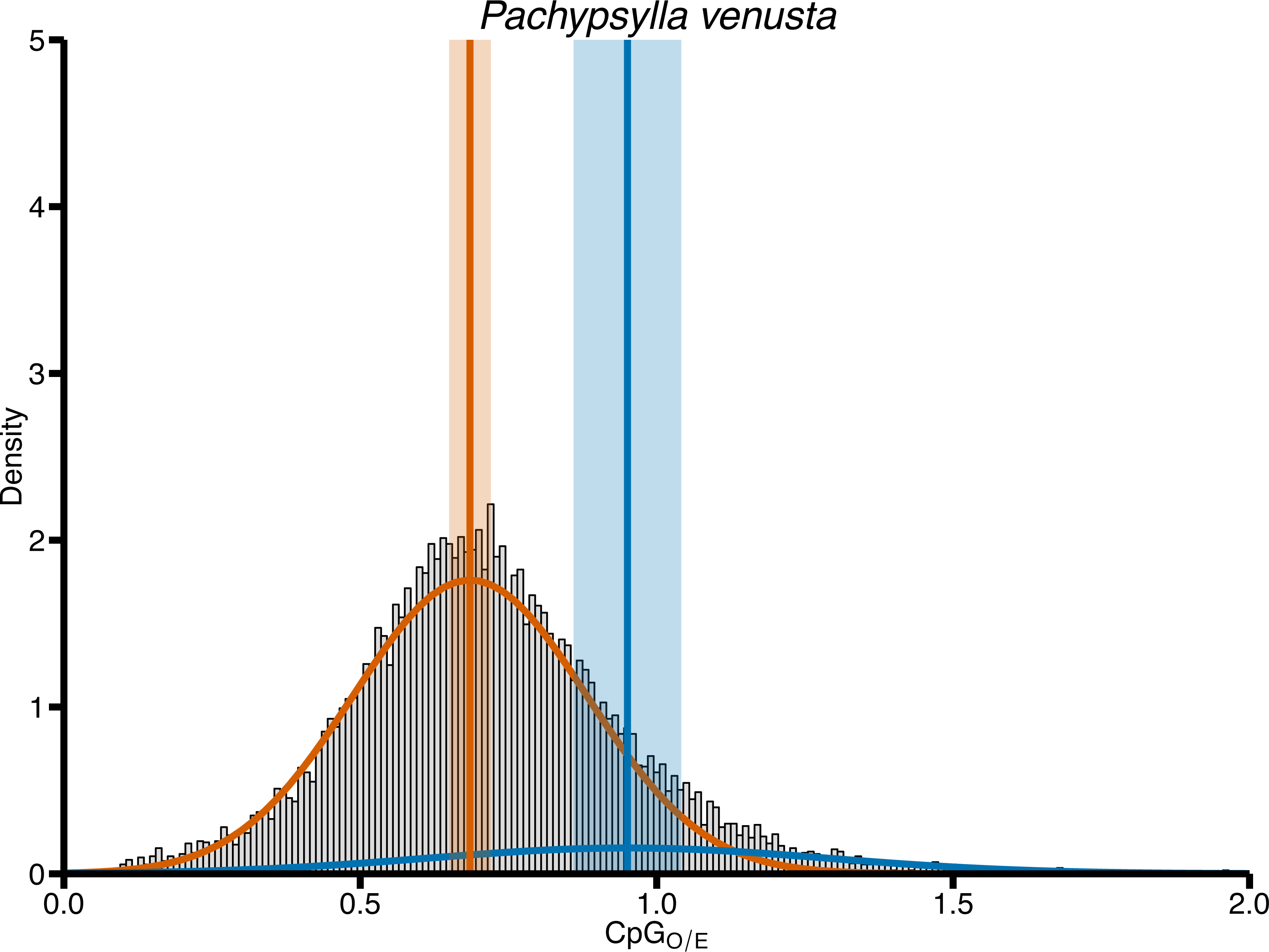

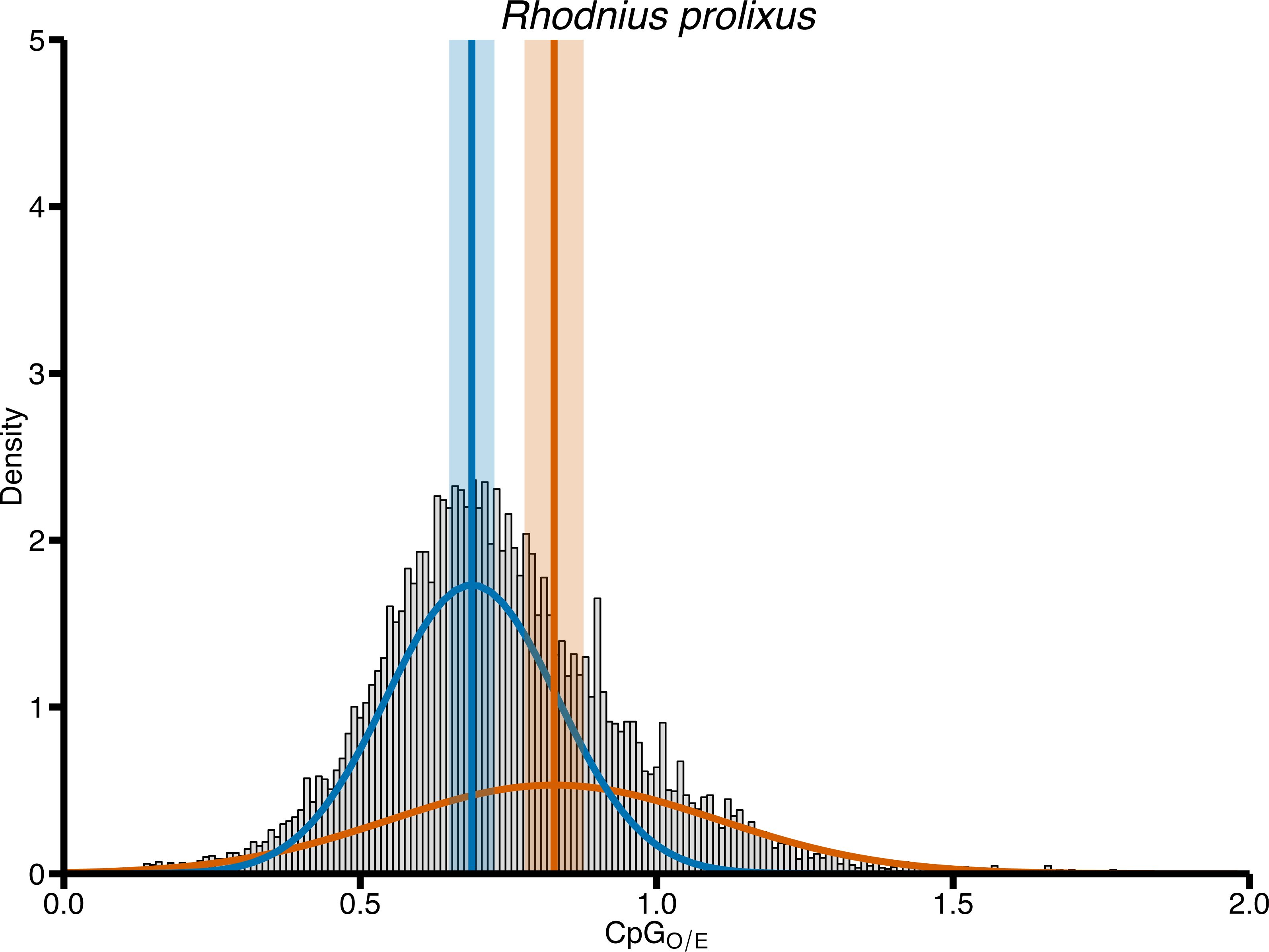

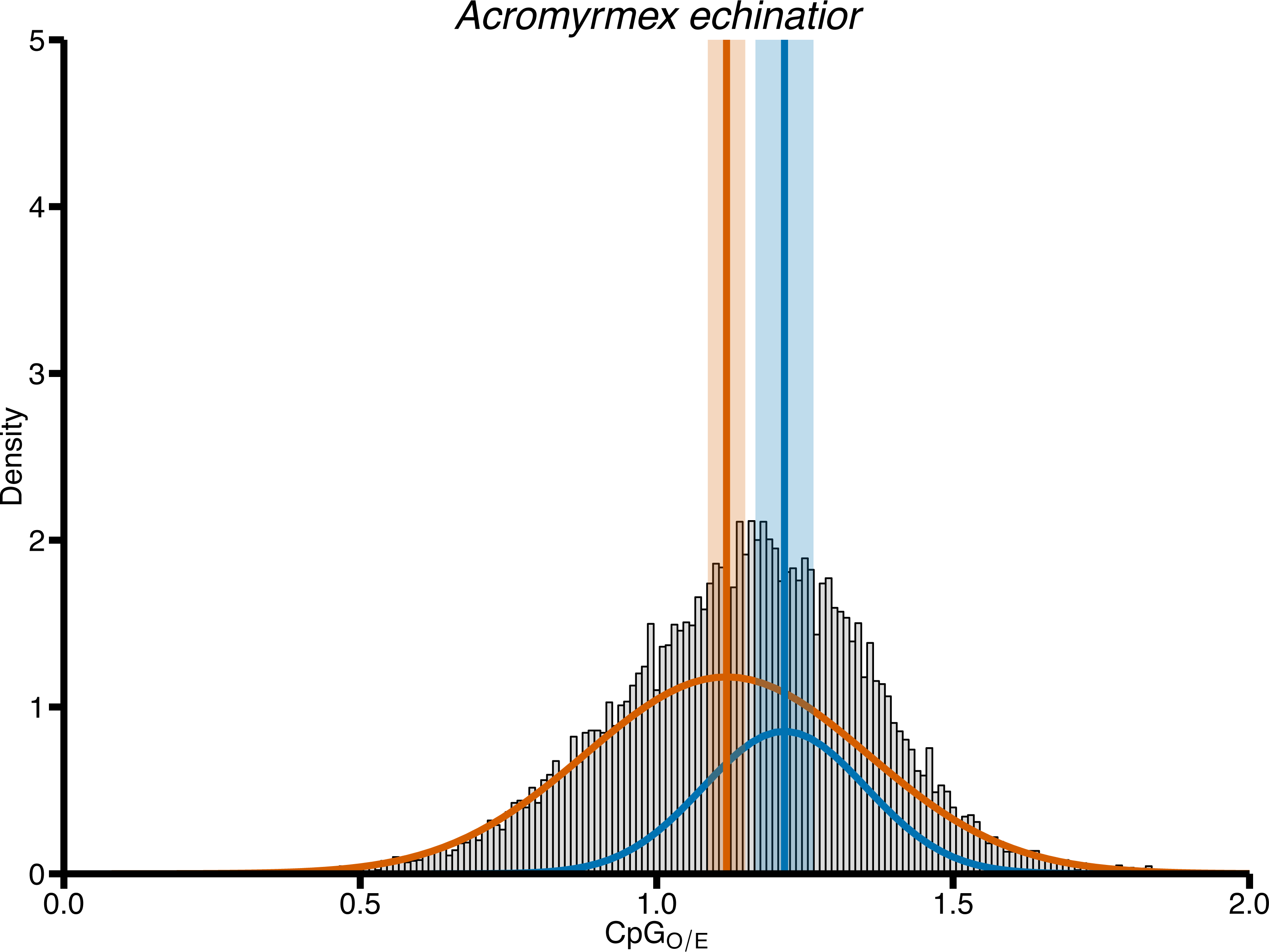

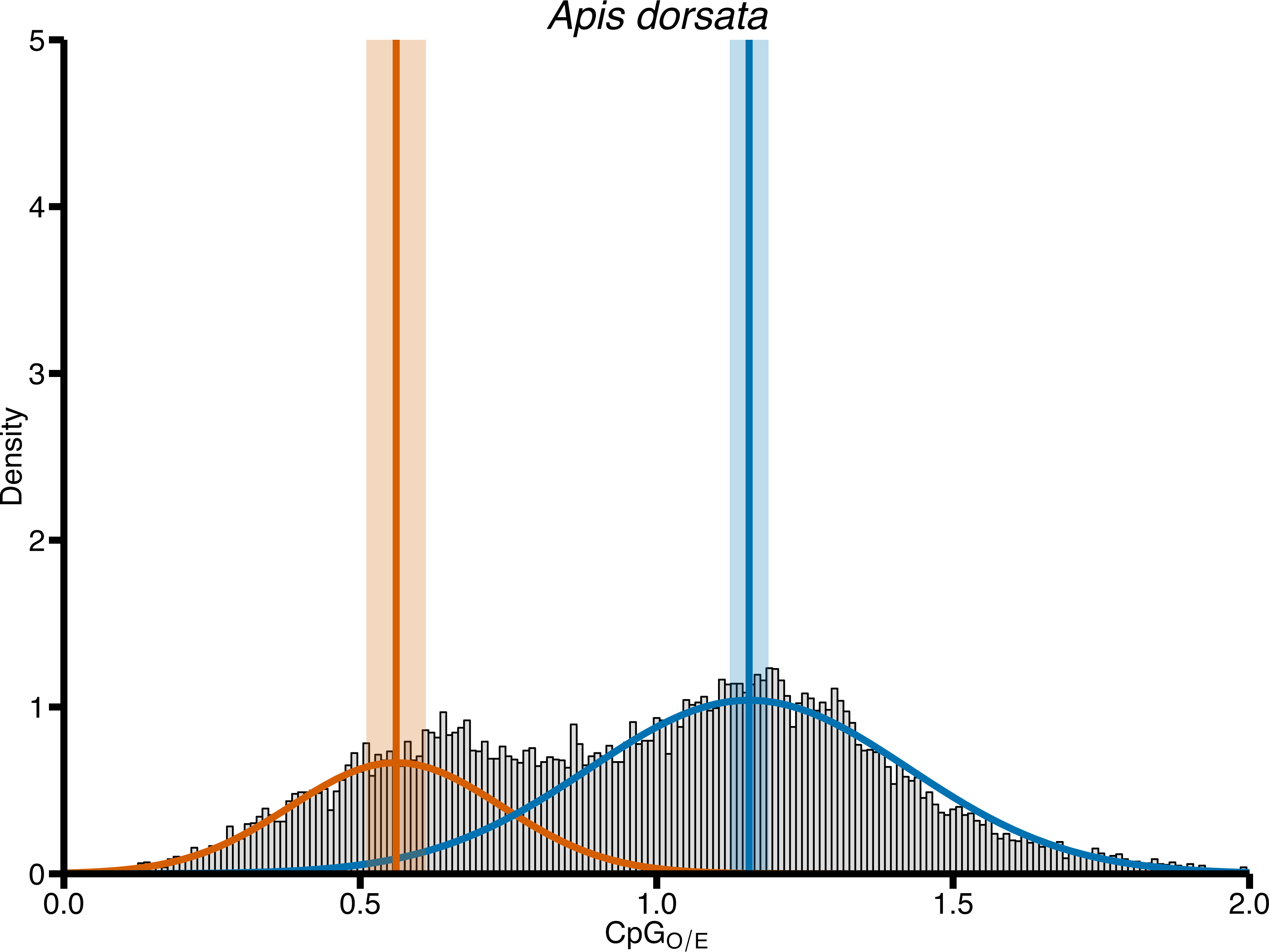

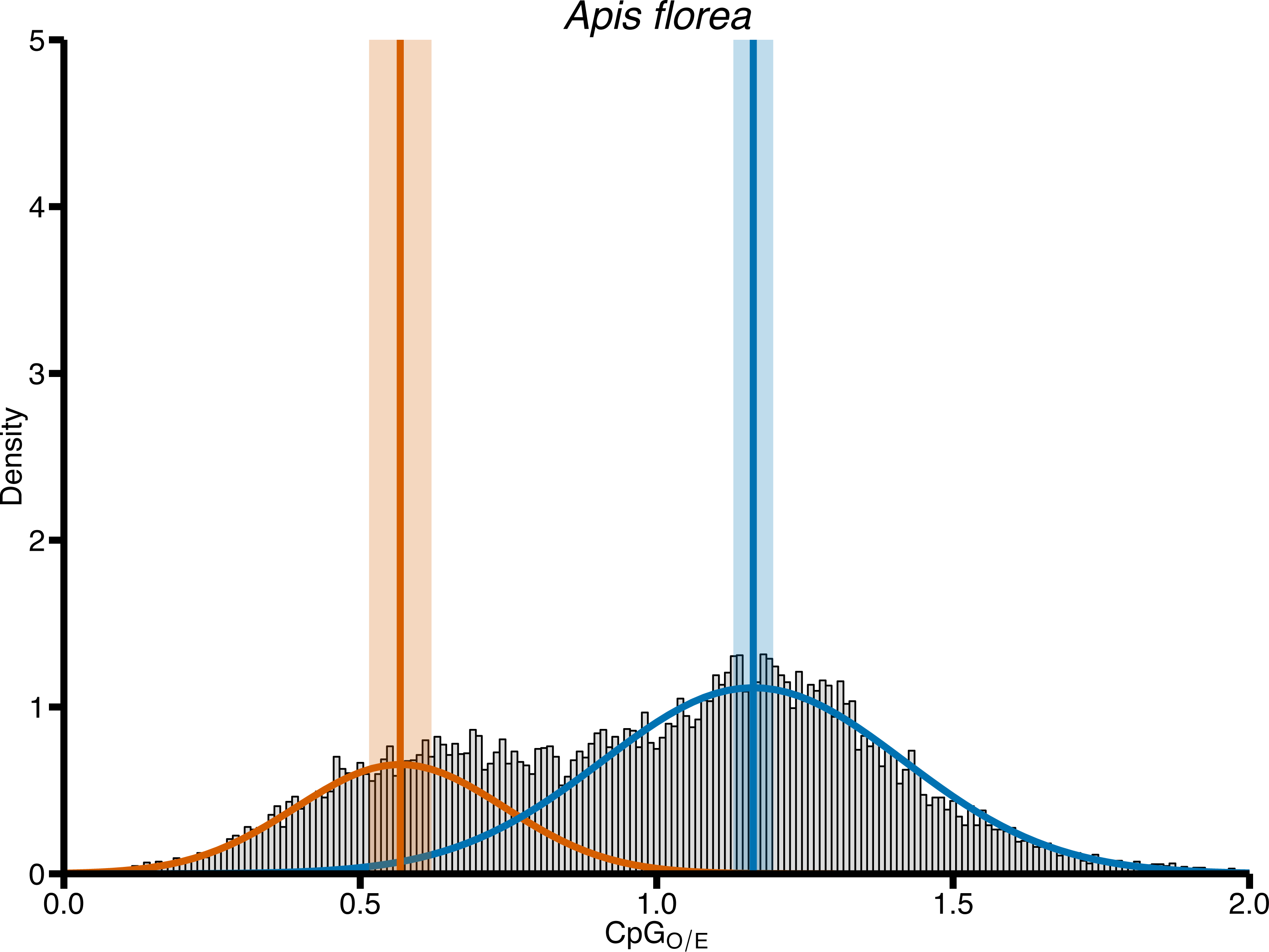

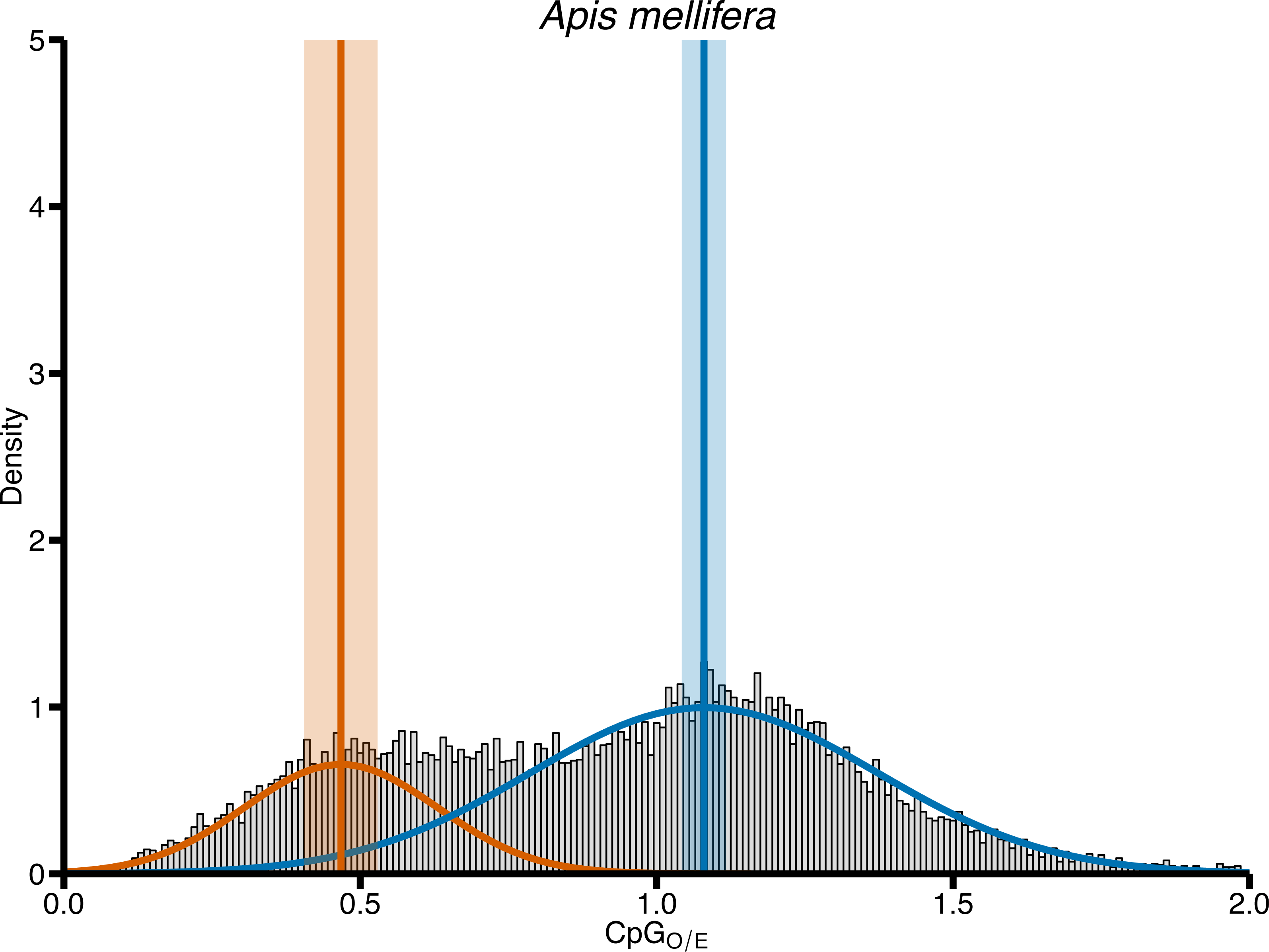

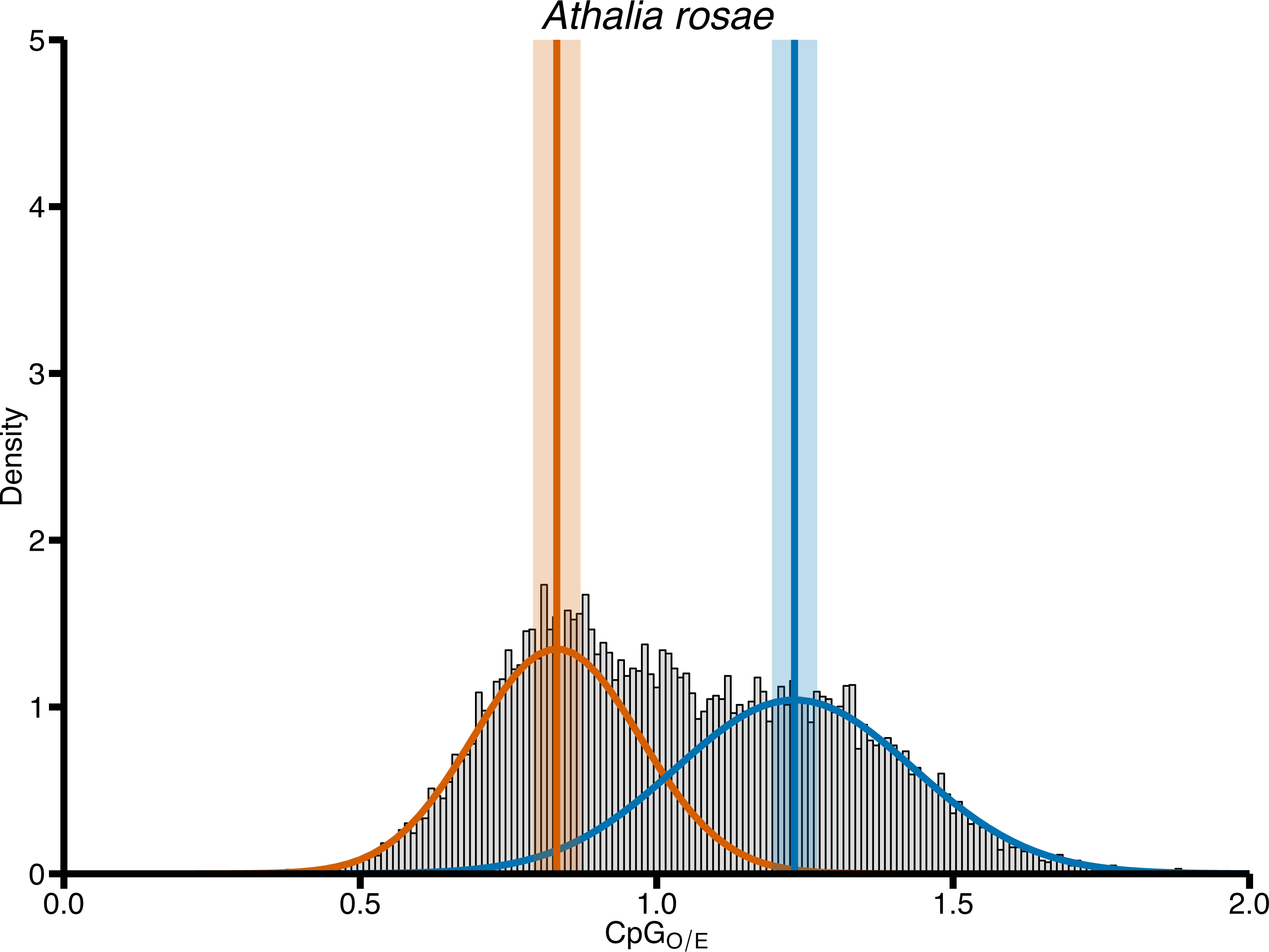

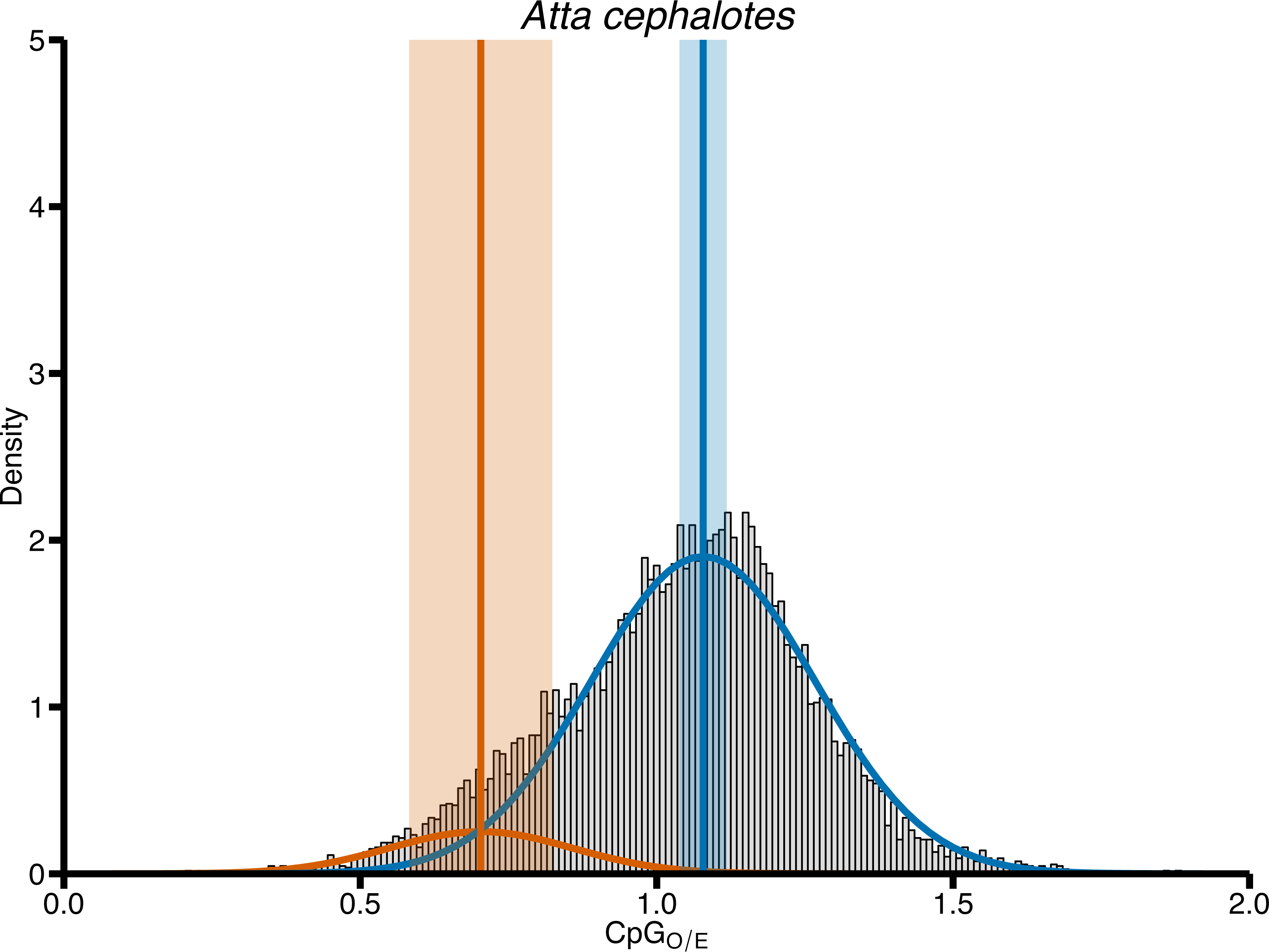

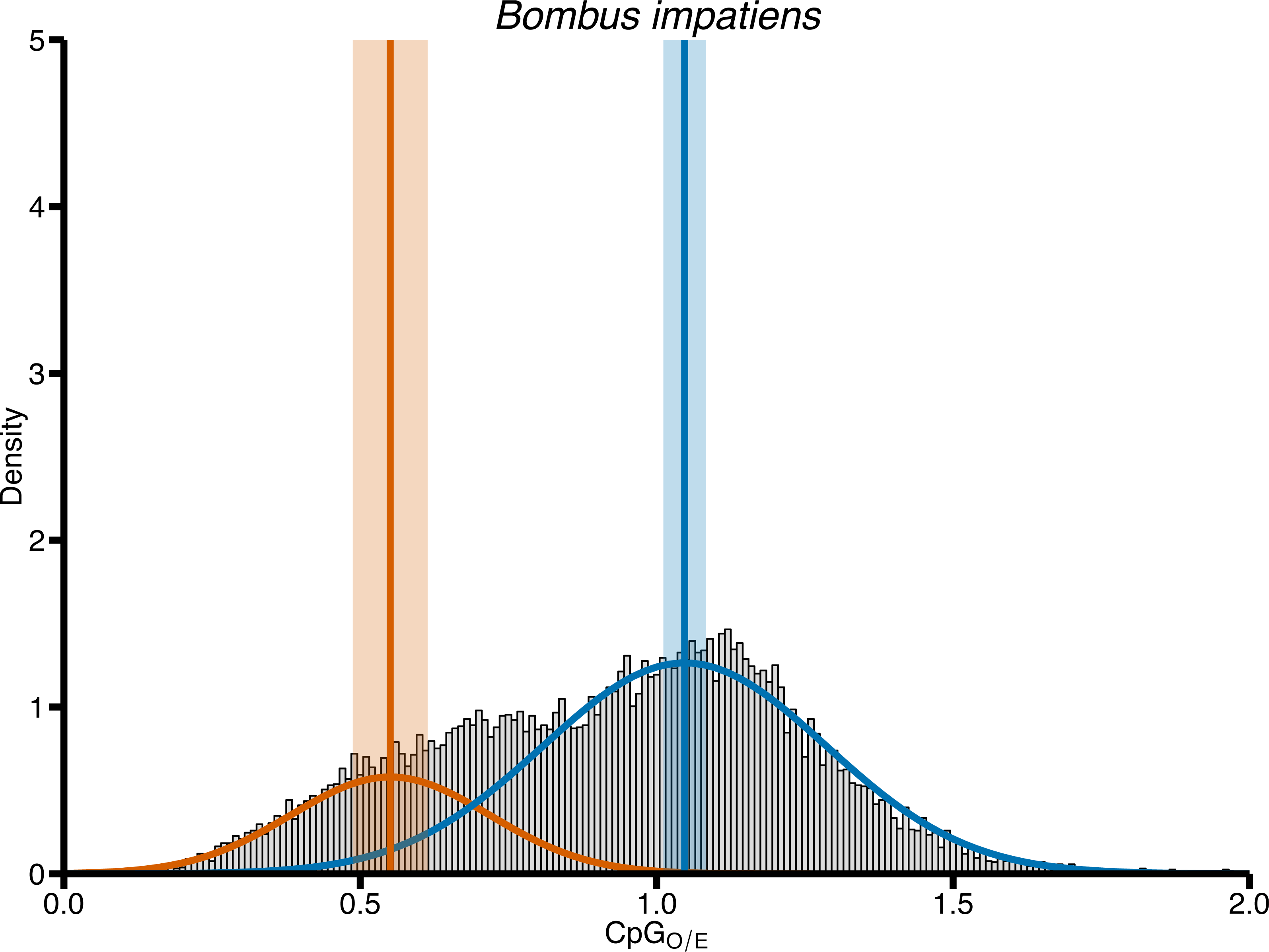

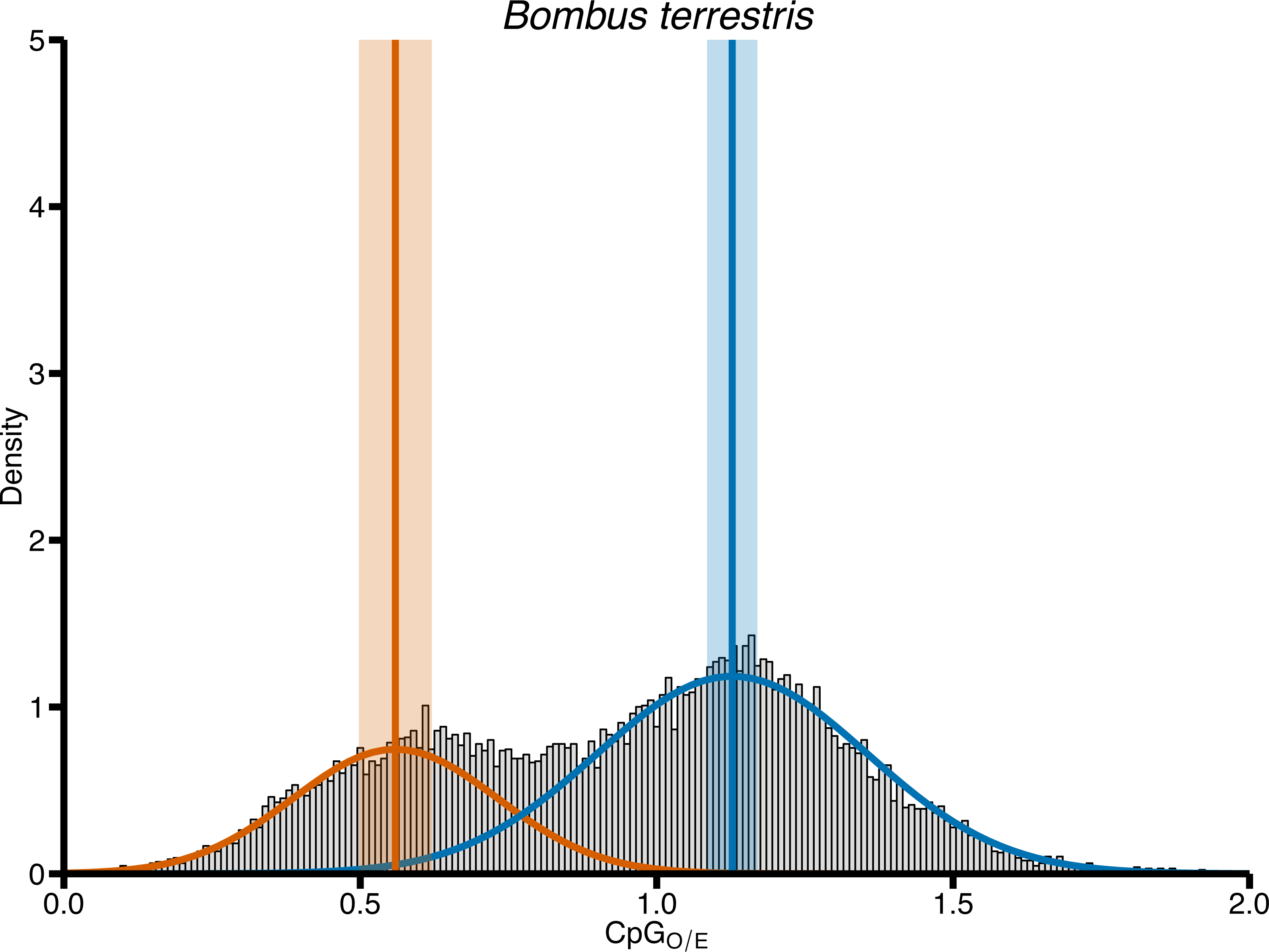

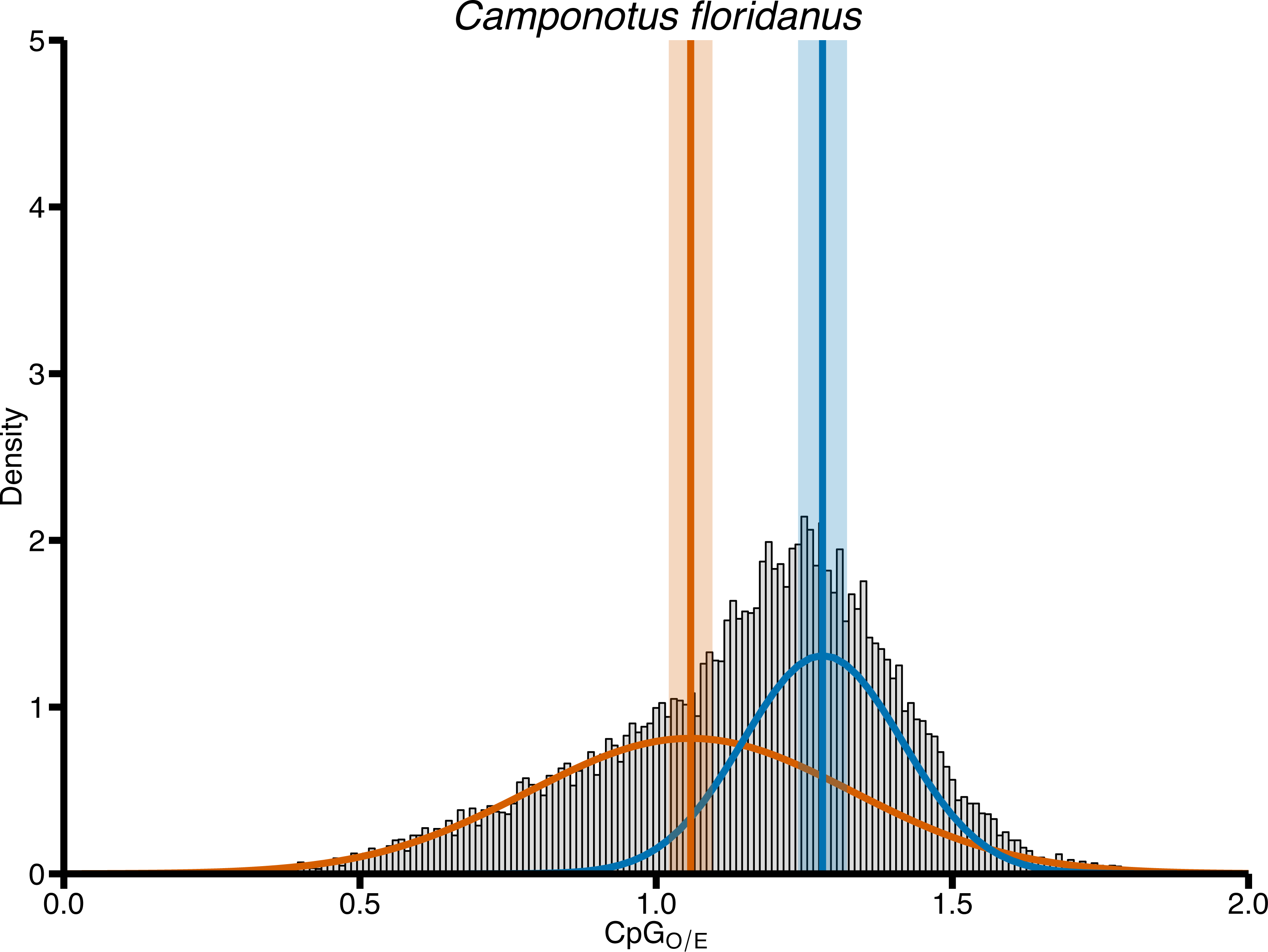

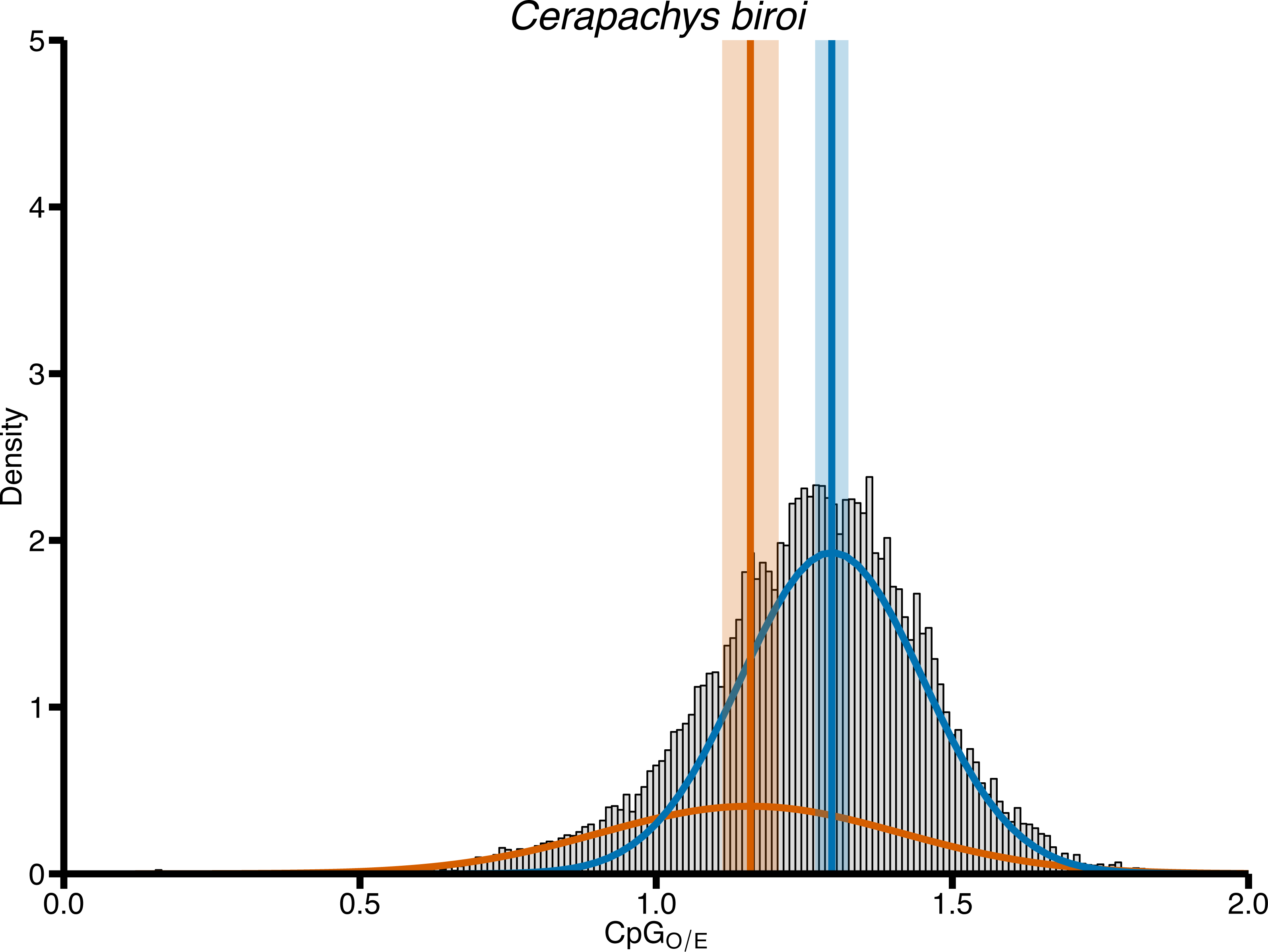

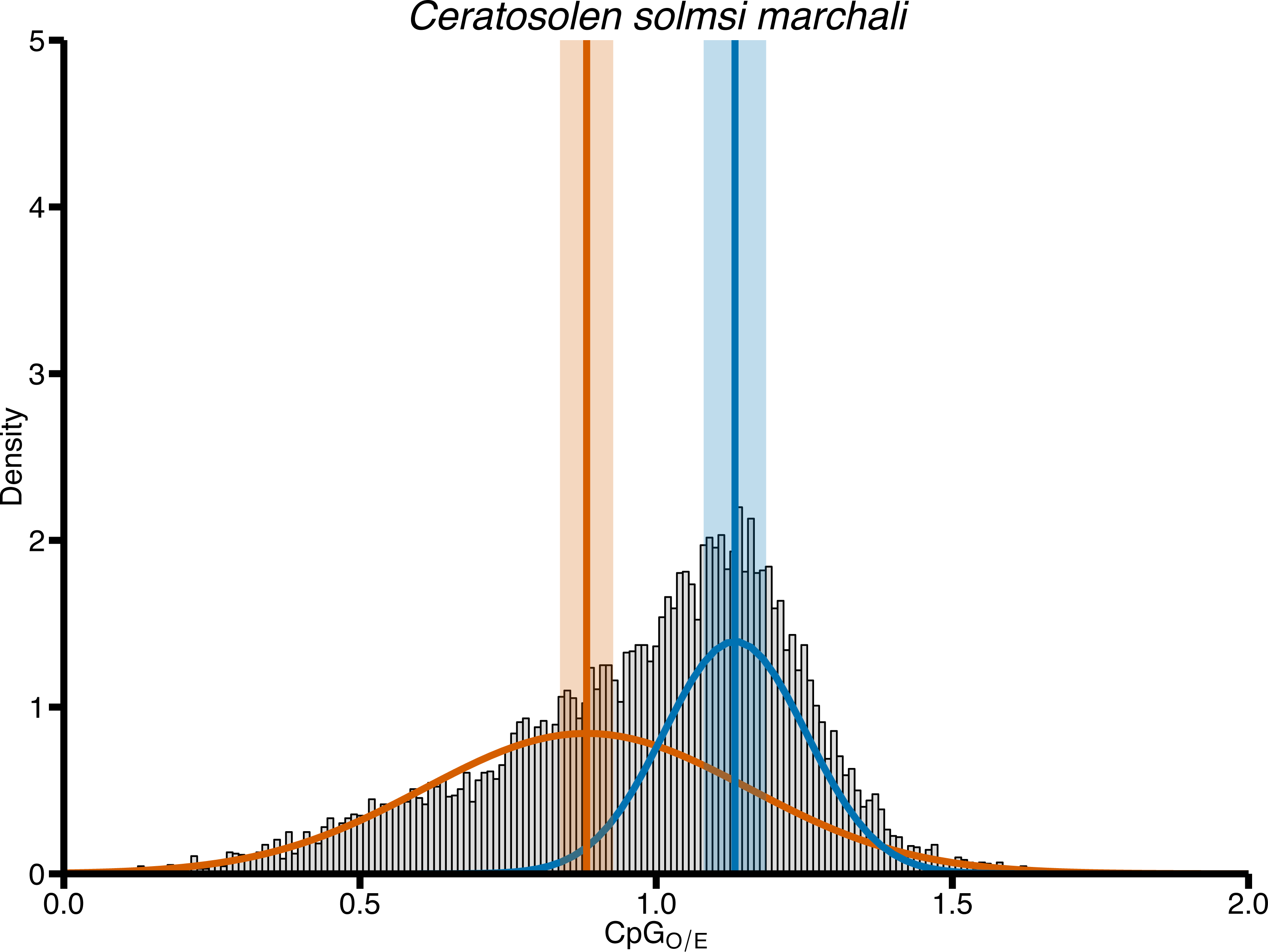

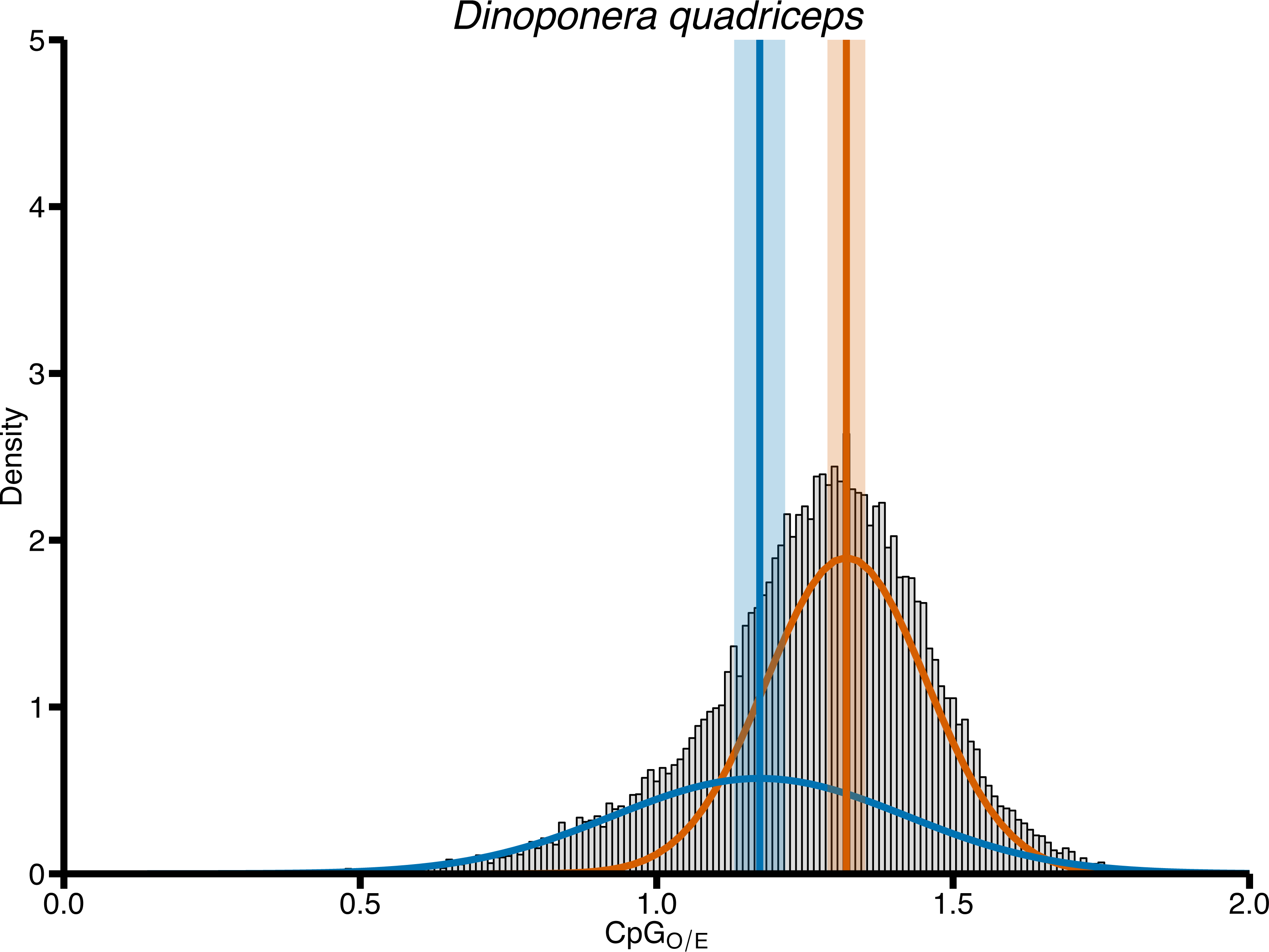

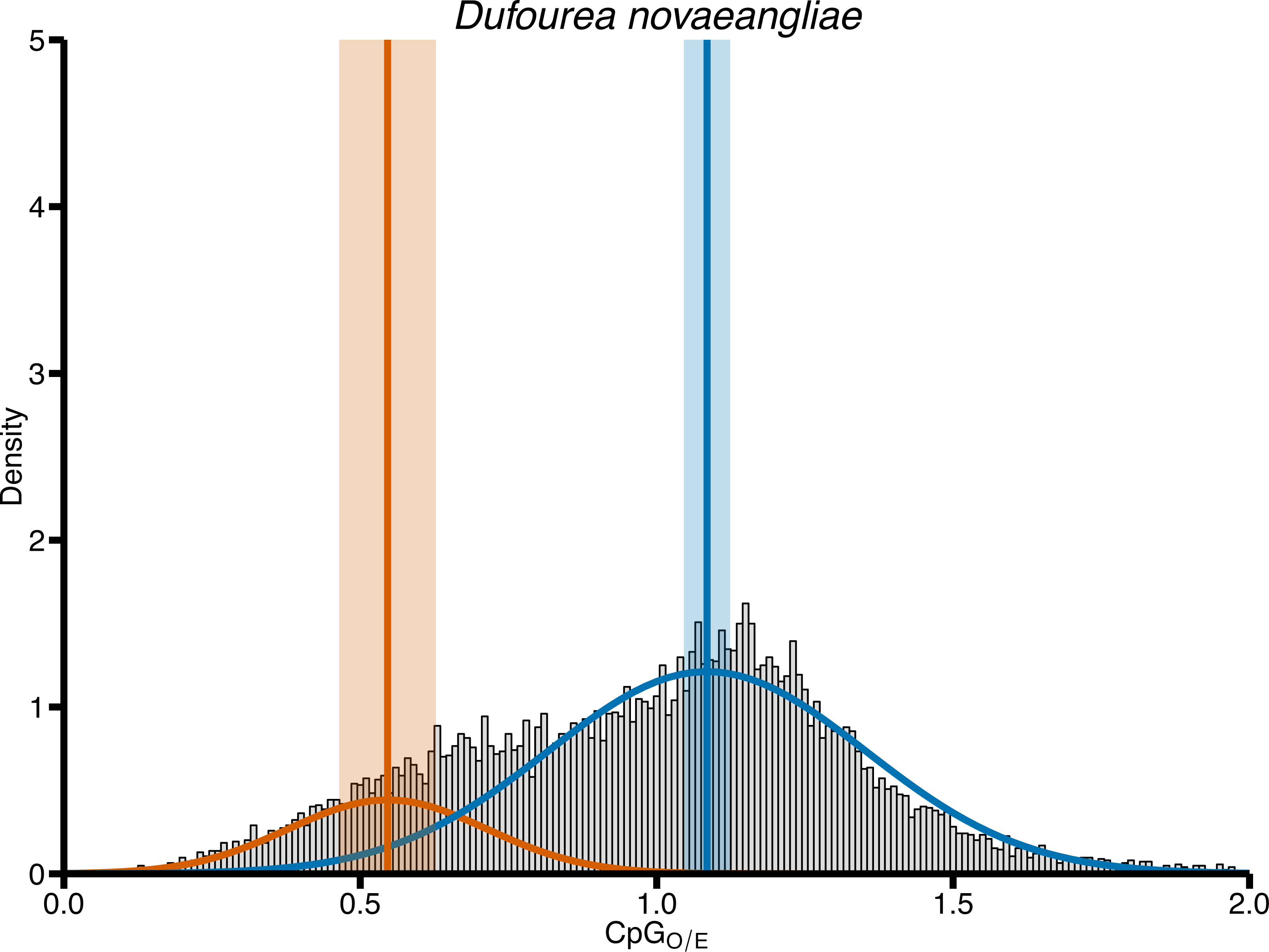

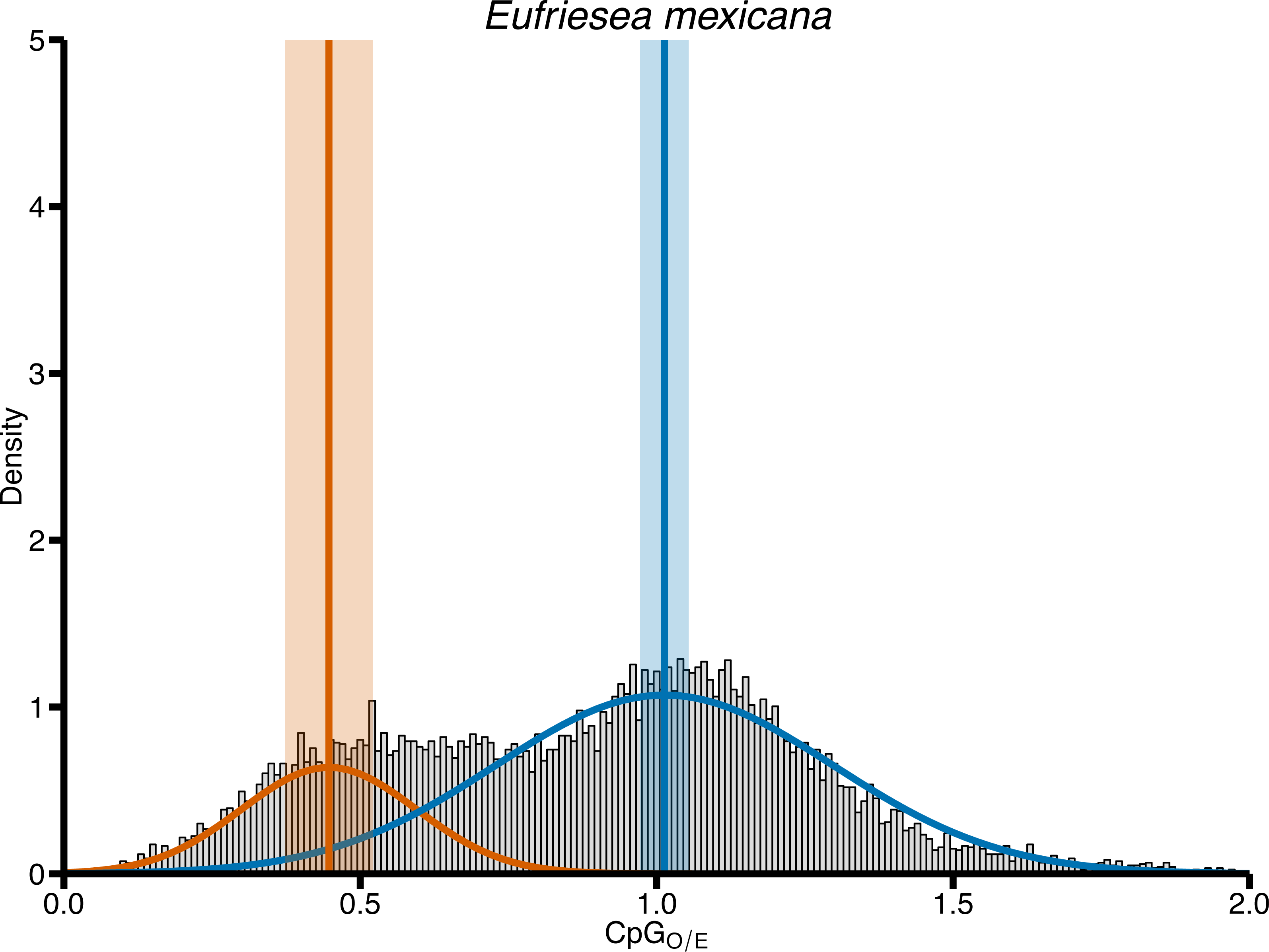

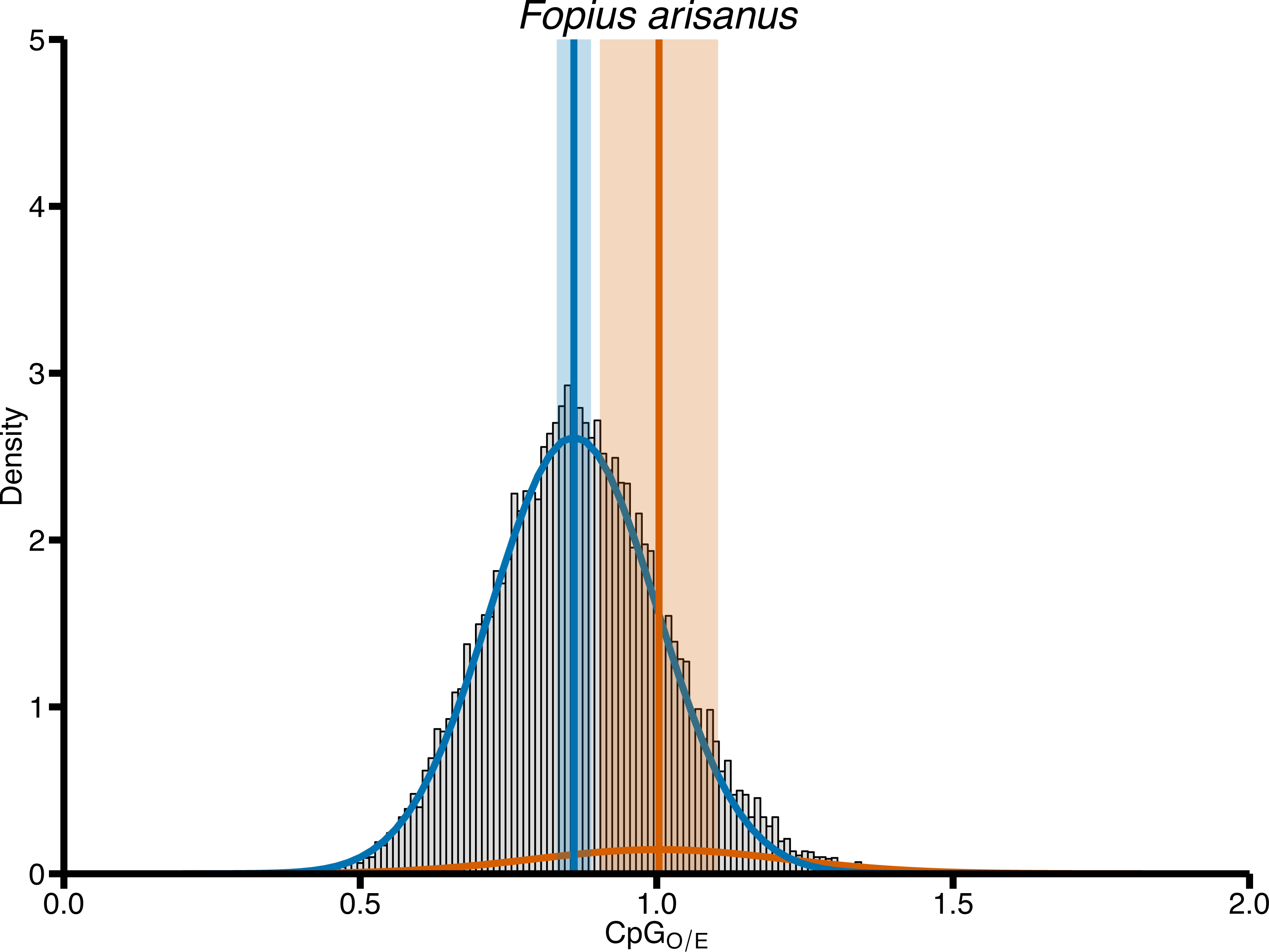

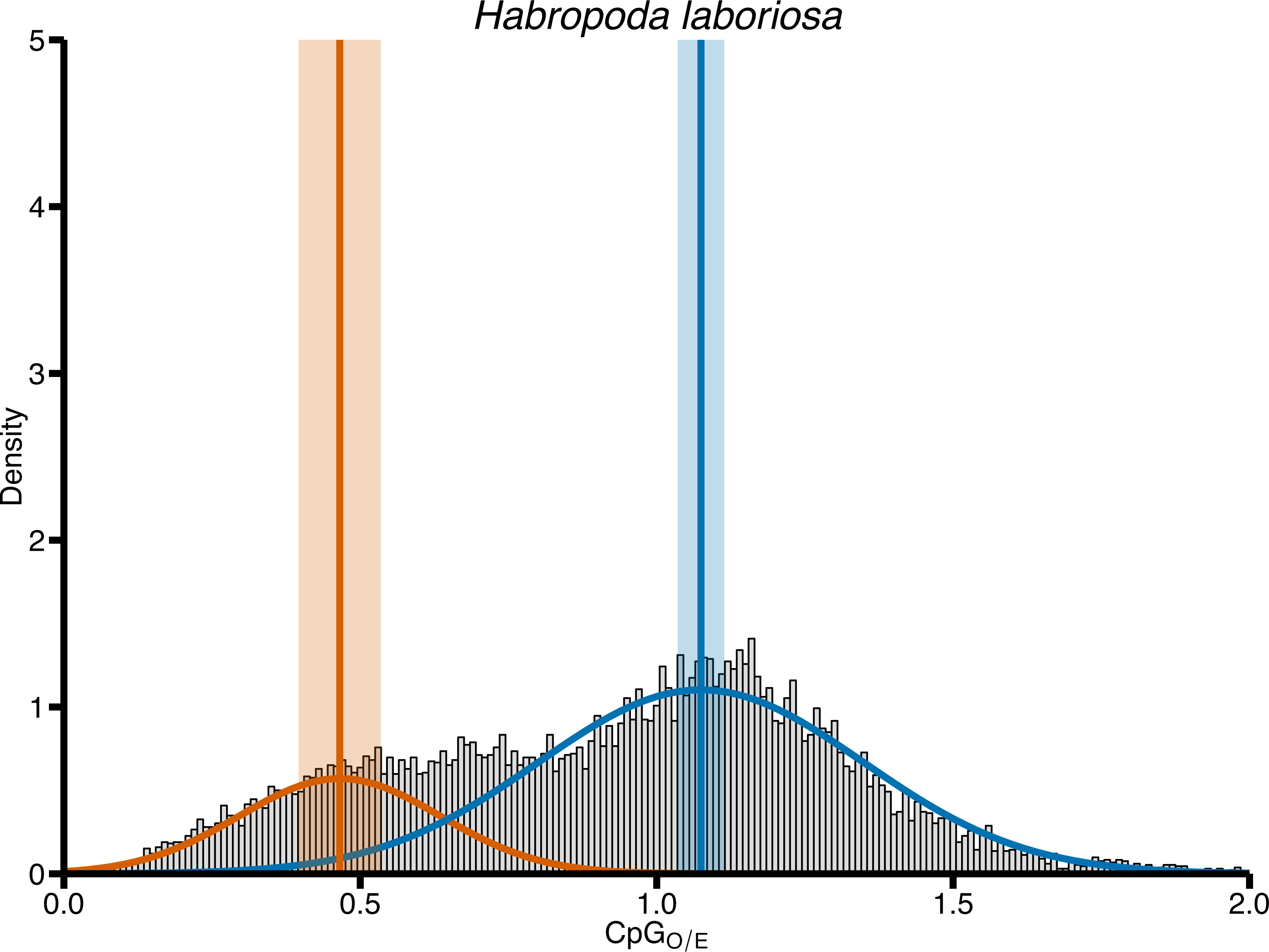

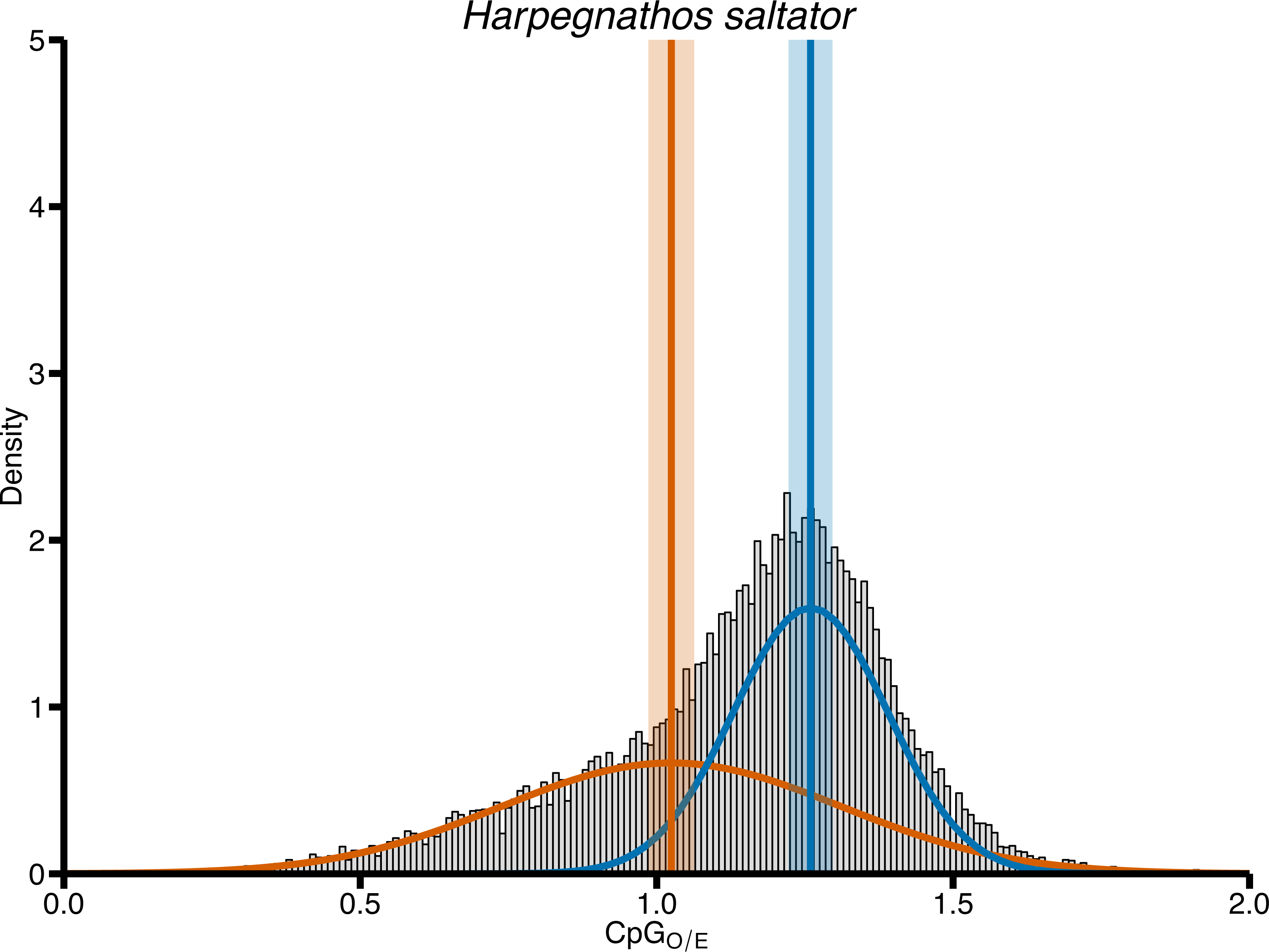

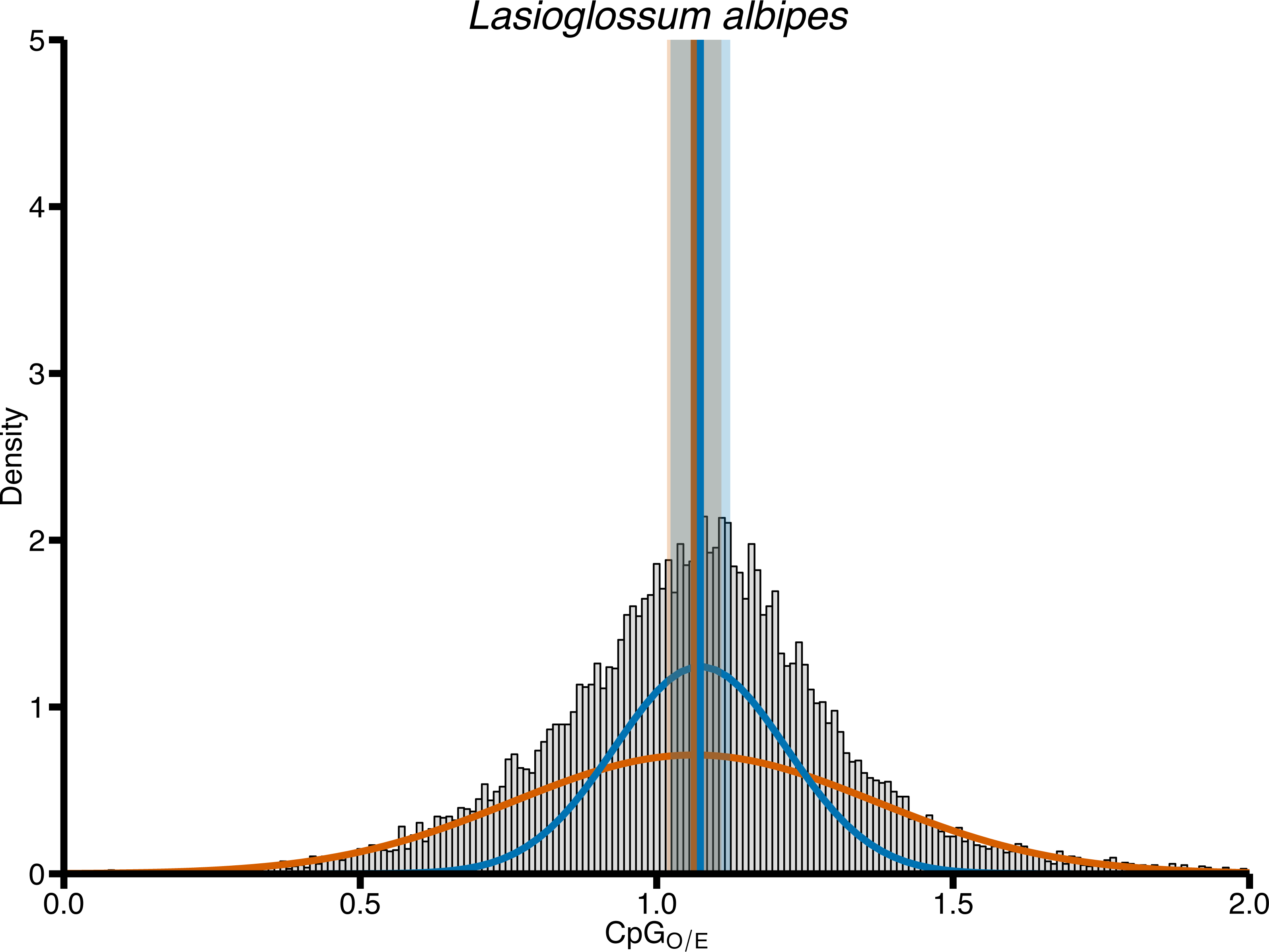

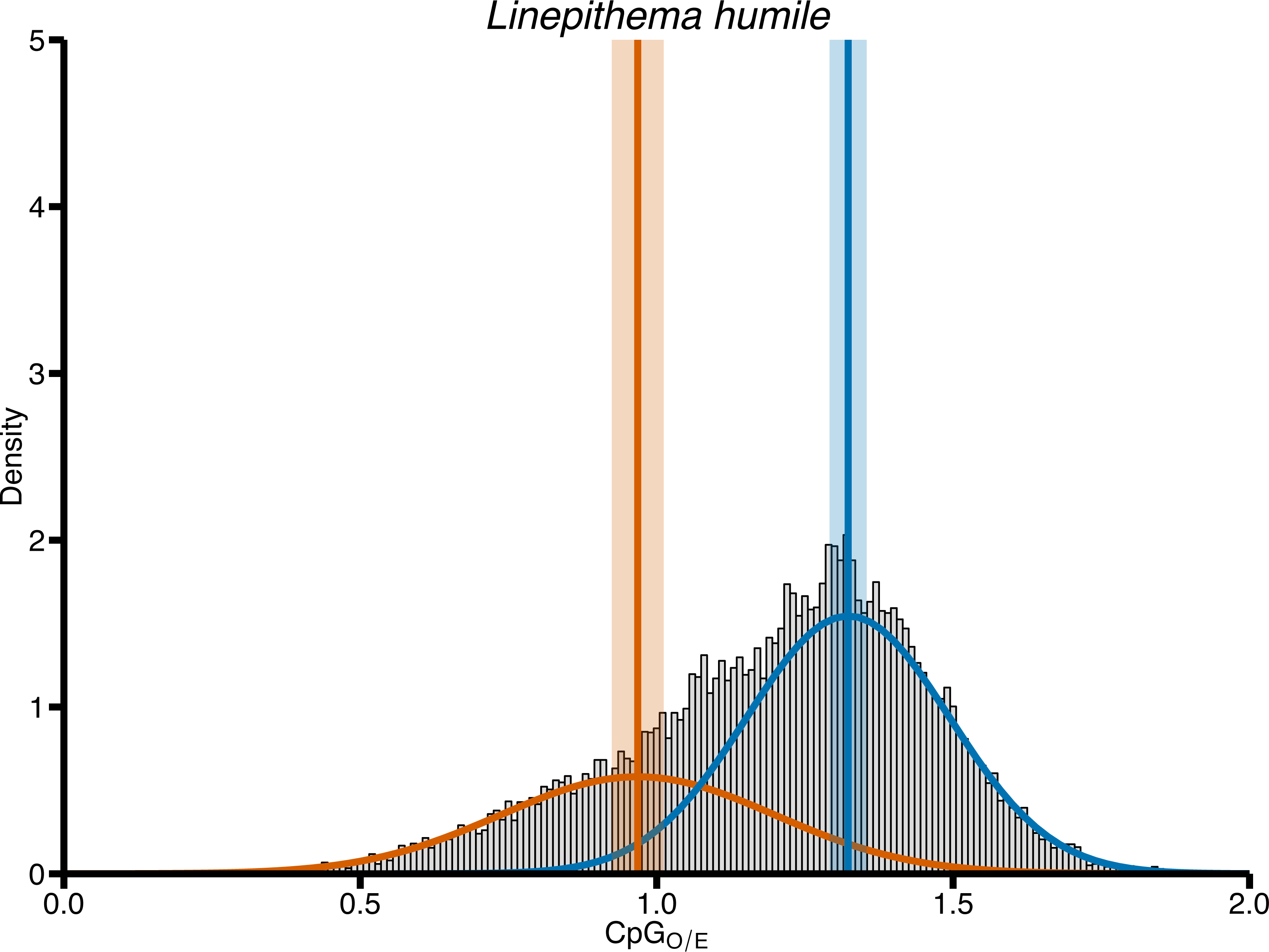

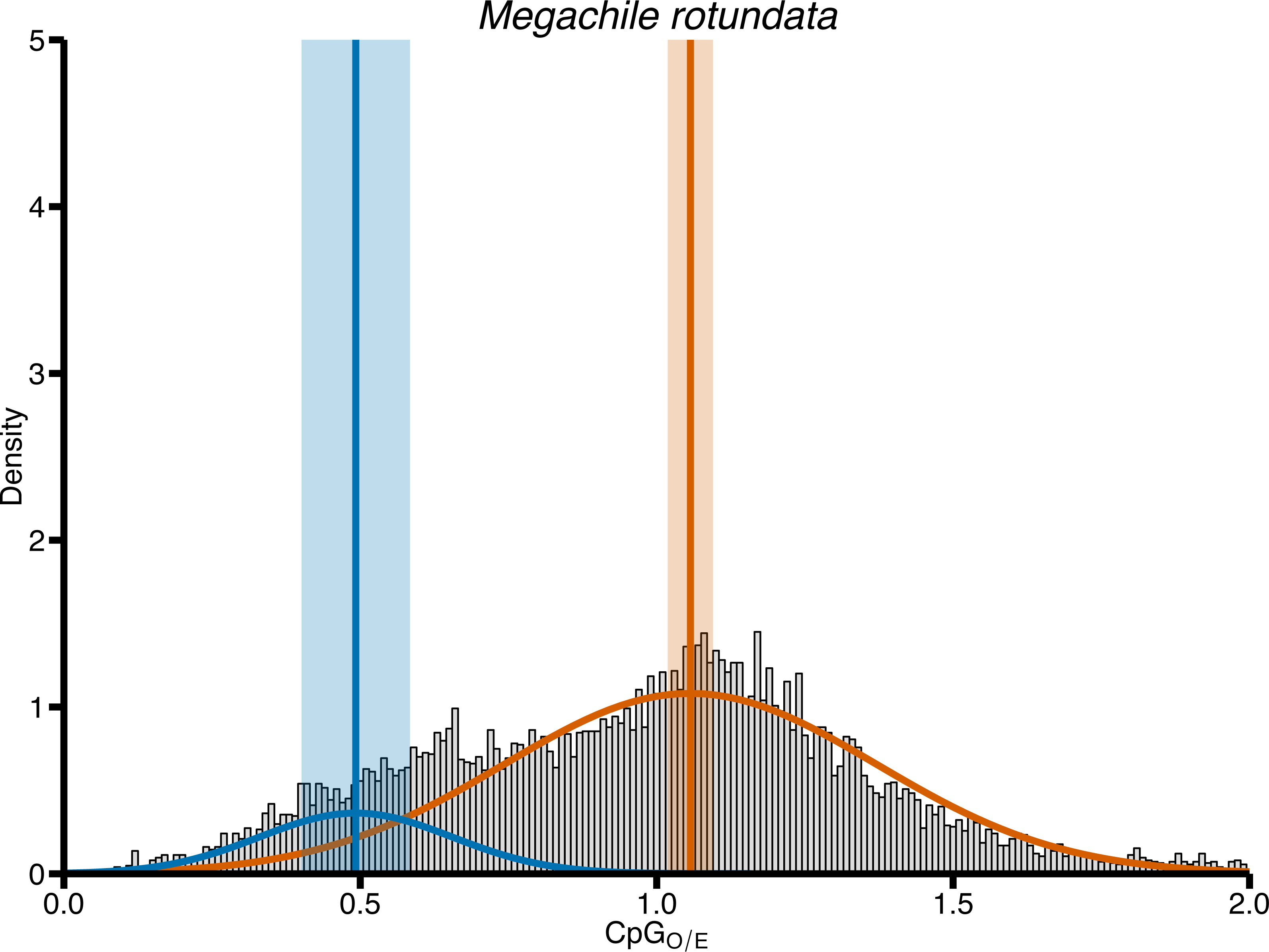

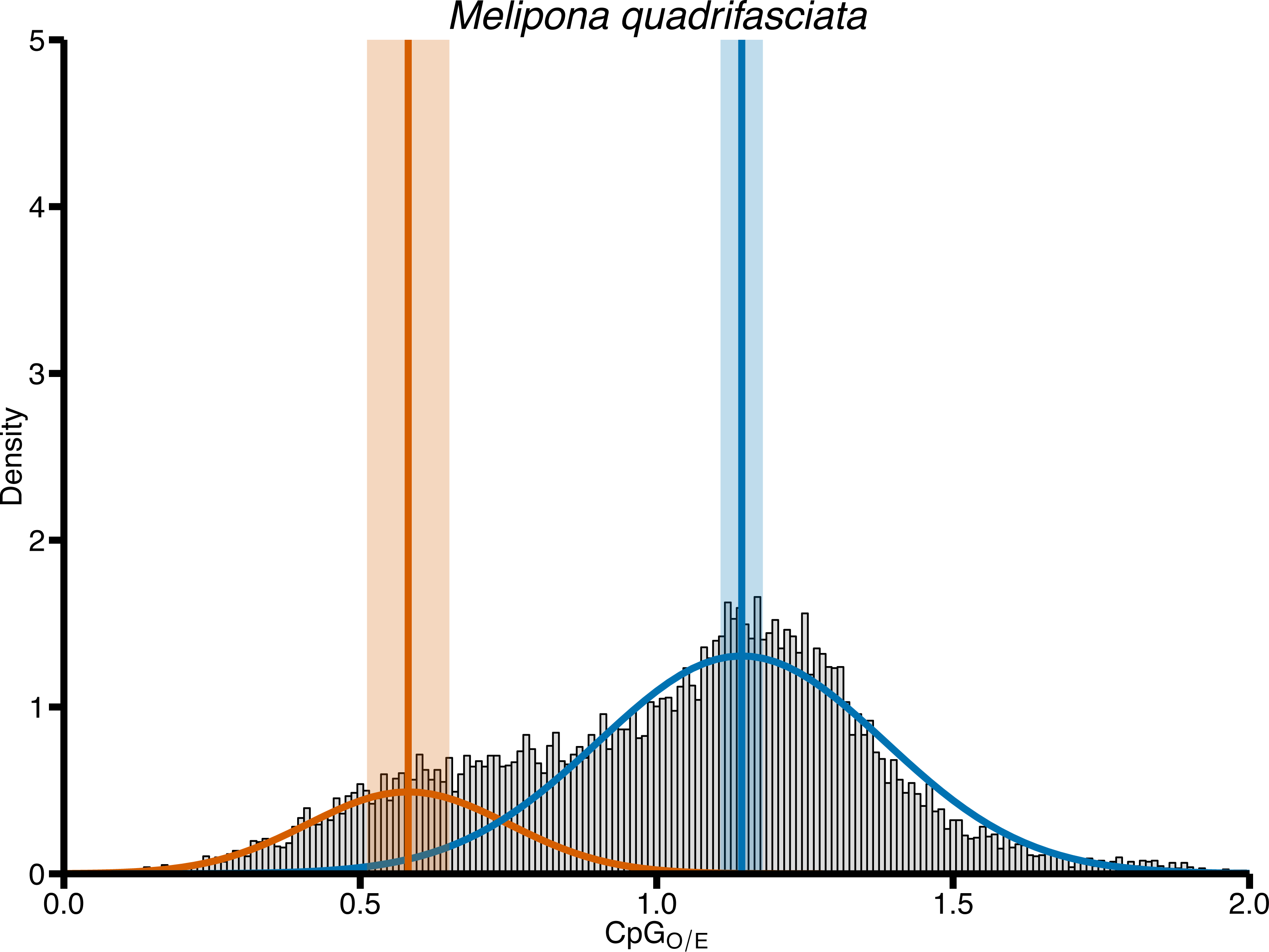

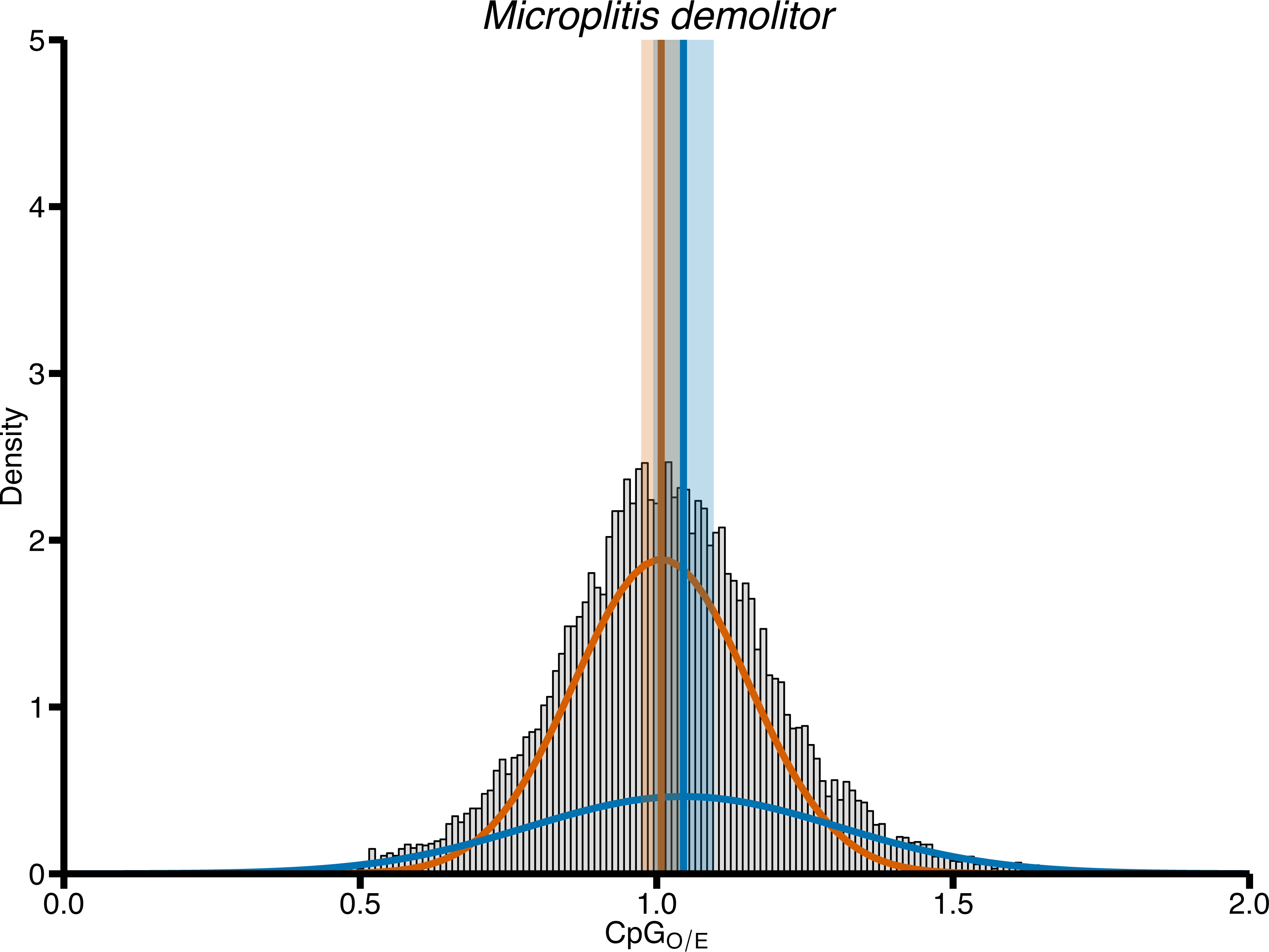

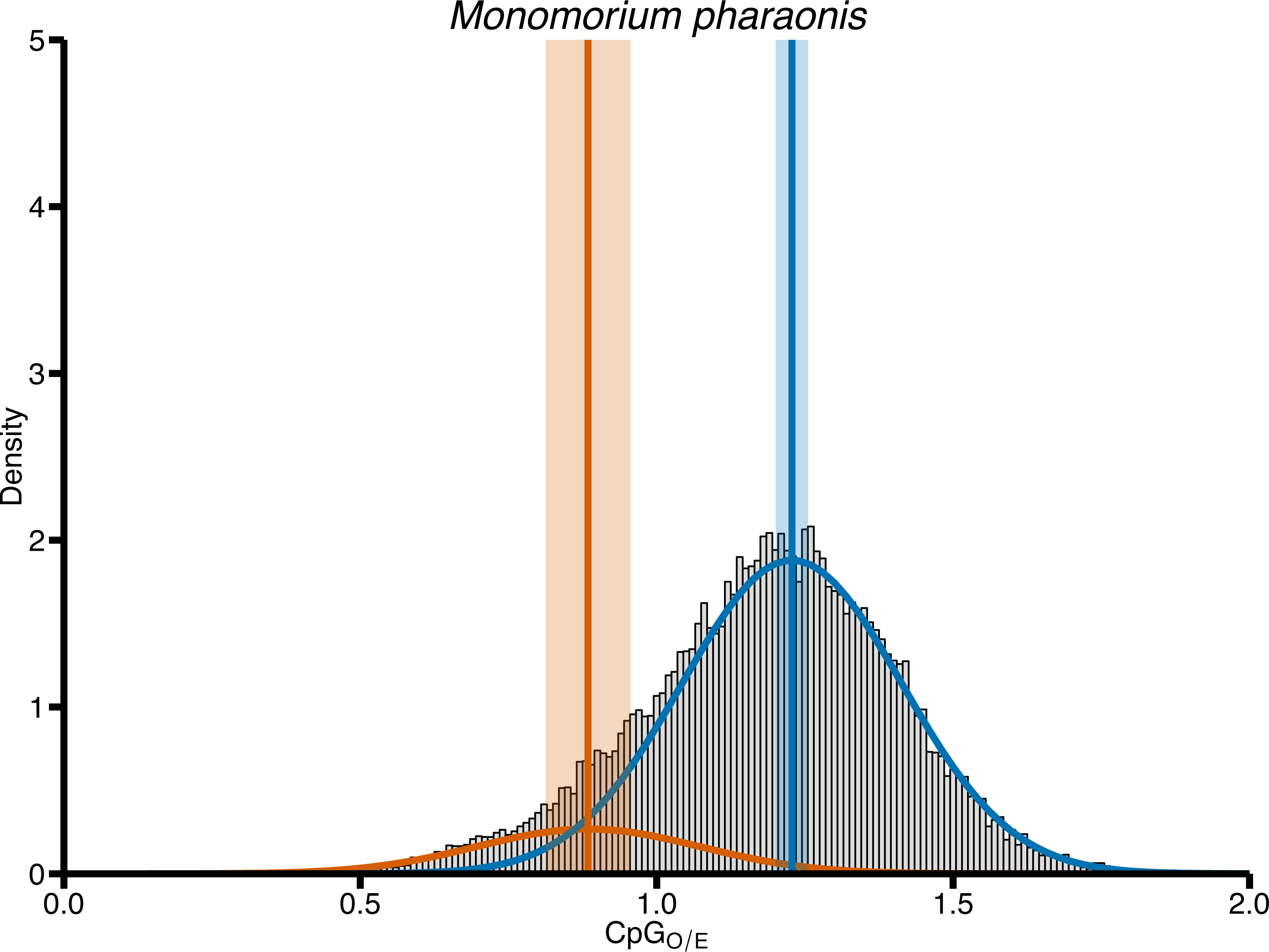

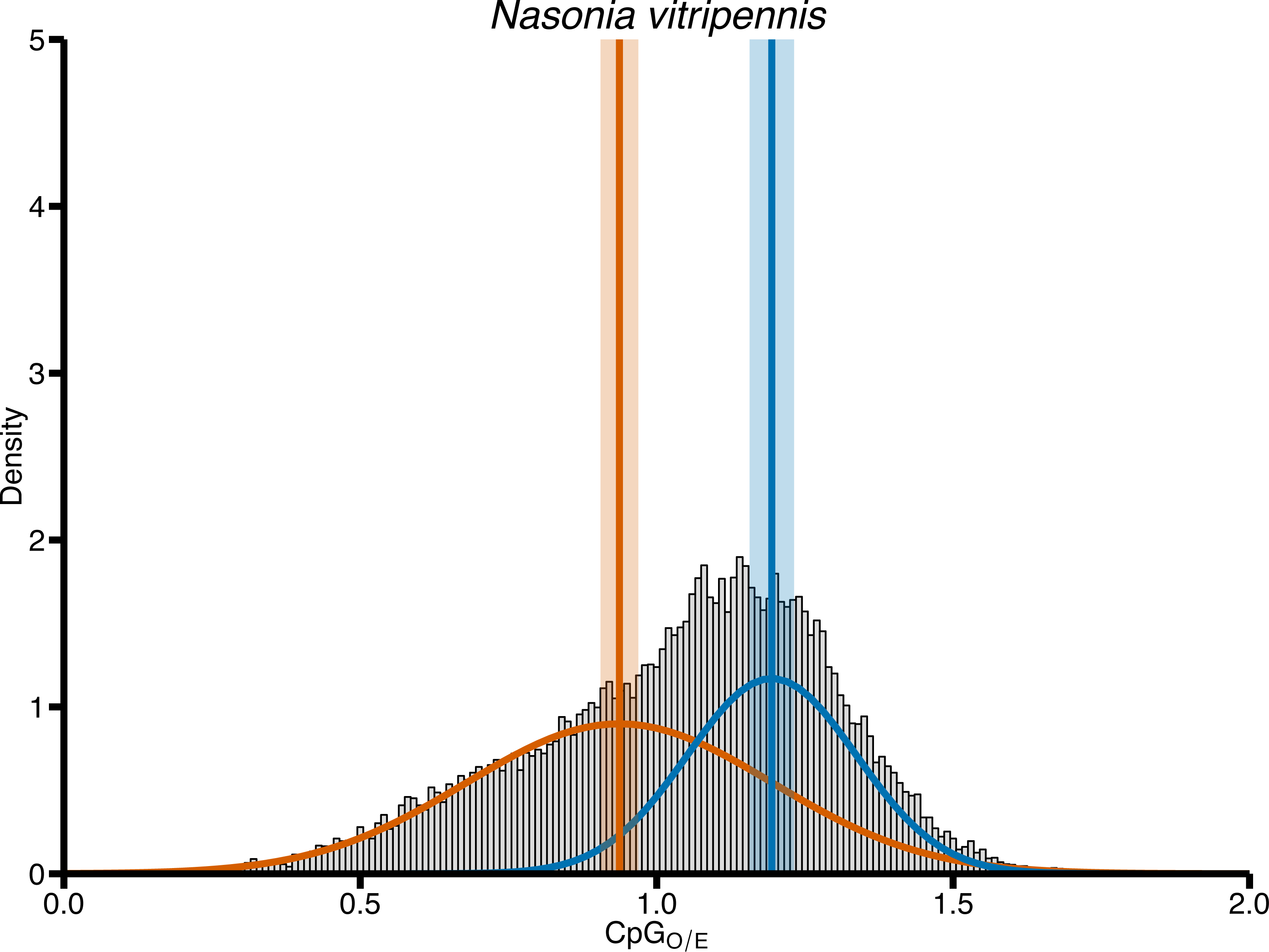

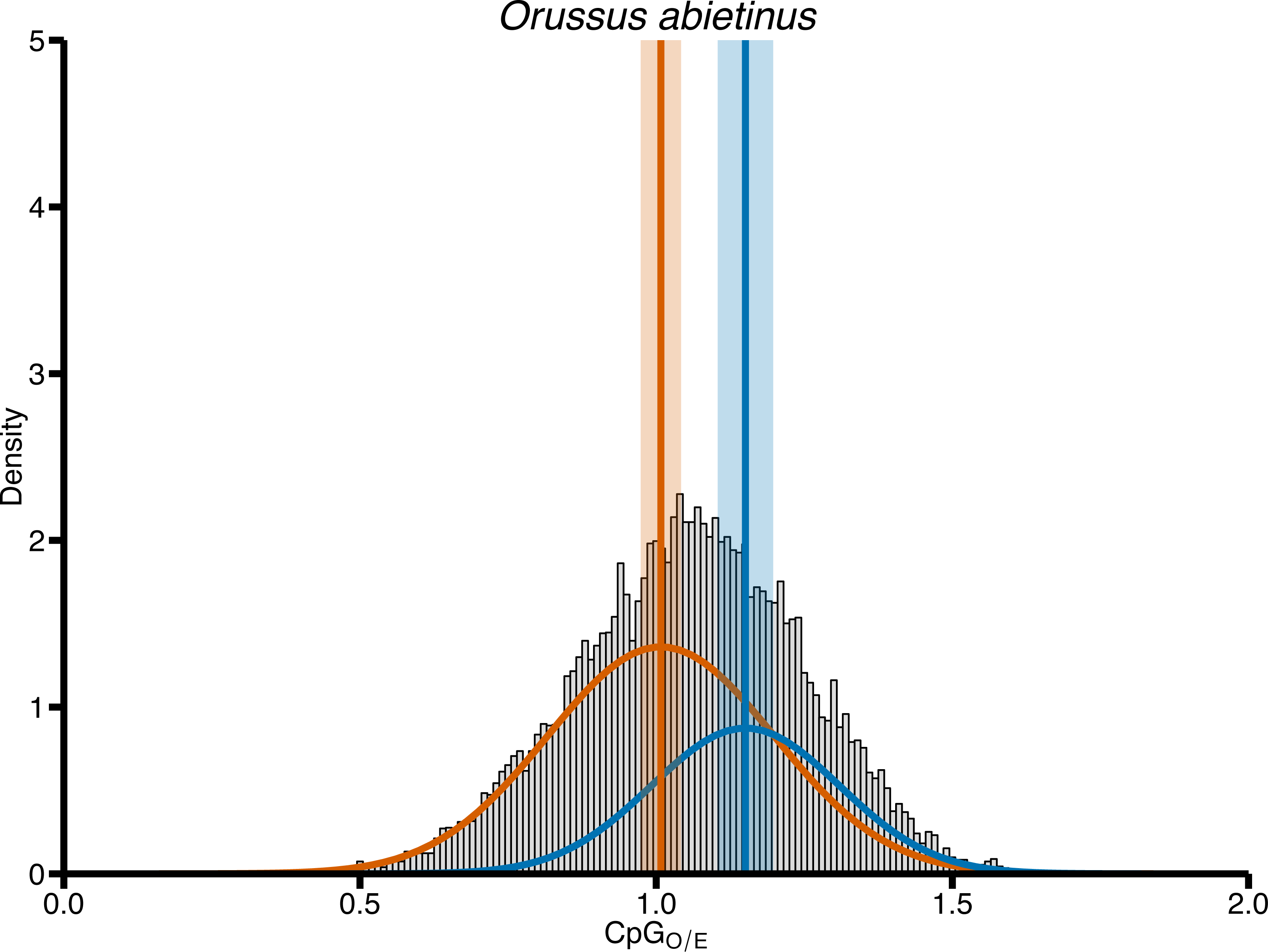

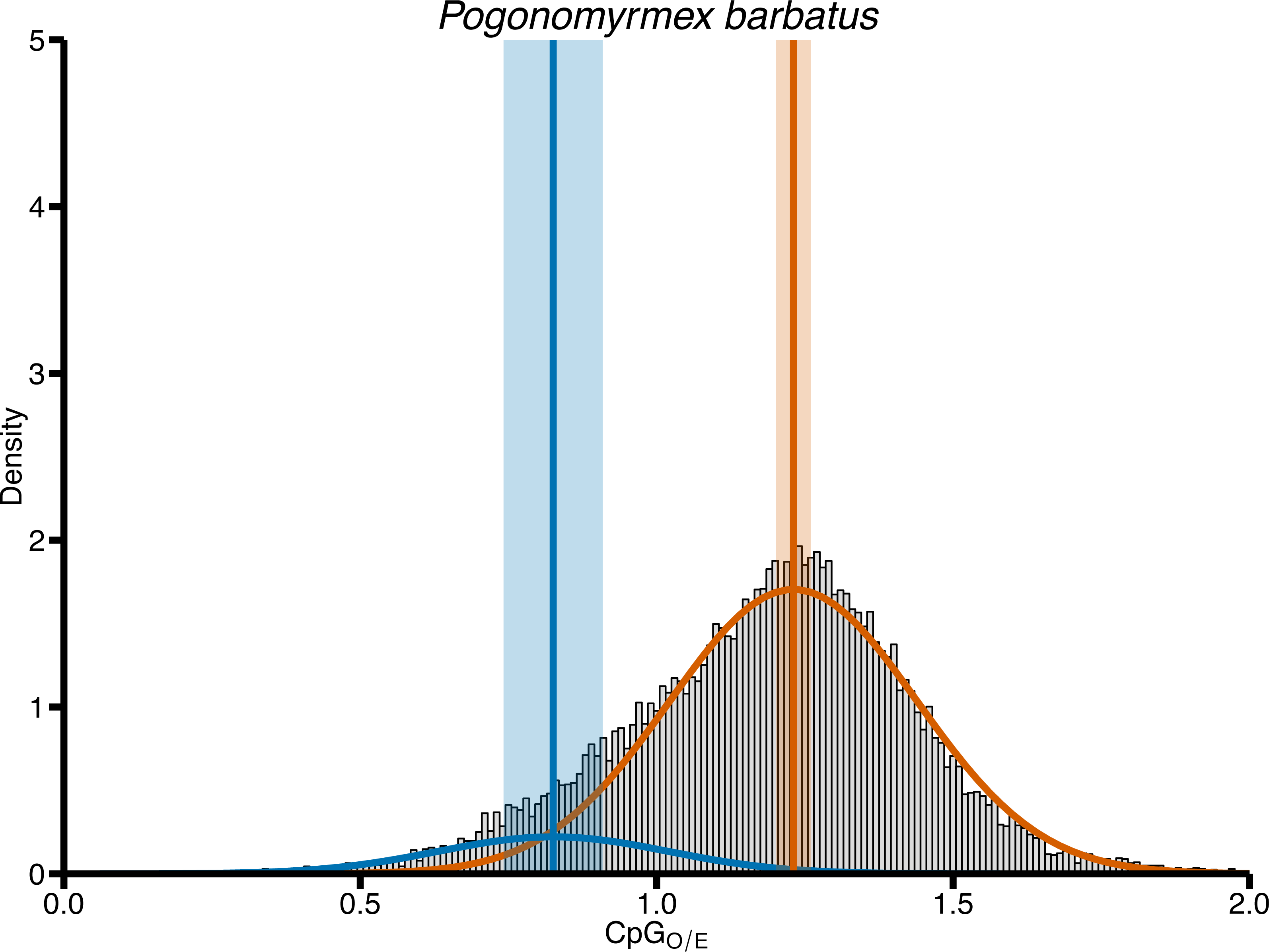

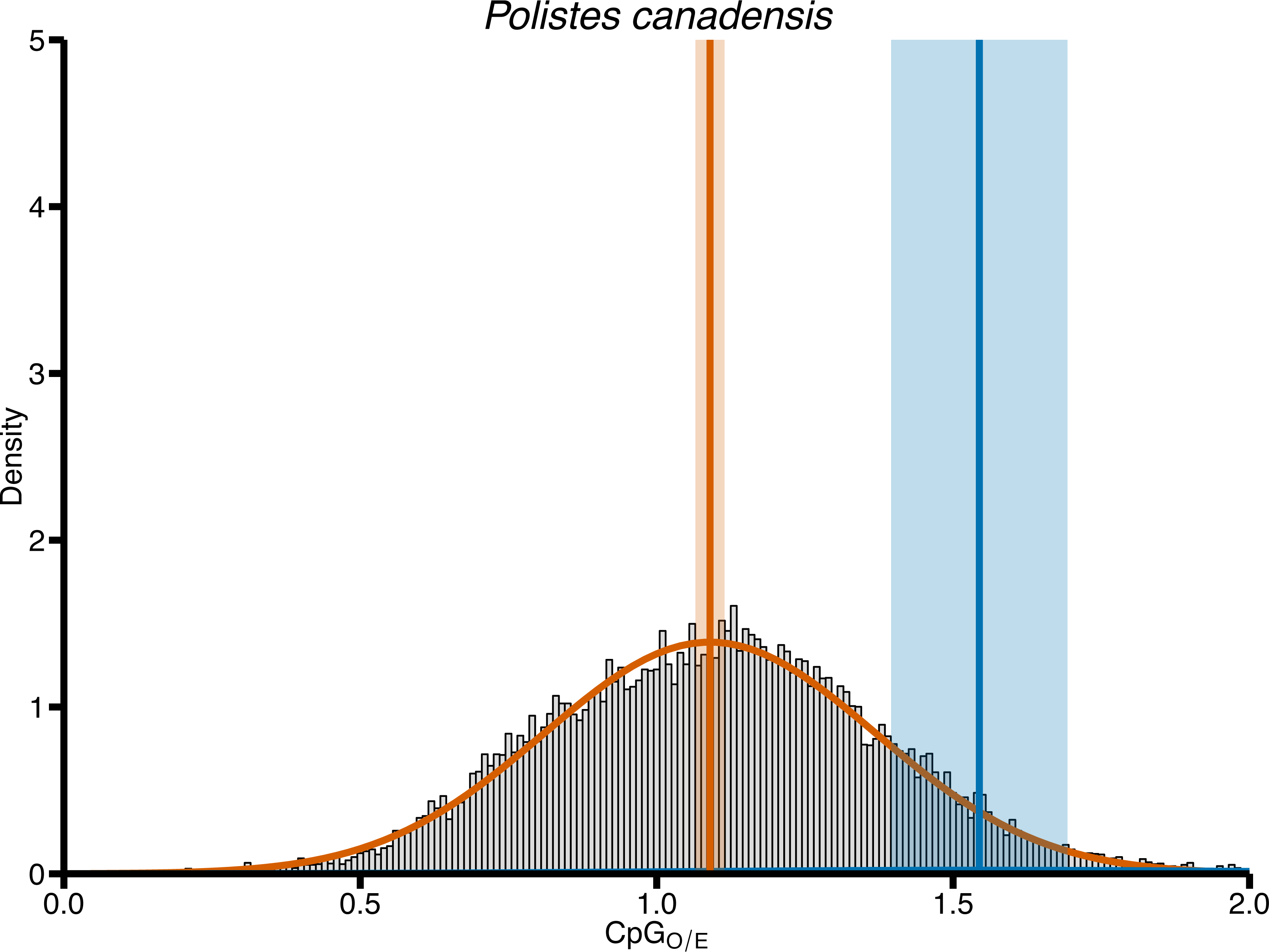

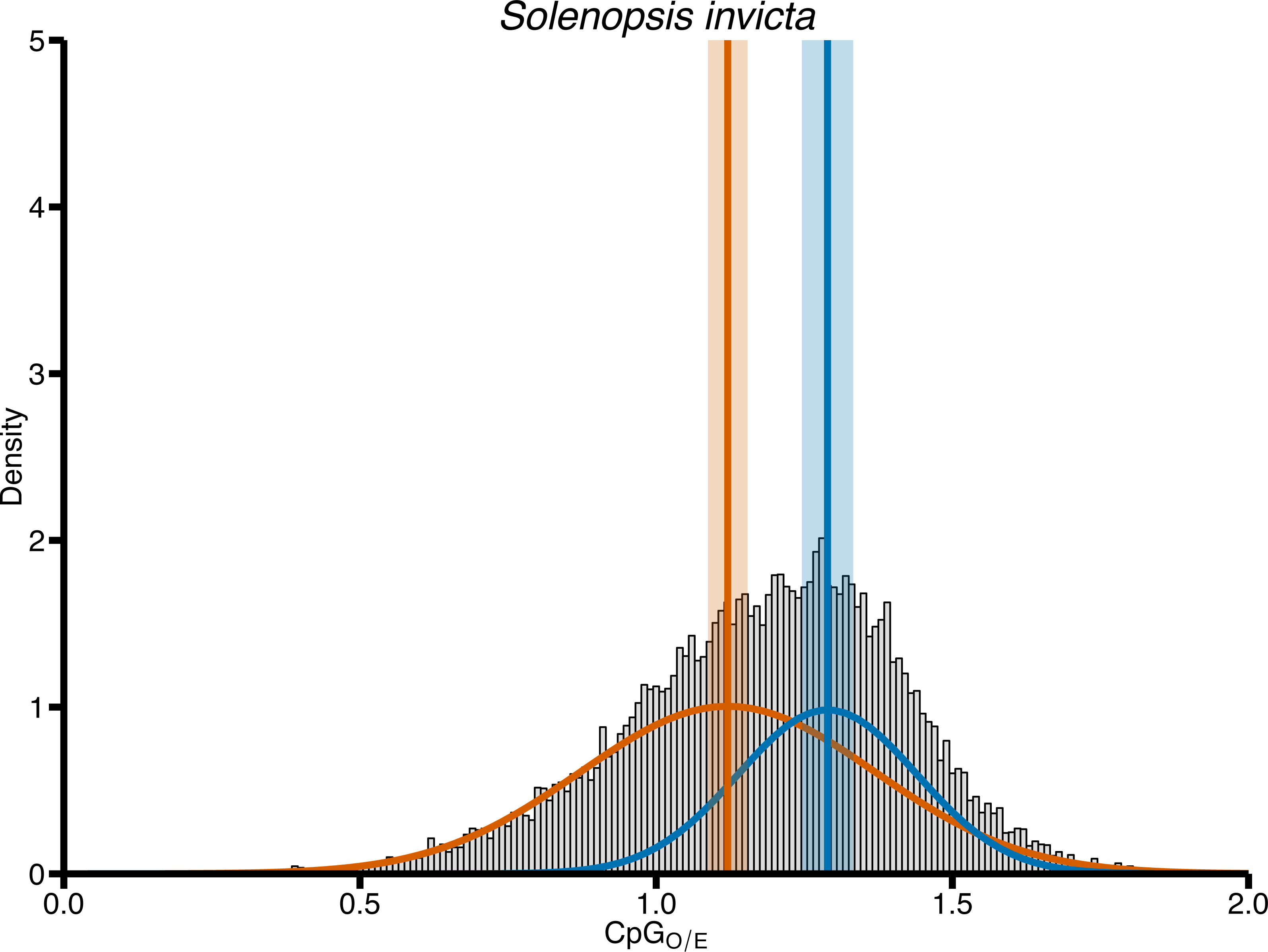

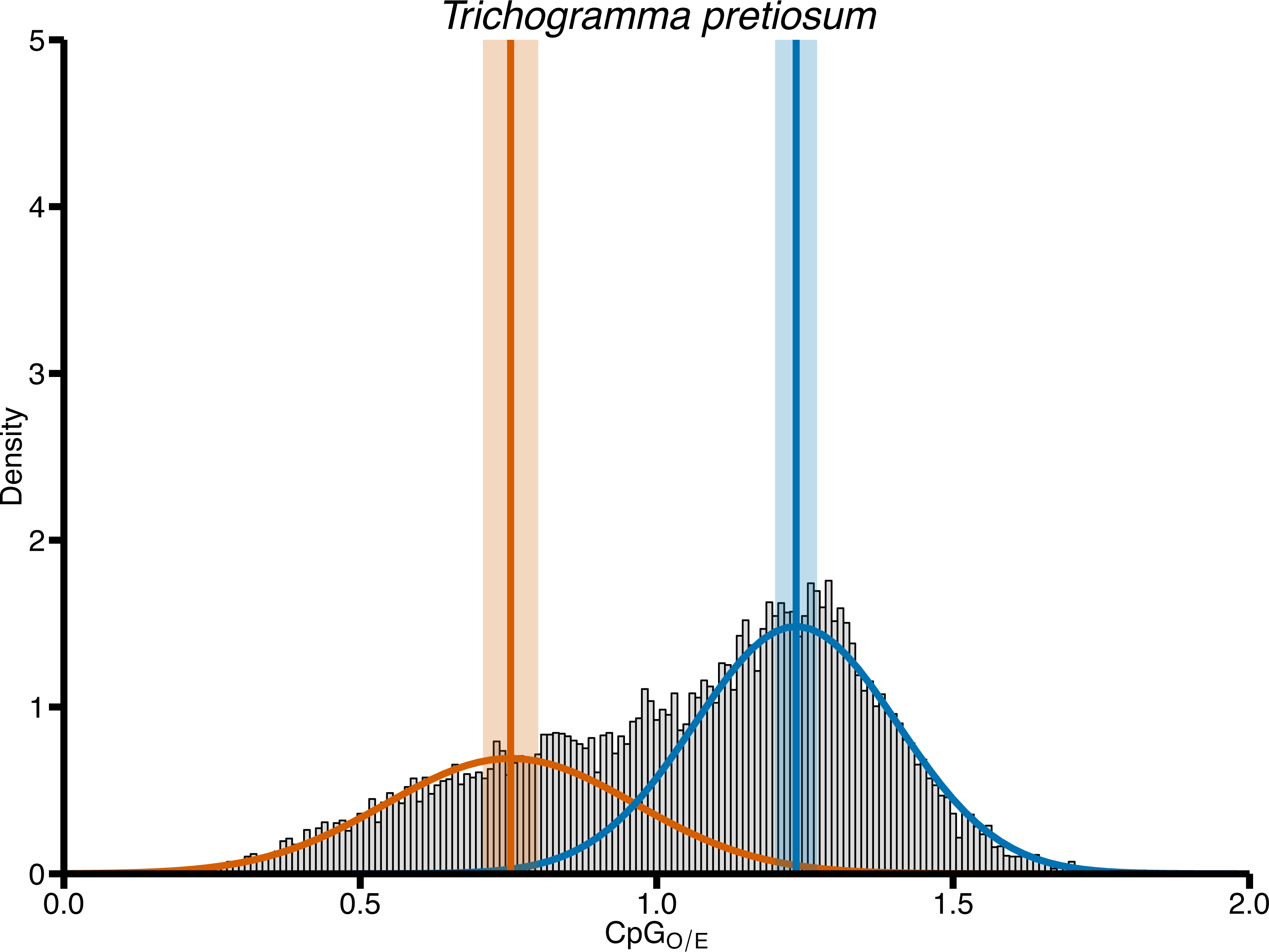

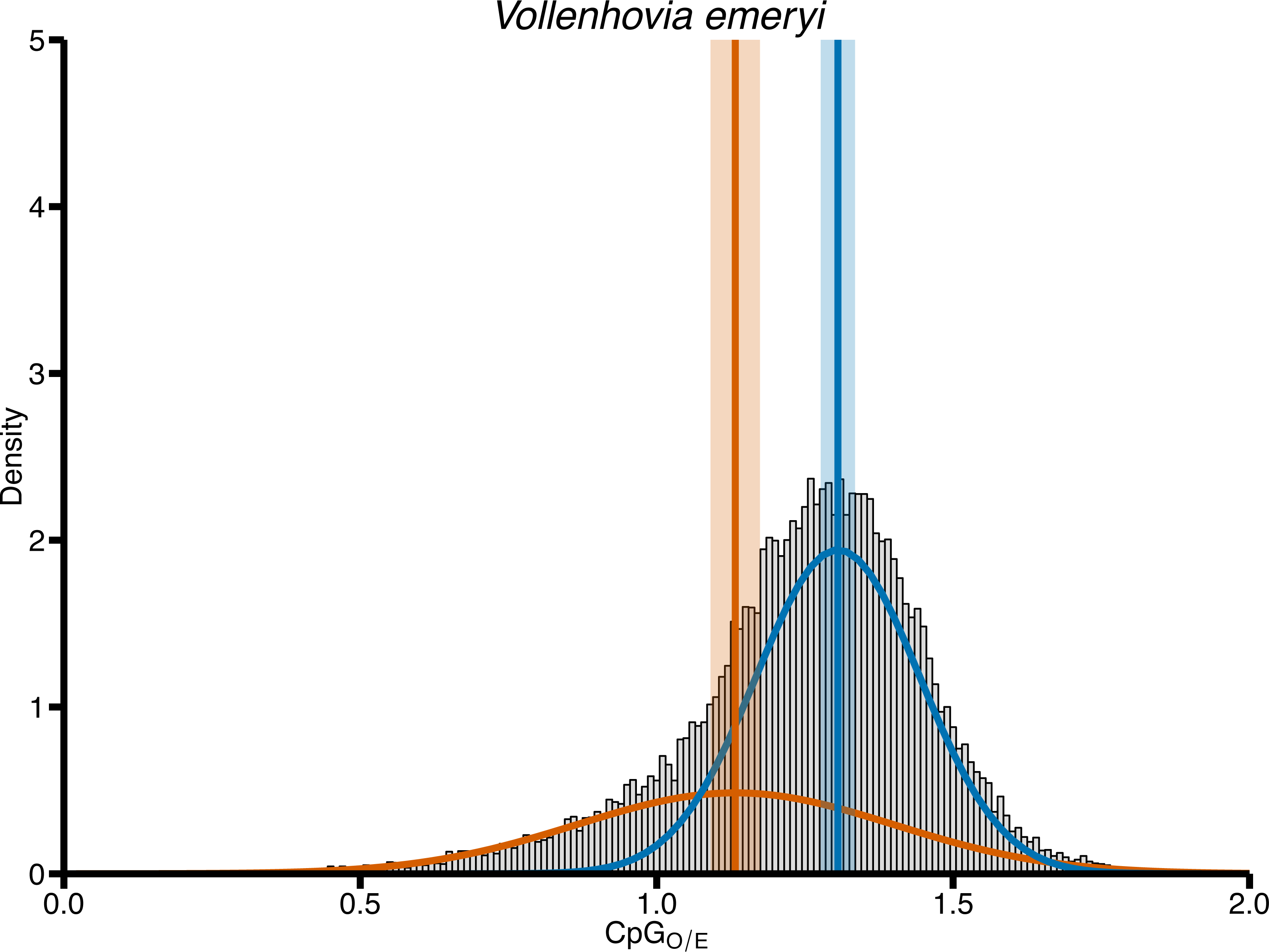

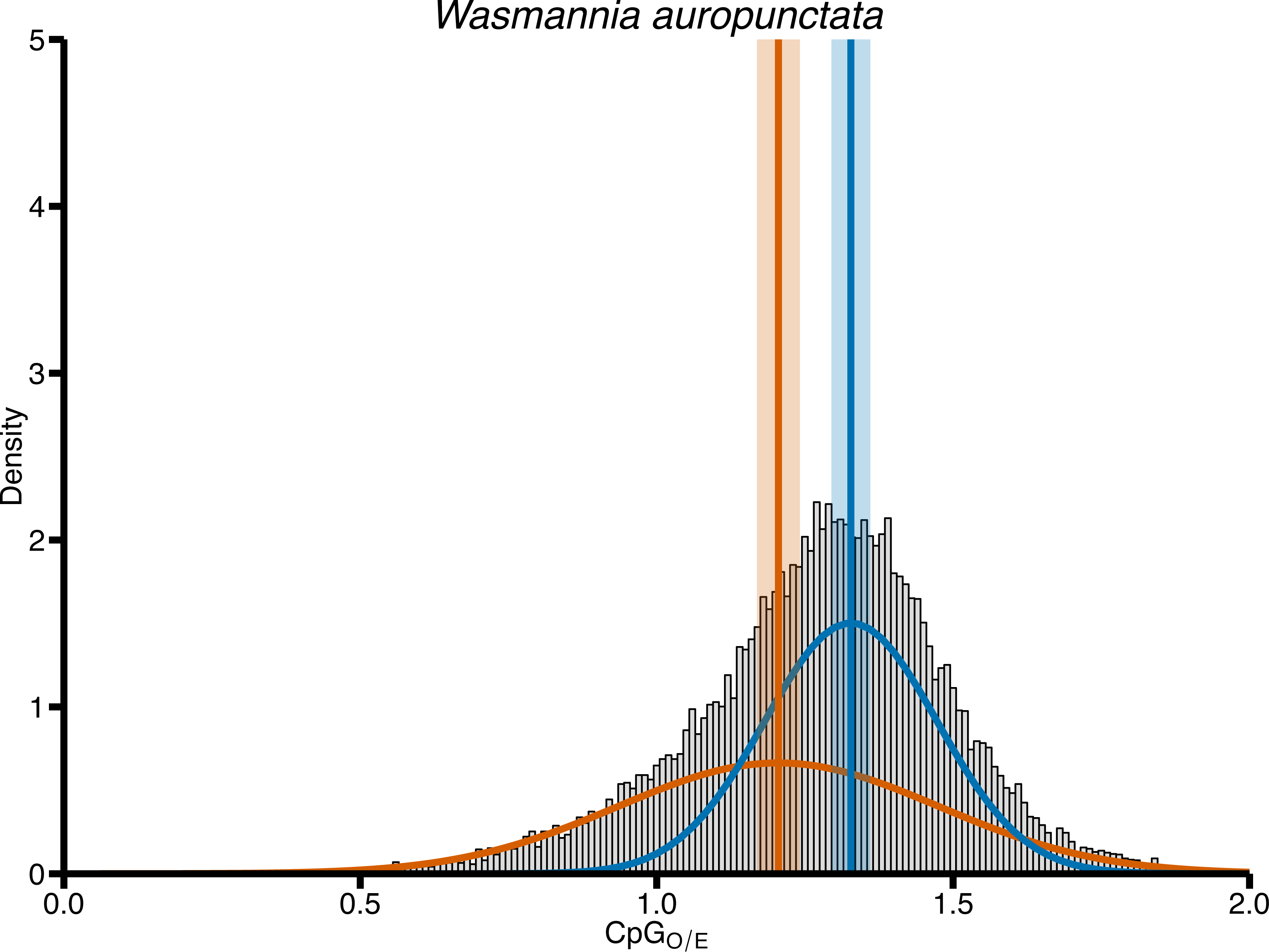

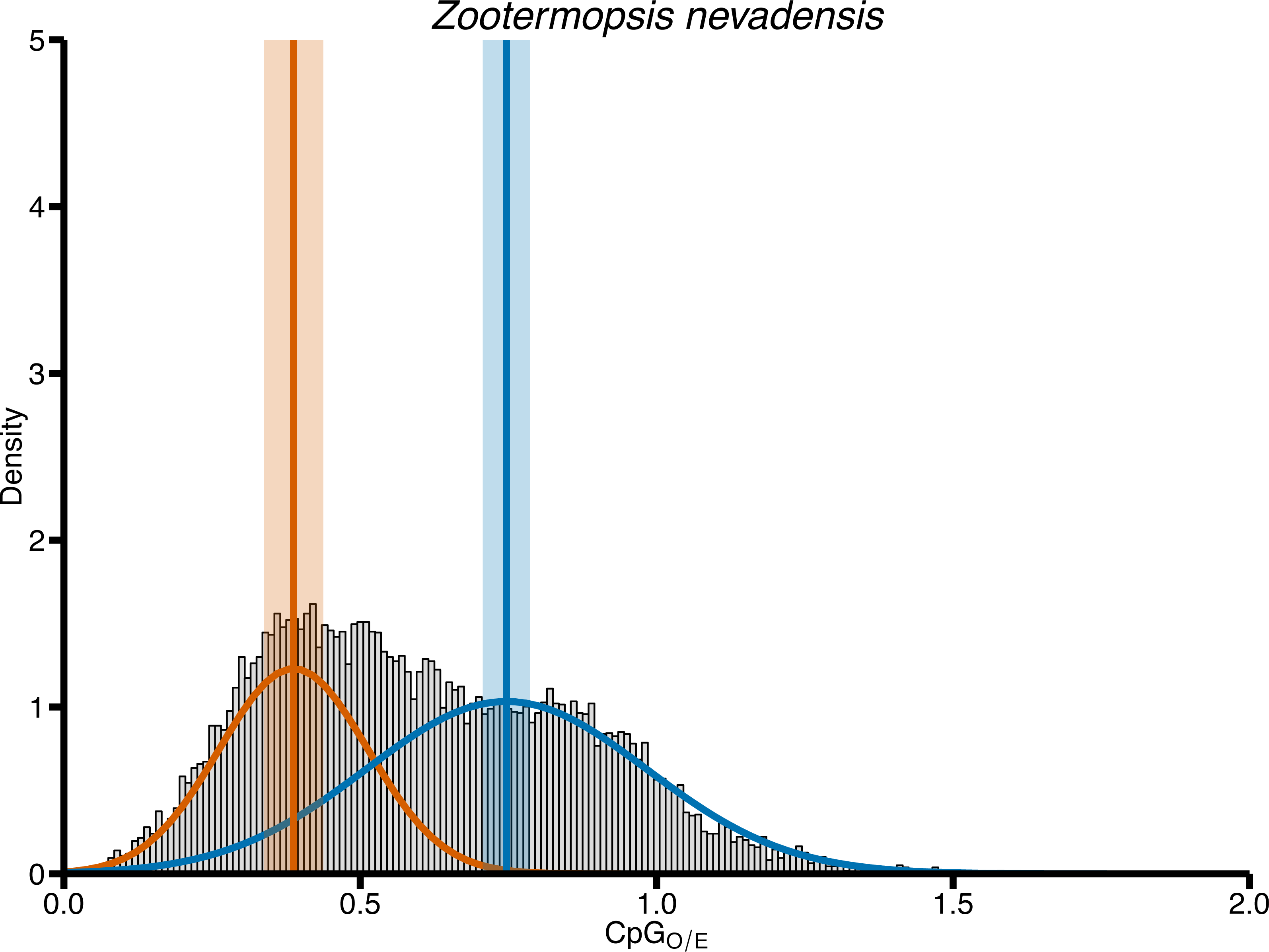

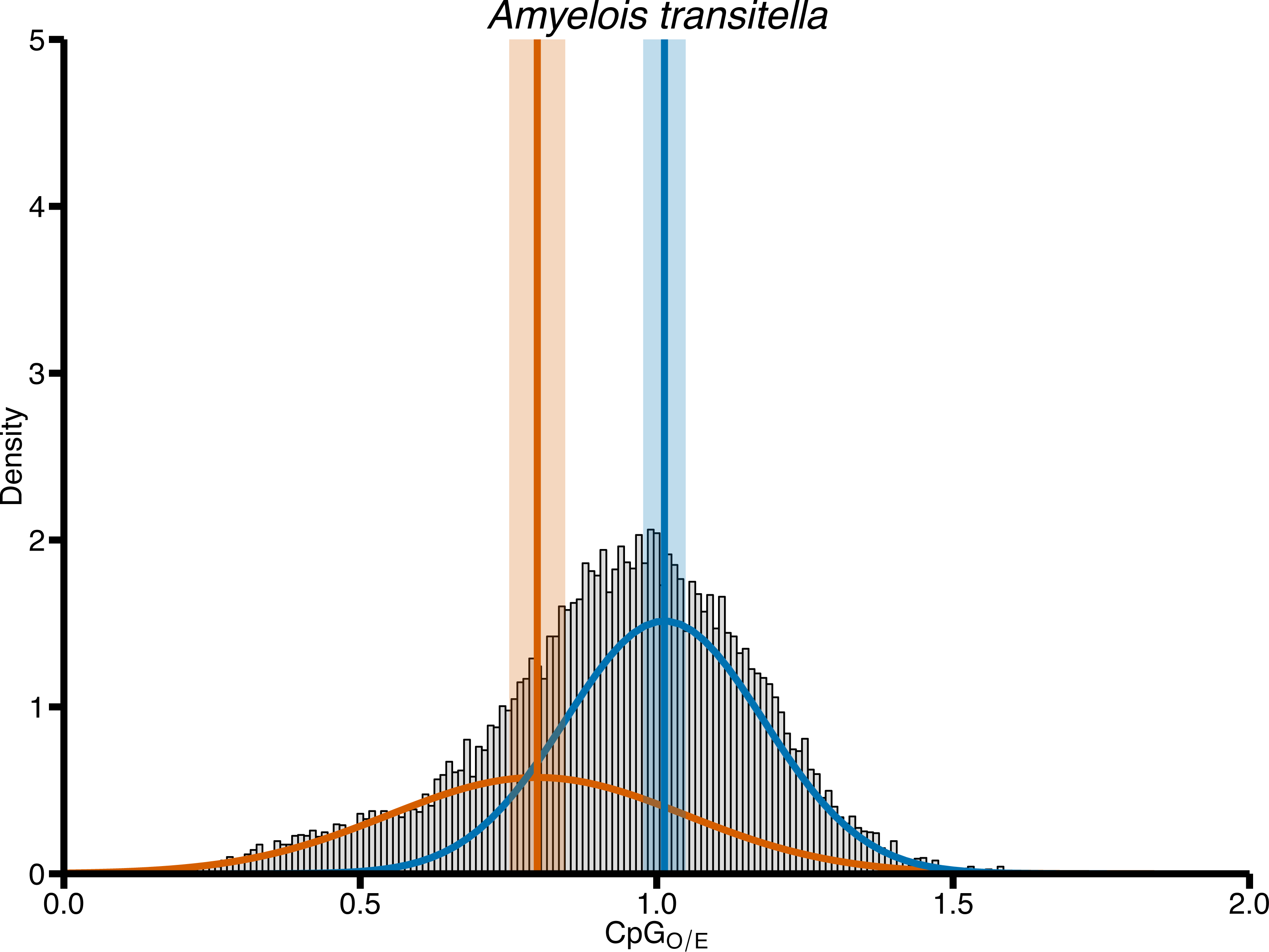

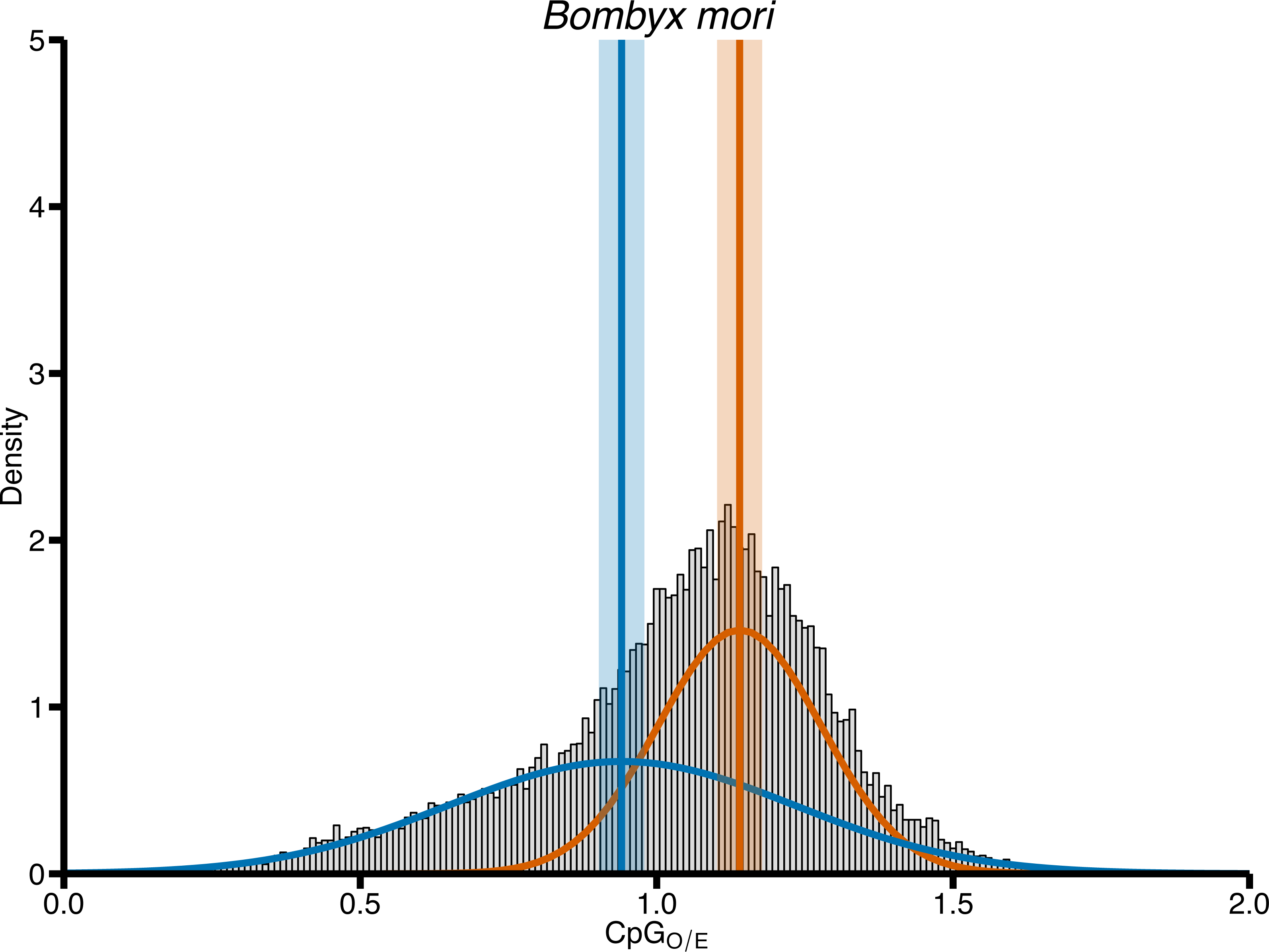

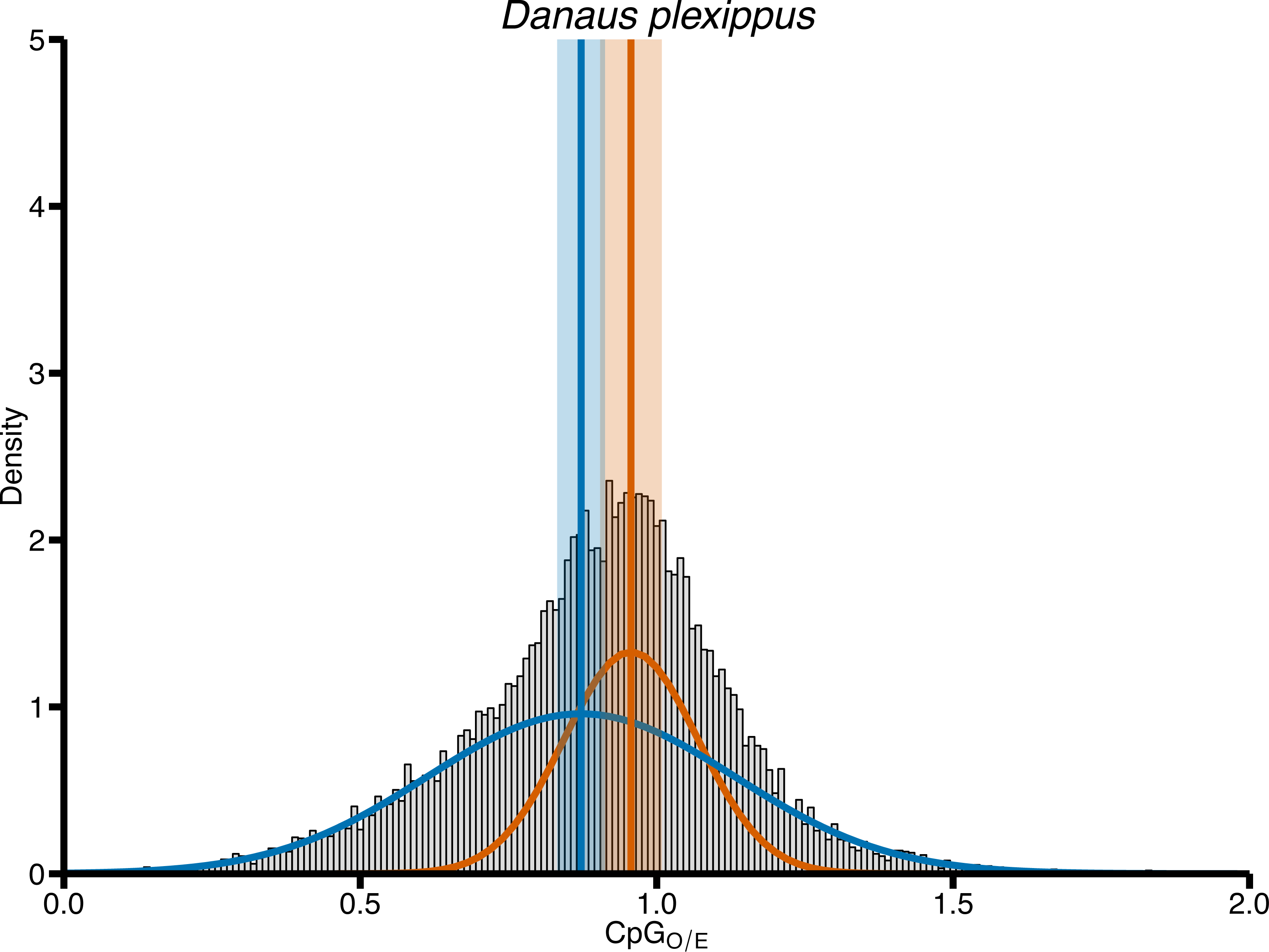

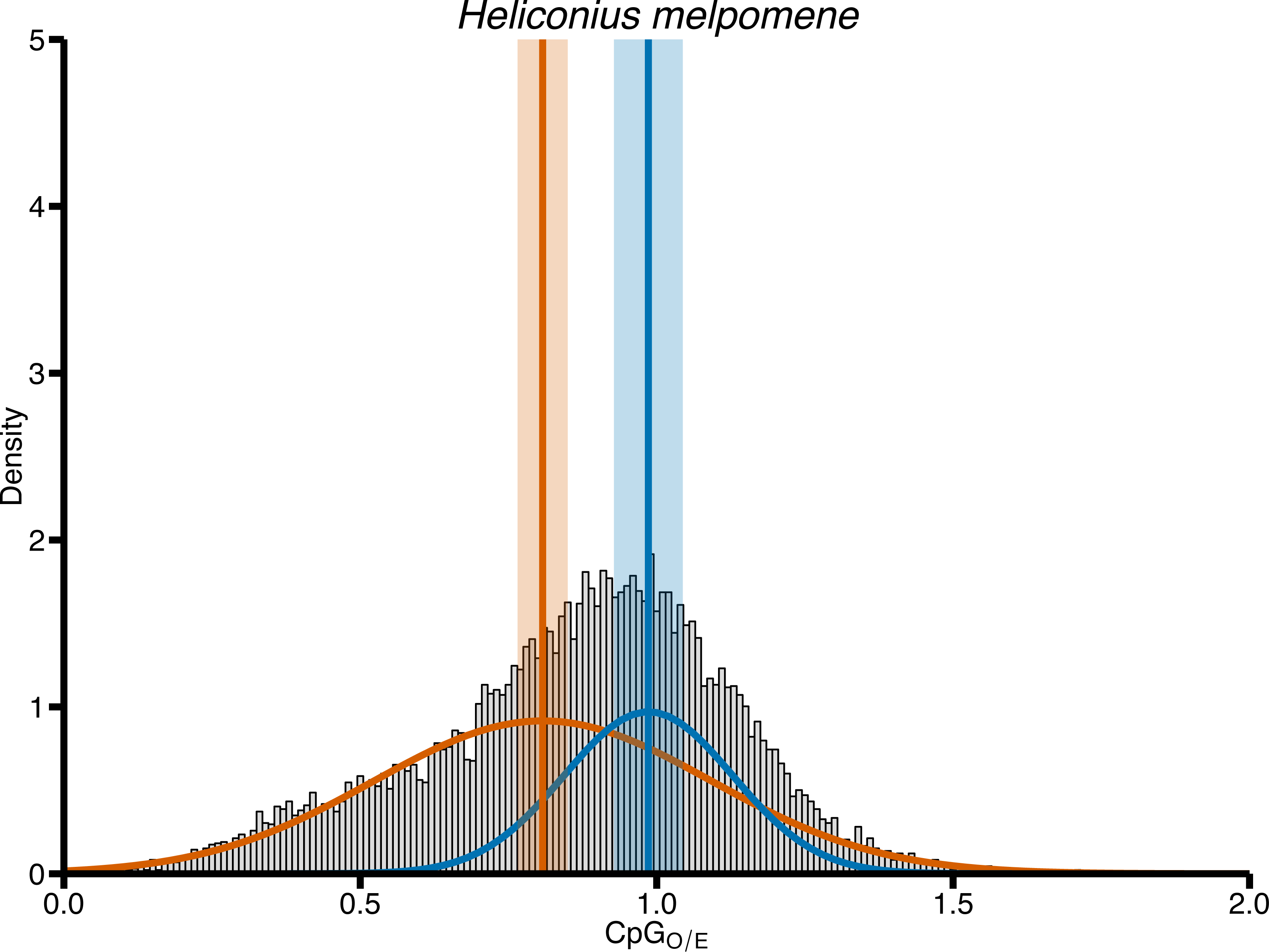

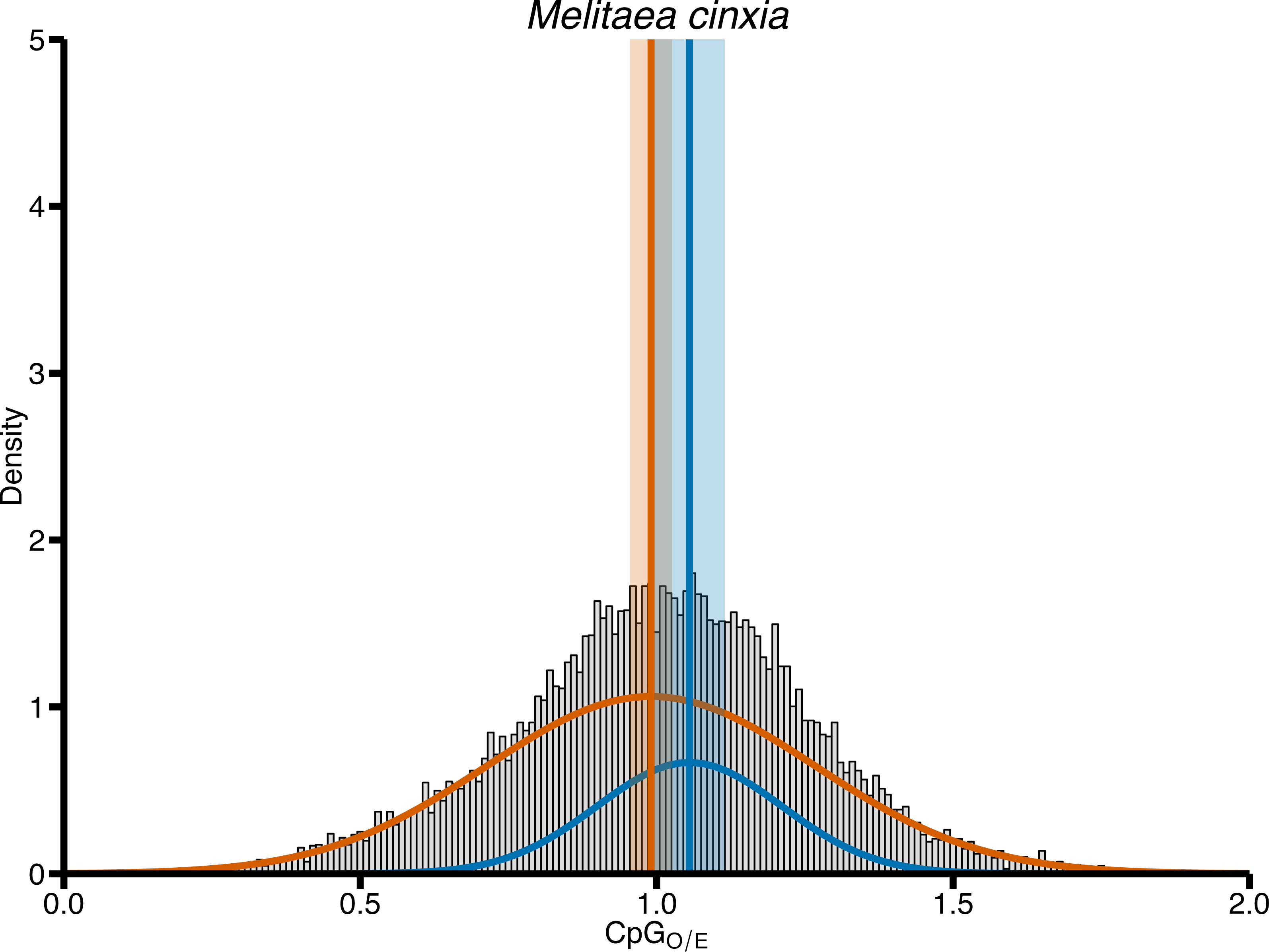

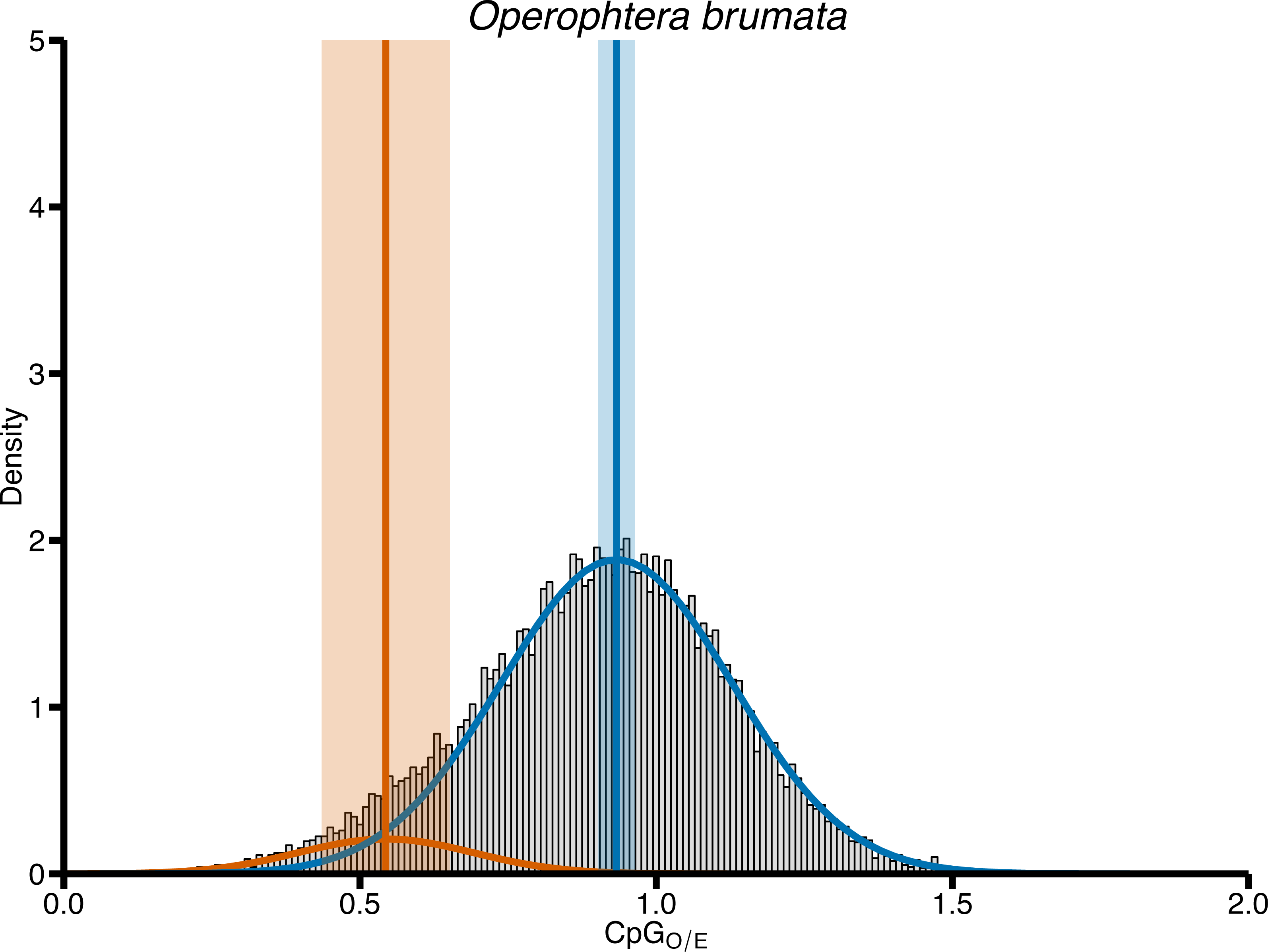

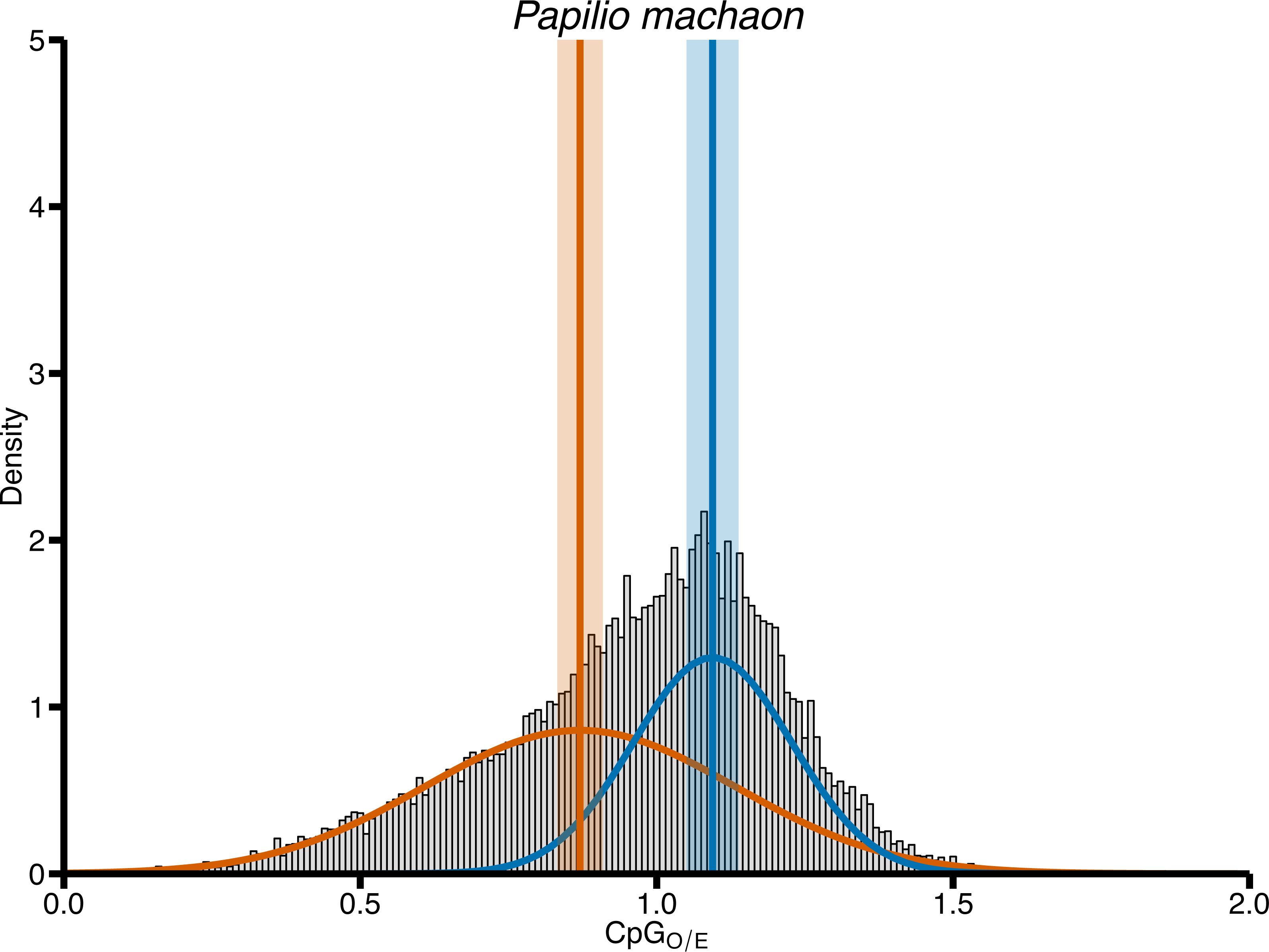

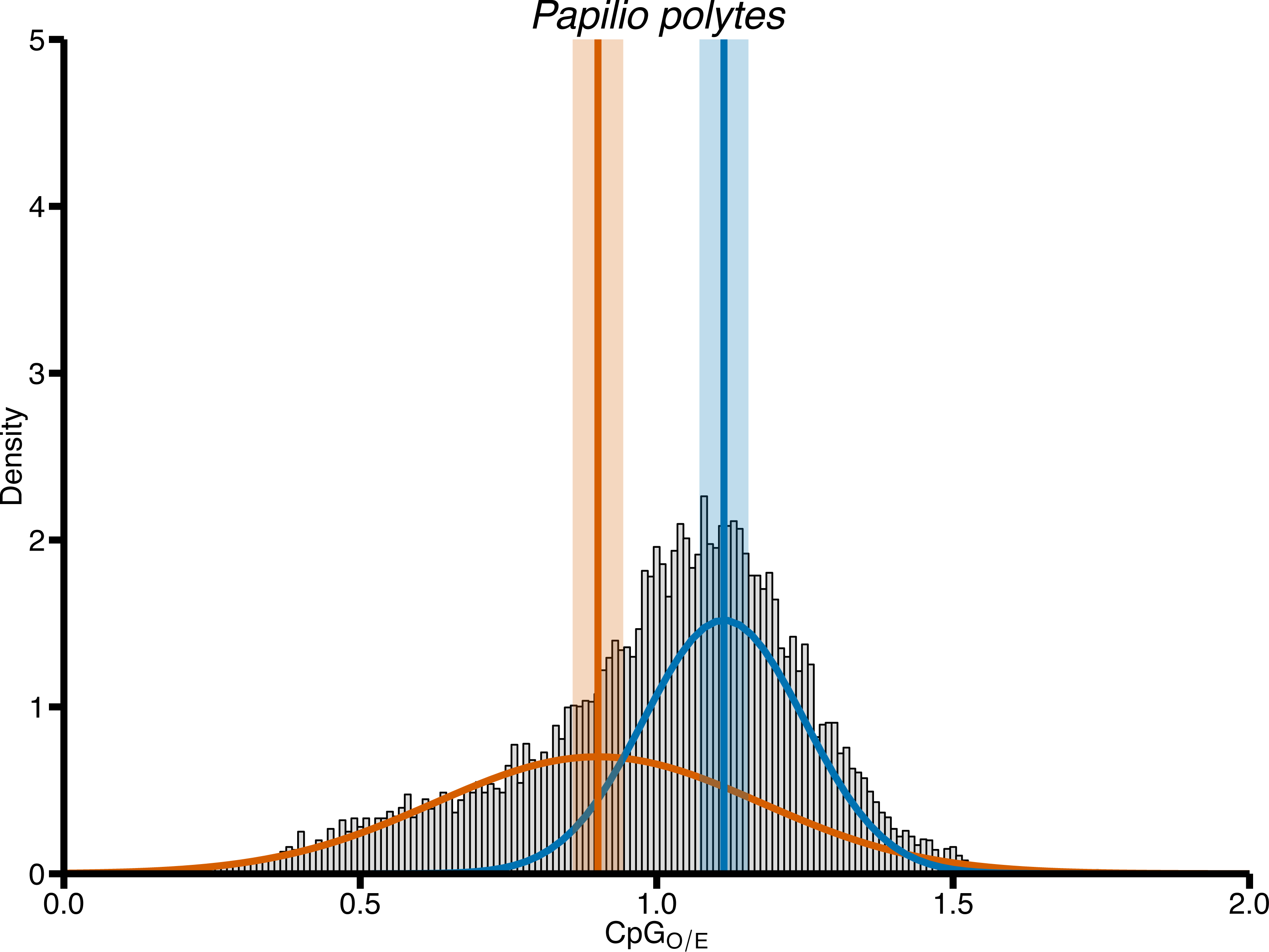

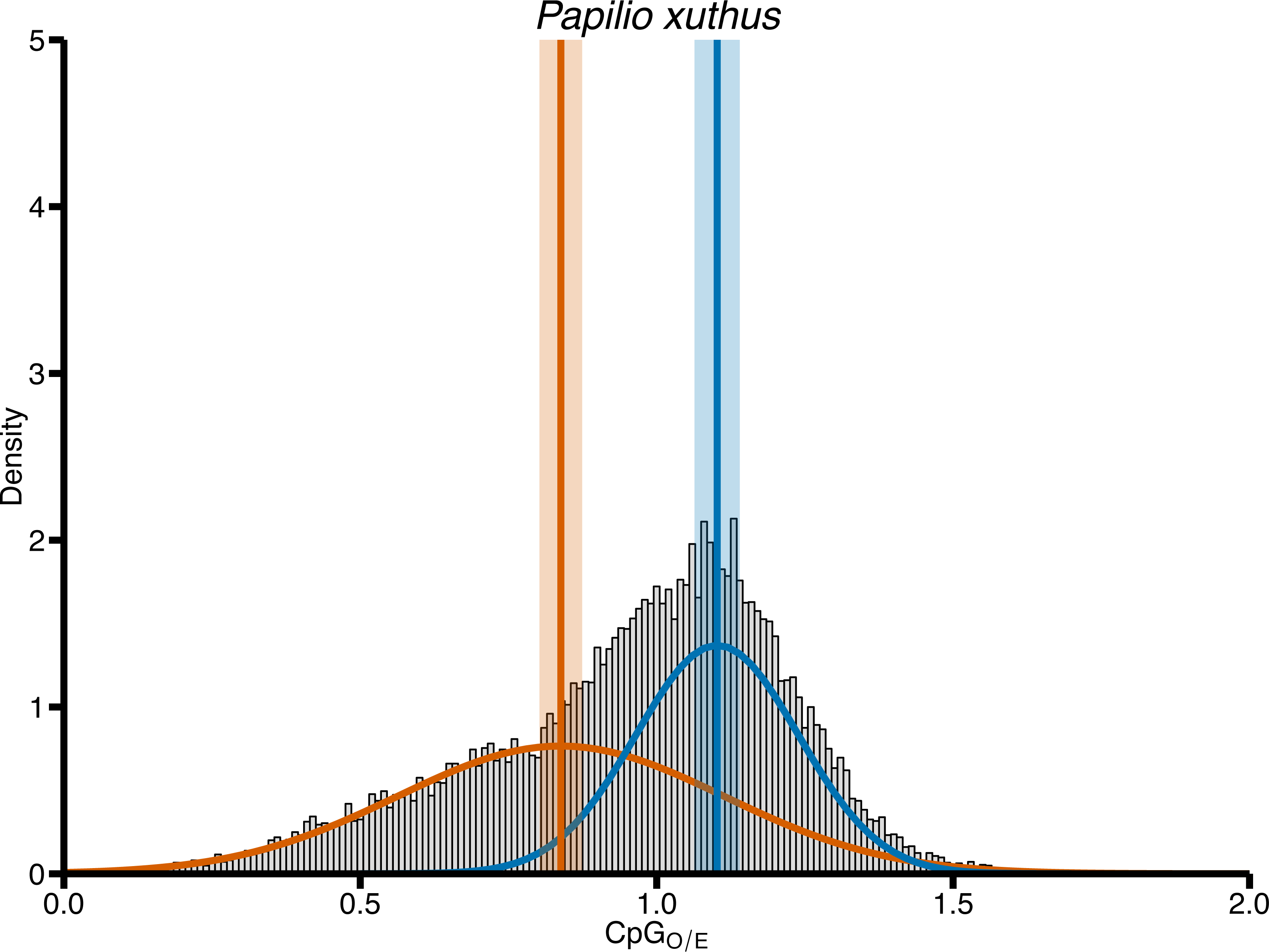

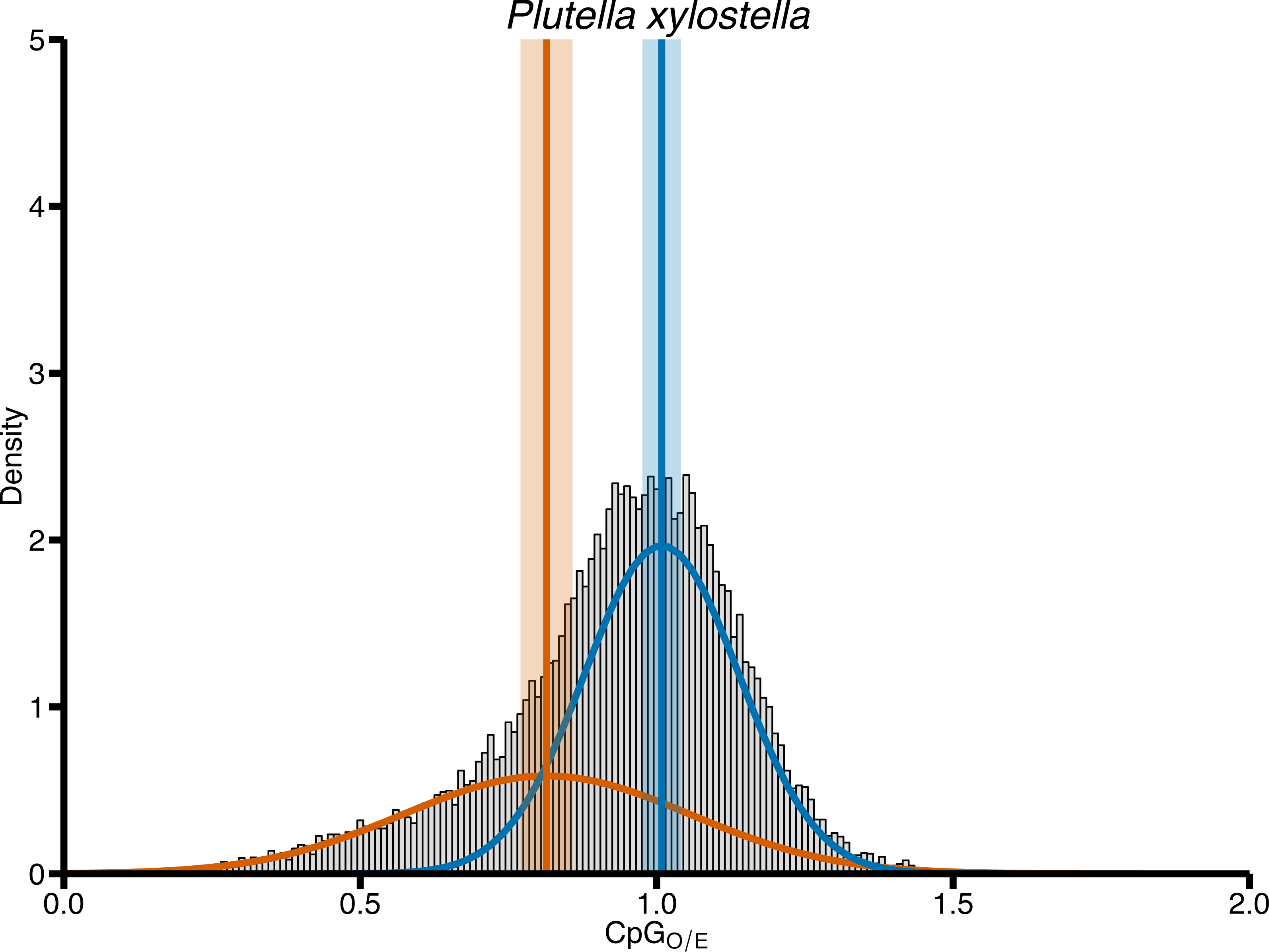

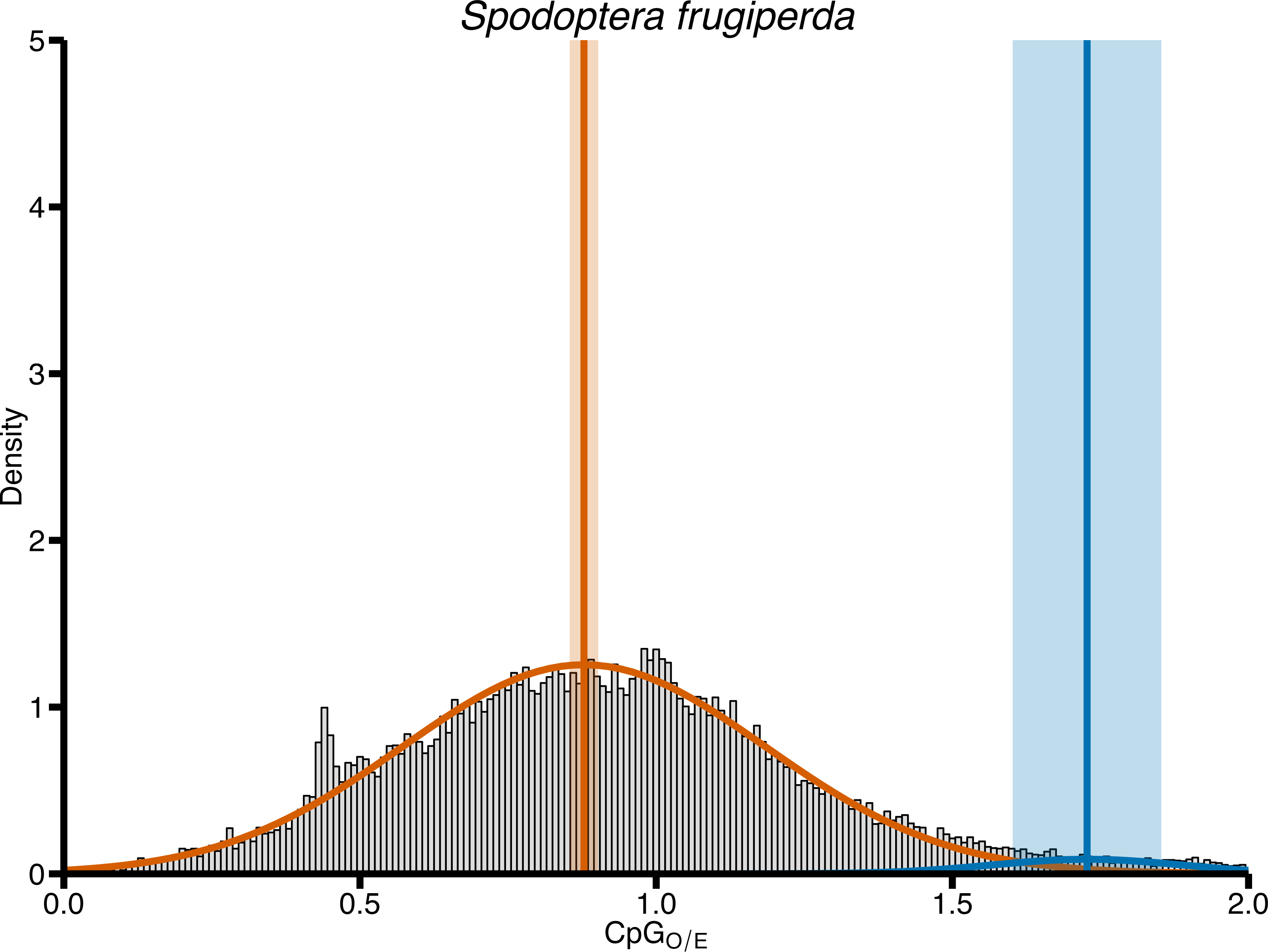

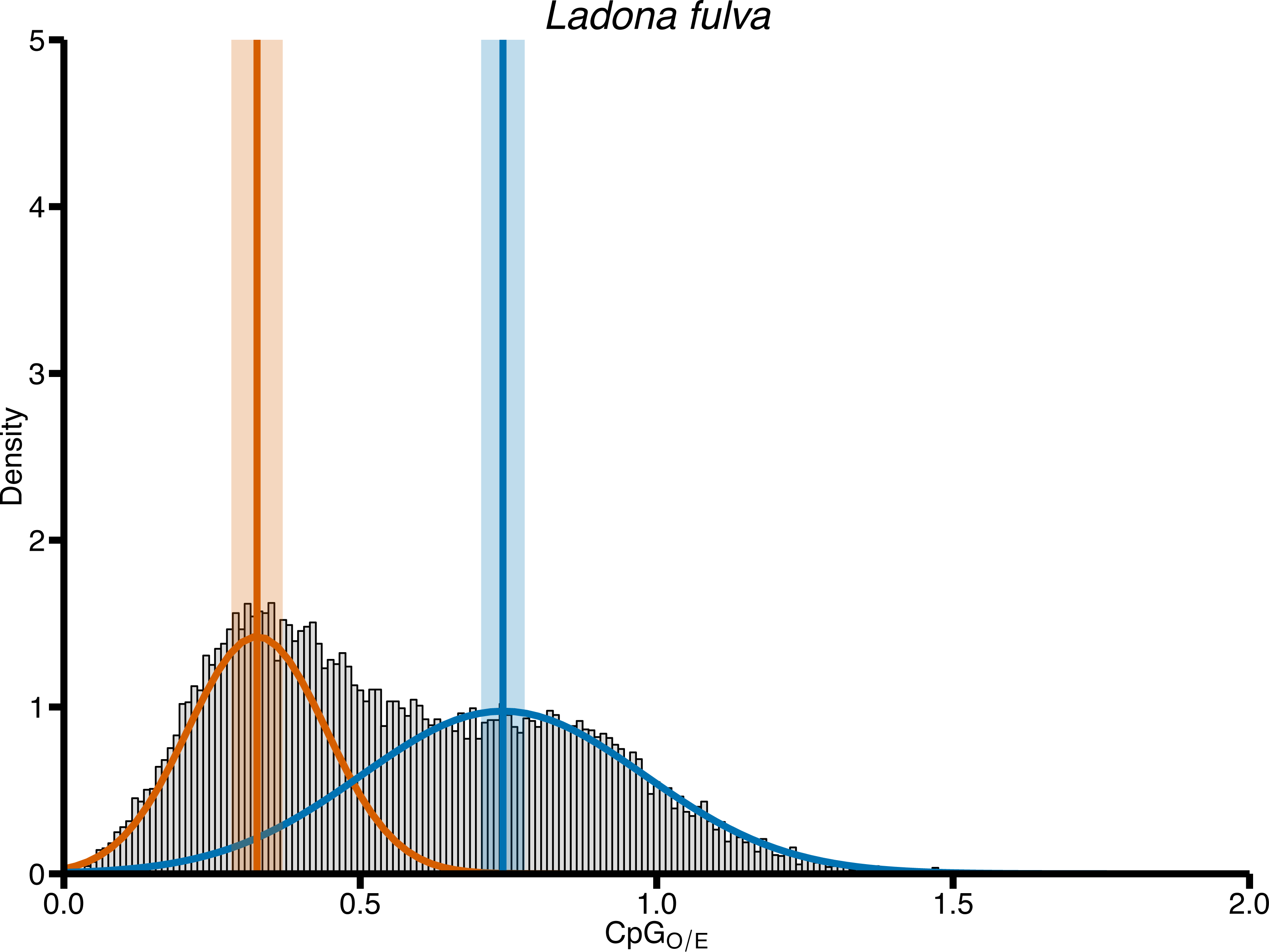

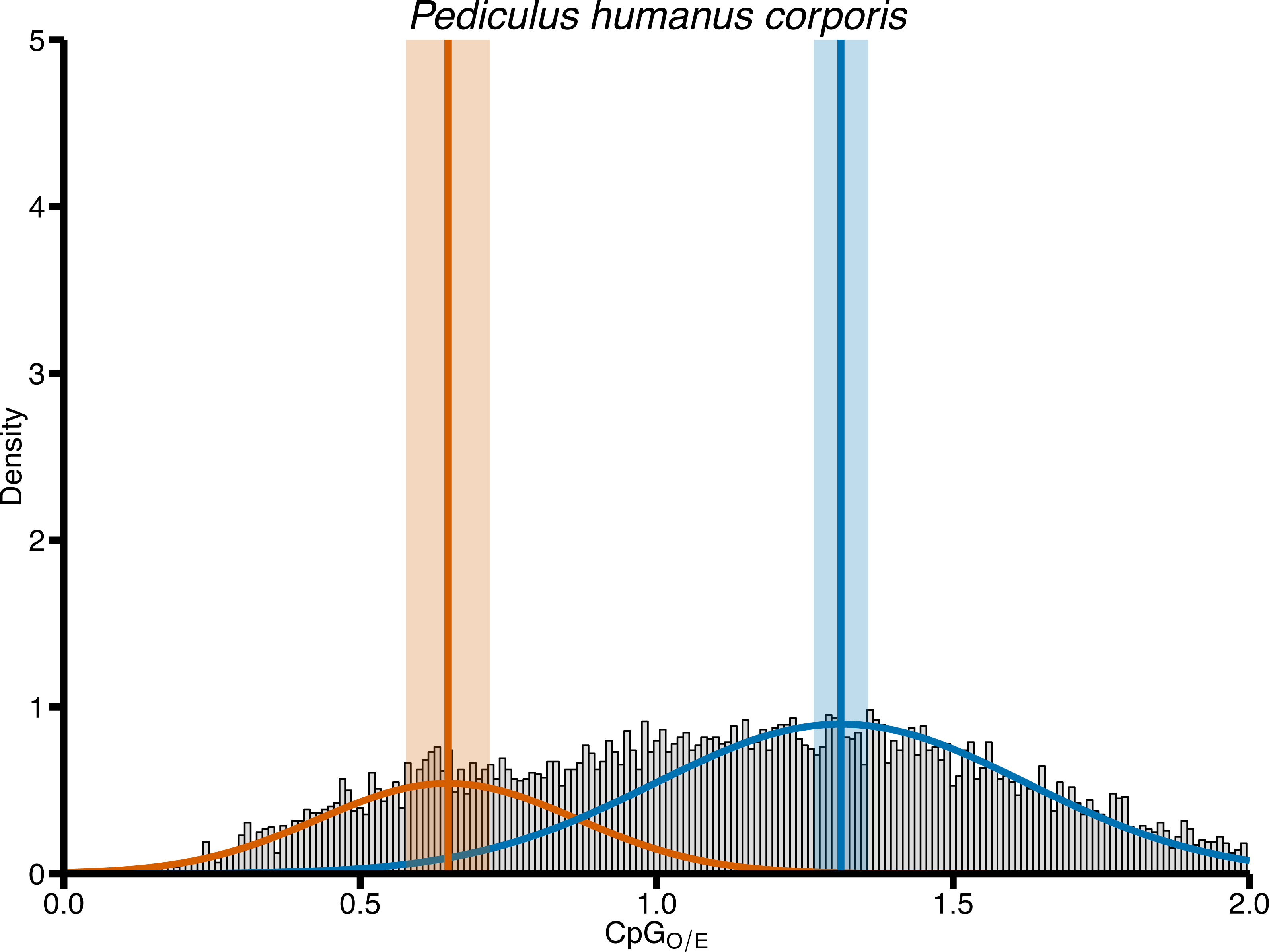

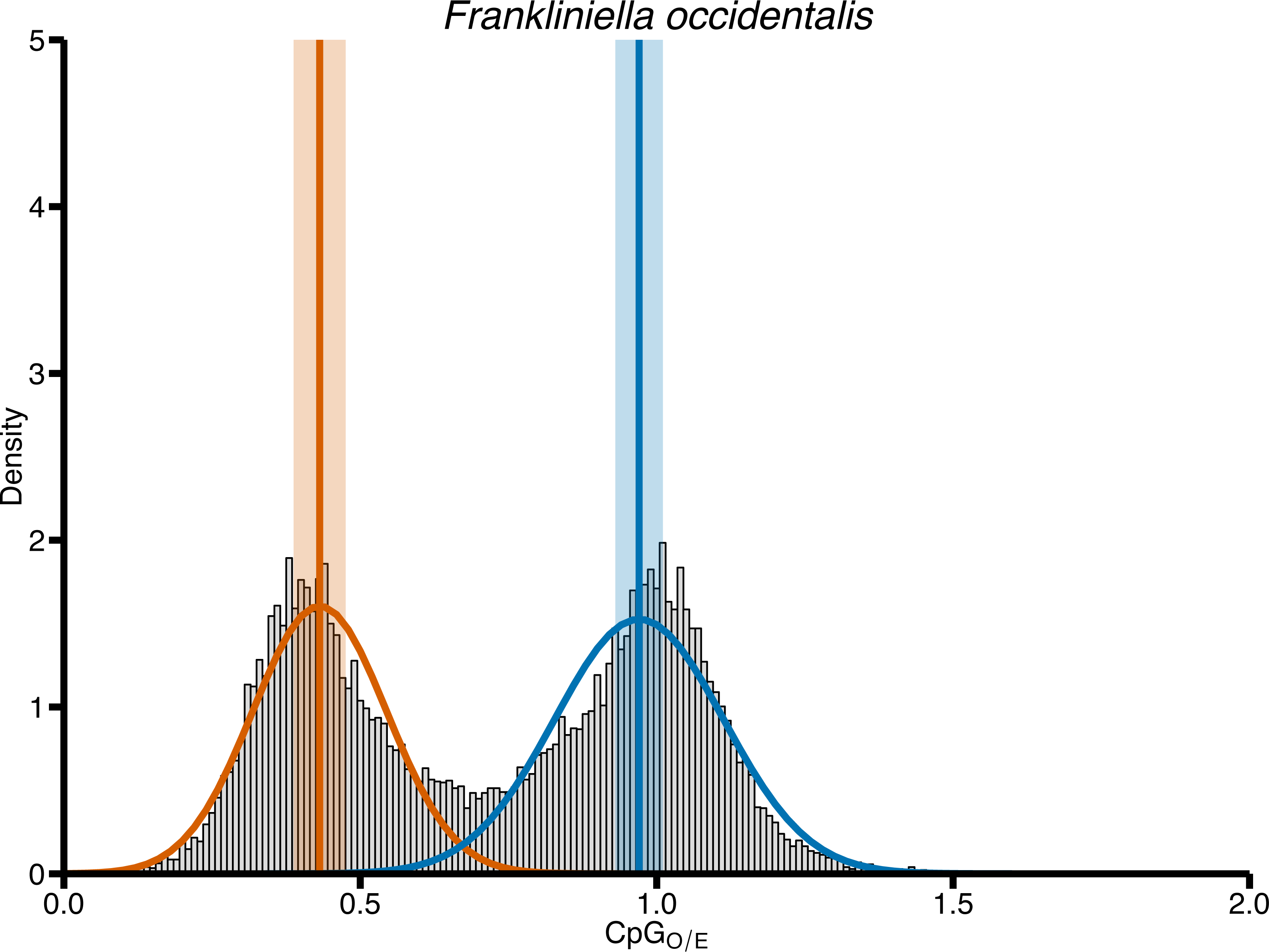

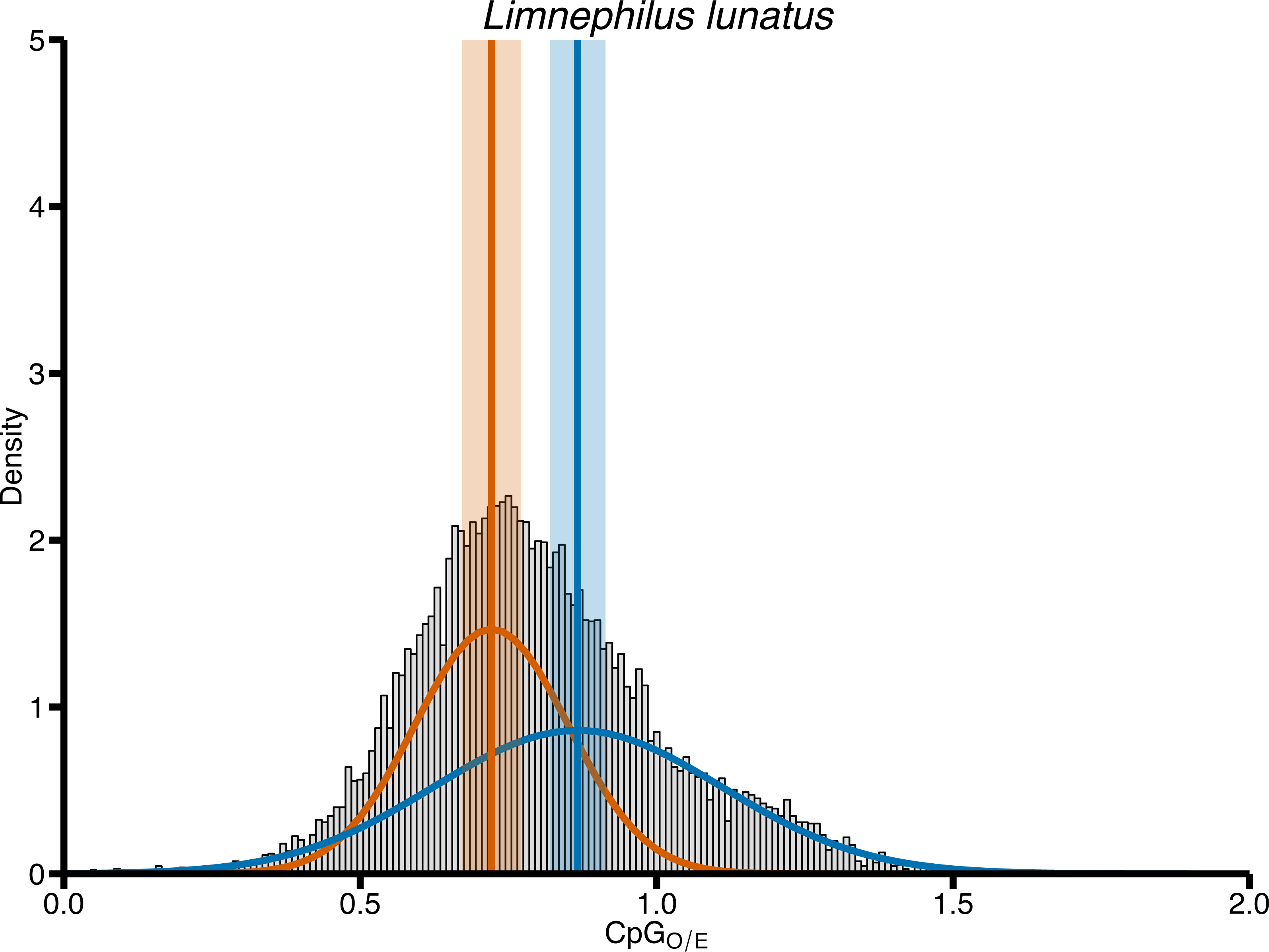

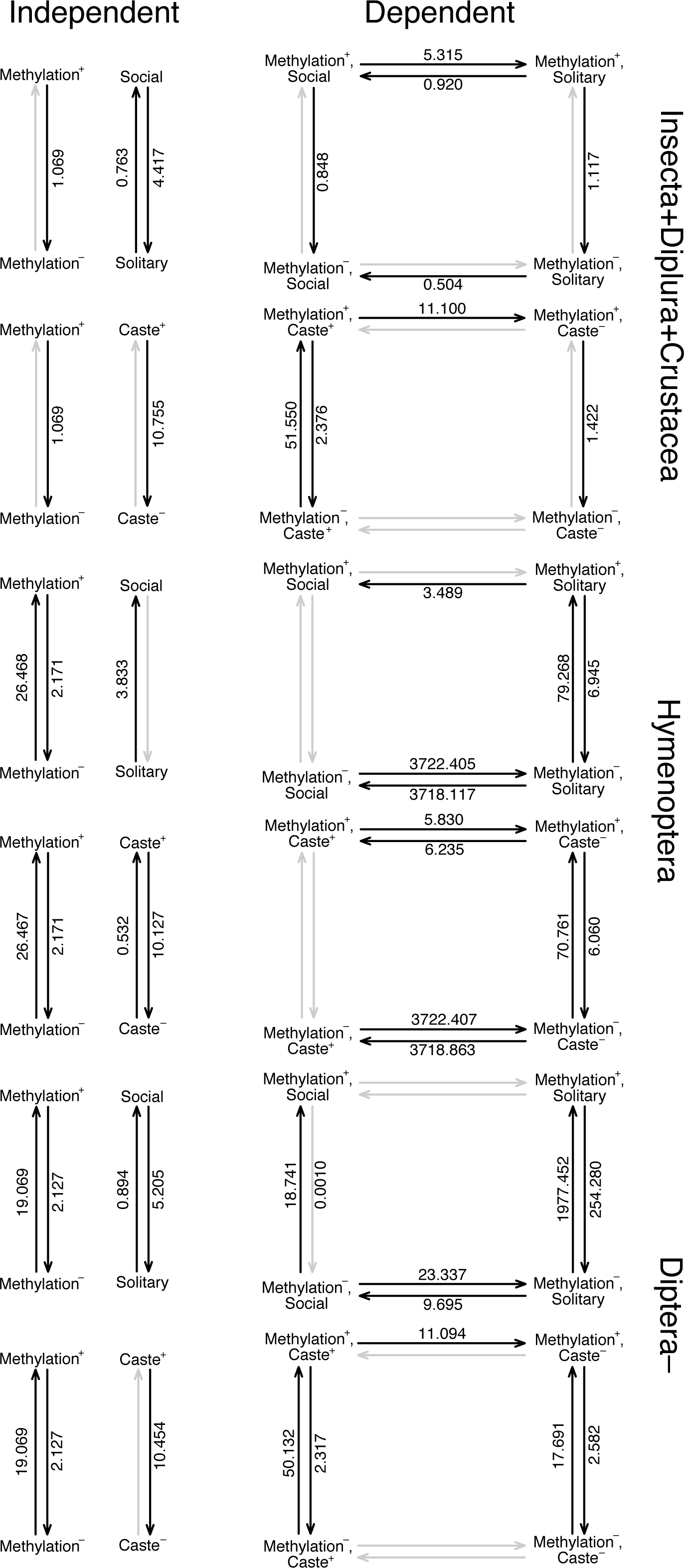

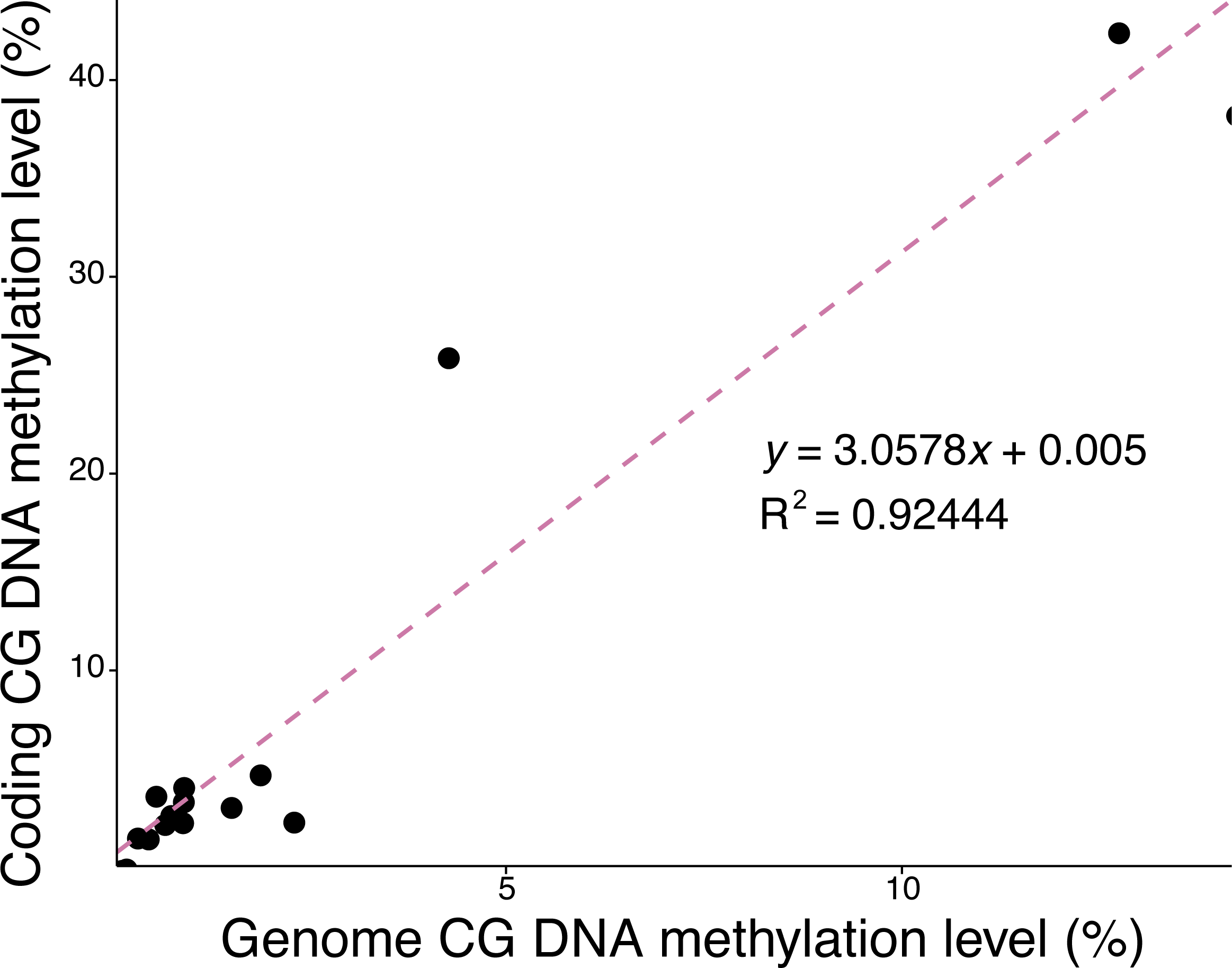

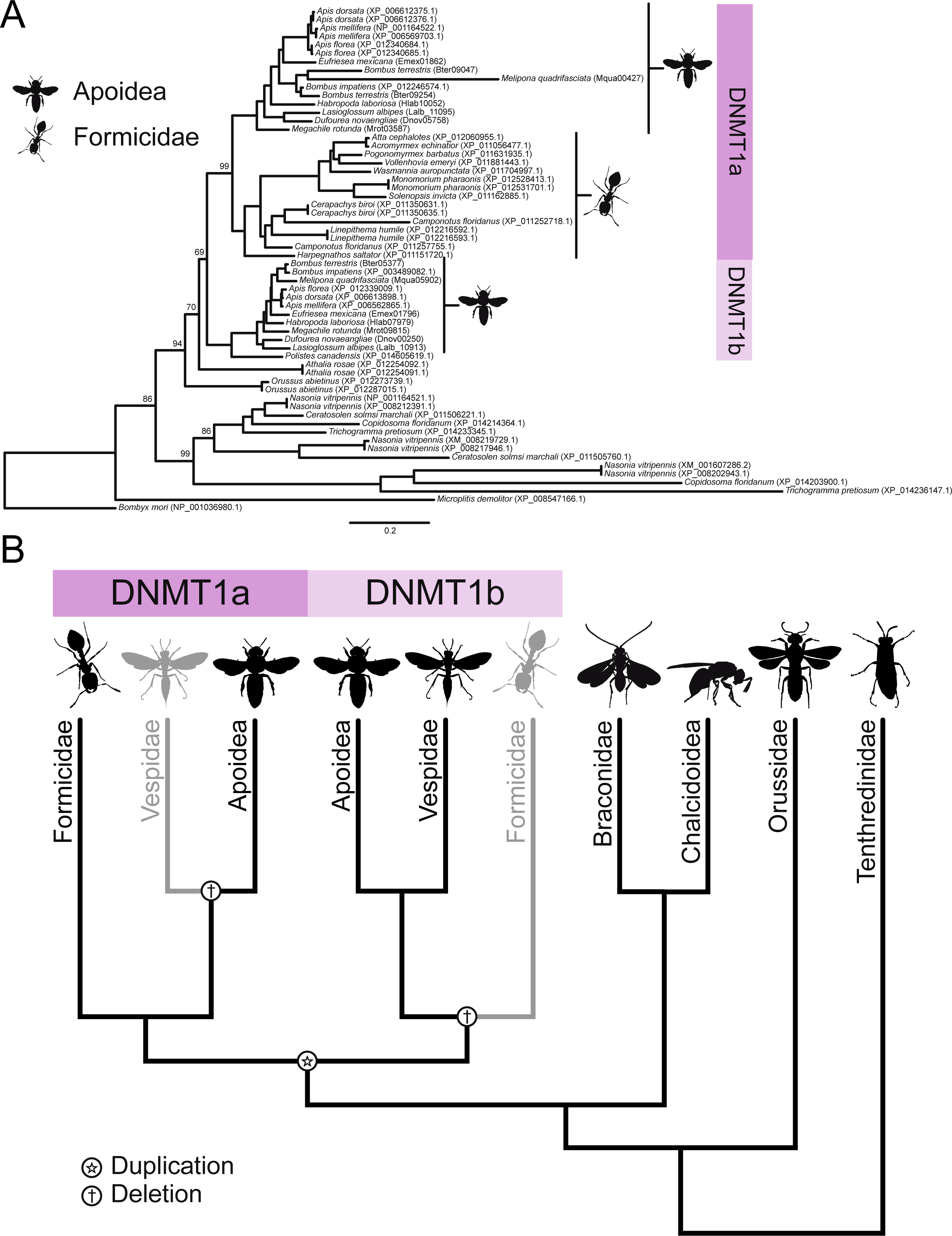

